# A cholesterol addiction creates an Achilles’ heel in *APC*-truncated colorectal cancer

**DOI:** 10.64898/2026.11.24.768807

**Authors:** Issam Boukich, Rubén A. Bartolomé, María Cuerda-López, Laura Pintado, Tania Calvo, María Estarás, Paraskevi Kousteridou, Pierre Bertrand, Pauline Perné, Delphine Debayle, Fabienne Marchai, Juan I. Imbaud, J. Ignacio Casal

**Affiliations:** Department of Biomedicine. Centro de Investigaciones Biológicas Margarita Salas. 28040 Madrid. Spain; Protein Alternatives SL. Tres Cantos Madrid. Spain; Centre de Recherche en Cancérologie de Marseille. INSERM 106. Marseille. France; CNRS-UMR 7275. France

**Keywords:** NPC2, cholesterol, auxotrophy, APC, metastasis, colorectal cancer

## Abstract

Cholesterol auxotrophy, dependence on exogenous cholesterol, is an emerging metabolic vulnerability of cancer that remains poorly understood. Here, we show that *APC*- truncated CRC cell lines displayed cholesterol auxotrophy, whereas *CTNNB1*-mutant cells were cholesterol-independent. The endolysosomal Niemann-Pick type C2 (NPC2) protein is a key transporter of cholesterol from lysosomes to cellular membranes. *NPC2* knockdown (KD) in auxotrophic *APC*-mutant cells depleted plasma-membrane cholesterol, disorganized lipid-raft scaffolding and signaling, and inhibited stemness, xenograft growth and metastasis. In contrast, NPC2 KD *CTNNB1*-mutant cells retained tumorigenic and metastatic capacity. Genome-wide transcriptional responses to NPC2 loss were nearly uncorrelated between *APC* and *CTNNB1* mutant cell lines revealing genotype-dependent adaptation and opposite cell signaling pathways alterations. NPC2 KD auxotrophs showed abortive compensatory cholesterol synthesis with no alternative egress route, whereas *CTNNB1*-mutant cells restored lysosomal cholesterol egress by inducing NPC1 and sustaining Wnt/stemness identity, AP-1/YAP/TAZ signaling and a pro-invasive secretome. Lipidomics confirmed the signaling divergences: sphingomyelin collapse, membrane-fluidity remodeling exclusive to the auxotrophs, and opposite- direction diacylglycerol signaling in CTNNB1 mutants. Notably, *APC*-truncated cells phenocopied selected Niemann–Pick type C hallmarks, linking lysosomal cholesterol export to tumor fitness. Thus, NPC2 constitutes a synthetic-lethal vulnerability specific to cholesterol-auxotrophic *APC*-mutant CRCs, providing a mechanistic rationale for targeting the secreted and recaptured NPC2 pool in this molecular subset of patients.

## INTRODUCTION

Cholesterol “addiction” has been linked to enhanced tumor cell growth, invasiveness, and therapeutic resistance ^1^. Metastatic cells, in particular, require substantial amounts of cholesterol to support aberrant proliferation, survival, and invasion. High proliferative rates combined with limited access to plasma cholesterol, as a consequence of the hypoxic and nutrient-restricted tumor microenvironment, force cancer cells to mobilize all available cholesterol sources, including cholesterol biosynthesis and uptake ^2^. Intracellular cholesterol levels are tightly regulated through three different pathways: cholesterol uptake from the extracellular environment (via low-density lipoprotein receptor (LDLR)), *de novo* cholesterol synthesis (via SREBPs) and cholesterol efflux (via ABCA1) ^3^. Disruptions in any of these processes will alter the delicate balance of cholesterol homeostasis compromising cell survival. Cholesterol biosynthetic pathway also supplies precursors for the synthesis of signaling sterols, including vitamin D, bile acids, corticosteroids, and sex hormones ^4^. Moreover, free cholesterol plays a critical role in the formation and function of lipid rafts, which serve as key platforms to amplify signaling cascades in carcinogenesis ^5,6^. For all these reasons, cholesterol homeostasis is getting increased attention as a metabolic vulnerability in cancer ^7^.

Cholesterol auxotrophy, defined as a dependence on exogenous cholesterol uptake, has emerged as a recurrent and potentially targetable metabolic vulnerability across multiple cancer types. In human cancers, glioblastoma cells cannot sustain <u>de novo</u> cholesterol synthesis and depend on LDLR–mediated uptake of lipoprotein-derived cholesterol for survival ^8^. Similarly, a subset of ALK+ anaplastic large cell lymphoma cell lines becomes cholesterol auxotrophic through loss of SQLE expression, creating a dependency on LDLR-mediated uptake ^9^. In clear cell renal cell carcinoma, biallelic VHL loss and constitutive HIF stabilization repress cholesterol-biosynthetic genes to rely on SCARB1-mediated HDL import, rendering SCARB1 inhibition selectively lethal ^7,10,11^. Extracellular LDL-derived cholesterol also contributes to the growth of advanced and castration-resistant prostate cancer ^12^. Additional mechanisms can generate or modulate this dependency. Loss of wild-type p53 relieves repression of the mevalonate pathway by reducing ABCA1 expression and permitting SREBP2 maturation in hepatocellular and breast cancers ^13,14^, whereas oncogenic PI3K/AKT/mTOR or AKT1 signaling can increase sterol demand beyond synthetic capacity, as reported in rhabdomyosarcoma and lung cancer ^15,16^. Likewise, USP28-mediated stabilization of SREBP2 promotes cholesterol-pathway activation in squamous carcinomas ^17^. Collectively, these findings indicate that cholesterol auxotrophy is a convergent metabolic phenotype arising through distinct genetic and signaling mechanisms, with dependence on LDL- or HDL-receptor- mediated uptake as a common feature. This metabolic vulnerability has not yet been systematically explored in colorectal cancer (CRC).

Recent proteomic analysis of the secretome of highly metastatic cells identified a molecular signature associated with poor prognosis in CRC ^18^, which included the Niemann-Pick type C2 (NPC2) protein. NPC2 was originally identified as HE1, a major secretory component of epididymal fluid ^19^, which carries M6P modifications on its two N-linked glycans (Asn58, Asn135) that mediate receptor-dependent uptake and lysosomal delivery ^20^. NPC2 is a soluble lysosomal cholesterol-binding protein that cooperates with the transmembrane transporter NPC1 to egress cholesterol from late endosomes and lysosomes to the cytosol and other cellular compartments ^21,22^, including the plasma membrane (PM) ^5^. When NPC2 or NPC1 are absent or mutated, patients suffer the lethal Niemann-Pick disease, characterized by toxic cholesterol accumulation in the late endosomes/lysosomes ^19^. NPC1/NPC2 loss additionally causes secondary storage of glycosphingolipids and sphingomyelin, mitochondrial dysfunction with reduced coenzyme Q, oxidative stress, and dysregulated mTORC1/autophagy signaling ^23^. Although NPC1 has been the subject of various studies in cancer research ^24^, NPC2 relevance in cancer, particularly in CRC, where cholesterol homeostasis intersects with Wnt signaling, remains poorly understood.

CRC progression usually initiates following the appearance of *APC* mutations, which are present in ⁓90% of the sporadic CRCs ^25^. *APC* is a key regulator of Wnt signaling ^26^. However, a small percentage of CRC patients (5-10%), generally MSI- associated, present mutations in the *CTNNB1* gene while maintaining wild-type *APC* ^27^. Interestingly, *APC* and *CTNNB1* mutations are mutually exclusive ^28^. Despite being a small percentage in patients, *CTNNB1* mutants tend to be overrepresented among metastatic CRC cell lines, being HCT116 and SW48 two examples of *CTNNB1*-mutated cell lines. Two recent studies have established a critical link between *APC* truncation and PM cholesterol activity in CRC. Erazo-Oliveras et al. ^3^ demonstrated that oncogenic APC increases PM free cholesterol and membrane rigidity, reshaping Wnt receptor (LRP6, Fzd7) nanodomains and enhancing their interactions with cholesterol and the cytosolic effector Dishevelled (Dvl). Sharma et al. ^29^ extended this finding by showing that inner-leaflet PM cholesterol is elevated in *APC*-truncated CRC cells, drives Wnt-independent signalosome formation through Dvl-cholesterol interaction, and that cholesterol-targeting inhibitors can selectively suppress *APC*-truncated CRC cells without intestinal toxicity.

Here, we observed that the *APC/CTNNB1* mutational context determines cholesterol auxotrophy and the response to NPC2 loss in CRC. Therefore, we investigated the effects of NPC2 depletion in cholesterol trafficking, signaling, clonogenicity, and metastatic capacity, integrating PM cholesterol quantification, transcriptomics, lipidomics, and *in vivo* models across *APC*-truncated and *CTNNB1*-mutant CRC cells. Defining these context-dependent mechanisms is a prerequisite for precision targeting of cholesterol metabolism via NPC2 in metastatic CRC.

## RESULTS

### *APC-*mutant CRC cell lines show cholesterol auxotrophy

First, we investigated cholesterol auxotrophy using five CRC cell lines: KM12SM, HT29, SW620, SW48 and HCT116, which represent different *APC/CTNNB1* mutational landscapes ^30^ (**Table 1**). APC truncations in KM12SM, HT29 and SW620 were confirmed by western blot (**Supp. Fig. S1A**). In contrast, SW48 and HCT116 exhibited full length APC, albeit the lower protein expression of APC and β-catenin in HCT116 cells **(Supp. Fig. S1A)**. Cholesterol auxotrophy was explored in the five metastatic cell lines using different culture conditions containing either 10% FBS, 10% delipidated serum (DLPS) or DLPS reconstituted with 10 µg/ml cholesterol. Remarkably, KM12SM, HT29 and SW620, harboring truncated *APC*, exhibited a significant proliferation arrest in DLPS, with significant effects on cell confluence visible by phase contrast microscopy (**Fig. 1A**). We also noticed a major presence of apoptosis in *APC* mutant cells cultured with DLPS (**Fig. 1B**). These effects on cell viability, proliferation and apoptosis were largely rescued following treatment with exogenous cholesterol, supporting a vulnerability of *APC*-truncated CRC cell lines to exogenous cholesterol. In contrast, *CTNNB1* mutants, HCT116 and SW48, did not show significant alterations in proliferation or apoptosis alterations following lipid removal or cholesterol treatment (**Fig. 1A, 1B**). Collectively, these results indicate that the presence of oncogenic, truncated *APC* leads to cholesterol auxotrophy in CRC. In contrast, *CTNNB1-*mutated cell lines did not rely on exogenous cholesterol for cell proliferation and survival.

**Figure 1.**
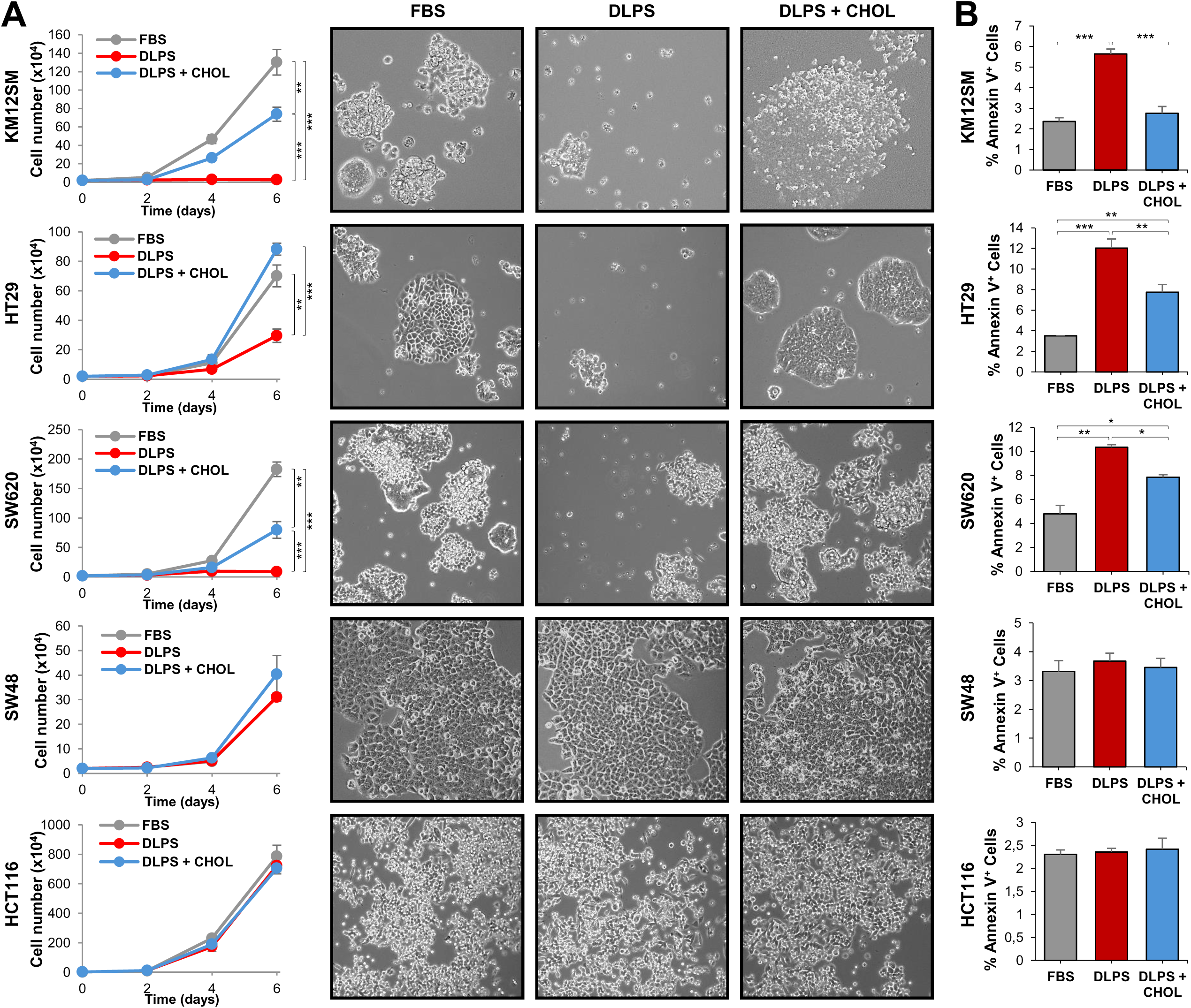
APC-mutant colorectal cancer cells display cholesterol auxotrophy. (A) Cholesterol dependency was assessed by culturing the indicated CRC cell lines in medium supplemented with fetal bovine serum (FBS) or delipidated serum (DLPS), with or without cholesterol (CHOL) supplementation. Cell growth was significantly reduced in DLPS medium and partially restored by CHOL addition (**, p < 0.01; **, p < 0.001). Representative images acquired after 6 days of culture are shown. (B) Apoptosis was analyzed under the same experimental conditions. The percentage of apoptotic cells was significantly increased in DLPS medium, whereas CHOL supplementation significantly reduced this effect. Statistical significance from three different experiments is indicated as (*, p < 0.05; **, p < 0.01; ***, p < 0.001).

**Table 1.**
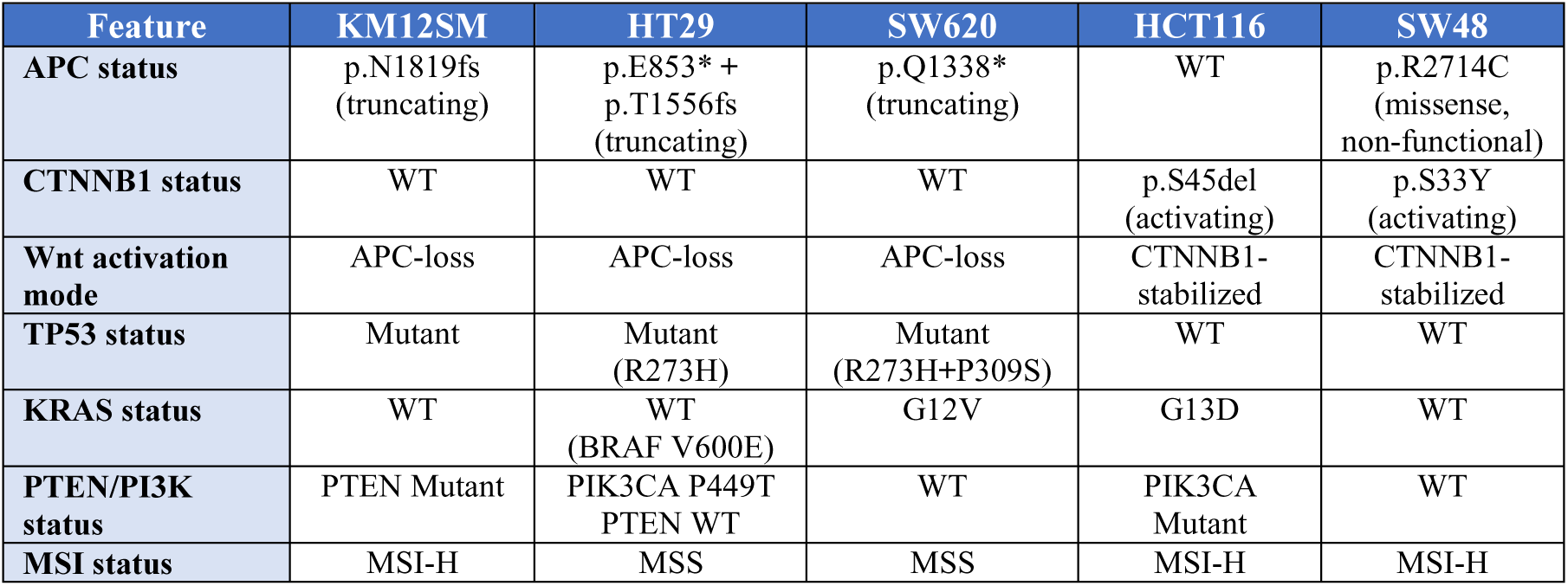
Genetic characteristics of CRC metastatic cell lines.

| **Feature** | **KM12SM** | **HT29** | **SW620** | **HCT116** | **SW48** |
|---|---|---|---|---|---|
| **APC status** | p.N1819fs (truncating) | p.E853* + p.T1556fs (truncating) | p.Q1338* (truncating) | WT | p.R2714C (missense, non-functional) |
| **CTNNB1 status** | WT | WT | WT | p.S45del (activating) | p.S33Y (activating) |
| **Wnt activation mode** | APC-loss | APC-loss | APC-loss | CTNNB1-stabilized | CTNNB1-stabilized |
| **TP53 status** | Mutant | Mutant<br>(R273H) | Mutant (R273H+P309S) | WT | WT |
| **KRAS status** | WT | WT<br>(BRAF V600E) | G12V | G13D | WT |
| **PTEN/PI3K status** | PTEN Mutant | PIK3CA P449T PTEN WT | WT | PIK3CA Mutant | WT |
| **MSI status** | MSI-H | MSS | MSS | MSI-H | MSI-H |

### Loss of NPC2 depletes plasma membrane cholesterol and promotes lysosomal cholesterol accumulation in an *APC*-dependent manner

Since auxotrophic cells rely on exogenous cholesterol delivered through the endolysosomal pathway, we examined whether NPC2 depletion alters cholesterol distribution between lysosomes and the PM. NPC2 expression in cell lysates showed higher expression of NPC2 in aggressive metastatic KM12SM and HCT116 cell lines (**Supp. Fig. S1B)**, harboring *APC* and *CTNNB1* mutations, respectively. Then, cells were stably depleted of NPC2, and KD efficiency was validated by qPCR (**Supp. Fig. S1C**) and western blotting (**Supp. Fig. S1D**). NPC2-KD and SCR control cells were subsequently analyzed by filipin staining to assess cholesterol distribution, using methyl- β-cyclodextrin (MβCD) as a positive control for PM cholesterol depletion (**Fig. 2A**). Filipin analysis showed reduced PM-associated cholesterol together with increased lysosomal cholesterol accumulation in both NPC2-KD cell lines, although to a different extent. KM12SM KD showed a 43% reduction in PM, while HCT116 showed a 18% reduction, which suggests different cholesterol availability for lipid raft-mediated signaling, associated with a concomitant accumulation of intracellular cholesterol (**Fig. 2B**). Next, we examined the lysosomal location of NPC2 by co-staining with LAMP1, a lysosomal marker. In control SCR cells, NPC2 staining revealed a diffuse intracellular distribution pattern, with partial LAMP1 overlap, while strong cytoplasmic staining was observed in HCT116, with little or no NPC2 colocalization with LAMP1 (**Fig. 2C**). Following NPC2 depletion, both cell lines exhibited prominent lysosomal accumulation and polarized lysosomal clustering (LAMP1-positive) (**Fig. 2D**). The retention of cholesterol in the lysosomes mirrors the lysosomal trafficking defects characteristic of Niemann–Pick disease^31^. Following these results, we investigated cholesterol auxotrophy in NPC2 KD cells. Whereas KM12SM NPC2 KD cells showed similar auxotrophy to the parental cells including increase of apoptosis in DLPS medium (**Fig. 2E, F**), HCT116 KD cells showed a modest reduction in proliferation in DLPS-containing medium and no alterations in apoptosis ratios, confirming significant cholesterol independence (**Fig. 2E, F**). Collectively, these findings indicate that NPC2 depletion redistributes cholesterol from the PM toward lysosomal compartments in both cell lines, with a more pronounced effect in KM12SM cells. NPC2 depletion does not alter the cholesterol auxotrophy status of both models.

**Figure 2.**
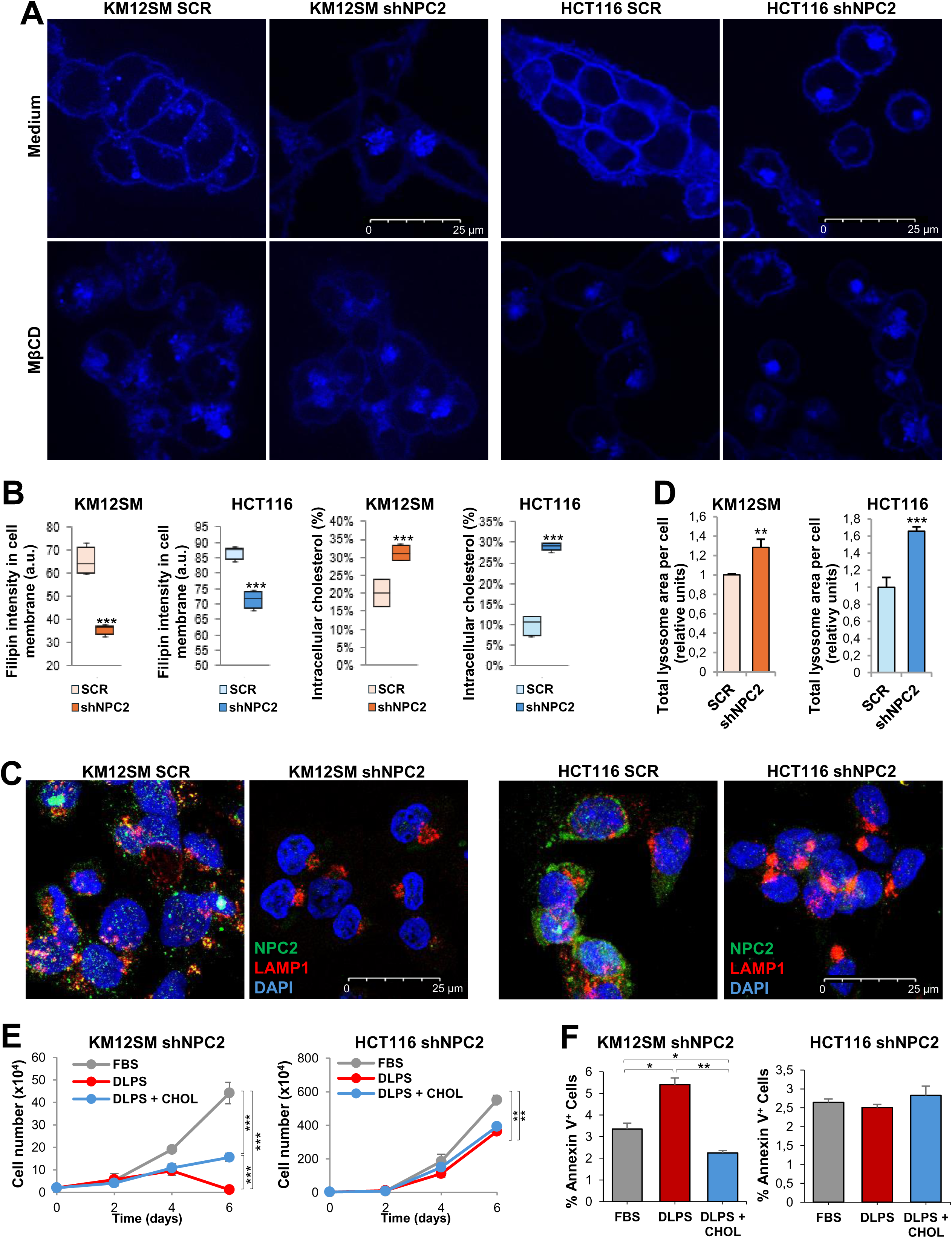
NPC2 depletion redistributes cholesterol from the plasma membrane to lysosomal compartments. (A) Representative filipin-stained images of the indicated CRC cell lines following NPC2 silencing, with or without MβCD treatment as a control for PM cholesterol depletion. (B) Quantification of PM filipin fluorescence, the PM-to-intracellular cholesterol ratio, and the fraction of intracellular cholesterol relative to total cellular cholesterol. NPC2 silencing significantly increased the intracellular cholesterol fraction in both cell lines (***p < 0.001). (C, D) Representative immunofluorescence images of the indicated cell lines stained with DAPI and antibodies against NPC2 and LAMP1. NPC2 depletion increased the LAMP1-positive lysosomal area (**, p < 0.01; ***, p < 0.001). (D) Total lysosomal area per cell quantification. (E) Auxotrophy assays in NPC2-depleted KM12SM and HCT116 cells as described in Fig.1 (**, p < 0.01; ***, p < 0.001). (F) apoptosis assays in the same conditions as in E (*, p < 0.05; **, p < 0.01). Statistical significance from three different experiments is indicated

### Loss of NPC2 differentially reduces metastatic competence, clonogenicity and spheroid formation depending on the APC mutational background

To investigate the functional impact of NPC2 depletion on the metastatic competence in cell culture, KM12SM and HCT116 cells were silenced with either shRNA for long-term experiments or siRNA for transient silencing. KD cells showed significantly reduced adhesion, migration and invasion in both cases (**Fig. 3A, Supp. Fig. S2**), whereas cell proliferation was inhibited only in KM12SM cells. Consistent with this differential proliferative phenotype, NPC2 depletion completely suppressed KM12SM tumor xenograft growth following subcutaneous implantation in NSG mice, while HCT116 KD tumors showed an initial growth delay, followed by recovery (**Fig. 3B**). Strikingly, NPC2 loss also produced opposite effects on clonogenicity, virtually abolishing colony formation in KM12SM cells, while significantly increasing colony formation in HCT116 (**Fig. 3C**). In 3D cultures, NPC2 depletion disrupted spheroid organization in both models, generating looser and more irregular aggregates with reduced Wadell circularity, consistent with impaired cell–cell cohesion and a less epithelial phenotype (**Fig. 3D**). Collectively, these findings indicate that NPC2 supports metastatic traits in both cell lines, but is differentially required for tumor growth and clonogenic fitness, with KM12SM cells displaying a markedly greater dependency. NPC2 also contributes to the maintenance of cohesive epithelial organization in 3D.

**Figure 3.**
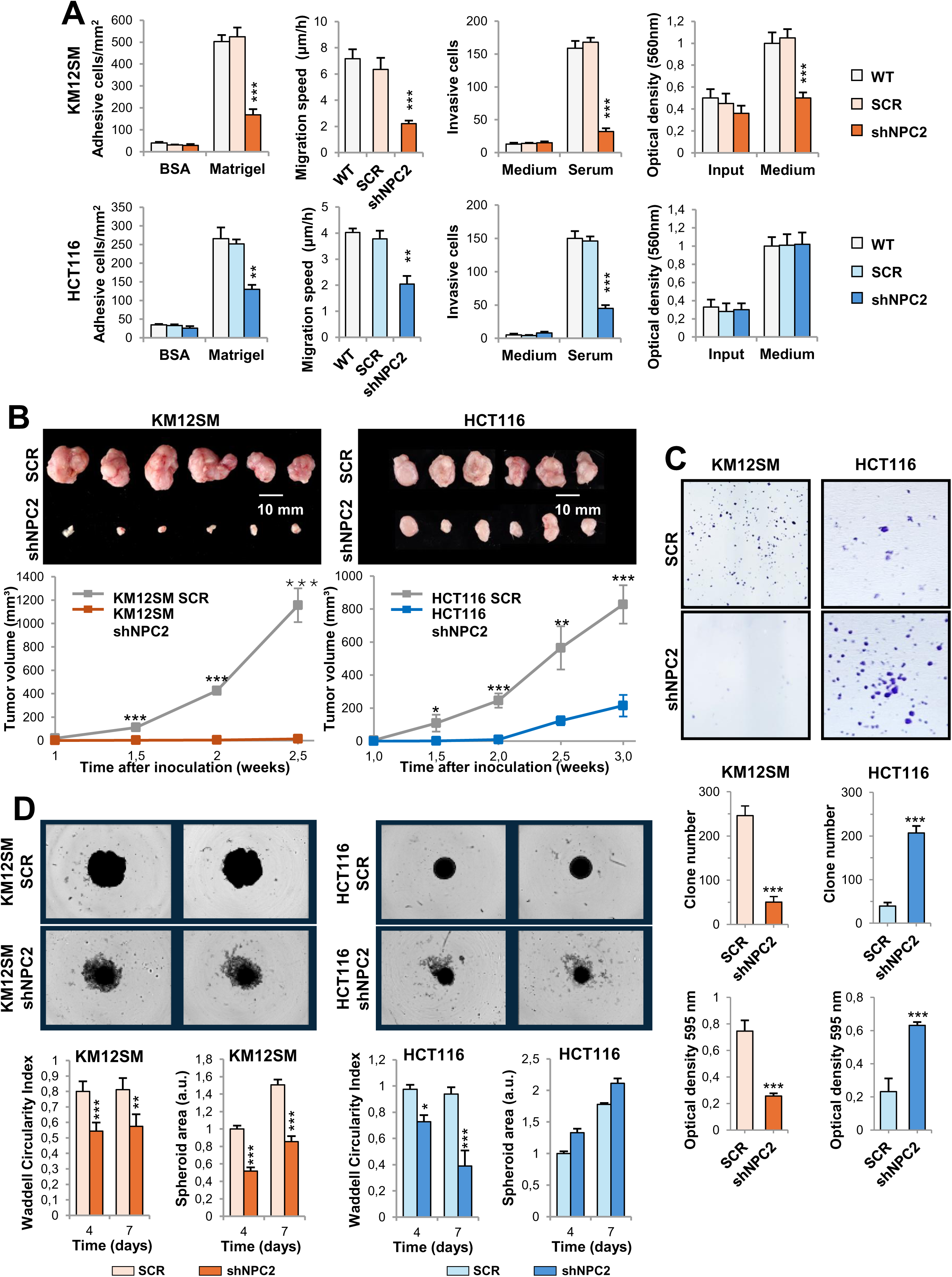
NPC2 loss regulates tumor growth, metastatic traits, and clonogenic fitness in colorectal cancer cells. (A) The indicated cell lines were analyzed for adhesion, migration by wound-healing assay, invasion, and proliferation. NPC2 silencing significantly reduced adhesion, migration, invasion and/or proliferation, depending on the cellular model (**p < 0.01; ***p < 0.001). (B) NPC2-KD and control cells were subcutaneously inoculated into NSG mice, and tumor volume was monitored twice weekly. NPC2 depletion significantly impaired tumor growth, with a markedly stronger effect in KM12SM than in HCT116 cells (*, p < 0.05; **p < 0.01; ***p < 0.001). Representative isolated tumors are shown. (C) Colony formation assays revealed that NPC2 depletion markedly reduced colony number and cell density in KM12SM cells, whereas both parameters increased in HCT116 cells (***p < 0.001). (D) The indicated cell lines were grown as 3D spheroids. Representative images and quantification of spheroid area and Waddell circularity are shown. NPC2 depletion significantly altered spheroid growth and morphology, reducing circularity and spheroid area (*, p < 0.05; **p < 0.01; ***p < 0.001).

### NPC2 depletion promotes *APC*-dependent cell signaling divergences

To assess the signaling consequences of NPC2 depletion, we compared control SCR and NPC2-KD cells after serum, IL-13, or EGF stimulation, which engage complementary growth factor- and cytokine-dependent signaling cascades ^32,33^. Following treatment with IL-13, EGF and/or serum, KM12SM KD cells showed lower phosphorylation of FAK(Y397), Src(Y416), AKT(S473) and ERK (Thr202/Tyr204) (**Fig. 4A**). Interestingly, HCT116 KD cells did not show significant signaling alterations respect to SCR cells after the various treatments. To confirm that the signaling effects were not shRNA-dependent, we repeated the analysis with an independent siRNA targeting NPC2. Consistently, the divergent signaling effects were largely reproduced (**Supp. Fig. S3**).

**Figure 4.**
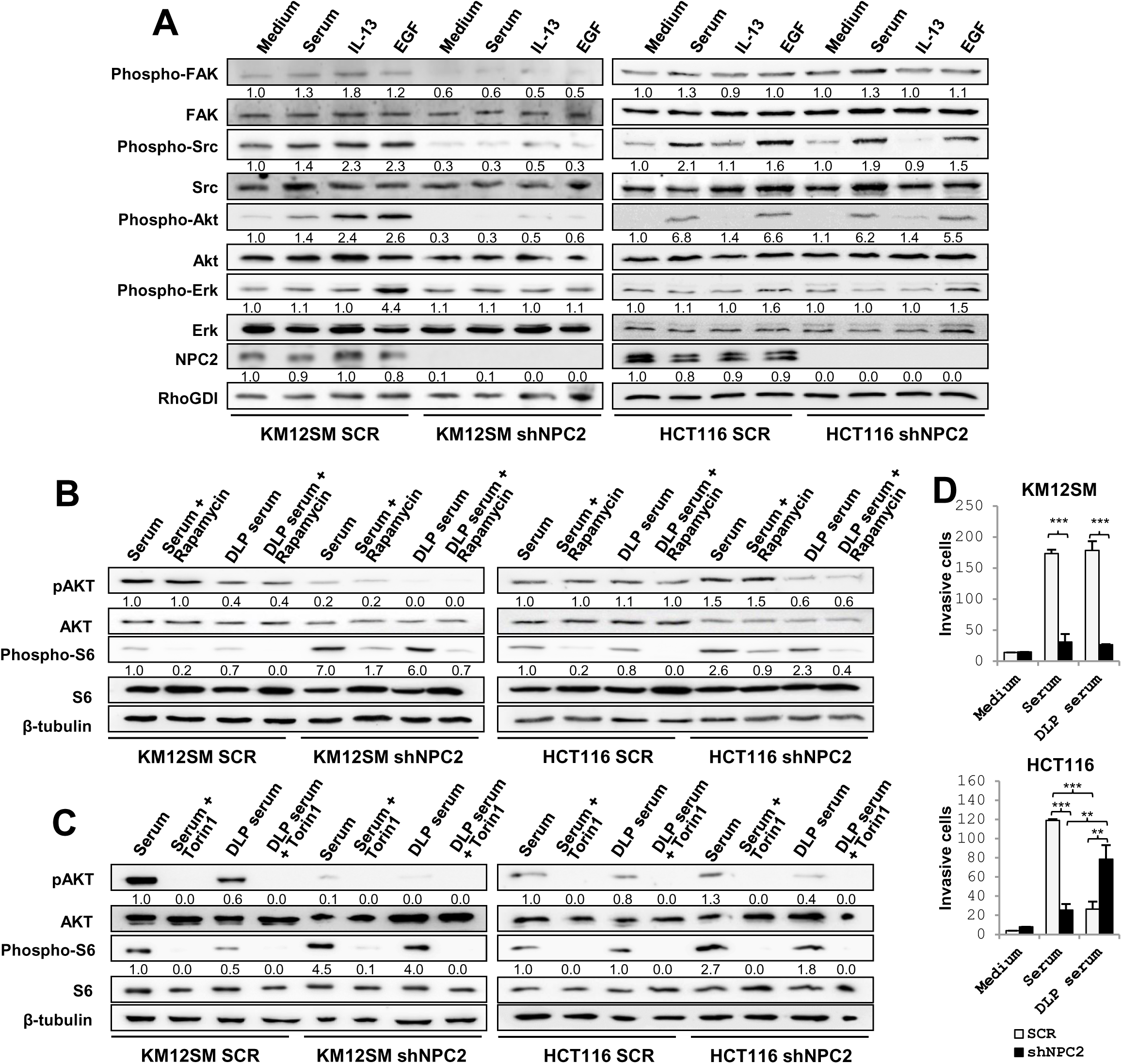
NPC2 depletion differentially modulates growth-factor and mTOR signaling according to cholesterol availability. (A) SCR and NPC2-KD cells were stimulated with serum, IL-13, or EGF, and cell lysates were analyzed by western blotting for the indicated signaling proteins. Band quantification is shown below the corresponding lanes. (B) The indicated cell lines were cultured in serum-containing or DLPS medium in the presence or absence of rapamycin (1 μM), followed by western blot analysis of the indicated proteins. (C) Cells were analyzed under the same conditions following treatment with Torin-1. (D) Cell invasion was assessed under serum or DLPS conditions (**p < 0.01; ***p < 0.001). Data are representative from three different experiments

In addition, given the central role of mTOR signaling in metabolic regulation, we examined the AKT–mTORC1–S6 axis following rapamycin (**Fig. 4B**) and Torin-1 (**Fig. 4C**) treatment. NPC2 depletion induced a pronounced uncoupling of AKT and S6 phosphorylation in KM12SM cells; pAKT was strongly reduced under both serum and DLPS conditions, whereas pS6 was strongly increased under both serum and DLPS (**Fig. 4B, C**). The pS6 elevation was largely abolished by Torin-1 (**Fig. 4C**). Since pAKT (Ser473) was already collapsed in NPC2-depleted KM12SM cells, and rapamycin did not further reduce pAKT in any condition whereas Torin-1 eliminated it, the sustained S6 phosphorylation occurs without AKT/mTORC2 input, identifying mTORC1/S6K as the main driver. Notably, elevated pS6 persisted under DLPS, indicating that mTORC1 activation is maintained independently of extracellular lipoprotein availability, consistent with lysosomal cholesterol accumulation driving Rag-dependent mTORC1 activation at the lysosomal surface. By contrast, after NPC2 depletion, HCT116 cells increased AKT phosphorylation under serum, and only partially reduced under DLPS, exhibiting a modest pS6 increase under serum and DLPS that remained fully sensitive to both rapamycin and Torin-1 (**Fig. 4B, C**).

Strikingly, cell invasion assays in serum or DLPS revealed marked differences between both models (**Fig. 4D**). NPC2 depletion completely inhibited invasion in KM12SM cells under both serum and DLPS conditions. In contrast, NPC2 silencing in HCT116 cells markedly reduced invasion under serum conditions, but enhanced invasion under lipoprotein-depleted conditions revealing a context-dependent response.

### Loss of NPC2 inhibits tumor homing and liver metastasis in APC-truncated CRCs

To determine whether the *in cellulo* effects of NPC2 depletion translated into altered metastatic competence *in vivo*, we assessed liver homing in Swiss nude mice and metastatic colonization following intrasplenic inoculation into NSG mice. NPC2 silencing significantly impaired liver homing by KM12SM cells, whereas no significant effect was observed in HCT116 cells (**Fig. 5A**). Consistently, mice inoculated with KM12SM NPC2-KD cells showed markedly prolonged survival and developed few metastatic nodules compared with SCR controls (**Fig. 5B, C**). By contrast, NPC2 depletion in HCT116 cells did not improve survival and was associated with extensive liver colonization and pronounced hepatomegaly (**Fig. 5B, C**), mirroring the hepatic alterations characteristic of Niemann–Pick disease ^34^. To validate the APC-truncated phenotype, we used SW620 cells, an additional *APC*-mutant model (**Fig. 5D**). Whereas 4 of 6 mice inoculated with SW620 SCR cells developed liver metastases, no detectable metastatic nodules or increase in liver weight was observed in mice receiving SW620 NPC2-KD cells (**Fig. 5D**). These findings, together with our previous observation regarding cholesterol auxotrophy, support a genotype-dependent requirement for NPC2 in metastatic progression. NPC2 appears essential for maintaining metastatic competence in cholesterol-dependent *APC*-mutant, but largely dispensable in *CTNNB1*-mutant tumors.

**Figure 5.**
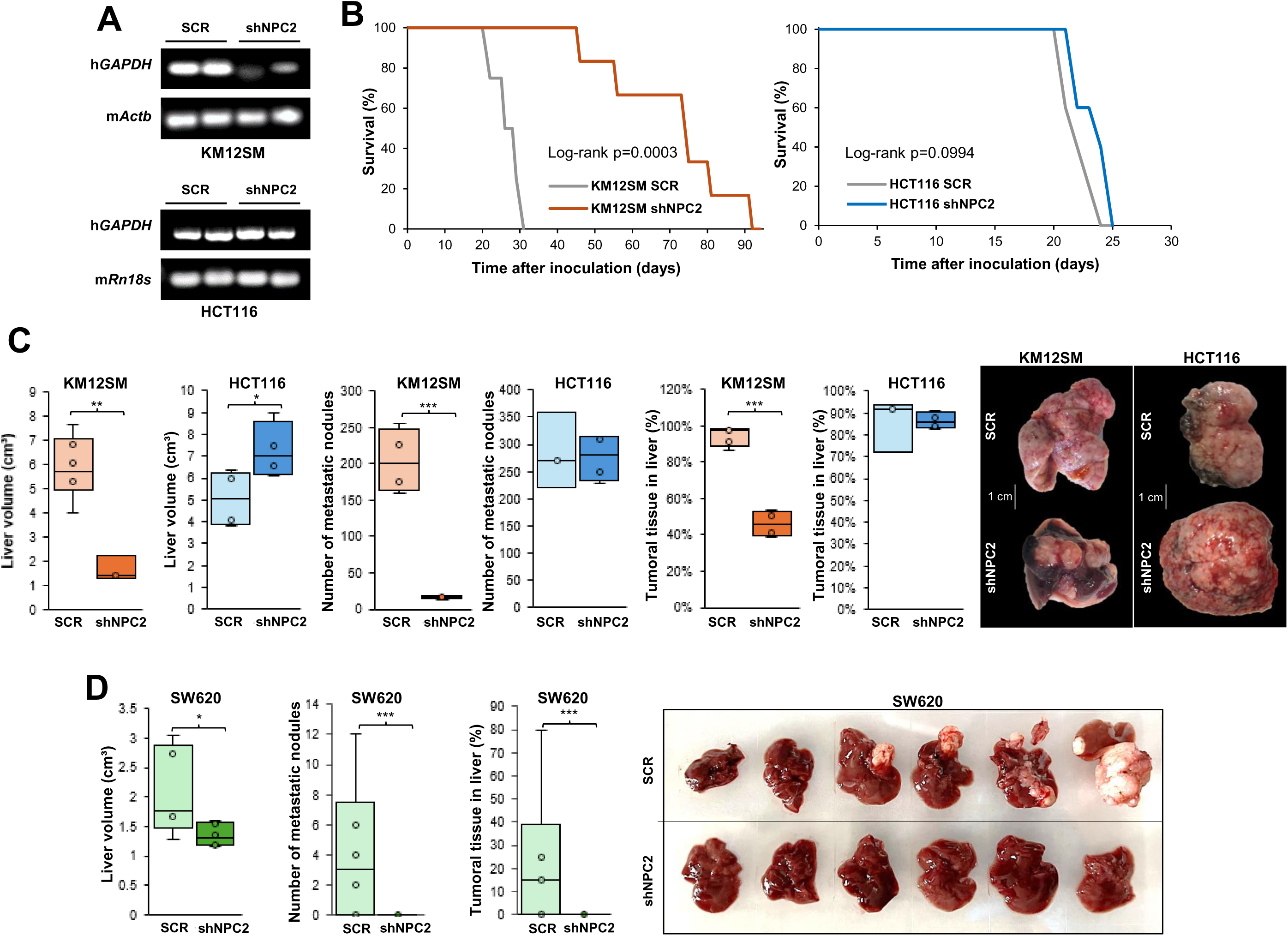
NPC2 depletion reduces liver metastasis and prolongs survival preferentially in APC-mutant colorectal cancer models. (A) KM12SM and HCT116 SCR and NPC2-KD cells were inoculated intrasplenically into Swiss nude mice. After 48 h, RNA was isolated from the livers and analyzed by RT-PCR for human *GAPDH* transcripts as a measure of liver homing. Mouse *Actb* or *Rn18S* transcripts were used as loading controls. (B) Kaplan–Meier survival analysis following intrasplenic inoculation of KM12SM and HCT116 SCR and NPC2-KD cells into NSG mice (n=6 per experiment). NPC2 depletion markedly prolonged survival in mice inoculated with KM12SM cells, whereas no survival benefit was observed with HCT116 cells. Representative images of metastatic livers are shown. (C) Analyses of liver volume, number of metastatic nodules, and percentage of tumor tissue in mouse livers from B. (D) Stable SCR and shNPC2 SW620 cells were inoculated intrasplenically into NSG mice (n=6 per experiment). NPC2 depletion markedly impaired metastatic progression, with prolonged survival and a significant reduction in metastatic nodules, tumor-occupied liver area, and liver weight compared with SCR controls (**p < 0.01; ***p < 0.001).

### NPC2 depletion exposes genotype-dependent cholesterol adaptation and survival programs

To clarify the molecular basis of the divergent effects of NPC2 depletion on metastatic competence, we performed global transcriptomic analyses in both models. Genome-wide comparison of NPC2-KD versus control SCR cells revealed a very low cross-line correlation between both models (r = 0.06) (**Fig. 6A**), indicating that the two models engage largely independent responses to NPC2 loss. KM12SM underwent more extensive reprogramming than HCT116 (1,228 vs 926 upregulated and 1,197 vs 652 downregulated genes (**Supp. Tables S1 and S2**). Gene set enrichment analysis (GSEA) identified gene alterations shared by both lines: enrichment of epithelial–mesenchymal transition (EMT), interferon-α/γ responses, TGFβ signaling, apoptosis and hypoxia, together with downregulation of bile-acid metabolism (**Fig. 6B, Supp. Fig. S4**). However, both cell lines diverged in cholesterol biosynthesis, IL6/JAK/STAT3, IL2/STA5, KRAS, Hedgehog signaling and in proliferation programs (mitotic spindle, G2/M checkpoint), which were suppressed in KM12SM and maintained or enriched in HCT116. KEGG and Reactome analyses confirmed this architecture (**Supp. Fig. S5A**), both lines showed enriched endocytosis, cytoskeletal organization, tight junctions and focal adhesions. However, KM12SM exhibited broad inhibition of PI3K/AKT, MAPK/RAS, RHO- GTPases, RTK and metabolic signaling with activation of DNA-damage, senescence, apoptosis and autophagy programs, whereas HCT116 showed enrichment of RHO- GTPases, PI3K/AKT, MAPK, mTORC1, inflammatory, NOTCH and focal-adhesion pathways, which correlate with an adaptive, pro-invasive and metabolically plastic phenotype. Furthermore, the divergence extended to the cholesterol homeostasis genes that connect NPC2 loss to auxotrophy (**Fig. 6C**). HCT116 KD cells engaged an effective compensatory response, showing increased LDLR and NPC1 induction, consistent with preserved uptake and restored lysosomal egress, whereas in KM12SM, LDLR and NPC1 were nearly abolished and ABCA1 decreased (**Fig. 6C, D**). Additionally, whereas KM12SM displayed a SREBP2-associated biosynthetic program (ACAT2, SQLE, IDI1), HCT116 suppressed downstream sterol-conversion enzymes (FDFT1, LSS, CYP51A1). QPCR (**Supp. Fig S5B**) and western blotting **(Fig. 6D**) confirmed most of these changes, revealing substantial post-transcriptional regulation. HCT116 thus adapted to NPC2 loss by increasing or preserving cholesterol acquisition and egress genes, whereas cholesterol- auxotrophic KM12SM fails to restore cholesterol trafficking, supporting the mechanistic basis of its synthetic lethality upon NPC2 depletion (**Supp. Fig. S6)**.

**Fig. 6.**
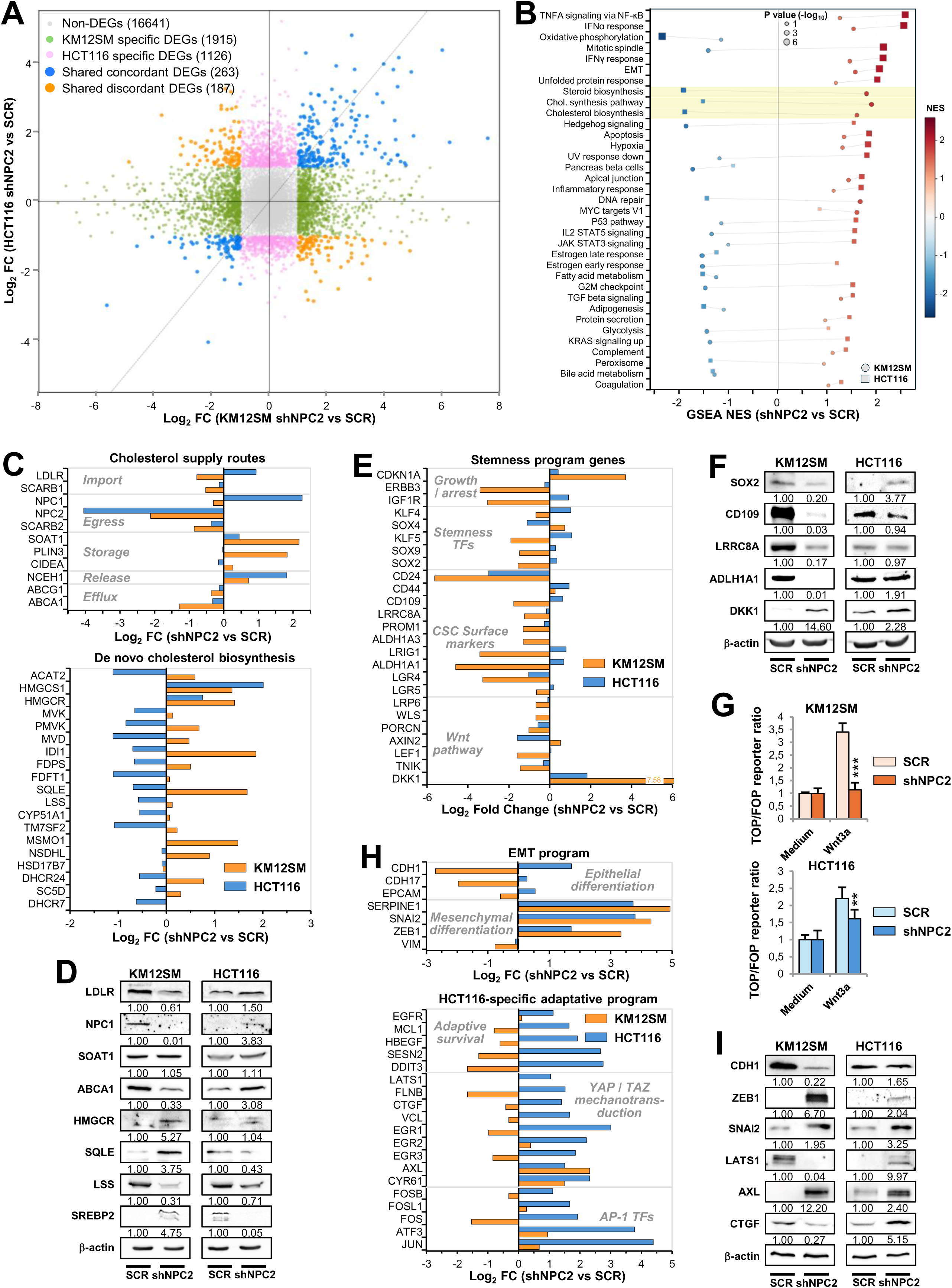
NPC2 silencing drives divergent cholesterol, stemness and adaptive transcriptional programs in colorectal cancer cells. (A) Scatter plot comparing genome-wide log_2_ FC of differentially expressed genes in NPC2-KD versus SCR cells between KM12SM and HCT116 (n = 3 biological replicates per condition). (B) GSEA of transcriptome analyses from KM12SM (circles) and HCT116 (squares) cells expressing NPC2 shRNA (shNPC2) relative to scrambled shRNA controls (SCR). Normalized enrichment scores (NES) are indicated by color (red, enriched; blue, depleted in shNPC2). Dot size represents the statistical significance (−log_10_ adjusted p value). Cholesterol-associated gene sets are highlighted in yellow. Only gene sets significant (adjusted p < 0.05) in at least one cell line are shown. (C) Log_2_ FC (shNPC2 *vs* SCR) of transcripts involved in cholesterol trafficking (top), grouped by function: import, egress, storage, release and efflux and *de novo* cholesterol-biosynthesis (bottom). (D) Western blot validation of cholesterol-related proteins; β-actin was used as loading control. (E) Log_2_ FC of genes associated with stemness program, CSC markers and Wnt pathway. (F) Western blot analysis of stemness-related proteins; β-actin as loading control. (G) TOP/FOP luciferase reporter ratio in SCR and shNPC2 cells under basal conditions or after stimulation with Wnt3a-conditioned medium (0.5 μg/mL). Data are mean ± s.e.m. of three independent experiments; **, p< 0.01, ***, p< 0.001. (H) Log_2_ FC of EMT- associated genes (top), grouped into epithelial and mesenchymal programs, and adaptive program genes (adaptive survival, YAP/TAZ mechanotransduction and AP-1 transcription factor groups). (I) Western blot validation of EMT and adaptive-response proteins; β-actin as loading control.

Additionally, NPC2 depletion disrupted the canonical Wnt/intestinal stem-cell (ISC) program in KM12SM, but largely preserved it in HCT116 (**Fig. 6E**). KM12SM KD downregulated multiple stemness genes (LGR4, LEF1, SOX2, SOX9, PROM1/CD133, LGR5), with SOX2 and ALDH1A1 proteins virtually abolished despite probe-dependent mRNA changes, and decreased Wnt activation genes at multiple levels (PORCN, WLS, LRP6, TNIK), while strongly inducing the antagonist DKK1 (**Fig. 6E**). HCT116 maintained or increased ALDH1A1, CD44, LRIG1 and SOX9, kept SOX2 mRNA unchanged with protein upregulation, and partially reduced CD133. Protein and gene alterations were confirmed by western blot and qPCR (**Fig. 6F, Supp. Fig S5C**). Functionally, NPC2 KD impaired Wnt3a-induced TCF/LEF transcriptional activity in both lines, to a markedly greater extent in KM12SM as determined by TOP/FOP assays (**Fig. 6G**), confirming the inhibition of Wnt responsiveness in the *APC*-mutant background.

Finally, EMT was induced in both KD lines with upregulated SNAI2, ZEB1 and SERPINE1, with E-cadherin and CDH17 showing opposite trends (**Fig. 6H, 6I, Supp. Fig. S5D**). However, only HCT116 coupled EMT to a productive survival and secretory program. HCT116 KD cells exhibited a coordinated increase of AP-1/YAP/TAZ response, immediate-early genes (EGR1–3), YAP/TAZ targets (CYR61, CTGF, AXL), and focal-adhesion genes (VCL, TLN1, PTK2), whereas KM12SM KD cells showed an unbalanced AP-1 response (strong JUN induction without FOS increase) and reduced CTGF protein, with opposite LATS1 regulation (**Fig. 6H, 6I, Supp. Fig. S5D**). HCT116 KD cells additionally engaged all three branches of the unfolded-protein response (UPR) (PERK, IRE1/XBP1 and ATF6; while KM12SM involved ATF6 only), induced the ferroptosis-protective SLC7A11–CoQ axis, and developed a pro-invasive secretome profile (metalloproteases, PLAUR, inflammatory cytokines and receptors) (**Supp. Table S2**). KM12SM instead suppressed growth-factor signaling (IGF1R, ERBB3, VEGFA, neuropilins) (**Supp. Table S1**). Altogether, NPC2 loss induces an EMT-like response in both models, but only HCT116 showed increased expression of genes involved in AP- 1/YAP/TAZ signaling, integrated stress adaptation, ferroptosis protection and a pro- invasive secretory program, displaying a transcriptional signature of its preserved survival and metastatic competence.

### Lipidomic analysis identify lipid-level mechanisms underlying *APC*-dependent divergent metastatic progression

To dissect the metabolic basis of synthetic lethality upon NPC2 depletion, we performed quantitative lipidomics. After normalization and principal component analysis, samples showed adequate reproducibility and sample separation according to experimental conditions (**Supp. Fig. S7A, 7B**). Heatmap analysis indicated distinct clusters of increased and decreased lipid species, highlighting significant differences between both cell lines (**Supp. Fig. S7C).** Still, cross-line lipid correlation (r = 0.41) substantially exceeded the transcriptome correlation (r = 0.06), consistent with tight biophysical constraints on membrane composition (**Supp. Fig. S8A**). NPC2 KD produced a 3.4-fold larger lipidomic perturbation in KM12SM, 536 differentially-expressed lipids (DELs), than HCT116 (158 DELs) (**Supp. Table S3, Supp. Fig. S8B**). Of 106 common DELs, 80 changed concordantly and 26 changed in opposite directions (**Fig. 7A)**. The cholesteryl-ester pool was selectively remodeled in KM12SM, which accumulated very- long-chain polyunsaturated fatty acids (VLC-PUFAs) such as ChE(24:5) (+3.84), ChE(26:8) (+3.48), ChE(22:5) (+2.42), ChE(22:4) (+2.04), a pattern mostly absent in HCT116 (**Fig. 7B**). This pattern is consistent with esterification of membrane-derived VLC-PUFAs with the limited available cholesterol as a compensatory response to preserve membrane fluidity ^35^. Conversely, shorter-chain ChE species were depleted in KM12SM, including ChE(18:1), with a similar, non-significant trend in HCT116, consistent with a limited supply of fatty-acyl substrates for cholesterol esterification at the endoplasmic reticulum. Free cholesterol showed a trend toward reduction in KM12SM, but was stable in HCT116 (**Fig. 7C**), consistent with the cholesterol auxotrophy model.

**Fig. 7.**
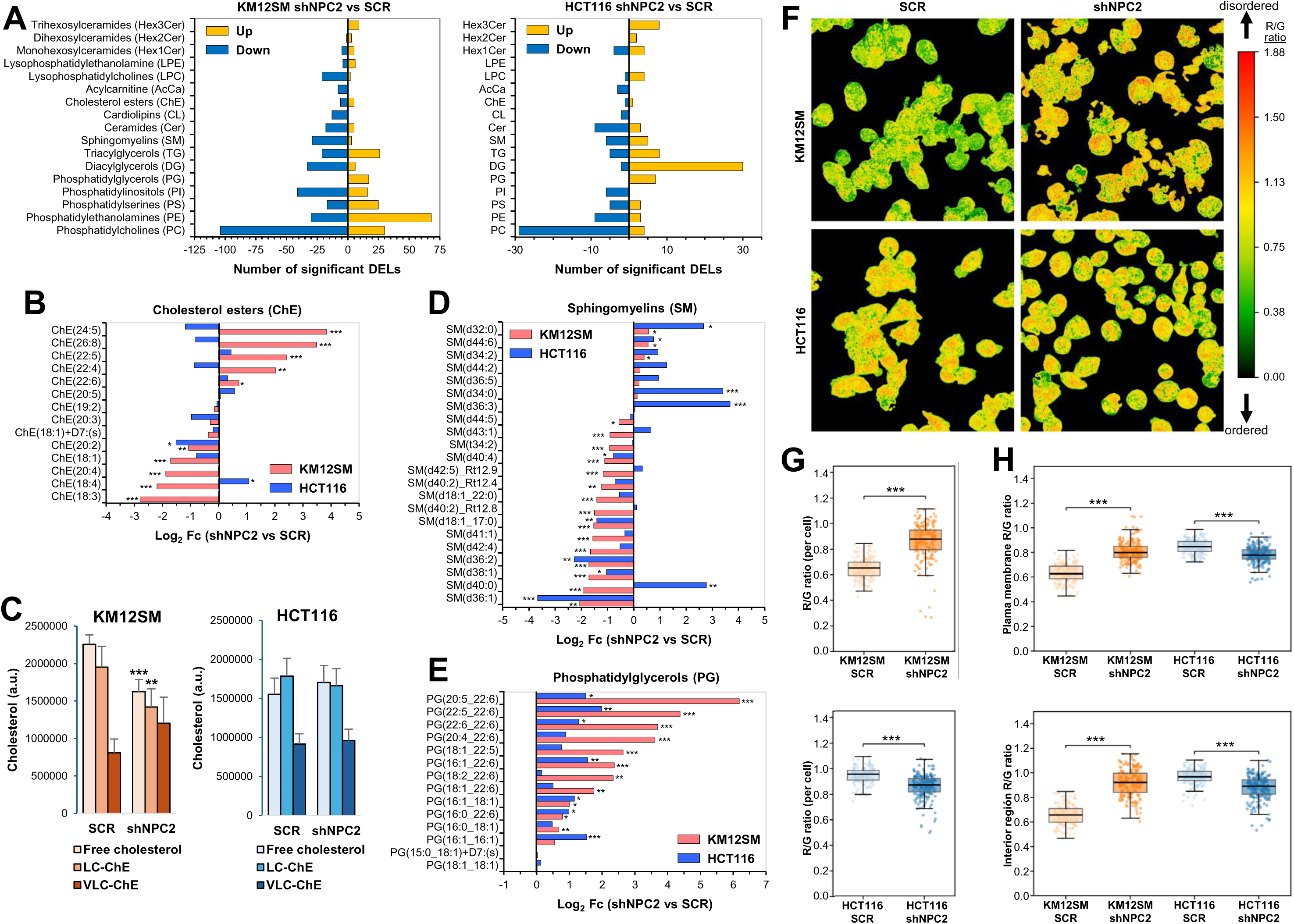
NPC2 knockdown differentially remodels the lipidome and membrane order in colorectal cancer cells. (A) Number of significantly different lipid species (DELs) per lipid class in KM12SM (left) and HCT116 (right) cells expressing shNPC2 relative to scrambled control (SCR). Differential lipids were defined by Benjamini–Hochberg- adjusted P < 0.05 and |log2 fold change| ≥ 0.585 (1.5-fold; n = 4 biological replicates per condition). Yellow, increased; blue, decreased. (B) Log₂ fold changes (shNPC2 vs SCR) of cholesteryl esters (ChE). Quantification of: (C) free cholesterol and long chain (LC) and very long chain (VLC) cholesteryl esters (ChE) in the indicated cells (**, p < 0.01; ***, p < 0.001), (D) sphingomyelins (SM), and (E) phosphatidylglycerols (PG). Lipid species are ordered according to decreasing log_2_ FC in KM12SM cells. Red, KM12SM; blue, HCT116. *, p< 0.05, **P < 0.01, ***, p< 0.001. +D7:(s) denotes the deuterated internal standard. (F) Representative ratiometric LipiORDER images of live SCR and NPC2-KD cells (shNPC2). The R/G ratio reports membrane order, with higher values (red/orange) indicating greater membrane disorder and lower values (green) indicating greater membrane order. (G) Quantification of the per-cell R/G ratio from the indicated number of cells obtained from three independent cultures. (H) Quantification of R/G ratio of the PM (upper) and interior region compartments from segmented images. Boxes indicate median and interquartile range; whiskers indicate the 5th–95th percentiles (***, p< 0.001).

Lipid rafts are cholesterol- and sphingolipid-enriched membrane microdomains that serve as signaling platforms. Their two lipid components were differentially disrupted: sphingomyelins were near-universally depleted in KM12SM (29/43 species down, 2 up) but balanced in HCT116 (5 up, 4 down) (**Fig. 7A, D**), indicating lipid raft collapse and supporting the loss of raft-associated signaling. The strongest discordance was observed in diacylglycerols (DG), 12 DG species, including DG(18:0_18:1), DG(18:0_20:2), DG(18:0_22:3) and DG(26:0_18:1) accumulated in HCT116 while were selectively lost in KM12SM (**Supp. Fig. S9A**). Accordingly, HCT116 maintained raft- dependent phospholipase C (PLC) signaling by depleting phosphatidylinositol (PI) substrate (6 species down, none up), while KM12SM lost both, the PI substrate (42/85 down) and the signaling output. The PKCα-encoding gene PRKCA was downregulated in KM12SM (−0.63, FDR = 0.016) with a non-significant increasing trend in HCT116 (**Supp. Tables S1, S2**). Thus, SM collapse and opposite DG regulation mirrored, at the lipid level, the divergent raft-signaling phenotypes observed experimentally.

Mitochondrial lipids followed the same divergence. Cardiolipins (CL), essential for inner membrane integrity, respiratory supercomplex assembly and cytochrome c retention were depleted in KM12SM (11/20 species down) but essentially preserved in HCT116 (2/20) (**Supp. Fig. S9B**). The CL precursor phosphatidylglycerol (PG) accumulated in KM12SM (18/26 species up; 5/26 in HCT116) (**Fig. 7E**), indicating failed CL biosynthesis and/or remodeling, a Barth syndrome-like signature ^36^, and supporting mitochondrial dysfunction in the auxotroph, consistent with its pro-apoptotic transcriptomic signature (BAX up, MCL1 down). Sphingolipid metabolism diverged accordingly, HCT116 accumulated C24 dihydroceramides (Cer(d18:0_24:0) +2.43, Cer(d18:0_24:1) +1.28, Cer(d18:0_22:0) +1.12), cytoprotective precursors of mature ceramides ^37^, whereas KM12SM failed to accumulate them and showed severe depletion of sphingosine (−2.80) and the stress-signaling phytoceramide Cer(t18:0_24:1) (−4.84) (**Supp. Fig. S9C**). Lysophosphatidylcholines were broadly depleted in KM12SM (22/53 down) but maintained or increased in HCT116 (**Supp. Fig. S9D**), a pattern associated with membrane-bilayer disruption, cholesterol oxidation and transfer of PM cholesterol into intracellular compartments, as observed in KM12SM (**Fig. 2A**).

To validate the biophysical consequences of this remodeling, we performed membrane lipid order imaging in live cells using the solvatochromic probe LipiORDER, whose red/green (R/G) emission ratio is low in liquid-ordered (Lo) and high in liquid- disordered (Ld) membranes (**Fig. 7F**). The two cell lines showed distinct baseline states: KM12SM maintained a uniformly ordered membrane system (cortex R/G 0.63, interior 0.66), whereas HCT116 was more disordered, particularly in its interior (cortex 0.85, interior 0.97). NPC2 KD again acted in opposite directions (**Fig. 7G)**. In KM12SM, order collapsed in both compartments, most severely in the interior (cortex 0.63 → 0.80; interior 0.66 → 0.92; interior–cortex difference Δ = 0.03 → 0.12, r = 0.56), consistent with depletion of the saturated SM scaffold and PM cholesterol (**Fig. 7H)**). In HCT116, order was maintained or mildly increased in both compartments (cortex 0.85 → 0.78; interior 0.97 → 0.89), consistent with compensatory accumulation of saturated sphingolipid species (C24 dihydroceramides up to +2.43; 8/9 Hex3Cer species increased). Therefore, NPC2 depletion causes a pronounced loss of membrane order in the cholesterol-auxotrophic model, while the cholesterol-adaptable line maintains membrane organization. Collectively, the lipidome and membrane biophysics discriminate the two phenotypes. In APC-mutant KM12SM, NPC2 KD cells produced raft lipid collapse, global loss of membrane order, CE-pool remodeling without net storage, mitochondrial CL failure and sphingolipid depletion, converging on stemness loss, cell-cycle arrest and pro-apoptotic programming. In HCT116 NPC2 KD, NPC1 compensation maintains cholesterol flux, raft integrity, DG signaling and protective dihydroceramide accumulation, preserving membrane order, viability and tumorigenicity.

### NPC2-deficient CRC cells recapitulate selected Niemann–Pick type C hallmarks in an APC-dependent manner

We next assessed whether NPC2 depletion in CRC cells recapitulated the cellular hallmarks associated with Niemann–Pick type C disease by comparing nine predefined hallmarks with the lipidomic and transcriptomic profiles of NPC2-KD cells using a scorecard (**Supp. Fig. S10)** derived from the quantitative tables (**Supp. Tables S1, S2 and S3**). Glycosphingolipid accumulation emerged as the most robust shared phenotype, with marked elevation of Hex3Cer/Gb3 species, a recognized secondary storage lipid in NPC disease, in both KM12SM and HCT116 cells, whereas upstream glycosphingolipid intermediates showed heterogeneous changes. In contrast, several metabolic and signaling features were predominantly restricted to KM12SM cells, including extensive sphingomyelin loss, sustained cholesterol-biosynthetic activity, CoQ depletion, oxidative-stress responses and activation of Rag-dependent nutrient sensing. These changes were largely absent in HCT116 cells, which instead suppressed cholesterol synthesis and preserved CoQ levels. Thus, NPC2 depletion induces a common NPC-like glycosphingolipid-storage phenotype, while the accompanying metabolic-stress response is strongly influenced by the cellular context and is most pronounced in cholesterol- auxotrophic APC-mutant cells, providing a rationale for evaluating NPC-targeted strategies in a genotype-dependent context.

### NPC2 expression associates with poor prognosis in CRC and other tumors

To further investigate the clinical relevance of NPC2 we performed *<u>in silico</u>* analysis of human CRC datasets. NPC2 showed higher expression in tumoral tissues than in normal mucosa samples (**Fig. 8A**), being preferentially expressed in the aggressive mesenchymal subtype CRIS-B, (**Fig. 8B**). The prognostic value of NPC2 was supported at the single- gene level using the Kaplan–Meier Plotter tool, where high NPC2 expression was associated with poorer relapse-free survival (RFS) and overall survival (OS) in a cohort of 1,336 CRC patients (**Fig. 8C**). Consistently, analysis of the independent GSE39582 cohort revealed a significant association between high NPC2 expression and poorer outcome in premetastatic stages (**Fig. 8D**). Association with lower survival was also preferentially observed in the *KRAS* wild-type subgroup *vs KRAS* mutant (p = 0.038; **Fig. 8E**). Additional prognostic analyses revealed that high NPC2 expression was associated with poorer outcome across several CRC intrinsic subtypes, including CRIS-B, CRIS-C and CRIS-D (**Fig. 8F**), which are respectively characterized by prominent TGFβ/EMT features, enhanced RTK signaling, and high Wnt activity with stem cell-like properties. Although, the downstream pathway differs among subtypes, all three rely heavily on signaling processes that are sensitive to membrane cholesterol organization. Beyond CRC, KM Plotter analysis of RNA-seq datasets revealed that elevated NPC2 expression also associated with reduced overall survival in esophageal squamous cell carcinoma, kidney renal clear cell carcinoma, liver hepatocellular carcinoma, rectum adenocarcinoma, stomach adenocarcinoma and thymoma (**Supp. Fig. S11**). Collectively, these *in silico* analyses support a broad adverse prognostic association for high NPC2 expression in CRC and several additional tumor types.

**Figure 8.**
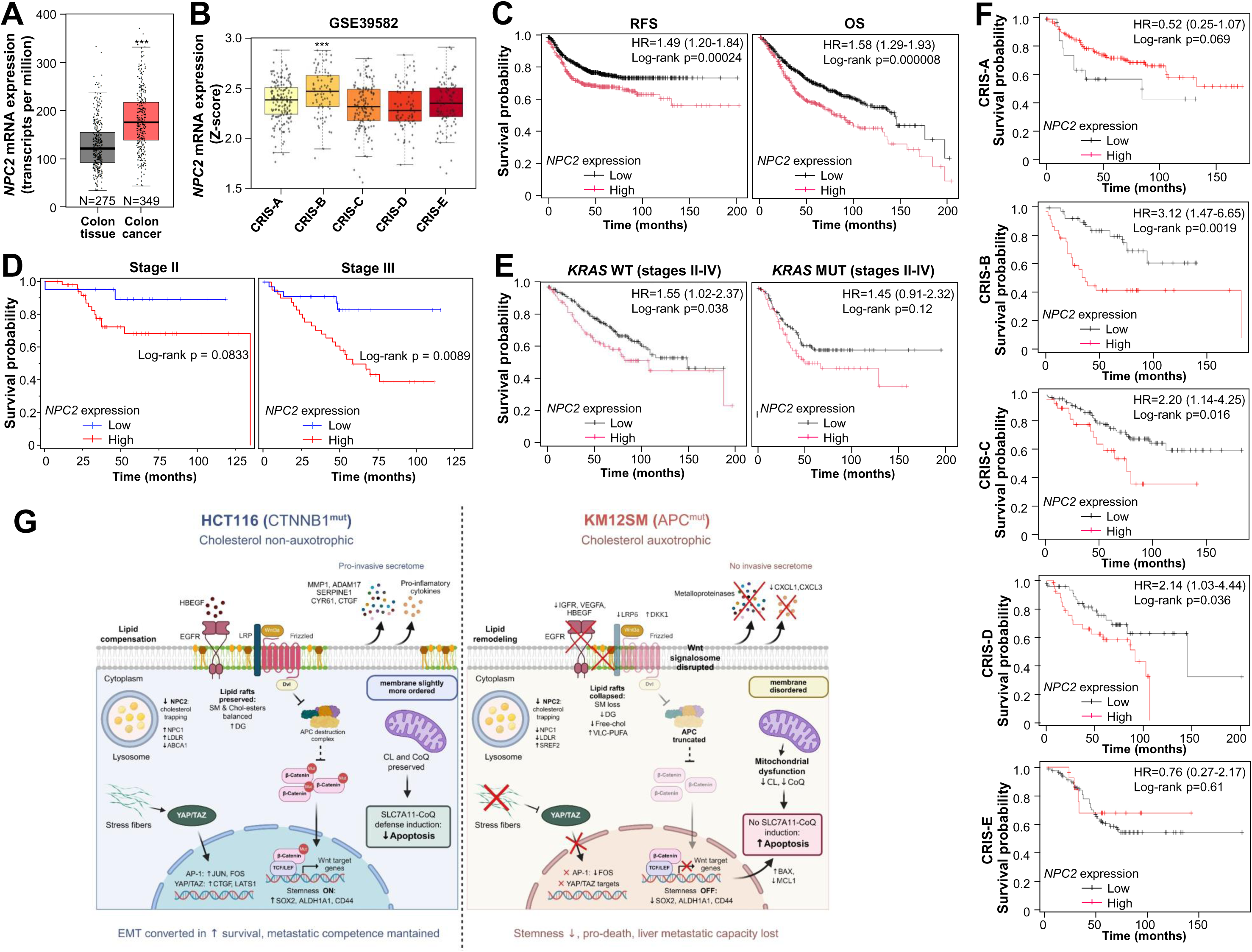
High NPC2 expression associates with aggressive colorectal cancer features and poor clinical outcome. (A) NPC2 mRNA expression in normal colon and CRC tissues from the TCGA-COAD cohort. NPC2 expression was significantly increased in tumor samples (***, p< 0.001). (B) NPC2 expression across CRIS molecular subtypes in the GSE39582 CRC cohort, showing significantly higher expression in the mesenchymal CRIS-B subtype (***, p< 0.001). (C) Kaplan–Meier analysis of relapse-free survival (RFS) and overall survival (OS) according to NPC2 expression using the KM-Plotter platform. Log-rank P values and hazard ratios are indicated within the plots. (D) Overall survival analysis of stage II–III CRC patients stratified according to NPC2 expression in the GSE39582 cohort. (E) Kaplan–Meier overall survival analysis according to NPC2 expression within the indicated CRIS molecular subtypes. (F) Kaplan–Meier survival analysis of stage II–IV CRC patients stratified according to NPC2 expression and KRAS mutational status. (G) Cholesterol auxotrophy determines the response to NPC2 loss and metastatic outcome. NPC2 depletion impairs lysosomal cholesterol export and restricts cholesterol delivery to PM in both models. In CTNNB1-mutant, cholesterol-adaptable cells, this defect is compensated by metabolic and signaling rewiring, including a pro- invasive secretory program, allowing xenograft growth and metastatic competence to be maintained. In contrast, APC-mutant, cholesterol-auxotrophic cells cannot restore cholesterol egress. The resulting membrane disorder disrupts raft-dependent signaling, collapses Wnt/stemness programs, and is accompanied by failure of AP-1/YAP–TAZ adaptation, ultimately suppressing tumor growth and liver metastasis.

## DISCUSSION

Cholesterol auxotrophy is emerging as a potential metabolic vulnerability in tumors requiring exogenous cholesterol for maintaining tumor growth and metastasis ^38^. Here, we identify a marked cholesterol-auxotrophic phenotype in *APC*-truncated CRC cells, in contrast to *CTNNB1*-mutant models, and show that this difference determines their dependency on NPC2-mediated lysosomal cholesterol export. NPC2 depletion in *APC*- mutant cells caused PM cholesterol loss and lysosomal cholesterol accumulation, recapitulating key features of Niemann–Pick disease ^39^, and was accompanied by profound impairment of proliferation, clonogenicity, invasion and metastatic competence. Multi-omic analyses revealed two distinct adaptive outcomes: cholesterol- auxotrophic *APC*-mutant cells failed to restore cholesterol homeostasis, leading to loss of membrane organization and receptor-dependent signaling, whereas *CTNNB1*-mutant cells exhibited pro-survival and pro-invasive rewiring (**Fig. 8G**). These findings define NPC2 as a selective vulnerability of cholesterol-auxotrophic CRC and provide a mechanistic basis for the differential response of *APC*- and *CTNNB1*-mutant tumors to disruption of lysosomal cholesterol trafficking ^7,40^. Consistent with this biological role, high NPC2 expression showed association with poor outcome in aggressive CRC subtypes and in several additional cancer types.

Our data identify the *APC/CTNNB1* mutational context as the primary determinant of NPC2 dependency in CRC. Differences in microsatellite stability (KM12SM/HCT116, MSI; SW620, MSS) and KRAS status exclude both as primary determinants of auxotrophy, and although the auxotrophic lines harbor TP53 mutations, the mechanistic evidence points to *APC*-driven Wnt deregulation. The entire cascade operates through cholesterol trafficking, PM composition and raft-dependent Wnt signalosomes rather than p53-mediated mevalonate regulation ^13^. *APC* and *CTNNB1* NPC2 KD cell lines also diverge in AKT/mTORC1 regulation. APC-mutant displays profound AKT inhibition coexisting with strong, rapamycin- and Torin-sensitive S6 phosphorylation, indicating AKT-independent mTORC1 activation through the Rag–Ragulator pathway ^41^. *CTNNB1*- mutant retains AKT activation with only modest pS6 elevation and restores cholesterol egress likely through NPC1, without requiring induction of the lysosomal sensor SLC38A9 ^42^. The result is a dysregulated state in which mTORC1 activity persists despite trapped lysosomal cholesterol, resembling the cholesterol-decoupled mTORC1 hyperactivation of Niemann–Pick type C disease ^23^, yet cannot substitute for AKT- dependent survival signaling.

The reduced migration and invasion of NPC2-deficient HCT116 cells *in vitro* contrasts with their preserved metastatic competence *in vivo*, which likely reflects the different biological processes measured by the two assays. *In vitro* migration and invasion primarily assess cell-autonomous motility, whereas metastatic progression also depends on survival, extravasation, adaptation to the distant microenvironment and subsequent tumor growth. Indeed, the increased invasion of HCT116 NPC2-KD cells under lipoprotein-depleted conditions indicates that their invasive behavior is highly dependent on extracellular lipid availability. *In vivo*, access to circulating lipids and microenvironmental signals, together with the AP-1/YAP–TAZ-associated inflammatory and pro-invasive program induced by NPC2 loss, may therefore support survival and metastatic colonization despite reduced basal motility. Thus, NPC2 depletion appears to shift cholesterol-adaptable HCT116 cells toward a metastasis program driven more by metabolic adaptation, niche interactions and colonization capacity than by intrinsic motility. In contrast, cholesterol-auxotrophic KM12SM cells fail to compensate for NPC2 loss, resulting in concordant impairment of both *in vitro* invasive properties and *in vivo* metastatic progression due to large PM-associated signaling alterations.

A major consequence of NPC2 depletion in cholesterol-auxotrophic KM12SM cells was the disruption of membrane-associated signaling. Cholesterol-rich membrane domains provide organized platforms for the assembly and activation of receptors, kinases and downstream signaling complexes ^43^. Accordingly, loss of plasma-membrane cholesterol and membrane order in KM12SM cells was accompanied by inhibition of several pathways that depend on these domains for efficient signal propagation. This was particularly evident for canonical Wnt signaling, whose activation requires cholesterol- dependent organization of FZD/LRP5/6–DVL signaling complexes at the plasma membrane ^3,44^. NPC2 depletion also reduced Src/FAK activation, consistent with previous evidence linking membrane cholesterol and lipid microdomains to integrin– FAK/Src signaling and showing that delivery of LDL-derived cholesterol from endolysosomal compartments promotes FAK activation and cell adhesion ^45,46^. Transcriptomic analyses further revealed broad suppression of RTK–RAS–PI3K/AKT and related growth-survival pathways, supporting a generalized loss of membrane- dependent signaling competence. By contrast, HCT116 cells largely preserved membrane order and maintained Src/FAK and PI3K/AKT signaling under serum-replete conditions despite reduced plasma-membrane cholesterol, consistent with compensatory cholesterol trafficking and lipid remodeling that preserve signaling-competent membrane domains. Thus, NPC2 loss differentially affects the organization of cholesterol-dependent signaling platforms: these are profoundly disrupted in cholesterol-auxotrophic KM12SM cells but largely maintained in cholesterol-adaptable HCT116 cells.

Lipidomics provided independent support for the alterations in cell signaling pathways identified by transcriptomic profiling. In cholesterol-auxotrophic KM12SM cells, NPC2 depletion caused extensive membrane lipid remodeling, with loss of SM, DG and mitochondrial CL, consistent with disruption of cholesterol-rich signaling domains and mitochondrial dysfunction ^47^. In contrast, HCT116 cells largely preserved SM, maintained cholesteryl-ester turnover and accumulated signaling and cytoprotective lipids, including DG and dihydroceramides, indicating a more effective metabolic adaptation to NPC2 loss ^35,48^. Accordingly, membrane order was maintained in HCT116, likely through compensatory sphingolipid remodeling ^49,50^, whereas KM12SM showed marked membrane disorder together with cardiolipin and CoQ depletion, consistent with increased apoptotic and ferroptotic vulnerability ^51–53^. Notably, Gb3 accumulated in both models, as reported in Niemann–Pick disease ^54,55^, indicating that lipid storage is a common consequence of NPC2 loss, whereas failure of membrane and metabolic adaptation is restricted to the cholesterol-auxotrophic context

Interestingly, transcriptomics and lipidomics revealed some commonalities between CRC NPC2-depleted cells and Niemann–Pick disease or other lysosomal storage disorders ^31,56^. These include enrichment of inflammatory pathways, increased authophagy ^31^ and reduced bile acid metabolism ^57^. The hepatomegaly observed following HCT116 KD inoculation in mice, also observed in NPC2 patients ^58^, suggests that these similarities extend to tissue-level pathology. Moreover, NPC2 depletion induced an upregulation of genes involved in lysosomal acidification, including ATP6V subunits ^59^, vesicular trafficking (LYST, BLOC1 complex) ^60^, and lipid metabolism at different extent in each cell line, suggesting an adaptive lysosomal remodeling program. Notably, several of these genes are implicated in other lysosomal storage disorders, including Chediak– Higashi syndrome (LYST) ^61^, sialidosis (NEU1) ^62^, and Hermansky–Pudlak syndrome (BLOC1 complex) ^63^. These findings suggest that NPC2-deficient CRC models may provide a useful system for investigating broader mechanisms of lysosomal dysfunction.

Several NPC2-independent, or partially redundant, traffic pathways may help to explain the divergent capacity of HCT116 and KM12SM to compensate for NPC2 loss. LIMP-2/SCARB2, a lysosomal integral membrane protein, operates in parallel with the NPC1/NPC2 axis, binding and delivering cholesterol to the limiting membrane at a slower rate ^64^, while oxysterol-binding protein (OSBP)-related proteins (ORP) family members redistribute cholesterol from endolysosomal compartments to the ER and PM through membrane contact sites ^65–67^. Our transcriptomic data revealed that HCT116 mounted a coordinated bypass response including upregulation of NPC1, OSBP, ORP2, and VAPA, whereas KM12SM failed to induce NPC1 and shows reduced expression of several alternative trafficking components, including LIMP-2/SCARB2, and ORP family members. This cascading failure of all compensatory egress routes, in a cholesterol- auxotrophic background that cannot reuse *de novo* synthesis, explains NPC2 loss as synthetically lethal in KM12SM, but tolerable in HCT116.

The preferential association of high NPC2 expression with poor prognosis in CRIS-B, CRIS-C and CRIS-D tumors is consistent with the mechanistic functions identified in our experimental models. Although these subtypes are biologically distinct, they are characterized by prominent TGFβ/EMT and invasive programs (CRIS-B), RTK signaling (CRIS-C), or Wnt-driven stemness (CRIS-D), all of which depend to varying degrees on membrane organization and cholesterol-sensitive signaling. Our findings suggest that NPC2 may provide a common lipid-trafficking support system for these aggressive tumor states by sustaining receptor signaling, focal- adhesion/mechanotransduction pathways and Wnt-dependent stemness. Thus, the adverse prognostic association of NPC2 across CRIS-B-C-D may reflect a broader requirement for efficient cholesterol trafficking in tumors with high signaling and phenotypic plasticity.

In summary, these findings identify NPC2-dependent cholesterol trafficking as a critical determinant of tumor growth and metastatic fitness in *APC*-mutant CRC and support its potential therapeutic exploitation in this genetic context. Our model is consistent with recent evidences showing that inner-leaflet plasma membrane cholesterol is elevated in *APC*-truncated CRC cells and promotes Wnt-independent signalosome assembly ^29^. In contrast, *CTNNB1*-mutant cells induce compensatory mechanisms that preserve cholesterol homeostasis and sustain an adaptive, pro-invasive state, highlighting the existence of distinct genotype-specific responses to impaired cholesterol trafficking. From a therapeutic perspective, NPC2 represents an attractive target because, as a soluble lysosomal protein with extracellular availability, it may be amenable to strategies such as antibody-based neutralization or small molecules targeting its cholesterol-binding pocket.

### Limitations of the study

Although our findings strongly support a major role for APC mutations in determining cholesterol auxotrophy, CRC is a highly heterogeneous disease with many different combinations of genetic alterations. Therefore, we cannot exclude that, in some molecular backgrounds, additional mutations may modify or even override the influence of the APC/TP53 genotype on cholesterol dependency. Further studies using a broader panel of CRC models will be needed to establish the generalizability of this relationship.

## Materials and methods

Further detailed methodology can be found in **Supplementary Information**

### Cell culture conditions and stable silencing with lentiviruses

KM12SM cells were obtained from the MD Anderson Cancer Center (USA). SW620, HT29 and SW48 cells were obtained from the ATCC (USA), whereas HCT116 were from DSMZ (Germany). They were regularly authenticated by SPR determination. Cells were grown in DMEM supplemented with fetal bovine serum (10%) and antibiotics, under standard conditions (37°C, humidified atmosphere with 5% CO₂), and used within 6 months of passaging for all experiments. Cells were regularly tested for mycoplasma contamination. NPC2 KD was achieved by generating stable cell lines using two independent shRNAs (Merck): #21 TRCN0000029321 and #23 TRCN0000029323. Scrambled vector was generated with pGFP-C-shLenti #TR30021. Lentiviruses were generated in HEK293T cells using Lenti-X™ Packaging Single Shots, VSV-G (#631275, Takara, USA) according to the manufactureŕs protocol. For transient silencing, siRNAs targeting NPC2 (SASI-HS01_00239264) or control-scrambled siRNA (5’- AUUGUAUGCGAUCGCAGACCdTdT-3’) were purchased from Merck (Darmstadt, Germany) and transfected in cancer cells using JetPrime Reagent PolyPlus (Illkirch, France).

### Auxotrophy and cell viability assay

For testing cell auxotrophy, 2×10^4^ cells of each cell line/condition were seeded in triplicate on 6-well plates and supplemented with either DMEM containing 10% FBS, 10% DLPS (S181L-500; Biowest), 10% DLP plus 10 µg/ml cholesterol (C8667; Merck). Every two days, cells were trypsinized and counted with a TC10 Automated Cell Counter (Bio-Rad). For cell viability, cells were seeded at 1 × 10^4^ cells/well on 96-well plates and incubated for 48 h at 37°C in DMEM with 0.5% serum, followed by 1 h incubation with thiazolyl blue tetrazolium bromide (MTT; 0.5 mg/mL; Sigma-Aldrich). Cell viability was determined by A_560_ nm and comparison with control cells collected *ad initium* (input).

### Filipin staining

For filipin staining, 2×10^4^ cells were plated on a Matrigel-coated coverslip per well in 24- well plates. As a positive control for PM cholesterol depletion, we used MβCD (Methyl- β-cyclodextrin) solution (1:50) incubated at 37°C for 15 min. Cells were fixed, washed with PBS and then incubated with filipin (#SAE0088, Merck) for 30 min, blocked (4% BSA) at RT for 30 min. Finally, re-incubated with filipin for 30 min at RT, and mounted in slides. Filipin staining was viewed by fluorescence microscopy using a DAPI filter. Images were acquired with a laser-scanning confocal microscope (LSM700, Zeiss; Germany) using Plan Apochromat ×63 oil immersion objective lens and analysed with Zen Lite 2012 software.

### Lipid Extraction

Total lipids were extracted using a modified Folch method ^68^. Briefly, 1 × 10⁶ cells were lysed in 200 μL ice-cold lysis buffer and cleared by centrifugation (14,000 rpm, 10 min, 4°C). Supernatants were subjected to biphasic extraction with chloroform:methanol (2:1, v/v). Following centrifugation (14,000 rpm, 10 min), the organic phase was collected, vacuum-dried at 35°C, and stored at −20°C under desiccated conditions to prevent oxidation. Solvent volumes were scaled proportionally to cell number.

### Statistical analyses

All experiments were performed in triplicate, and data are presented as the mean ± standard deviation (SD). Correlations between gene expression levels were evaluated using Pearson’s correlation coefficient. Kaplan–Meier survival curves were compared using the log-rank test. Data normality was assessed using the Kolmogorov–Smirnov test, and homogeneity of variances was evaluated using the F-test. Comparisons between two groups were performed using a Student’s *t*-test, whereas comparisons involving more than two groups were analysed by one-way ANOVA followed by Tukey’s multiple comparison test. In all statistical analyses, the minimum acceptable level of significance was p < 0.05.

## Supporting information

Supplementary methodology

## ACKNOWLEDGEMENTS

We thank Francisco Casado for technical assistance. Language polishing was made with the help of ChatGPT. IB and MC were supported by IND2022/BMD-23554 and IND2024/BMD-34329 fellowship grants of the Comunidad de Madrid. This project was supported by grants PID2021-122227OB-I00 and PID2024-156301OB-I00 (MICIU) with funds MCIN/AEI/10.13039/501100011033 and “Next-Generation EU”/PRTR.

## AUTHOŔS CONTRIBUTIONS

JIC supervised the experiments. IB, RAB, MC, LP, TC, ME participated in the experiments. PK, PB, PP, DD and FM performed and supervised the lipidomic experiments. IB, RAB, FM, JII and JIC analyzed and interpreted the experiments. RAB and JIC wrote the manuscript. JIC and JII obtained the funding.

## REFERENCES

1. Lauridsen, A.R., Skorda, A., Winther, N.I., Bay, M.L., and Kallunki, T. (2024). Why make it if you can take it: review on extracellular cholesterol uptake and its importance in breast and ovarian cancers. J Exp Clin Cancer Res 43, 254. 10.1186/s13046-024-03172-y.

2. Liu, X., Lv, M., Zhang, W., and Zhan, Q. (2023). Dysregulation of cholesterol metabolism in cancer progression. Oncogene 42, 3289–3302. 10.1038/s41388-023-02836-x.

3. Erazo-Oliveras, A., Munoz-Vega, M., Mlih, M., Thiriveedi, V., Salinas, M.L., Rivera-Rodriguez, J.M., Kim, E., Wright, R.C., Wang, X., Landrock, K.K., et al. (2023). Mutant APC reshapes Wnt signaling plasma membrane nanodomains by altering cholesterol levels via oncogenic beta-catenin. Nat Commun 14, 4342. 10.1038/s41467-023-39640-w.

4. Nes, W.D. (2011). Biosynthesis of cholesterol and other sterols. Chem Rev 111, 6423–6451. 10.1021/cr200021m.

5. Cruz, P.M., Mo, H., McConathy, W.J., Sabnis, N., and Lacko, A.G. (2013). The role of cholesterol metabolism and cholesterol transport in carcinogenesis: a review of scientific findings, relevant to future cancer therapeutics. Front Pharmacol 4, 119. 10.3389/fphar.2013.00119.

6. Simons, K., and Toomre, D. (2000). Lipid rafts and signal transduction. Nat Rev Mol Cell Biol 1, 31–39. 10.1038/35036052.

7. Riscal, R., Bull, C.J., Mesaros, C., Finan, J.M., Carens, M., Ho, E.S., Xu, J.P., Godfrey, J., Brennan, P., Johansson, M., et al. (2021). Cholesterol Auxotrophy as a Targetable Vulnerability in Clear Cell Renal Cell Carcinoma. Cancer Discov 11, 3106–3125. 10.1158/2159-8290.CD-21-0211.

8. Villa, G.R., Hulce, J.J., Zanca, C., Bi, J., Ikegami, S., Cahill, G.L., Gu, Y., Lum, K.M., Masui, K., Yang, H., et al. (2016). An LXR-Cholesterol Axis Creates a Metabolic Co-Dependency for Brain Cancers. Cancer Cell 30, 683–693. 10.1016/j.ccell.2016.09.008.

9. Garcia-Bermudez, J., Baudrier, L., Bayraktar, E.C., Shen, Y., La, K., Guarecuco, R., Yucel, B., Fiore, D., Tavora, B., Freinkman, E., et al. (2019). Squalene accumulation in cholesterol auxotrophic lymphomas prevents oxidative cell death. Nature 567, 118–122. 10.1038/s41586-019-0945-5.

10. Stone, L. (2021). Cutting cholesterol curbs clear cell RCC. Nat Rev Urol 18, 509. 10.1038/s41585-021-00510-z.

11. Du, W., Zhang, L., Brett-Morris, A., Aguila, B., Kerner, J., Hoppel, C.L., Puchowicz, M., Serra, D., Herrero, L., Rini, B.I., et al. (2017). HIF drives lipid deposition and cancer in ccRCC via repression of fatty acid metabolism. Nat Commun 8, 1769. 10.1038/s41467-017-01965-8.

12. Raftopulos, N.L., Washaya, T.C., Niederprum, A., Egert, A., Hakeem-Sanni, M.F., Varney, B., Aishah, A., Georgieva, M.L., Olsson, E., Dos Santos, D.Z., et al. (2022). Prostate cancer cell proliferation is influenced by LDL-cholesterol availability and cholesteryl ester turnover. Cancer Metab 10, 1. 10.1186/s40170-021-00278-1.

13. Moon, S.H., Huang, C.H., Houlihan, S.L., Regunath, K., Freed-Pastor, W.A., Morris, J.P.t., Tschaharganeh, D.F., Kastenhuber, E.R., Barsotti, A.M., Culp-Hill, R., et al. (2019). p53 Represses the Mevalonate Pathway to Mediate Tumor Suppression. Cell 176, 564–580 e519. 10.1016/j.cell.2018.11.011.

14. Liang, B., Chen, R., Song, S., Wang, H., Sun, G., Yang, H., Jing, W., Zhou, X., Fu, Z., Huang, G., and Zhao, J. (2019). ASPP2 inhibits tumor growth by repressing the mevalonate pathway in hepatocellular carcinoma. Cell death & disease 10, 830. 10.1038/s41419-019-2054-7.

15. You, W., Su, L., Weng, S., Li, J., Wang, X., Liang, B., Li, D., Li, L., and Chen, H. (2025). SREBP-2 promotes cancer progression through the mevalonate-Akt pathway in non-small cell lung cancer. Sci Rep 15, 23103. 10.1038/s41598-025-07437-0.

16. Codenotti, S., Asperti, M., Poli, M., Lorenzi, L., Pietrantoni, A., Cassandri, M., Marampon, F., and Fanzani, A. (2025). Synthetic inhibition of SREBP2 and the mevalonate pathway blocks rhabdomyosarcoma tumor growth in vitro and in vivo and promotes chemosensitization. Mol Metab 92, 102085. 10.1016/j.molmet.2024.102085.

17. Maier, C.R., Hartmann, O., Prieto-Garcia, C., Al-Shami, K.M., Schlicker, L., Vogel, F.C.E., Haid, S., Klann, K., Buck, V., Munch, C., et al. (2023). USP28 controls SREBP2 and the mevalonate pathway to drive tumour growth in squamous cancer. Cell Death Differ 30, 1710–1725. 10.1038/s41418-023-01173-6.

18. Robles, J., Pintado-Berninches, L., Boukich, I., Escudero, B., de Los Rios, V., Bartolomé, R.A., Jaén, M., Martín-Regalado, Á., Fernandez-Aceñero, M.J., Imbaud, J.I., and Casal, J.I. (2022). A prognostic six-gene expression risk-score derived from proteomic profiling of the metastatic colorectal cancer secretome. The journal of pathology. Clinical research 8, 495–508. 10.1002/cjp2.294.

19. Naureckiene, S., Sleat, D.E., Lackland, H., Fensom, A., Vanier, M.T., Wattiaux, R., Jadot, M., and Lobel, P. (2000). Identification of HE1 as the second gene of Niemann-Pick C disease. Science 290, 2298–2301. 10.1126/science.290.5500.2298.

20. Chikh, K., Vey, S., Simonot, C., Vanier, M.T., and Millat, G. (2004). Niemann-Pick type C disease: importance of N-glycosylation sites for function and cellular location of the NPC2 protein. Mol Genet Metab 83, 220–230. 10.1016/j.ymgme.2004.06.013.

21. Vanier, M.T. (2015). Complex lipid trafficking in Niemann-Pick disease type C. J Inherit Metab Dis 38, 187–199. 10.1007/s10545-014-9794-4.

22. Ikonen, E., and Olkkonen, V.M. (2023). Intracellular Cholesterol Trafficking. Cold Spring Harb Perspect Biol 15. 10.1101/cshperspect.a041404.

23. Davis, O.B., Shin, H.R., Lim, C.Y., Wu, E.Y., Kukurugya, M., Maher, C.F., Perera, R.M., Ordonez, M.P., and Zoncu, R. (2021). NPC1-mTORC1 Signaling Couples Cholesterol Sensing to Organelle Homeostasis and Is a Targetable Pathway in Niemann-Pick Type C. Dev Cell 56, 260–276 e267. 10.1016/j.devcel.2020.11.016.

24. Skorda, A., Lauridsen, A.R., Wu, C., Huang, J., Mrackova, M., Winther, N.I., Jank, V., Sztupinszki, Z., Strauss, R., Bilgin, M., et al. (2023). Activation of invasion by oncogenic reprogramming of cholesterol metabolism via increased NPC1 expression and macropinocytosis. Oncogene 42, 2495–2506. 10.1038/s41388-023-02771-x.

25. Kinzler, K.W., and Vogelstein, B. (1996). Lessons from hereditary colorectal cancer. Cell 87, 159–170. 10.1016/s0092-8674(00)81333-1.

26. Giles, R.H., van Es, J.H., and Clevers, H. (2003). Caught up in a Wnt storm: Wnt signaling in cancer. Biochim Biophys Acta 1653, 1–24. 10.1016/s0304-419x(03)00005-2.

27. Segditsas, S., and Tomlinson, I. (2006). Colorectal cancer and genetic alterations in the Wnt pathway. Oncogene 25, 7531–7537. 10.1038/sj.onc.1210059.

28. Shitoh, K., Furukawa, T., Kojima, M., Konishi, F., Miyaki, M., Tsukamoto, T., and Nagai, H. (2001). Frequent activation of the beta-catenin-Tcf signaling pathway in nonfamilial colorectal carcinomas with microsatellite instability. Genes Chromosomes Cancer 30, 32–37. 10.1002/1098-2264(2000)9999:9999<::aid-gcc1065>3.0.co;2-i.

29. Sharma, A., Zalejski, J., Bendre, S.V., Kavrokova, S., Hasdemir, H.S., Ozgulbas, D.G., Sun, J., Pathmasiri, K.C., Shi, R., Aloulou, A., et al. (2025). Cholesterol-targeting Wnt-beta-catenin signaling inhibitors for colorectal cancer. Nat Chem Biol 21, 1376–1386. 10.1038/s41589-025-01870-y.

30. Ilyas, M., Tomlinson, I.P., Rowan, A., Pignatelli, M., and Bodmer, W.F. (1997). Beta-catenin mutations in cell lines established from human colorectal cancers. Proc Natl Acad Sci U S A 94, 10330–10334. 10.1073/pnas.94.19.10330.

31. Liedtke, M., Volkner, C., Jurs, A.V., Peter, F., Rabenstein, M., Hermann, A., and Frech, M.J. (2021). Pathophysiological In Vitro Profile of Neuronal Differentiated Cells Derived from Niemann-Pick Disease Type C2 Patient-Specific iPSCs Carrying the NPC2 Mutations c.58G>T/c.140G>T. Int J Mol Sci 22. 10.3390/ijms22084009.

32. Barderas, R., Bartolomé, R.A., Fernandez-Aceñero, M.J., Torres, S., and Casal, J.I. (2012). High expression of IL-13 receptor α2 in colorectal cancer is associated with invasion, liver metastasis, and poor prognosis. Cancer Res 72, 2780–2790. 10.1158/0008-5472.can-11-4090.

33. Bartolome, R.A., Martin-Regalado, A., Pintado-Berninches, L., Robles, J., Ramirez-Gonzalez, M.A., Boukich, I., Sanchez-Gomez, P., Balyasnikova, I.V., and Casal, J.I. (2023). Schnurri-3 drives tumor growth and invasion in cancer cells expressing interleukin-13 receptor alpha 2. Cell death & disease 14, 742. 10.1038/s41419-023-06255-4.

34. Shimano, H., Horton, J.D., Hammer, R.E., Shimomura, I., Brown, M.S., and Goldstein, J.L. (1996). Overproduction of cholesterol and fatty acids causes massive liver enlargement in transgenic mice expressing truncated SREBP-1a. J Clin Invest 98, 1575–1584. 10.1172/JCI118951.

35. Vona, R., Iessi, E., and Matarrese, P. (2021). Role of Cholesterol and Lipid Rafts in Cancer Signaling: A Promising Therapeutic Opportunity? Front Cell Dev Biol 9, 622908. 10.3389/fcell.2021.622908.

36. Raja, V., Reynolds, C.A., and Greenberg, M.L. (2017). Barth syndrome: A life-threatening disorder caused by abnormal cardiolipin remodeling. J Rare Dis Res Treat 2, 58–62. 10.29245/2572-9411/2017/2.1087.

37. Lachkar, F., Ferre, P., Foufelle, F., and Papaioannou, A. (2021). Dihydroceramides: their emerging physiological roles and functions in cancer and metabolic diseases. Am J Physiol Endocrinol Metab 320, E122–E130. 10.1152/ajpendo.00330.2020.

38. Riscal, R., Skuli, N., and Simon, M.C. (2019). Even Cancer Cells Watch Their Cholesterol! Mol Cell 76, 220–231. 10.1016/j.molcel.2019.09.008.

39. Shin, S., Han, M.J., Patel, I., Welsh, A.M., Sverdlov, M., Cho, W., Cologna, S.M., Roux, P.P., and Yoon, S.O. (2025). mTOR inhibition reprograms cellular lipid homeostasis by inducing alternative lipid uptake and promoting cholesterol transport. Mol Cell 85, 3486–3504 e3487. 10.1016/j.molcel.2025.08.021.

40. Rattani, A., Anderson, C., Trim, W.V., Garita, E., Imada, S., Khawaled, S., Shin, H., Zhang, J., Yilmaz, O.H., and Kirschner, M. (2025). The Wnt/APC destruction complex targets SREBP2 in a beta-catenin-independent pathway to control cholesterol metabolism. bioRxiv. 10.1101/2025.10.12.681896.

41. Sancak, Y., Bar-Peled, L., Zoncu, R., Markhard, A.L., Nada, S., and Sabatini, D.M. (2010). Ragulator-Rag complex targets mTORC1 to the lysosomal surface and is necessary for its activation by amino acids. Cell 141, 290–303. 10.1016/j.cell.2010.02.024.

42. Castellano, B.M., Thelen, A.M., Moldavski, O., Feltes, M., van der Welle, R.E., Mydock-McGrane, L., Jiang, X., van Eijkeren, R.J., Davis, O.B., Louie, S.M., et al. (2017). Lysosomal cholesterol activates mTORC1 via an SLC38A9-Niemann-Pick C1 signaling complex. Science 355, 1306–1311. 10.1126/science.aag1417.

43. Sezgin, E., Levental, I., Mayor, S., and Eggeling, C. (2017). The mystery of membrane organization: composition, regulation and roles of lipid rafts. Nat Rev Mol Cell Biol 18, 361–374. 10.1038/nrm.2017.16.

44. Sheng, R., Kim, H., Lee, H., Xin, Y., Chen, Y., Tian, W., Cui, Y., Choi, J.C., Doh, J., Han, J.K., and Cho, W. (2014). Cholesterol selectively activates canonical Wnt signalling over non-canonical Wnt signalling. Nat Commun 5, 4393. 10.1038/ncomms5393.

45. Baillat, G., Siret, C., Delamarre, E., and Luis, J. (2008). Early adhesion induces interaction of FAK and Fyn in lipid domains and activates raft-dependent Akt signaling in SW480 colon cancer cells. Biochim Biophys Acta 1783, 2323–2331. 10.1016/j.bbamcr.2008.08.008.

46. Takahashi, K., Kanerva, K., Vanharanta, L., Almeida-Souza, L., Lietha, D., Olkkonen, V.M., and Ikonen, E. (2021). ORP2 couples LDL-cholesterol transport to FAK activation by endosomal cholesterol/PI(4,5)P(2) exchange. EMBO J 40, e106871. 10.15252/embj.2020106871.

47. Cooke, M., and Kazanietz, M.G. (2022). Overarching roles of diacylglycerol signaling in cancer development and antitumor immunity. Sci Signal 15, eabo0264. 10.1126/scisignal.abo0264.

48. Kraft, M.L. (2016). Sphingolipid Organization in the Plasma Membrane and the Mechanisms That Influence It. Front Cell Dev Biol 4, 154. 10.3389/fcell.2016.00154.

49. Levental, K.R., Malmberg, E., Symons, J.L., Fan, Y.Y., Chapkin, R.S., Ernst, R., and Levental, I. (2020). Lipidomic and biophysical homeostasis of mammalian membranes counteracts dietary lipid perturbations to maintain cellular fitness. Nat Commun 11, 1339. 10.1038/s41467-020-15203-1.

50. Ernst, R., Ballweg, S., and Levental, I. (2018). Cellular mechanisms of physicochemical membrane homeostasis. Curr Opin Cell Biol 53, 44–51. 10.1016/j.ceb.2018.04.013.

51. Choi, S.Y., Gonzalvez, F., Jenkins, G.M., Slomianny, C., Chretien, D., Arnoult, D., Petit, P.X., and Frohman, M.A. (2007). Cardiolipin deficiency releases cytochrome c from the inner mitochondrial membrane and accelerates stimuli-elicited apoptosis. Cell Death Differ 14, 597–606. 10.1038/sj.cdd.4402020.

52. Doll, S., Freitas, F.P., Shah, R., Aldrovandi, M., da Silva, M.C., Ingold, I., Goya Grocin, A., Xavier da Silva, T.N., Panzilius, E., Scheel, C.H., et al. (2019). FSP1 is a glutathione-independent ferroptosis suppressor. Nature 575, 693–698. 10.1038/s41586-019-1707-0.

53. Bersuker, K., Hendricks, J.M., Li, Z., Magtanong, L., Ford, B., Tang, P.H., Roberts, M.A., Tong, B., Maimone, T.J., Zoncu, R., et al. (2019). The CoQ oxidoreductase FSP1 acts parallel to GPX4 to inhibit ferroptosis. Nature 575, 688–692. 10.1038/s41586-019-1705-2.

54. Chang, T.Y., Reid, P.C., Sugii, S., Ohgami, N., Cruz, J.C., and Chang, C.C. (2005). Niemann-Pick type C disease and intracellular cholesterol trafficking. J Biol Chem 280, 20917–20920. 10.1074/jbc.R400040200.

55. Wheeler, S., and Sillence, D.J. (2020). Niemann-Pick type C disease: cellular pathology and pharmacotherapy. J Neurochem 153, 674–692. 10.1111/jnc.14895.

56. Platt, N., Speak, A.O., Colaco, A., Gray, J., Smith, D.A., Williams, I.M., Wallom, K.L., and Platt, F.M. (2016). Immune dysfunction in Niemann-Pick disease type C. J Neurochem 136 Suppl 1, 74–80. 10.1111/jnc.13138.

57. Nicoli, E.R., Huebecker, M., Smith, D., Morris, L., and Platt, F.M. (2017). Differential response of the liver to bile acid treatment in a mouse model of Niemann-Pick disease type C. Wellcome Open Res 2, 76. 10.12688/wellcomeopenres.12431.2.

58. Millat, G., Chikh, K., Naureckiene, S., Sleat, D.E., Fensom, A.H., Higaki, K., Elleder, M., Lobel, P., and Vanier, M.T. (2001). Niemann-Pick disease type C: spectrum of HE1 mutations and genotype/phenotype correlations in the NPC2 group. Am J Hum Genet 69, 1013–1021. 10.1086/324068.

59. Colacurcio, D.J., and Nixon, R.A. (2016). Disorders of lysosomal acidification-The emerging role of v-ATPase in aging and neurodegenerative disease. Ageing Res Rev 32, 75–88. 10.1016/j.arr.2016.05.004.

60. Dell’Angelica, E.C. (2004). The building BLOC(k)s of lysosomes and related organelles. Curr Opin Cell Biol 16, 458–464. 10.1016/j.ceb.2004.05.001.

61. Turner, M.E., Che, J., Mirhaidari, G.J.M., Kennedy, C.C., Blum, K.M., Rajesh, S., Zbinden, J.C., Breuer, C.K., Best, C.A., and Barker, J.C. (2024). The lysosomal trafficking regulator “LYST”: an 80-year traffic jam. Front Immunol 15, 1404846. 10.3389/fimmu.2024.1404846.

62. Khan, A., and Sergi, C. (2018). Sialidosis: A Review of Morphology and Molecular Biology of a Rare Pediatric Disorder. Diagnostics (Basel) 8. 10.3390/diagnostics8020029.

63. Bowman, S.L., Bi-Karchin, J., Le, L., and Marks, M.S. (2019). The road to lysosome-related organelles: Insights from Hermansky-Pudlak syndrome and other rare diseases. Traffic 20, 404–435. 10.1111/tra.12646.

64. Heybrock, S., Kanerva, K., Meng, Y., Ing, C., Liang, A., Xiong, Z.J., Weng, X., Ah Kim, Y., Collins, R., Trimble, W., et al. (2019). Lysosomal integral membrane protein-2 (LIMP-2/SCARB2) is involved in lysosomal cholesterol export. Nat Commun 10, 3521. 10.1038/s41467-019-11425-0.

65. Lim, C.Y., Davis, O.B., Shin, H.R., Zhang, J., Berdan, C.A., Jiang, X., Counihan, J.L., Ory, D.S., Nomura, D.K., and Zoncu, R. (2019). ER-lysosome contacts enable cholesterol sensing by mTORC1 and drive aberrant growth signalling in Niemann-Pick type C. Nat Cell Biol 21, 1206–1218. 10.1038/s41556-019-0391-5.

66. Zhao, K., Foster, J., and Ridgway, N.D. (2020). Oxysterol-binding protein-related protein 1 variants have opposing cholesterol transport activities from the endolysosomes. Mol Biol Cell 31, 793–802. 10.1091/mbc.E19-12-0697.

67. Dong, J., Du, X., Wang, H., Wang, J., Lu, C., Chen, X., Zhu, Z., Luo, Z., Yu, L., Brown, A.J., et al. (2019). Allosteric enhancement of ORP1-mediated cholesterol transport by PI(4,5)P(2)/PI(3,4)P(2). Nat Commun 10, 829. 10.1038/s41467-019-08791-0.

68. Folch, J., Lees, M., and Sloane Stanley, G.H. (1957). A simple method for the isolation and purification of total lipides from animal tissues. J Biol Chem 226, 497–509.

