## Supplementary methodology for "A cholesterol addiction creates an Achilles’ heel in *APC*-truncated colorectal cancer"

### Antibodies

For NPC2, we used 19888-1-AP (Proteintech, Rosemont, IL, USA) for Western blot, and AF8644 (R&D Systems, Minneapolis, MN, USA) for immunofluorescence. LAMP-1 (sc-20011) was purchased from SCBT (Dallas, Tx, USA) for immunofluorescence. APC #MAB3785 (Millipore Sigma-Aldrich, St. Louis, USA), LGR5 #144-10545-50 (Raybiotech, Atlanta, USA), β-catenin (#8814), pERK1/2 (#9106), ERK (#4695), pAKT (#3787), AKT (#2920), pSrc (#2101), pS6 (#4857), S6 (#2217), CD109 (#24765), LRRC8A (#24979) were from Cell Signaling Technologies (Danvers, MA, USA). Also, FAK (sc-271126), β-tubulin (sc-55529), RhoGDI (sc-373724), ABCG2 (sc-377176), NPC1 (sc-271334), SREBP-2 (sc-271616), and c-JUN (sc-74543) were from SCBT. SOX2 (E-AB-18159), and SQLE (E-AB-63722) were from Elabscience Biotechnology (Wuhan, China). Phospho-FAK (611722) was from BD Biosciences, and HGMCR (PA5-37367); from R&D Systems. We purchased SRC (AF3389), CDH1 (AF648), ALDH1A1 (60171-1-Ig), LDLR (10785-1-AP), ABCA1 (26564-1-AP), SOAT1 (13202-1-AP) and SLUG (12129-1-AP) from Proteintech (Rosemont, IL, USA), ZEB1 (HPA027524, Sigma-Aldrich, St. Louis, MO, USA), while AXL (F0178), CTGF (F1126), and LATS1 (F0232) were from Selleckchem, Houston, TX, USA). β-actin (TA811000S), and GAPDH (TA802519S) were from Origene (Rockville, MD, USA). The antibodies against LSS (A6930) and DKK1 (AW5504), were from Abclonal (Wuhan, China) and Abcepta (San Diego, CA, USA), respectively.

### Immunofluorescence

Cells were cultured on Matrigel-coated coverslips to reach 70% confluence, then fixed in 4% paraformaldehyde, prepared in PBS and permeabilized with 1% Triton X-100 in PBS. Cells were subsequently incubated for 1h in blocking solution (40 μg/mL human γ-globulin in PBS), followed by incubation for 2h or O/N with anti-NPC2 and anti-LAMP1 antibodies (10 μg/mL). After washing, cells were incubated with Alexa Fluor 488-conjugated anti-rabbit IgG and Alexa Fluor 555-conjugated anti-mouse IgG secondary antibodies (Invitrogen). Cells were then washed and counterstained with 4,6-diamidino-2-phenylindole (DAPI). Visualization was performed on a Leica SP8 confocal laser scanning microscope (Leica Microsystems, Wetzlar, Germany). Lysosomal area and intracellular to PM cholesterol ratios were calculated from images containing at least 100 cells, using MultiGauge (FujiFilm Life Sciences, Tokyo, Japan).

**TOP/FOP reporter assays**

TOP/FOP reporter assays were performed to assess the activity of Wnt/β-catenin signaling. In brief, 8xTOPFlash and 8xFOPFlash plasmids were transfected in CRC cells. Following different treatments, cells were lysed, and the luciferase activities of Firefly and Renilla were measured using the Dual-Luciferase Reporter Assays System (Promega, Madison, WI), following the manufacturer's instructions. The firefly luciferase gene in TOPFlash is controlled by a promotor regulated by β-catenin, whereas such promotor is mutated in FOPFlash vector, setting the basal luminescence. Luciferase activity was normalized by transfection efficiency using the activity of Renilla luciferase encoded in the vectors under a constitutive promotor.

**Cell adhesion, invasion, migration**

For adhesion assays, CRC cells were serum-starved for 3 h, detached with EDTA/PBS, and labeled with BCECF-AM (Molecular Probes). Then, $6 \times 10^5$ cells in 100 μL were added to 96-well plates previously coated with 4 μL/mL Matrigel (Corning), and the plates were incubated for 25-30 min at 37 °C. Subsequently, non-adhered cells were removed by three washes with serum-free DMEM. Bound cells were lysed with 1% SDS in PBS, and the extent of the adhesion was quantified using POLARstar Galaxy (BMG LABTECH). For invasion, $6\times10^4$ cells were seeded onto 8 μm pore-size Transwell inserts (Corning, NY, USA) previously coated with 35 μL of Matrigel diluted 1:3 in DMEM. The lower chamber of the invasion assay was filled with 700 μL of DMEM supplemented or not with serum (5 %). After 48 h, invasive cells were fixed with 4 % paraformaldehyde (Merck), stained with Crystal Violet (0.1 %, Merck) and counted under a microscope. For migration or wound healing assay, 1-mm incision was made in cell monolayers incubated in medium containing 0.5% serum in 24-well plates, which were previously coated with 0.5 μL/mL matrigel. Pictures were taken immediately after the incision and after 16-24 h. Effective cell migration speed was calculated as the distance (in each flank) covered by the cells in 16-24 h.

**Annexin V-PI Apoptosis Assay**

Cell death was assessed using the FITC-Annexin V Apoptosis Detection Kit with PI (ANXVKF-200T; Immunostep) according to the manufacturer's instructions. Briefly, 2 x

$10^4$ cells per well were seeded in triplicate in 6-well plates under the indicated experimental conditions. Four days after plating, cells were stained with FITC Annexin V and PI following the manufacturer's protocol, and analyzed by flow cytometry with a Cytoflex-S flow cytometer (Beckman-Coulter, Brea, CA, USA).

**Clonogenic assay and spheroid formation**

For clonogenic assays, $7\times10^2$ scrambled and shNPC2 KD cells were seeded into a 6-well plate and cultured in DMEM media with 10% FBS. After 8-10 days, colonies were fixed in 4% PFA, stained with 0.5% Crystal violet, imaged, and quantified by ImageJ software. For alternative quantification, colonies were discoloured with 10% acetic acid and optic density was measured at $A_{600}$ nm. For spheroid formation assays, $6\text{x}10^3$ cells were seeded per well in ultra-low attachment 96-well plates (Corning, NY, USA) containing 100 µL of DMEM supplemented with 10% FBS. Spheroid growth was monitored for 4–5 days by acquiring bright-field images every day using IncuCyte™ S3 Live-Cell analysis system (Sartorius, Göttingen, Germany).

**RNA extraction from cell lines, cDNA synthesis and PCR analyses**

Total RNA was isolated from cellular cultures using TRIzol Reagent (Ambion, Austin, TX, USA) in a ratio of 1mL per $5\text{x}10^6$ cells. The mixture was incubated for 5 min at RT, after which 200 µL of chloroform was added and incubated again for 3 min at room temperature. The mixture was centrifuged at 12,000 x g for 15 min at 4 ° C, the aqueous fraction was collected and 1 vol of 70% ethanol was added. Then, the RNA was purified with the RneasyMini kit (Qiagen). First-strand cDNA synthesis was performed starting from 1 µg of total RNA using the Superscript III first-strand synthesis system kit (Thermo Fisher Scientific) with oligodT primers according to the manufacturer's instructions. The resulting cDNA was subsequently used for quantitative real-time PCR analysis (Bio-Rad, Hercules, CA, USA). GAPDH was amplified as loading control. PCR profile was 5 min at 95 °C, 40 step cycles (10s at 95 °C, 30s at 57 °C) followed by a melt curve from 57 °C to 95 °C for 380s. Relative gene expression was quantified using the 2−ΔΔCt method, with GAPDH as the endogenous normalization control. Primer sequences are listed in **Supp. Table S4**. For semi-quantitative PCR of mouse liver homing assays, the profile was: 5 min at 95 °C, 35 three-step cycles (30s at 94 °C; 30s at 57 °C and 30s at 72 °C) and 5 min at 72 °C. Mouse *Actb* gene was amplified as loading control.

**Transcriptomic profiling and differential expression**

Gene expression profiling of NPC2 (KD) and scrambled control (SCR) cells was performed for KM12SM and HCT116, using microarray-based transcriptomics (three biological replicates per condition) at the Genomics Unit of the Universidad Complutense (Madrid, Spain) using GeneChip® WT PLUS Reagent Kit (Applied Biosystems), hybridized with "Clariom™ S Array, human" (Applied Biosystems) and scanned with a "GeneChip® Scanner 3000 7G" (Applied Biosystems). Raw hybridization intensities were normalized and $log_2$-transformed by the genomics facility; probe-level expression values, gene symbol annotations, fold changes, and FDR-adjusted p-values were provided in tabular format. For each cell line, differential expression between NPC2-KD and SCR conditions was assessed by computing $log_2$ FC as the difference between mean $log_2$ expression of KD and SCR replicates. Multiple-testing correction was applied using the Benjamini–Hochberg procedure (FDR). Probes mapping to the same gene symbol were collapsed to a single entry per gene. Genes with FDR < 0.05 were classified as differentially expressed genes (DEGs). All analyses were performed in Python (pandas, numpy, scipy.stats, matplotlib) and/or R. Figures were generated using Python (matplotlib v.X, seaborn v.X) / R (ggplot2 v.X).

**Gene Set Enrichment Analysis (GSEA)**

Genome-wide GSEA was performed independently for each cell line using ranked gene lists ordered by $log_2$ FC. Gene sets from the MSigDB (Molecular Signatures Database) were evaluated, including the Hallmark collection (50 gene sets), KEGG pathway database, Reactome pathway database, and WikiPathways. Normalized Enrichment Scores (NES) and FDR-adjusted p-values were computed for each gene set in both cell lines. Enriched pathways were considered significant at FDR<0.05. Cross-line comparison of GSEA results was performed by examining concordant (same NES direction) versus discordant (opposite NES direction) pathways, with particular attention to pathways showing significant enrichment in one line and significant enrichment in the opposite direction in the other. Bioinformatic analysis was carried out in collaboration with EGO Genomics (Salamanca, Spain). Gene functional annotation and pathway enrichment analyses were performed in R (version 4.1.2) using the clusterProfiler (version 4.2.2) and rrvgo (version 1.6.0) packages, the latter used to reduce redundancy among enriched Gene Ontology terms. Genome-wide GSEA was performed independently for each cell line using ranked gene lists ordered by log2 FC, the MSigDB Hallmark collection (50 gene sets) and default parameters. Normalized Enrichment

Scores (NES) and FDR-adjusted p-values were computed for each gene set in both cell lines, and enriched pathways were considered significant at FDR < 0.05. In addition, the KEGG, Reactome and WikiPathways databases were evaluated as complementary pathway sources. Cross-line comparison of enrichment results was performed by examining concordant (same NES direction) versus discordant (opposite NES direction) pathways, with particular attention to pathways showing significant enrichment in one line and significant enrichment in the opposite direction in the other.

**Custom curated gene set analysis**

In addition to genome-wide GSEA, hypothesis-driven gene sets were manually curated to interrogate specific biological processes relevant to NPC2 function and cholesterol-dependent signaling. These included: (i) cholesterol synthesis and transport genes (20 genes spanning the mevalonate pathway, lanosterol-to-cholesterol conversion, LDLR/NPC1/NPC2 egress machinery, and ABCA1/SOAT1 efflux/storage); (ii) epithelial–mesenchymal transition (EMT) transcription factors and effectors (SNAI1/2, ZEB1/2, TWIST1, VIM, CDH1, SERPINE1); (iii) intestinal stemness markers (ALDH1A1, SOX2, SOX9, LGR5, LRIG1, PROM1, LEF1, TCF7, CD44, KLF4); (iv) AKT/mTOR pathway components (AKT1–3, MTOR, RPTOR, RICTOR, TSC1/2, RRAGA–D, LAMTOR1–5, SESN1–3, ULK1, SQSTM1, BECN1, ATG5, TFEB); (v) AP-1 and YAP/TAZ pathway components (141 genes covering core AP-1 factors, YAP/TAZ effectors and targets, Hippo kinases, mechanotransduction/focal adhesion, ECM/hypoxia, and EGFR axis); and (vi) lipid raft–associated signaling molecules (CAV1, ROR2, LRP6, IGF1R, PTEN, PTCH1, EGFR, FZD1, AKT1, YES1, NPC1). For each curated set, per-gene log2FC and FDR values were extracted for both cell lines, and the fraction of genes changing concordantly with the expected direction, the median log2FC, and the number of significant DEGs (up vs. down) were reported. Computational analyses were conducted with assistance from Biomni (Phylo) (1), an AI-based research analysis platform; all analyses were reviewed and validated by the authors.

**Signaling assays**

For signaling assays, cells were kept in starving conditions for 3h, detached with 2 mM EDTA and incubated with 5 % serum for 15 min, 10 ng/mL IL-13 (Peprotech, London, United Kingdom) for 12 min or 20 ng/mL EGF for 5 min (Peprotech). Cells were lysed in GST-FISH buffer, and the extracts analyzed by western blotting. For mTOR analyses, cells were incubated 18 h with serum (10%) or DLPS (10%) in the presence or absence of mTOR inhibitors, 1 μM Rapamycin (MedChemExpress, Sweden) or 2.5 μM Torin-1 (MedChemExpress), before lysis.

**Western blot**

For Western blot, cells were lysed in buffer containing 1% Igepal (Merck), 100 mM NaCl, 2 mM $MgCl_2$, 10% glycerol, and 50 mM Tris-HCl, supplemented with protease inhibitors (Roche, Basel, Switzerland) and phosphatase inhibitors (Sigma-Aldrich, St. Louis, MO, USA) and 50 μg of total protein were resolved in SDS-PAGE and transferred onto nitrocellulose membranes. Blots were incubated with primary antibodies for 16 h at 4°C followed by incubation with HRP-conjugated secondary antibodies for 2 h (diluted 1:2000, Thermo Fisher Scientific). Bands were visualized with SuperSignal West Pico Chemiluminescent Substrate (Thermo Fisher Scientific) in the Chemidoc Imaging System device (Bio-Rad) and quantified using MultiGauge software (Fujifilm, Santa Ana, CA, USA).

**Lipidomics**

Lipids were extracted according to a modified Bligh and Dyer protocol. 200 µL of water was added to the cell pellets. After vortexing (30s), the sample was transferred to a glass tube. We added to the samples 4 μL SPLASH LIPIDOMIX Mass Spec Standard + 2 μL SphingoSPLASH I (all from Avanti Polar Lipids). 500 µl of methanol was used to rinse the first tube and then transferred to a glass tube to which 250 µl of chloroform was finally added. The mixture was vortexed for 10 min before a new addition of 250 µl of water and 250 µl of chloroform. The mixture was vortexed again for 10 min and finally centrifuged (3000 rpm, 4°C, 15 minutes). Then, 400 µL of the organic phase was collected in a new glass tube and dried under a stream of nitrogen. The dried extract was resuspended in 60 µL of isopropanol/methanol/water 50/30/20 and transferred in an injection vial. Lipidomic profiling was performed using reverse-phase ultra-performance liquid chromatography (RP-UPLC) coupled with High Resolution mass spectrometry. Reverse phase liquid chromatography was selected for separation with an UPLC system (Ultimate 3000, Thermo Fisher). Lipid extracts were separated on an Accucore C18 (150x2.1, 2.5µm column) (Thermo Fisher). The column was maintained at 35°C and the mobile phase was delivered at a flow rate of 400 µl/min. The injection volume was set to 3 µL. The mobile phases consisted of (A) Acetonitrile/Water 1/1 (V/V) containing 10mM ammonium formate and 0.1% formic acid and (B) Isopropanol/Acetonitrile/Water 88/10/2 (V/V) containing 2mM ammonium formate and 0.02% formic acid. The gradient was: 35% mobile phase B at 0 minutes, 60%B at 4 minutes, 70% B at 8 minutes, 85% B at 16 minutes, 97% B at 25 minutes, 100% B at 25,1 minutes until 31 minutes, and back to 35% B for 4 more minutes. The UPLC system was coupled with a Q-exactive Orbitrap Mass Spectrometer (Thermo Fisher Scientific) equipped with a heated electrospray ionization (HESI) source. Spectra were acquired in data-dependent MS2 (dd-MS2) mode with top 15 ions being subjected to MS2 analyses, in positive and negative ion modes. MS1 resolution was 70 000 (at m/z 200) over a mass range of m/z 250-1200. The automatic gain control (AGC) target was set to $1 \times 10^6$ with a maximum injection time of 100 ms. Precursor isolation was performed with a 1.0 m/z isolation window, and fragmentation was achieved using higher-energy collisional dissociation (HCD) at normalized collision energy (NCE) of 25 and 30 eV. MS/MS spectra were acquired in the ion trap at a resolution of 35,000 (at m/z 200) with an AGC target of $1 \times 10^5$ and a maximum injection time of 80 ms. Raw data were reprocessed using LipidSearch 4.2.21 (Thermo Fisher Scientific). Lipid identification was performed in product search mode, based on accurate mass and MS/MS fragmentation patterns. The following parameters were applied: precursor ions mass tolerance of 5ppm, product tolerance 5 ppm; ion and adducts searched [H+], [NH4+] and [Na+]; alignment retention time tolerance 0.4 min; ID quality filters A, B and C.

Untargeted LC–MS lipidomics was performed in SCR and NPC2-knockdown cells from both cell lines using four biological replicates per condition. Lipids were analyzed in positive- and negative-ion modes. SPLASH-type class-specific internal standards were used for normalization when available; otherwise, median/total-signal normalization was applied. Features with a QC-pool coefficient of variation >30% were excluded, and one failed positive-mode injection was removed. Technical replicate injections were averaged prior to statistical analysis. Because the cholesteryl-ester internal standard showed relatively high variability, cholesterol-related lipids were additionally analyzed using median normalization as a sensitivity analysis, yielding highly concordant results. Differential abundance was assessed for each lipid species using Welch's *t*-test comparing NPC2-KD with SCR cells within each cell line. Fold changes were expressed as $\log_2$(KD/SCR), and *P* values were corrected for multiple testing using the Benjamini–Hochberg FDR procedure. Significant changes were defined as adjusted $P<0.05$ and $|\log2FC|\geq0.585$ (1.5-fold). Lipid-class enrichment analyses (ORA and LSEA) were performed by the lipidomics facility using LIPID MAPS tools. As untargeted lipidomics provides relative rather than absolute quantification, lipid-class trends were prioritized over individual low-abundance species. Additional statistical tests included Pearson correlation for cross-line log2FC comparisons, Mann–Whitney U test for comparing total double-bond counts between up- and downregulated lipid species, and paired Wilcoxon signed-rank test for comparing within-array percentile ranks of cholesterol synthesis enzymes between cell lines at baseline.

**Bioinformatic lipidomic analysis**

Bioinformatic analysis of the lipidomic profiling was performed using R (4.4.3). Technical replicates of positive analysis were averaged prior to downstream analyses in order to be able to merge the positive and negative data. To apply QC filtering on the LC-MS/MS peak areas, we selected lipids for which the mean of samples' intensity was greater than 2x compared to the blank mean (2). QC samples were used to calculate the coefficient of variation (CV) for each lipid. Lipids showing a CV greater than 30% across QC samples were excluded from further analyses. Following QC filtering, low-intensity values below a signal threshold of 10,000 were treated as missing values. Remaining missing values were imputed using half of the minimum detected peak area in the dataset. Next, the probabilistic quotient normalization (PQN) method (3) was performed for data normalization, using the "Control" samples as reference. The final normalized values were log2 transformed. For computation of the z-scores, each lipid was centered by the reference samples' mean and scaled by the reference samples' standard deviation. For the differential analysis, t-tests were used to compare peaks between groups and fold changes were calculated for each lipid. A cutoff threshold of 0.05 for p-value and fold change of 1.5 was used to select the most significant lipids. The Benjamini-Hochberg method was used to correct p-value for multiple testing. Hierarchical clustering was performed on the log transformed normalized data (mean centered and scaled by standard deviation). Clustering was performed using the Ward method and Pearson's correlation was used for distance calculation.

**Membrane lipid order imaging**

Membrane lipid order was assessed in SCR and NPC2-silenced KM12SM and HCT116 cells using LipiORDER (FDV-0041; Funakoshi Co., Tokyo, Japan) (4), a pyrene-based solvatochromic dye whose emission shifts from green to red as membrane lipid packing decreases. Cells were seeded on ibidi 8-well µ-Slides (ibidi GmbH, Gräfelfing, Germany), grown to reach 70% confluence, and incubated with 300 nM LipiORDER in serum-free, phenol red-free DMEM for 10 min at 37 °C. Ratiometric imaging was performed on a Leica SP8 confocal laser scanning microscope (Leica Microsystems, Wetzlar, Germany) with 405 nm excitation, collecting emission simultaneously in the green (500–550 nm; ordered, Lo membranes) and red (550–650 nm; disordered, Ld membranes) channels. R/G images were generated, and the per-cell R/G ratio was quantified in the plasma membrane (cortex) and interior compartments from segmented images. Image segmentation and quantification were performed using Fiji/ImageJ and Biomni (1), following established ratiometric analysis procedures. Images containing at least 100 cells per condition from three independent cultures were analyzed.

**NPC disease hallmark scoring**

To assess whether NPC2 depletion recapitulates cellular features of Niemann–Pick type C (NPC) disease, nine predefined hallmarks were evaluated against the transcriptomic and lipidomic data: (H1) lysosomal free-cholesterol retention, (H2) reduced cholesterol esterification, (H3) glycosphingolipid storage, (H4) sphingomyelin/sphingosine storage, (H5) mitochondrial dysfunction, (H6) SREBP2-driven cholesterol synthesis upregulation, (H7a) Rag-dependent mTORC1 nutrient sensing, (H7b) autophagy/TFEB dysregulation, (H8) oxidative stress/Nrf2 activation, and (H9) hepatosplenomegaly. Each hallmark was scored as MATCH, PARTIAL-MATCH, DIVERGENT, or NOT-TESTABLE based on the direction and magnitude of gene/lipid changes relative to the NPC disease expectation, using the curated gene sets and lipid class summaries described above. Seven of the nine hallmarks were measurable in cell culture; H1 (lysosomal free-cholesterol retention) and H9 (hepatosplenomegaly) require subcellular localization or in vivo readouts unavailable in this system. Comparative analysis was performed with Biomni.

**In vivo experiments**

The Ethical Committees of the CSIC and Community of Madrid approved the protocols used for the experimental work with animals (PROEX 206.4/25). In all cases, we followed the standard calculation of sample size in animal studies, employing the formula $n = 2\sigma^2 \cdot (Z_{\alpha/2}+Z_\beta)^2/d^2$ to calculate the number of mice for the assays. For homing experiments, mice were inoculated in the spleen, with $1{\times}10^6$ scrambled and NPC2-silenced KM12SM and HCT116 cells, and euthanized 48 h after inoculation. After that, RNA was isolated from the liver using Trizol purification and subjected to RT-PCR with M-MLV retrotranscriptase (Promega, Madison, WI, USA) and Taq DNA polymerase (Thermo Fisher Scientific) to amplify human GAPDH and, as a control, murine β-actin. For Kaplan–Meier survival curves, both CRCs Scrambled or NPC2-silenced cells were inoculated in NSG mice (n = 6). 24 h after inoculation, mice were subjected to splenectomy to avoid local growth of tumors. Mice were monitored daily for signs of disease including reduced mobility, loss of appetite, tumor detection by palpation, self-isolation, dermatitis, etc. After relevant sign detection, mice were euthanized and subjected to necropsy to verify the presence of liver metastases. For xenograft studies, 5 x $10^6$ cells were inoculated subcutaneously with 0.1% glucose in PBS in both flanks of 3 NSG mice. Tumor volumes were measured twice per week for 3-4 weeks.

**In silico prognostic analysis**

To evaluate the prognostic value of NPC2 across different solid tumors, independent external patient cohorts were analyzed. Prognostic analyses were performed using the GSE39582 dataset, which includes 585 colorectal cancer samples classified according to Colorectal Cancer Intrinsic Subtypes (CRIS). NPC2 mRNA expression values were normalized by calculating z-scores, and patients were stratified into high- and low-expression groups using the optimal cut-off determined by the best cut-off method. Overall survival was assessed using the Kaplan-Meier method and compared with the log-rank test. Associations between NPC2 expression, mutational status, and molecular subtypes were also evaluated using the GSE39582 cohort. In addition, publicly available web-based platforms were used for independent validation. GEPIA2 (http://gepia2.cancer-pku.cn) was employed to analyze transcriptomic data from colon adenocarcinoma (COAD, n = 308), while KM Plotter (https://kmplot.com/analysis/) was used to assess the prognostic significance of NPC2 expression in colorectal cancer (n=513), including subgroup analyses according to mutational status.

**References**

1. Huang K, Zhang S, Wang H, Qu Y, Lu Y, Li R, et al. Autonomous biomedical research with an artificial intelligence agent. Science. 2026:eadz4351.
2. Smirnov D, Mazin P, Osetrova M, Stekolshchikova E, Khrameeva E. The Hitchhiker's Guide to Untargeted Lipidomics Analysis: Practical Guidelines. Metabolites. 2021;11(11).

3. Dieterle F, Ross A, Schlotterbeck G, Senn H. Probabilistic quotient normalization as robust method to account for dilution of complex biological mixtures. Application in 1H NMR metabonomics. Anal Chem. 2006;78(13):4281-90.
4. Valanciunaite J, Kempf E, Seki H, Danylchuk DI, Peyrieras N, Niko Y, et al. Polarity Mapping of Cells and Embryos by Improved Fluorescent Solvatochromic Pyrene Probe. Anal Chem. 2020;92(9):6512-20.

| Table S4. Primers sequences for validations | | |
|---|---|---|
| **Gene** | **FW** | **RV** |
| NPC2 | TCCTGGCAGCTACATTCCTG | TGGTGAAGGTGACATTGACG |
| LDLR | CAGGAGACATCCACCGTCAG | CGAGGAGCACTTGGTGAGAC |
| NPC1 | ATTACCACACGCACTTTCTG | TCATAGCCTCCCAACACAAG |
| SOAT1 | AAGTTTGCACAGGTCTTTGG | ACATAGGACCAGAACACGAG |
| ABCA1 | TGAAAGGGCTCTCTGAGAAG | CTTTCTCTGCATTCCACCTG |
| HMGCR | TAAGCCGAATCCTGTAACTC | CATCCAGTCCTAATGAAACC |
| SREBP2 | CAGATTCCCTTGTTTTGACC | CATCATTACAGGCATTGTGG |
| KLF4 | ACCAAGAGCTCATGCCACCC | GTGTAGGTTTTGCCGCAGCC |
| SOX2 | GAAAAACGAGGGAAATGGG | GGATGGGATTGGTGTTCTC |
| SOX9 | CCAGCAAGAACAAGCCGCAC | AGTCTCCAGAGCTTGCCCAG |
| CD44 | GCCTGGCGCAGATCGATTTG | TCCATCTGGGCCATTGTGGG |
| CD133 | GACTGAGACCCAACATCATC | CTGGTCAGACTGCTGCTAAG |
| ALDH1A1 | TGACCCTGGAGCTTGGAGGA | CAACACTGGCCCTGGTGGTA |
| LGR4 | TGTCTGGGTTGAAAGAACTC | GCATCTAAACGCAAAGACTG |
| CDH1 | CTGACACACCCCCTGTTGGT | CCGTTGGATGACACAGCGTG |
| SNAI2 | TCATACCACAACCAGAGATC | CCCAAAGATGAGGAGTATCC |
| ZEB1 | GCACAAGAAGAGCCACAAGTA | GCAAGACAAGTTCAAGGGTTC |
| VIM | TGCGCCTCCGGGAGAAATTG | TCAAGACGTGCCAGAGACGC |
| FOS1 | GGTGGAACAGTTATCTCCAG | CTTGGAGTGTATCAGTCAGC |
| JUN | AGCATGACCCTGAACCTGGC | TTATCAGGCGCTCCAGCTCG |
