## Supplementary Figures for "A cholesterol addiction creates an Achilles’ heel in *APC*-truncated colorectal cancer"

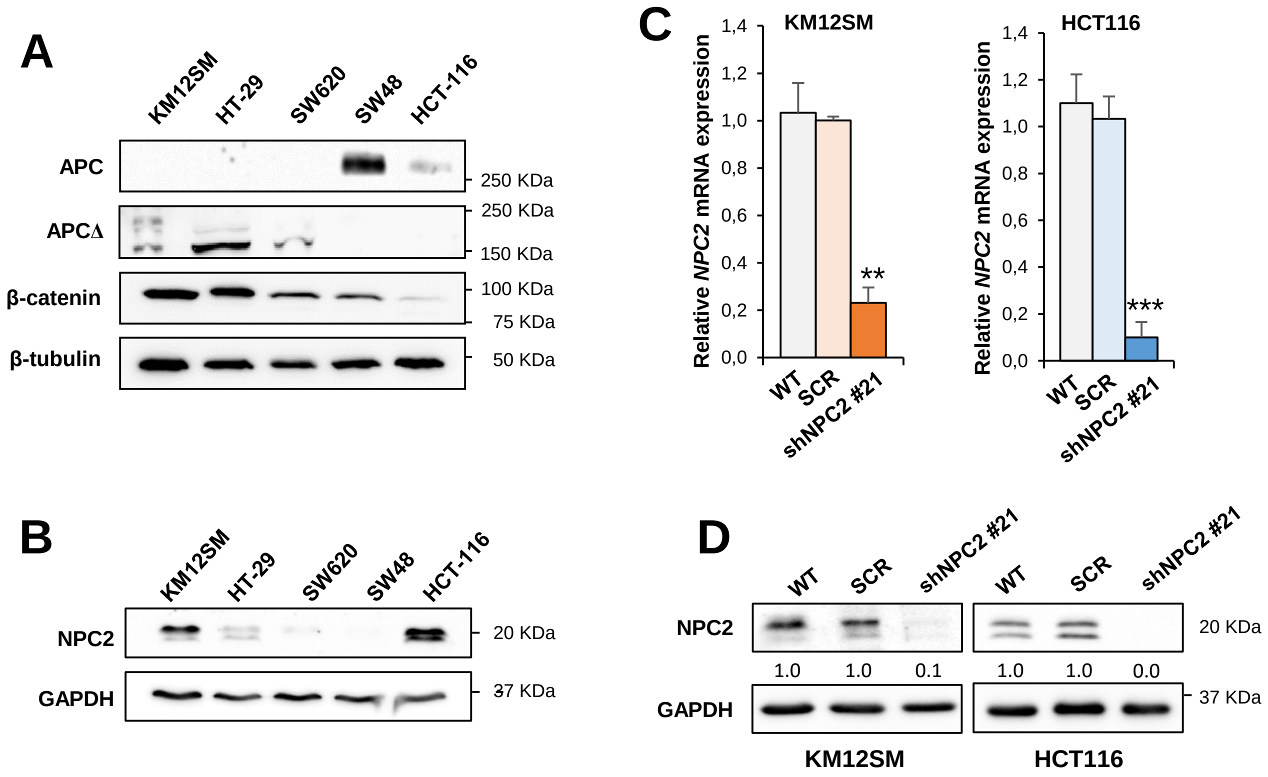


**Figure S1. Expression of APC, β-catenin and NPC2 and validation of NPC2 KD in CRC cell lines.** (A, B) The indicated cell lines were lysed and the extracts analyzed by western blot for APC and β-catenin (A) or total NPC2 (B). β-tubulin or GAPDH were used as a loading control. (C) The indicated cell lines were transduced with lentivirus expressing either a scrambled shRNA (SCR) or an NPC2-targeting shRNA (shNPC2), and selected with puromycin for 2 weeks. NPC2 mRNA levels in wild-type (WT), SCR and shNPC2 cells were quantified by RT-qPCR. NPC2 expression was significantly reduced in shNPC2 cells (**, $p < 0.01$; ***, $p < 0.001$). (D) Protein extracts from the same cells were analyzed by western blot to confirm NPC2 depletion. GAPDH was used as a loading control. Band quantification is shown below the corresponding lanes.

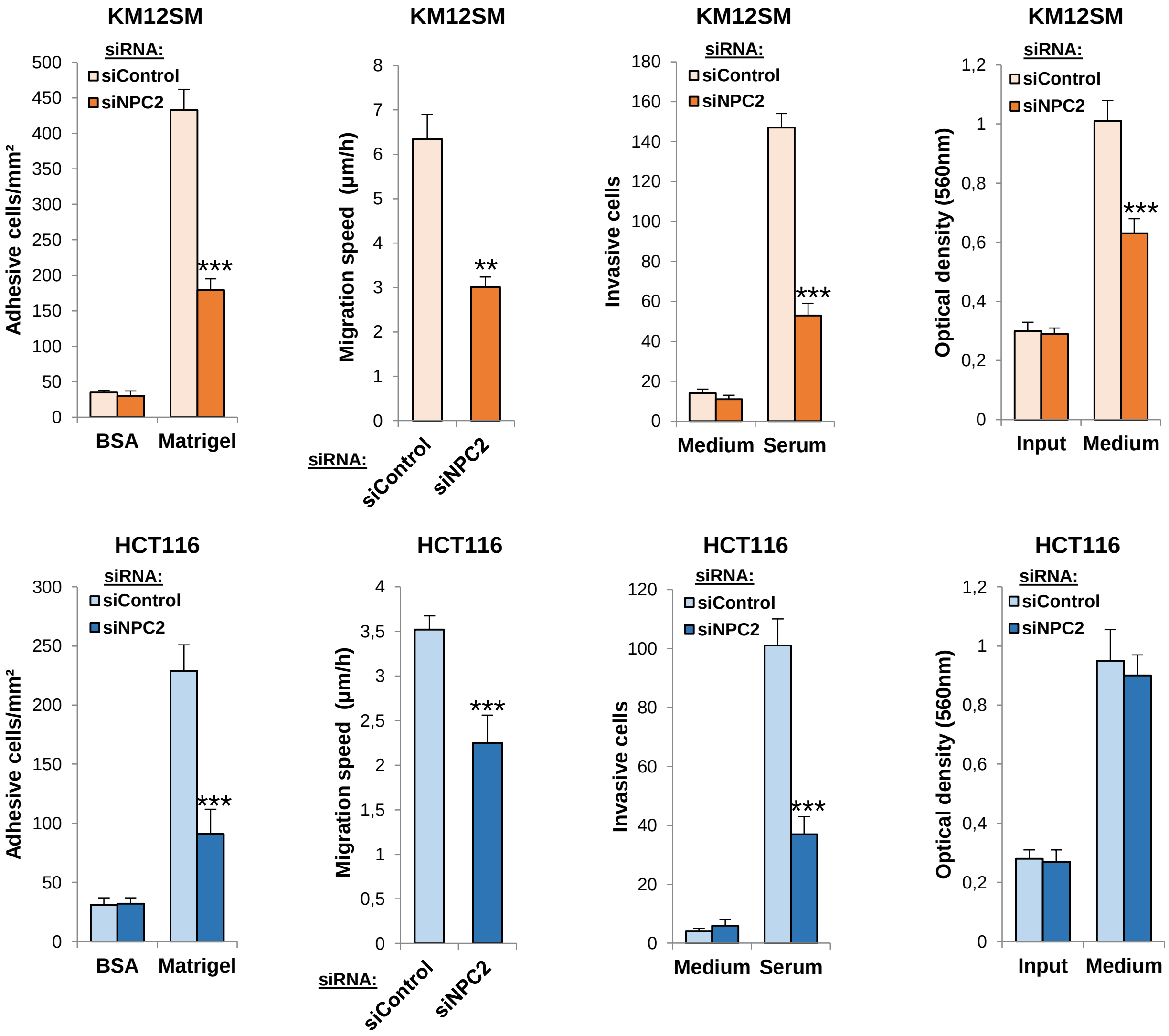


**Figure S2. NPC2 silencing reduces adhesion, migration and invasion in colorectal cancer cells.** The indicated parental cells and stable transfectants were analyzed using Matrigel adhesion assays, wound-healing assays, Matrigel invasion assays toward serum, and MTT proliferation assays. NPC2 depletion significantly reduced adhesion, migration and invasion, with effects on proliferation depending on the cellular model (**, p < 0.01; ***, p < 0.001).

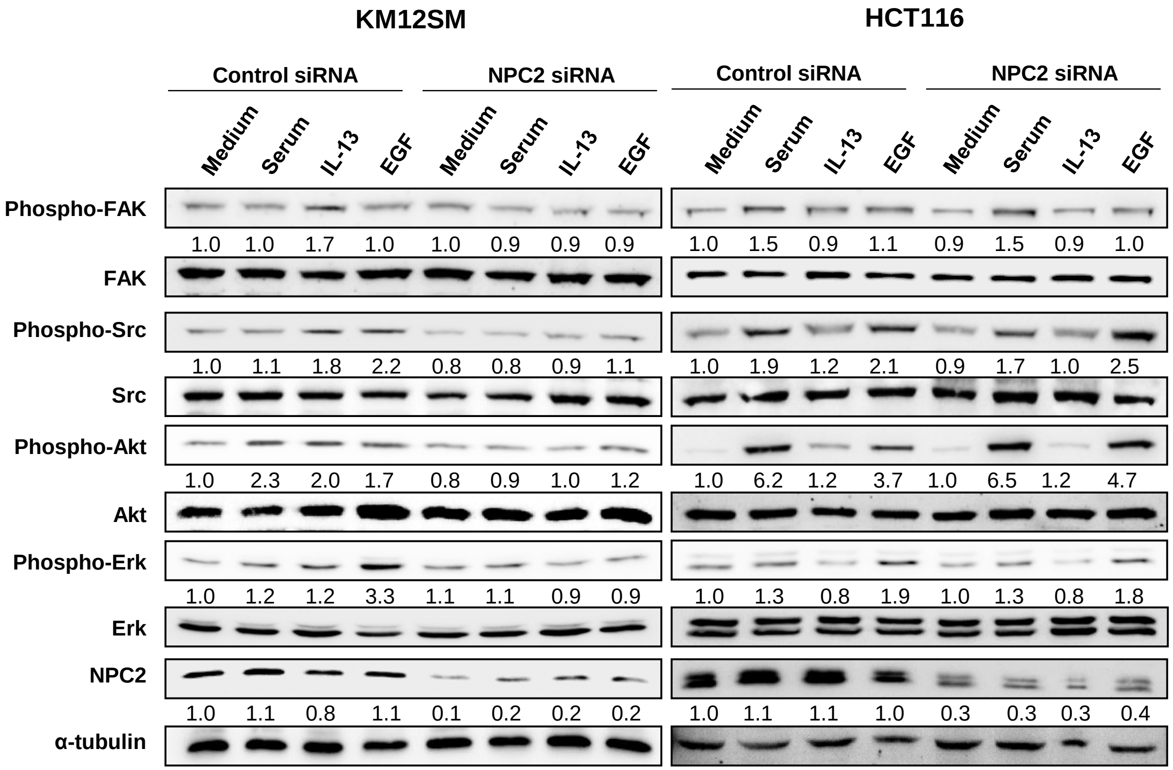


**Figure S3. NPC2 silencing attenuates cytokine/growth factor-induced signaling in KM12SM cells.** KM12SM cells were transfected with control or NPC2-targeting siRNAs. After 48 h, cells were serum-starved and subsequently stimulated with serum (5%, 15 min), IL-13 (10 ng/mL, 12 min), or EGF (20 ng/mL, 5 min). Cell lysates were analyzed by western blotting for the indicated total and phosphorylated proteins, with α-tubulin used as loading control. Quantification of phosphoprotein levels normalized to the corresponding total protein is shown below each lane.

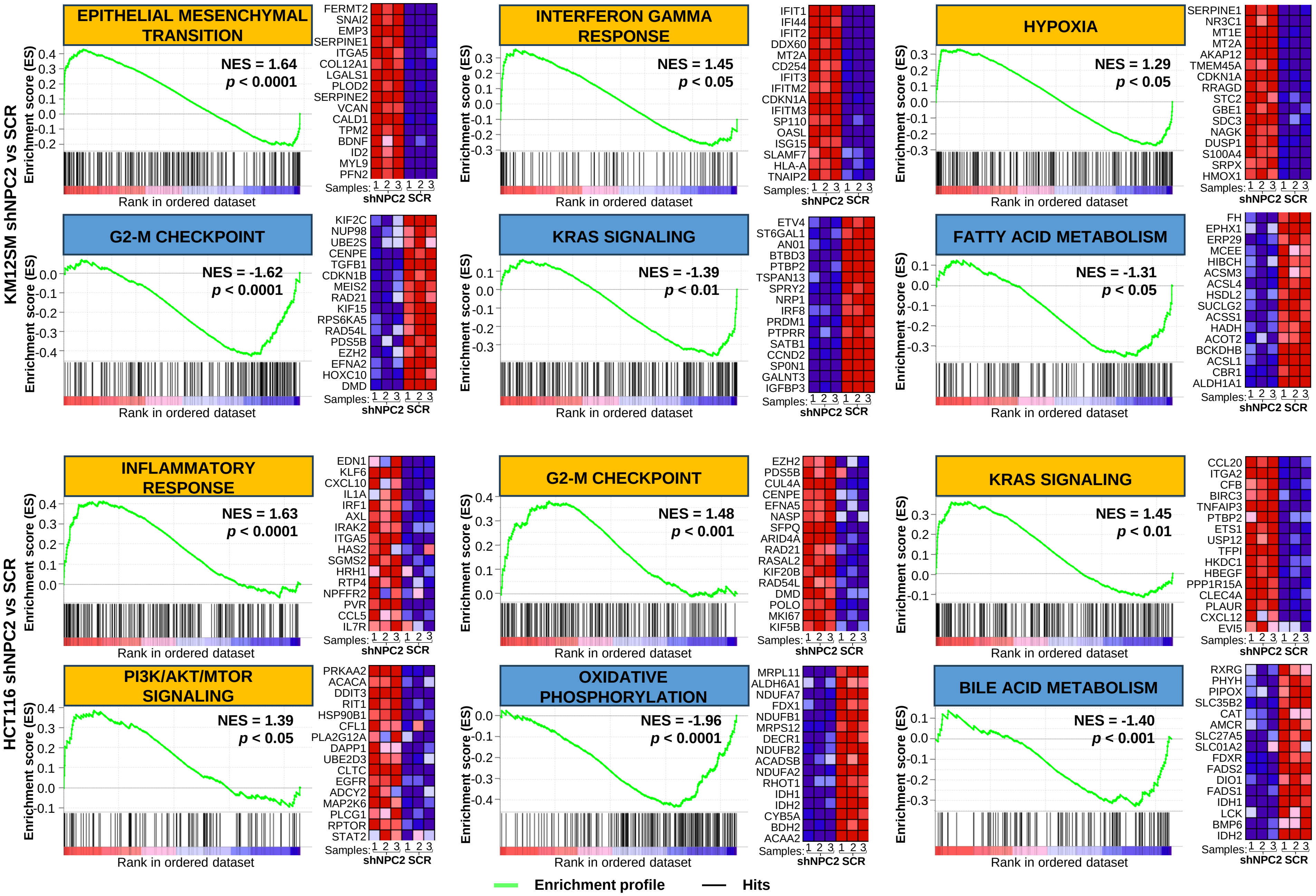


**Figure S4. Hallmark pathway enrichment following NPC2 depletion.** Gene set enrichment analysis (GSEA) of Hallmark gene sets significantly altered after NPC2 silencing in the indicated cell lines. Normalized enrichment scores (NES) and false discovery rate (FDR) P values are shown within each panel. Positive NES values indicate enrichment in NPC2-KD cells (orange), whereas negative NES values indicate depletion (blue). Heatmaps on the right show the most differentially expressed genes contributing to each Hallmark signature.

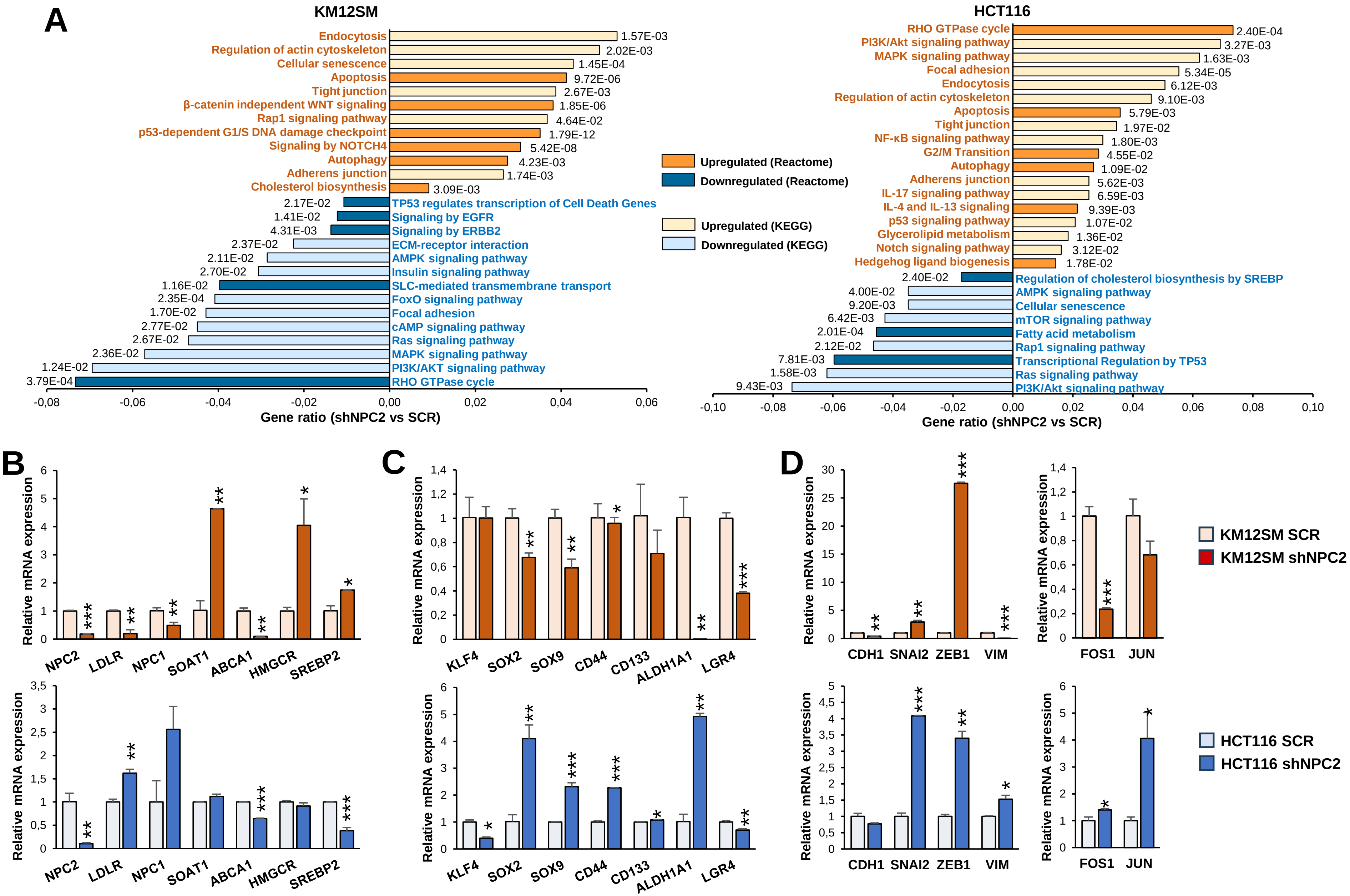


**Figure S5. NPC2 silencing alters gene expression profile in colorectal cancer cells.** (A) Pathways significantly dysregulated in shNPC2 cells according to KEGG and Reactome analyses. FDR is shown near each bar. Quantitative PCR analyses of selected genes involved in cholesterol homeostasis (B), stemness (C), and EMT and AP-1 transcription factor (D). NPC2 depletion significantly modulates the expression of the indicated genes (*, $p < 0.05$; **, $p < 0.01$; ***, $p < 0.001$).

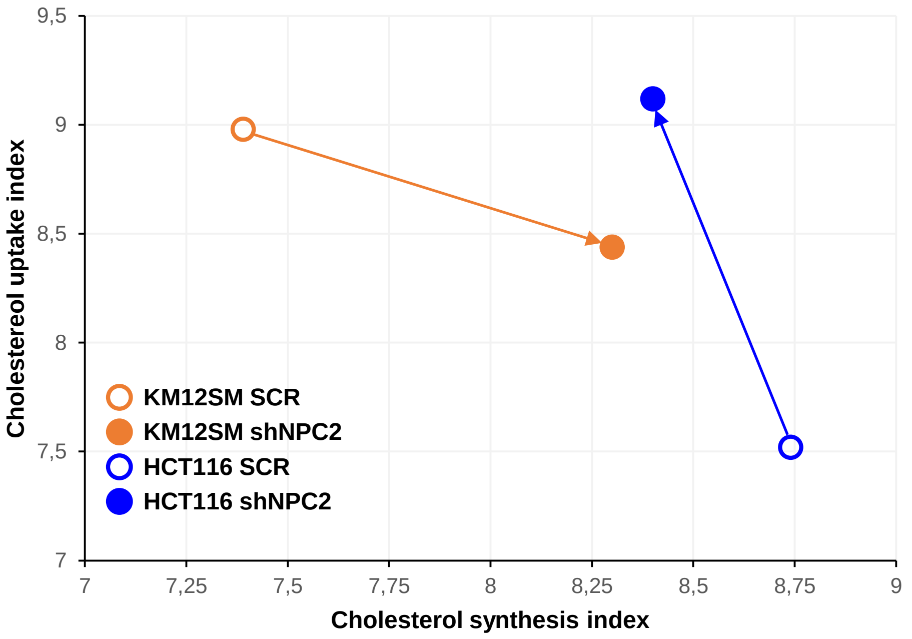


**Figure S6. Differential remodeling of cholesterol synthesis and uptake after NPC2 depletion.** Changes in cholesterol homeostasis are represented as a two-dimensional trajectory from scrambled control (SCR) to NPC2-silenced (shNPC2) cells. The cholesterol uptake index was calculated as the mean log2 expression intensity of LDLR and NPC1, whereas the cholesterol synthesis index was calculated from the mean $log_2$ expression intensity of HMGCR, HMGCS1, MVK, PMVK, MVD, SQLE, IDI1, FDPS, and FDFT1. HCT116 cells shifted predominantly toward increased cholesterol uptake (uptake index, +1.60), whereas KM12SM cells shifted toward increased synthesis with reduced uptake (synthesis index, +0.91; uptake index, −0.54), highlighting distinct compensatory responses to NPC2 depletion.

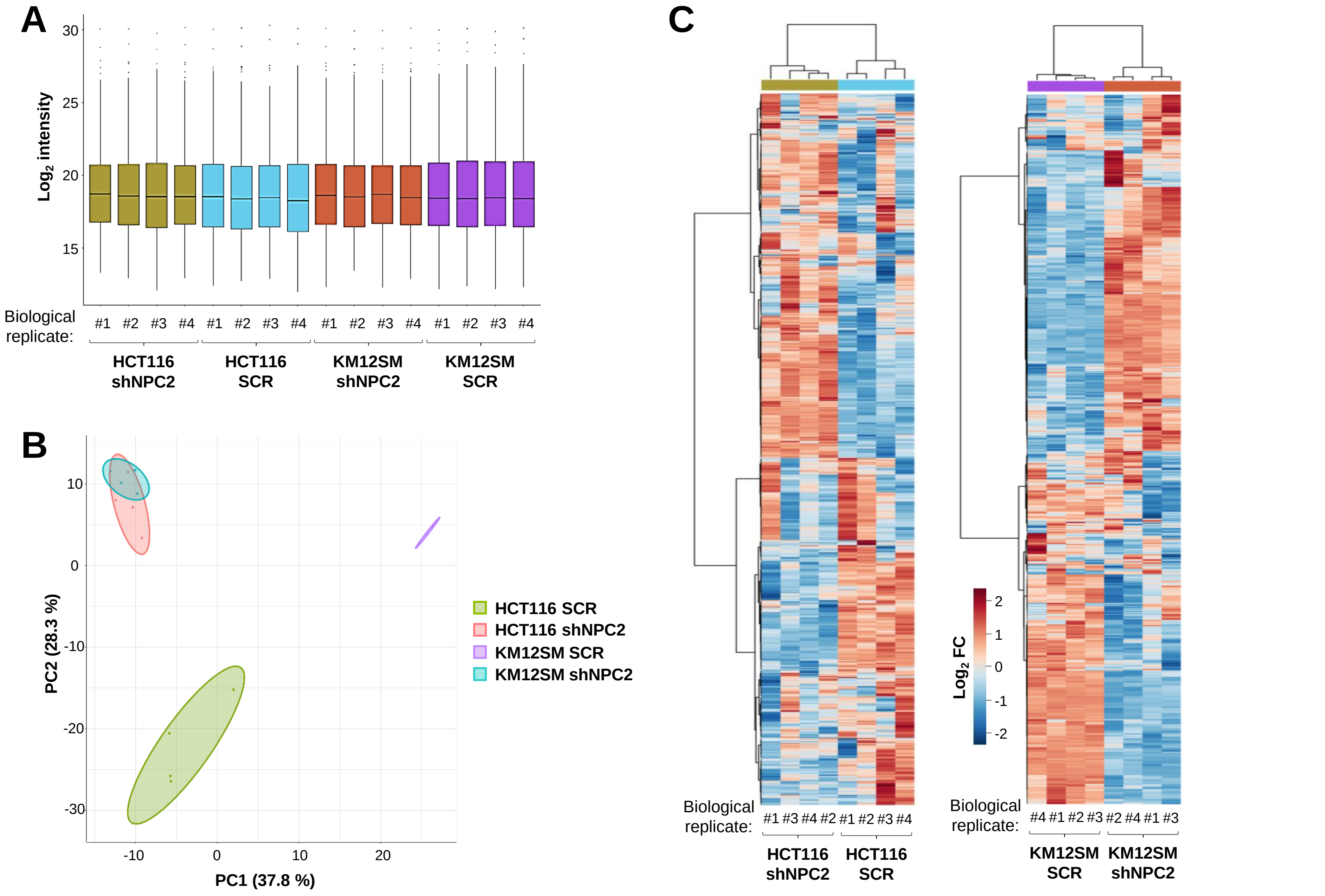


**Figure S7. Normalization, PCA and heatmap analyses of lipidomic profiles.** (A) Distribution of lipid intensities across biological replicates of HCT116 and KM12SM SCR and NPC2-KD cells, shown as $\log_2$-transformed intensity values. (B) Principal component analysis (PCA) of the lipidomic profiles showing separation of HCT116 and KM12SM samples according to cell line and experimental condition. PC1 and PC2 account for 37.8% and 28.3% of the total variance, respectively. (C) Heatmap of lipidomic changes represented as $\log_2$ fold changes (NPC2-KD vs SCR), showing distinct clusters of increased and decreased lipid species in HCT116 and KM12SM cells and highlighting their divergent lipidomic responses to NPC2 depletion.

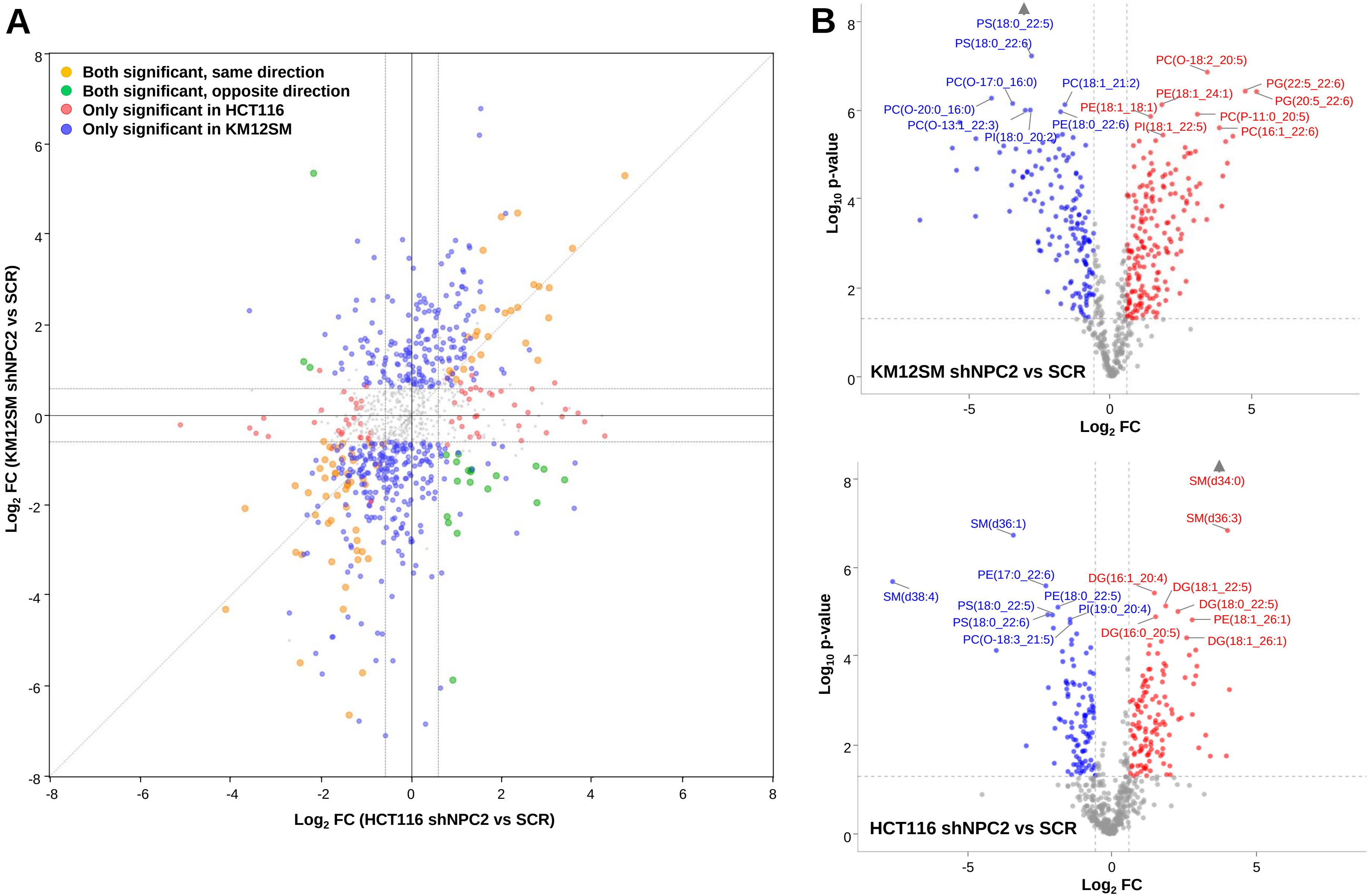


**Figure S8. Differential lipidomic responses to NPC2 depletion in KM12SM and HCT116 cells.** (A) Scatter plot comparing log2 fold changes (NPC2-KD vs SCR) of all detected lipid species in KM12SM and HCT116 cells. KM12SM cells showed a greater number of significantly altered lipid species than HCT116 cells. (B) Volcano plots showing differentially regulated lipid species in KM12SM NPC2-KD versus SCR cells (left) and HCT116 NPC2-KD versus SCR cells (right). KM12SM cells exhibited a broader lipidomic response, with prominent changes in phosphatidylserine (PS) and phosphatidylcholine (PC) species, whereas HCT116 cells showed a more restricted profile, mainly involving sphingomyelin (SM) and diacylglycerol (DG) species. Selected significantly altered lipid species are indicated ($p < 0.05$).

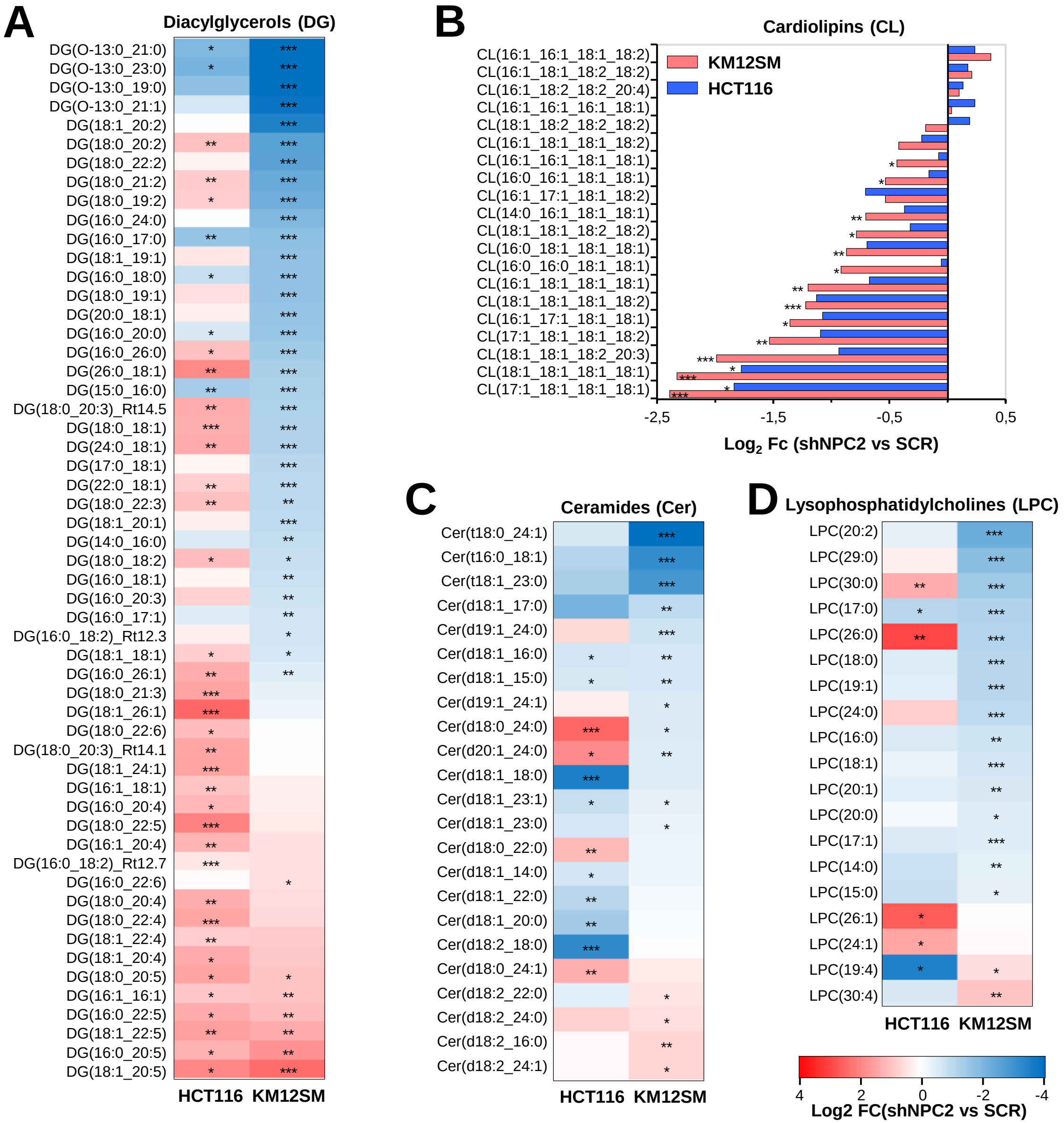


**Figure S9. NPC2 depletion differentially alters diacylglycerol, cardiolipin, ceramide and lysophosphatidylcholine species.** (A) Heatmap showing the $\log_2$ fold changes of detected diacylglycerols in HCT116 and KM12SM NPC2-KD cells compared with SCR controls. (B) Representation of $\log_2$ fold changes of detected cardiolipins in the same cell lines. (C, D) Heatmaps showing the $\log_2$ fold changes of detected ceramides (C) and lysophosphatidylcholines (D) in the same cell lines. The indicated lipid species showed significant changes following NPC2 depletion (*, $p < 0.05$; **, $p < 0.01$; ***, $p < 0.001$).

| Process | NPC2 disease expectation | KM12SM | HCT116 |
|---|---|---|---|
| *Free-cholesterol in lysosomal trap* | ↑ Lysosomal free cholesterol | **MATCH**<br>PM Ch -0.46<br>Lysosome area +0.28 | **MATCH**<br>PM Ch -0.15<br>Lysosome area +0.66 |
| *Reduced esterification* | ↓ Cholesteryl esters | **WEAK MATCH**<br>ChE -0.51 | **WEAK DIVERGENCE**<br>ChE -0.10 |
| *Glycosphingolipid storage* | ↑ Glucosylceramides, Globoceramides | **STRONG MATCH**<br>Gb3Cer +3.25 | **STRONG MATCH**<br>Gb3Cer +2.95 |
| *Sphingomyelin/ Sphingosine storage* | ↑ Sphingosines, sphingomyelins | **DIVERGENCE**<br>SM -1.30 | **PARTIAL DIVERGENCE**<br>SM +0.55 |
| *Mitochondrial dysfunction* | ↓ CoQ, Cardiolipins, ↑ Phosphatidyl-glycerols | **STRONG MATCH**<br>CoQ -2.49<br>CL -1.44<br>PG +3.07 | **WEAK MATCH**<br>CoQ -0.39<br>CL -1.81<br>PG +1.44 |
| *SREBP2 synthesis upregulation* | ↑ Cholesterol synthesis genes | **STRONG MATCH**<br>SQLE +1.48<br>HMGCR +1.41 | **STRONG DIVERGENCE**<br>SREBF2 -0.81<br>LDLR +0.93 |
| *mTORC1 Rag-sensing* | ↑ RRAG, LAMTOR | **MATCH**<br>RRAGD +2.24<br>RRAGC +1.43 | **DIVERGENCE**<br>PRAGD -0.62<br>LAMTOR1 -0.79 |
| *Autophagy / TFEB* | ↑ p62, ATG impaired flux | **PARTIAL MATCH**<br>SQSTM1 +0.99<br>GABARAPL1 +1.37<br>ATG5 +0.91 | **WEAK DIVERGENCE**<br>SQSTM1 +0.90 |
| *Oxidative stress / Nrf2* | ↑ HMOX1, GCLM, Nrf2 | **MATCH**<br>HMOX1 +1.83<br>GCLM +1.82 | **PARTIAL DIVERGENCE**<br>HMOX1 -0.09<br>GCLM +0.26 |
| *Hepatosplenomegaly* | ↑ Liver/spleen foam cells | **NO TESTABLE**<br>Low metastatic ability of shNPC2 cells | **MATCH**<br>Severe hepatomegaly in mice |

- **STRONG MATCH**
- **MATCH**
- **PARTIAL MATCH**
- **WEAK MATCH**
- **WEAK DIVERGENCE**
- **PARTIAL DIVERGENCE**
- **DIVERGENCE**
- **STRONG DIVERGENCE**

**Figure S10. Niemann-Pick disease resemblance scorecard.** Nine characteristic features of Niemann–Pick disease were compared with the phenotypes observed after NPC2 depletion in CRC cells. Accumulation of intracellular cholesterol and globo-series Hex3Cer was observed in both cell lines, whereas increased cholesterol synthesis, mitochondrial dysfunction, oxidative stress and Rag-related alterations were predominantly restricted to KM12SM cells**.**

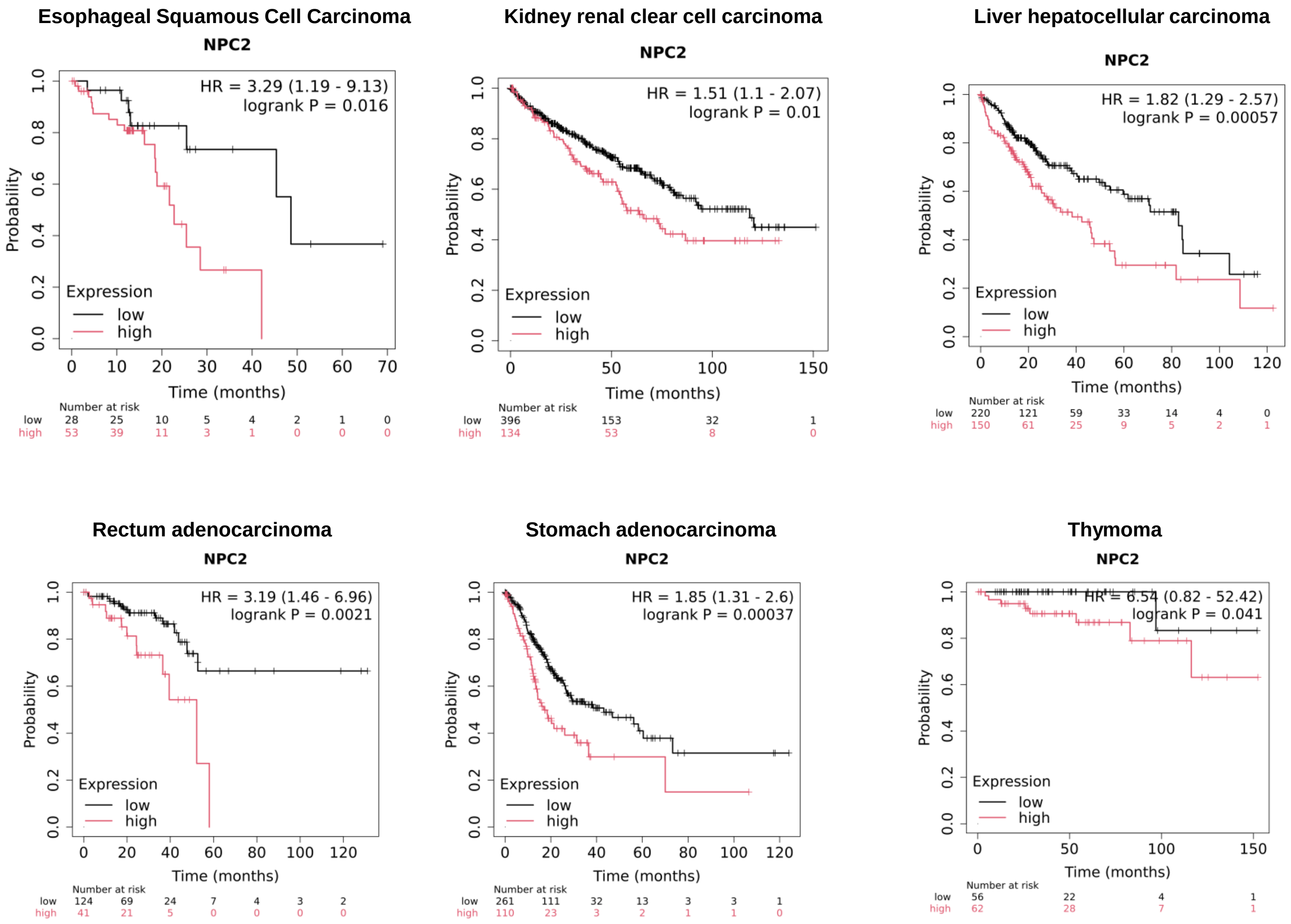


**Figure S11. High NPC2 expression is associated with poor survival across multiple cancer types.** Kaplan–Meier survival analyses of patients with the indicated cancer types stratified according to NPC2 mRNA expression using the Kaplan–Meier Plotter platform. Patients were classified into low- and high-NPC2 expression groups using the best-performing cutoff. Log-rank P values and hazard ratios are indicated in the corresponding plots.
