## Supplementary material for "A cholesterol addiction creates an Achilles’ heel in *APC*-truncated colorectal cancer": Supp Table S3

**Table S3. Lipidomic analysis of shNPC2 vs SCR cells**

| LipidID | HCT116 shNPC2 #1 | HCT116 shNPC2 #2 | HCT116 shNPC2 #3 | HCT116 shNPC2 #4 | HCT116 SCR #1 | HCT116 SCR #2 | HCT116 SCR #3 | HCT116 SCR #4 | KM12SM shNPC2 #1 | KM12SM shNPC2 #2 | KM12SM shNPC2 #3 | KM12SM shNPC2 #4 | KM12SM SCR #1 | KM12SM SCR #2 | KM12SM SCR #3 | KM12SM SCR #4 |
|---|---|---|---|---|---|---|---|---|---|---|---|---|---|---|---|---|
| Cer(d18:0_14:0) | 1.62E+04 | 2.81E+04 | 5.97E+04 | 2.67E+04 | 1.24E+04 | 9.26E+03 | 4.90E+03 | 1.33E+04 | 3.28E+04 | 2.03E+04 | 3.56E+04 | 1.44E+04 | 1.93E+04 | 1.47E+04 | 1.82E+04 | 2.31E+04 |
| Cer(d18:1_14:0) | 1.28E+05 | 2.79E+05 | 4.25E+05 | 2.57E+05 | 4.16E+05 | 3.36E+05 | 3.17E+05 | 6.03E+05 | 3.51E+05 | 3.18E+05 | 3.03E+05 | 2.55E+05 | 3.52E+05 | 3.25E+05 | 3.37E+05 | 3.94E+05 |
| Cer(d18:2_14:0) | 4.57E+03 | 1.72E+04 | 3.18E+04 | 1.45E+04 | 1.25E+04 | 8.16E+03 | 1.35E+04 | 2.94E+04 | 2.45E+04 | 2.13E+04 | 1.82E+04 | 1.69E+04 | 1.24E+04 | 1.25E+04 | 1.44E+04 | 1.61E+04 |
| Cer(d18:1_15:0) | 6.99E+04 | 1.58E+05 | 2.87E+05 | 1.60E+05 | 2.92E+05 | 1.96E+05 | 2.47E+05 | 4.94E+05 | 1.96E+05 | 1.74E+05 | 2.01E+05 | 1.04E+05 | 2.47E+05 | 2.26E+05 | 2.76E+05 | 2.95E+05 |
| Cer(d18:1_16:0) | 2.18E+06 | 3.23E+06 | 4.55E+06 | 3.46E+06 | 6.18E+06 | 5.04E+06 | 4.68E+06 | 8.27E+06 | 4.12E+06 | 3.25E+06 | 3.98E+06 | 2.86E+06 | 5.82E+06 | 5.57E+06 | 6.02E+06 | 6.48E+06 |
| Cer(d18:2_16:0) | 2.57E+05 | 4.11E+05 | 8.09E+05 | 5.18E+05 | 4.29E+05 | 2.91E+05 | 4.08E+05 | 6.54E+05 | 4.88E+05 | 5.27E+05 | 5.18E+05 | 4.14E+05 | 3.35E+05 | 2.80E+05 | 3.08E+05 | 4.19E+05 |
| Cer(d18:1_17:0) | 2.41E+04 | 4.89E+04 | 9.55E+04 | 3.54E+04 | 2.26E+05 | 1.63E+05 | 1.56E+05 | 3.34E+05 | 6.05E+04 | 3.96E+04 | 5.38E+04 | 3.55E+04 | 1.06E+05 | 9.23E+04 | 9.16E+04 | 8.96E+04 |
| Cer(d18:1_18:0) | 5.29E+04 | 1.83E+05 | 2.11E+05 | 8.92E+04 | 1.35E+06 | 1.23E+06 | 1.16E+06 | 2.04E+06 | 1.80E+05 | 1.08E+05 | 1.26E+05 | 8.56E+04 | 1.56E+05 | 1.65E+05 | 1.73E+05 | 1.67E+05 |
| Cer(d18:2_18:0) | 1.27E+04 | 3.64E+04 | 4.33E+04 | 2.49E+04 | 2.83E+05 | 2.19E+05 | 2.57E+05 | 2.91E+05 | 3.89E+04 | 2.80E+04 | 2.99E+04 | 2.68E+04 | 3.92E+04 | 4.05E+04 | 4.27E+04 | 4.93E+04 |
| Cer(d18:1_20:0) | 2.42E+04 | 5.41E+04 | 8.75E+04 | 3.39E+04 | 1.13E+05 | 1.02E+05 | 1.03E+05 | 2.20E+05 | 6.67E+04 | 3.92E+04 | 5.89E+04 | 3.51E+04 | 7.92E+04 | 6.47E+04 | 8.22E+04 | 6.95E+04 |
| Cer(d18:0_22:0) | 4.57E+04 | 3.85E+04 | 6.75E+04 | 3.54E+04 | 2.28E+04 | 1.43E+04 | 1.67E+04 | 2.83E+04 | 3.81E+04 | 2.22E+04 | 5.70E+04 | 1.86E+04 | 5.79E+04 | 4.36E+04 | 5.18E+04 | 4.93E+04 |
| Cer(d18:1_22:0) | 2.36E+05 | 4.84E+05 | 9.24E+05 | 3.57E+05 | 8.43E+05 | 6.55E+05 | 8.07E+05 | 1.71E+06 | 6.72E+05 | 4.17E+05 | 6.56E+05 | 3.61E+05 | 9.29E+05 | 7.03E+05 | 8.27E+05 | 6.74E+05 |
| Cer(d16:1_24:1) | 3.32E+04 | 6.59E+04 | 1.36E+05 | 5.87E+04 | 1.33E+05 | 9.82E+04 | 8.32E+04 | 2.61E+05 | 1.05E+05 | 6.42E+04 | 1.04E+05 | 5.15E+04 | 8.95E+04 | 7.74E+04 | 9.18E+04 | 1.01E+05 |
| Cer(d18:2_22:0) | 3.75E+04 | 7.60E+04 | 1.41E+05 | 6.30E+04 | 9.13E+04 | 6.66E+04 | 9.94E+04 | 1.74E+05 | 9.97E+04 | 7.10E+04 | 1.10E+05 | 6.24E+04 | 8.60E+04 | 7.00E+04 | 8.40E+04 | 7.63E+04 |
| Cer(d18:1_23:0) | 7.51E+04 | 1.33E+05 | 2.51E+05 | 1.14E+05 | 1.72E+05 | 1.37E+05 | 1.93E+05 | 3.49E+05 | 1.53E+05 | 1.17E+05 | 1.67E+05 | 1.09E+05 | 2.25E+05 | 1.94E+05 | 2.13E+05 | 1.73E+05 |
| Cer(d18:1_23:1) | 2.70E+04 | 5.28E+04 | 1.05E+05 | 4.73E+04 | 9.38E+04 | 7.02E+04 | 7.08E+04 | 1.89E+05 | 7.88E+04 | 5.78E+04 | 8.50E+04 | 4.62E+04 | 8.27E+04 | 7.60E+04 | 9.14E+04 | 8.42E+04 |
| Cer(d18:0_24:0) | 1.28E+05 | 9.07E+04 | 2.17E+05 | 1.25E+05 | 2.47E+04 | 1.89E+04 | 1.74E+04 | 4.21E+04 | 1.17E+05 | 5.57E+04 | 1.75E+05 | 5.16E+04 | 1.70E+05 | 1.40E+05 | 1.80E+05 | 1.80E+05 |
| Cer(d18:0_24:1) | 1.23E+05 | 1.01E+05 | 2.50E+05 | 1.07E+05 | 4.29E+04 | 2.62E+04 | 9.56E+03 | 6.15E+04 | 1.52E+05 | 6.72E+04 | 1.96E+05 | 3.83E+04 | 1.22E+05 | 1.08E+05 | 1.18E+05 | 1.78E+05 |
| Cer(d18:1_24:0) | 1.09E+06 | 1.94E+06 | 3.74E+06 | 1.83E+06 | 1.73E+06 | 1.52E+06 | 1.97E+06 | 3.96E+06 | 2.50E+06 | 1.88E+06 | 3.02E+06 | 1.85E+06 | 3.39E+06 | 2.67E+06 | 3.42E+06 | 2.37E+06 |
| Cer(d18:1_24:1) | 9.46E+05 | 1.41E+06 | 3.02E+06 | 1.48E+06 | 2.24E+06 | 1.89E+06 | 1.36E+06 | 3.97E+06 | 2.38E+06 | 1.61E+06 | 2.42E+06 | 1.41E+06 | 2.39E+06 | 2.18E+06 | 2.61E+06 | 2.28E+06 |
| Cer(d18:2_24:0) | 2.42E+05 | 5.42E+05 | 1.18E+06 | 4.68E+05 | 2.86E+05 | 1.78E+05 | 3.59E+05 | 6.29E+05 | 7.12E+05 | 5.61E+05 | 7.26E+05 | 5.11E+05 | 5.20E+05 | 3.94E+05 | 4.53E+05 | 3.77E+05 |
| Cer(d18:2_24:1) | 1.68E+05 | 3.19E+05 | 7.72E+05 | 3.35E+05 | 3.39E+05 | 2.26E+05 | 2.30E+05 | 6.70E+05 | 4.83E+05 | 3.84E+05 | 5.55E+05 | 3.14E+05 | 2.76E+05 | 2.32E+05 | 2.62E+05 | 3.17E+05 |
| Cer(d19:1_24:0) | 4.12E+04 | 6.96E+04 | 1.66E+05 | 6.72E+04 | 4.49E+04 | 3.50E+04 | 4.67E+04 | 9.30E+04 | 8.46E+04 | 7.56E+04 | 9.43E+04 | 7.74E+04 | 1.60E+05 | 1.35E+05 | 1.81E+05 | 1.23E+05 |
| Cer(d19:1_24:1) | 3.10E+04 | 6.50E+04 | 1.61E+05 | 5.89E+04 | 5.91E+04 | 4.24E+04 | 3.23E+04 | 1.16E+05 | 1.04E+05 | 9.31E+04 | 1.05E+05 | 8.04E+04 | 1.44E+05 | 1.28E+05 | 1.51E+05 | 1.25E+05 |
| Cer(d20:1_24:0) | 9.53E+04 | 2.18E+05 | 4.19E+05 | 1.79E+05 | 4.44E+04 | 3.88E+04 | 5.69E+04 | 1.17E+05 | 2.64E+05 | 2.17E+05 | 2.56E+05 | 2.42E+05 | 4.18E+05 | 3.60E+05 | 4.52E+05 | 2.91E+05 |
| Cer(d20:1_24:1) | 1.12E+05 | 2.19E+05 | 4.19E+05 | 1.78E+05 | 6.96E+04 | 6.31E+04 | 4.05E+04 | 1.34E+05 | 3.49E+05 | 2.93E+05 | 3.33E+05 | 2.55E+05 | 3.07E+05 | 2.39E+05 | 3.07E+05 | 2.79E+05 |
| Cer(t16:0_18:1) | 1.31E+04 | 1.80E+04 | 1.25E+04 | 1.22E+04 | 3.02E+04 | 3.71E+04 | 3.40E+04 | 2.50E+04 | 1.22E+04 | 1.11E+04 | 1.33E+04 | 1.24E+04 | 1.27E+05 | 1.11E+05 | 9.52E+04 | 1.17E+05 |
| Cer(t18:1_23:0) | 2.19E+03 | 3.23E+03 | 3.68E+03 | 2.37E+03 | 7.06E+03 | 5.76E+03 | 5.81E+03 | 1.04E+04 | 4.89E+03 | 2.78E+03 | 5.96E+03 | 4.40E+03 | 3.53E+04 | 3.14E+04 | 3.65E+04 | 3.00E+04 |
| Cer(t18:0_24:1) | 1.94E+03 | 9.02E+03 | 1.05E+04 | 6.12E+03 | 1.09E+04 | 6.96E+03 | 1.20E+04 | 1.32E+04 | 4.87E+03 | 3.29E+03 | 8.41E+03 | 6.81E+03 | 1.72E+05 | 1.59E+05 | 1.77E+05 | 1.62E+05 |

| | | | | | | | | | | | | | | | | |
|---|---|---|---|---|---|---|---|---|---|---|---|---|---|---|---|---|
| CL(14:0_16:1_18:1_18:1) | 4.32E+04 | 6.17E+04 | 7.86E+04 | 4.06E+04 | 5.46E+04 | 4.67E+04 | 9.31E+04 | 9.52E+04 | 4.52E+04 | 3.44E+04 | 4.82E+04 | 4.46E+04 | 6.02E+04 | 6.28E+04 | 8.05E+04 | 7.75E+04 |
| CL(16:1_16:1_16:1_18:1) | 8.81E+04 | 9.51E+04 | 1.12E+05 | 9.27E+04 | 6.86E+04 | 3.93E+04 | 1.08E+05 | 1.14E+05 | 9.68E+04 | 4.63E+04 | 8.76E+04 | 4.42E+04 | 5.46E+04 | 6.73E+04 | 7.57E+04 | 7.09E+04 |
| CL(16:0_16:0_18:1_18:1) | 5.80E+04 | 5.33E+04 | 1.16E+05 | 5.47E+04 | 6.42E+04 | 3.92E+04 | 7.23E+04 | 1.17E+05 | 3.87E+04 | 3.88E+04 | 6.25E+04 | 5.35E+04 | 7.44E+04 | 8.00E+04 | 1.31E+05 | 8.08E+04 |
| CL(16:0_16:1_18:1_18:1) | 9.23E+04 | 7.12E+04 | 1.89E+05 | 1.04E+05 | 1.07E+05 | 9.46E+04 | 1.31E+05 | 1.78E+05 | 8.25E+04 | 6.89E+04 | 1.07E+05 | 1.12E+05 | 1.24E+05 | 1.17E+05 | 1.52E+05 | 1.43E+05 |
| CL(16:1_16:1_18:1_18:1) | 2.12E+05 | 2.41E+05 | 3.60E+05 | 2.36E+05 | 2.69E+05 | 1.93E+05 | 3.23E+05 | 3.22E+05 | 2.49E+05 | 1.56E+05 | 2.83E+05 | 2.18E+05 | 2.78E+05 | 2.76E+05 | 3.72E+05 | 2.99E+05 |
| CL(16:1_16:1_18:1_18:2) | 1.07E+05 | 1.62E+05 | 2.27E+05 | 1.33E+05 | 1.12E+05 | 9.40E+04 | 1.26E+05 | 2.03E+05 | 1.34E+05 | 9.29E+04 | 1.52E+05 | 1.22E+05 | 8.67E+04 | 9.34E+04 | 1.23E+05 | 8.39E+04 |
| CL(16:1_17:1_18:1_18:1) | 2.14E+04 | 3.85E+04 | 7.61E+04 | 1.94E+04 | 8.21E+04 | 5.05E+04 | 8.36E+04 | 1.11E+05 | 2.60E+04 | 2.55E+04 | 2.56E+04 | 2.92E+04 | 7.14E+04 | 4.55E+04 | 9.30E+04 | 6.24E+04 |
| CL(16:1_17:1_18:1_18:2) | 9.70E+03 | 2.66E+04 | 4.60E+04 | 1.52E+04 | 3.75E+04 | 2.64E+04 | 4.04E+04 | 5.51E+04 | 1.76E+04 | 1.69E+04 | 1.81E+04 | 2.32E+04 | 2.87E+04 | 2.62E+04 | 3.89E+04 | 1.60E+04 |
| CL(16:0_18:1_18:1_18:1) | 4.35E+04 | 4.19E+04 | 9.05E+04 | 4.37E+04 | 9.44E+04 | 8.69E+04 | 7.63E+04 | 9.78E+04 | 3.08E+04 | 3.05E+04 | 4.94E+04 | 4.76E+04 | 6.30E+04 | 7.46E+04 | 8.59E+04 | 6.60E+04 |
| CL(16:1_18:1_18:1_18:1) | 1.44E+05 | 1.85E+05 | 3.06E+05 | 1.42E+05 | 3.14E+05 | 2.46E+05 | 3.16E+05 | 3.63E+05 | 8.31E+04 | 1.29E+05 | 1.49E+05 | 1.85E+05 | 3.02E+05 | 2.82E+05 | 3.51E+05 | 3.19E+05 |
| CL(16:1_18:1_18:1_18:2) | 2.16E+05 | 2.99E+05 | 5.04E+05 | 2.58E+05 | 3.51E+05 | 2.81E+05 | 3.14E+05 | 5.44E+05 | 2.60E+05 | 2.25E+05 | 2.96E+05 | 3.27E+05 | 3.34E+05 | 3.74E+05 | 4.60E+05 | 3.13E+05 |
| CL(16:1_18:1_18:2_18:2) | 7.34E+04 | 1.31E+05 | 2.14E+05 | 9.70E+04 | 1.02E+05 | 9.27E+04 | 8.55E+04 | 1.76E+05 | 9.93E+04 | 8.71E+04 | 1.04E+05 | 1.22E+05 | 8.92E+04 | 8.41E+04 | 1.11E+05 | 7.32E+04 |
| CL(17:1_18:1_18:1_18:1) | 7.03E+03 | 1.10E+04 | 4.02E+04 | 7.83E+03 | 6.41E+04 | 4.44E+04 | 5.55E+04 | 7.20E+04 | 9.18E+03 | 6.87E+03 | 1.05E+04 | 1.16E+04 | 4.43E+04 | 4.69E+04 | 6.05E+04 | 4.83E+04 |
| CL(17:1_18:1_18:1_18:2) | 1.32E+04 | 2.35E+04 | 5.97E+04 | 2.35E+04 | 6.63E+04 | 4.76E+04 | 5.14E+04 | 9.05E+04 | 1.53E+04 | 2.21E+04 | 1.86E+04 | 2.92E+04 | 6.02E+04 | 6.06E+04 | 8.04E+04 | 4.55E+04 |
| CL(18:1_18:1_18:1_18:1) | 2.93E+04 | 3.38E+04 | 5.86E+04 | 1.97E+04 | 1.48E+05 | 8.55E+04 | 1.00E+05 | 1.49E+05 | 2.16E+04 | 2.26E+04 | 9.93E+03 | 1.93E+04 | 9.74E+04 | 7.90E+04 | 1.06E+05 | 8.69E+04 |
| CL(18:1_18:1_18:1_18:2) | 7.20E+04 | 1.17E+05 | 1.43E+05 | 6.34E+04 | 2.40E+05 | 1.87E+05 | 1.35E+05 | 3.01E+05 | 7.81E+04 | 7.70E+04 | 8.23E+04 | 1.15E+05 | 1.97E+05 | 2.15E+05 | 2.20E+05 | 1.91E+05 |
| CL(18:1_18:1_18:2_18:2) | 8.19E+04 | 1.47E+05 | 2.99E+05 | 9.38E+04 | 1.97E+05 | 1.40E+05 | 1.40E+05 | 3.01E+05 | 1.09E+05 | 1.32E+05 | 1.09E+05 | 1.69E+05 | 2.15E+05 | 2.08E+05 | 2.86E+05 | 1.88E+05 |
| CL(18:1_18:2_18:2_18:2) | 2.66E+04 | 6.96E+04 | 1.27E+05 | 3.79E+04 | 5.38E+04 | 4.78E+04 | 3.79E+04 | 8.92E+04 | 3.21E+04 | 5.07E+04 | 3.62E+04 | 7.37E+04 | 6.01E+04 | 5.34E+04 | 5.95E+04 | 4.66E+04 |
| CL(16:1_18:2_18:2_20:4) | 3.02E+04 | 5.37E+04 | 8.72E+04 | 3.70E+04 | 4.38E+04 | 3.19E+04 | 3.80E+04 | 7.59E+04 | 3.97E+04 | 3.17E+04 | 3.72E+04 | 4.65E+04 | 3.44E+04 | 3.38E+04 | 4.41E+04 | 3.24E+04 |
| CL(18:1_18:1_18:2_20:3) | 1.02E+04 | 2.10E+04 | 4.46E+04 | 8.93E+03 | 4.23E+04 | 3.37E+04 | 2.77E+04 | 5.82E+04 | 1.25E+04 | 1.82E+04 | 1.18E+04 | 1.94E+04 | 6.64E+04 | 5.60E+04 | 7.39E+04 | 4.93E+04 |
| DGDG(16:0_16:1) | 1.56E+04 | 1.14E+04 | 1.31E+04 | 1.59E+04 | 8.90E+02 | 7.03E+02 | 1.83E+03 | 1.94E+03 | 2.46E+04 | 1.56E+04 | 3.42E+04 | 1.90E+04 | 5.88E+04 | 6.09E+04 | 6.09E+04 | 7.14E+04 |
| DGDG(18:0_16:1) | 1.16E+04 | 8.67E+03 | 7.64E+03 | 1.04E+04 | NP | NP | NP | NP | 3.16E+04 | 2.22E+04 | 2.94E+04 | 2.13E+04 | 1.33E+05 | 1.10E+05 | 1.21E+05 | 1.24E+05 |

| | | | | | | | | | | | | | | | | |
|---|---|---|---|---|---|---|---|---|---|---|---|---|---|---|---|---|
| LPC(14:0) | 1.40E+04 | 2.40E+04 | 4.42E+04 | 2.79E+04 | 1.73E+04 | 4.38E+04 | 5.31E+04 | 7.87E+04 | 1.94E+04 | 2.14E+04 | 2.72E+04 | 2.78E+04 | 2.20E+04 | 2.96E+04 | 3.45E+04 | 4.14E+04 |
| LPC(15:0) | 7.30E+03 | 7.78E+03 | 2.35E+04 | 1.14E+04 | 1.14E+04 | 1.80E+04 | 2.20E+04 | 4.08E+04 | 7.55E+03 | 9.94E+03 | 1.28E+04 | 1.24E+04 | 1.00E+04 | 1.12E+04 | 1.63E+04 | 1.83E+04 |
| LPC(16:0) | 2.18E+05 | 3.79E+05 | 7.34E+05 | 3.69E+05 | 2.58E+05 | 5.02E+05 | 7.22E+05 | 1.19E+06 | 4.98E+05 | 4.63E+05 | 5.12E+05 | 4.31E+05 | 7.95E+05 | 6.53E+05 | 8.14E+05 | 6.90E+05 |
| LPC(16:1) | 2.50E+04 | 4.56E+04 | 1.15E+05 | 6.94E+04 | 3.69E+04 | 6.79E+04 | 7.89E+04 | 1.31E+05 | 5.68E+04 | 5.84E+04 | 7.45E+04 | 6.97E+04 | 7.43E+04 | 6.20E+04 | 9.73E+04 | 1.56E+05 |
| LPC(17:0) | 5.35E+03 | 8.06E+03 | 1.73E+04 | 7.23E+03 | 9.93E+03 | 1.52E+04 | 2.10E+04 | 3.27E+04 | 1.31E+04 | 9.48E+03 | 1.16E+04 | 7.91E+03 | 2.69E+04 | 2.16E+04 | 2.25E+04 | 1.98E+04 |
| LPC(17:1) | 4.20E+03 | 7.96E+03 | 2.13E+04 | 1.08E+04 | 8.96E+03 | 1.35E+04 | 1.29E+04 | 3.13E+04 | 1.12E+04 | 9.80E+03 | 1.22E+04 | 9.77E+03 | 1.12E+04 | 1.28E+04 | 1.77E+04 | 1.77E+04 |
| LPC(18:0) | 1.08E+05 | 2.18E+05 | 3.06E+05 | 1.39E+05 | 1.29E+05 | 1.90E+05 | 3.22E+05 | 4.00E+05 | 3.42E+05 | 1.99E+05 | 2.58E+05 | 1.63E+05 | 5.56E+05 | 4.17E+05 | 4.54E+05 | 4.36E+05 |
| LPC(18:1) | 1.51E+05 | 2.28E+05 | 5.72E+05 | 2.85E+05 | 1.82E+05 | 2.70E+05 | 3.13E+05 | 7.77E+05 | 3.77E+05 | 2.76E+05 | 3.79E+05 | 3.11E+05 | 3.40E+05 | 4.62E+05 | 5.83E+05 | 6.05E+05 |
| LPC(18:1)+D7:(s) | 9.64E+05 | 1.02E+06 | 2.06E+06 | 1.10E+06 | 1.10E+06 | 1.10E+06 | 1.07E+06 | 2.22E+06 | 1.08E+06 | 9.16E+05 | 9.98E+05 | 1.06E+06 | 9.18E+05 | 9.45E+05 | 1.10E+06 | 1.16E+06 |
| LPC(18:2) | 9.62E+03 | 1.56E+04 | 4.53E+04 | 2.37E+04 | 1.28E+04 | 2.29E+04 | 1.31E+04 | 4.96E+04 | 1.40E+04 | 2.06E+04 | 2.38E+04 | 2.59E+04 | 1.38E+04 | 1.97E+04 | 3.08E+04 | 3.16E+04 |
| LPC(20:0) | 4.46E+03 | 1.38E+04 | 1.42E+04 | 7.18E+03 | 4.13E+03 | 6.31E+03 | 1.83E+04 | 1.40E+04 | 2.04E+04 | 1.41E+04 | 1.49E+04 | 1.23E+04 | 2.07E+04 | 1.83E+04 | 1.93E+04 | 1.79E+04 |
| LPC(20:1) | 1.07E+04 | 2.27E+04 | 3.49E+04 | 1.70E+04 | 1.26E+04 | 1.85E+04 | 3.17E+04 | 4.36E+04 | 3.10E+04 | 1.87E+04 | 2.33E+04 | 1.86E+04 | 3.42E+04 | 3.25E+04 | 4.04E+04 | 4.12E+04 |
| LPC(20:4) | 1.86E+04 | 2.40E+04 | 5.14E+04 | 2.45E+04 | 2.54E+04 | 5.73E+04 | 2.00E+04 | 4.90E+04 | 3.06E+04 | 3.21E+04 | 3.27E+04 | 3.32E+04 | 3.29E+04 | 2.42E+04 | 2.75E+04 | 2.95E+04 |
| LPC(22:5) | 6.98E+03 | 1.23E+04 | 1.85E+04 | 9.53E+03 | 7.79E+03 | 1.76E+04 | 1.11E+04 | 1.51E+04 | 1.47E+04 | 1.04E+04 | 1.39E+04 | 1.20E+04 | 1.44E+04 | 1.26E+04 | 1.72E+04 | 1.81E+04 |
| LPC(22:6) | 7.42E+03 | 1.52E+04 | 2.46E+04 | 1.39E+04 | 1.42E+04 | 2.55E+04 | 1.90E+04 | 3.20E+04 | 1.79E+04 | 1.63E+04 | 1.49E+04 | 1.64E+04 | 2.57E+04 | 2.02E+04 | 2.32E+04 | 2.23E+04 |
| LPC(26:0) | 4.39E+03 | 3.48E+04 | 3.63E+04 | 1.91E+04 | 4.65E-03 | 3.24E-02 | 5.80E+03 | 1.92E+03 | 2.99E+04 | 2.84E+04 | 2.20E+04 | 3.19E+04 | 6.79E+04 | 5.73E+04 | 5.56E+04 | 5.33E+04 |
| LPC(26:1) | 5.46E+03 | 3.36E+04 | 3.13E+04 | 1.34E+04 | 2.13E+03 | 1.09E+03 | 4.64E+03 | 3.48E+03 | 4.16E+04 | 3.32E+04 | 2.16E+04 | 2.93E+04 | 3.17E+04 | 2.15E+04 | 2.64E+04 | 2.56E+04 |
| LPC(O-16:0) | 1.12E+04 | 6.50E+03 | 1.63E+04 | 1.18E+04 | 3.39E+03 | 4.93E+03 | 1.83E+04 | 1.62E+04 | 1.49E+04 | 6.41E+03 | 1.91E+04 | 5.82E+03 | 1.16E+05 | 1.08E+05 | 1.27E+05 | 1.12E+05 |
| LPC(O-16:1) | 1.97E+04 | 2.70E+04 | 5.60E+04 | 3.03E+04 | 2.19E+04 | 1.56E+04 | 3.21E+04 | 4.30E+04 | 5.22E+04 | 3.45E+04 | 4.50E+04 | 2.88E+04 | 4.56E+04 | 3.68E+04 | 4.12E+04 | 3.11E+04 |
| LPC(O-18:1) | 1.08E+04 | 2.23E+04 | 4.38E+04 | 2.14E+04 | 1.50E+04 | 1.10E+04 | 1.52E+04 | 2.50E+04 | 3.73E+04 | 2.96E+04 | 3.25E+04 | 2.13E+04 | 3.73E+04 | 3.30E+04 | 3.22E+04 | 3.13E+04 |
| LPE(16:0) | 7.28E+04 | 1.40E+05 | 2.38E+05 | 1.31E+05 | 8.71E+04 | 2.06E+05 | 2.92E+05 | 3.82E+05 | 1.89E+05 | 1.38E+05 | 1.99E+05 | 1.51E+05 | 1.22E+05 | 1.12E+05 | 1.38E+05 | 1.61E+05 |
| LPE(16:1) | 1.71E+04 | 3.32E+04 | 5.66E+04 | 3.89E+04 | 1.38E+04 | 3.12E+04 | 3.20E+04 | 5.19E+04 | 3.36E+04 | 2.99E+04 | 4.54E+04 | 3.66E+04 | 1.47E+04 | 1.92E+04 | 2.40E+04 | 4.15E+04 |
| LPE(17:0) | 5.66E+03 | 9.44E+03 | 1.86E+04 | 9.08E+03 | 1.09E+04 | 1.46E+04 | 2.79E+04 | 3.21E+04 | 1.39E+04 | 1.04E+04 | 1.19E+04 | 1.13E+04 | 1.75E+04 | 1.40E+04 | 1.57E+04 | 1.51E+04 |
| LPE(17:1) | 3.27E+03 | 5.86E+03 | 1.38E+04 | 7.20E+03 | 3.88E+03 | 8.57E+03 | 7.65E+03 | 1.59E+04 | 7.32E+03 | 5.76E+03 | 7.92E+03 | 8.44E+03 | 3.83E+03 | 3.90E+03 | 5.73E+03 | 5.83E+03 |
| LPE(18:0) | 1.90E+05 | 3.79E+05 | 6.22E+05 | 3.32E+05 | 2.25E+05 | 3.71E+05 | 7.08E+05 | 8.81E+05 | 6.68E+05 | 4.28E+05 | 4.97E+05 | 4.16E+05 | 5.38E+05 | 4.34E+05 | 5.50E+05 | 5.75E+05 |
| LPE(18:1) | 1.42E+05 | 2.75E+05 | 6.00E+05 | 3.30E+05 | 1.69E+05 | 3.13E+05 | 2.98E+05 | 5.12E+05 | 3.68E+05 | 2.94E+05 | 4.14E+05 | 3.83E+05 | 1.73E+05 | 2.17E+05 | 2.77E+05 | 3.51E+05 |
| LPE(18:1)+D7:(s) | 3.22E+05 | 3.04E+05 | 6.48E+05 | 3.24E+05 | 3.45E+05 | 3.52E+05 | 3.34E+05 | 7.47E+05 | 3.47E+05 | 3.41E+05 | 3.52E+05 | 3.65E+05 | 3.21E+05 | 3.30E+05 | 3.79E+05 | 3.82E+05 |
| LPE(18:2) | 7.91E+03 | 1.07E+04 | 2.91E+04 | 1.87E+04 | 1.05E+04 | 2.27E+04 | 1.45E+04 | 3.46E+04 | 1.36E+04 | 1.56E+04 | 2.22E+04 | 2.05E+04 | 7.93E+03 | 1.20E+04 | 1.78E+04 | 1.88E+04 |
| LPE(20:0) | 5.27E+03 | 1.35E+04 | 1.89E+04 | 1.02E+04 | 5.29E+03 | 7.66E+03 | 2.13E+04 | 1.51E+04 | 1.62E+04 | 1.04E+04 | 1.39E+04 | 1.07E+04 | 1.31E+04 | 1.11E+04 | 1.22E+04 | 1.40E+04 |
| LPE(20:1) | 8.60E+03 | 1.62E+04 | 2.82E+04 | 1.61E+04 | 1.61E+04 | 3.74E+04 | 3.68E+04 | 4.14E+04 | 2.63E+04 | 1.77E+04 | 2.33E+04 | 1.95E+04 | 1.45E+04 | 1.35E+04 | 1.73E+04 | 1.89E+04 |
| LPE(20:4) | 1.02E+05 | 1.06E+05 | 2.60E+05 | 1.42E+05 | 1.06E+05 | 3.73E+05 | 8.19E+04 | 1.84E+05 | 8.79E+04 | 1.48E+05 | 1.67E+05 | 1.31E+05 | 3.27E+04 | 8.68E+04 | 1.01E+05 | 9.15E+04 |
| LPE(22:5) | 2.66E+04 | 2.14E+04 | 4.27E+04 | 3.04E+04 | 4.71E+04 | 2.06E+05 | 2.95E+04 | 6.93E+04 | 2.37E+04 | 2.53E+04 | 3.85E+04 | 2.34E+04 | 4.19E+04 | 3.23E+04 | 5.48E+04 | 5.11E+04 |
| LPE(22:6) | 2.55E+04 | 3.69E+04 | 6.51E+04 | 3.32E+04 | 5.95E+04 | 2.53E+05 | 5.93E+04 | 1.27E+05 | 3.56E+04 | 3.71E+04 | 5.13E+04 | 4.14E+04 | 4.00E+04 | 5.42E+04 | 6.69E+04 | 6.92E+04 |

| | | | | | | | | | | | | | | | | |
|---|---|---|---|---|---|---|---|---|---|---|---|---|---|---|---|---|
| LPE(24:0) | 1.73E+04 | 2.54E+04 | 4.61E+04 | 2.18E+04 | 8.72E+03 | 6.73E+03 | 2.31E+04 | 1.81E+04 | 3.83E+04 | 2.75E+04 | 3.68E+04 | 2.61E+04 | 4.63E+04 | 3.65E+04 | 4.21E+04 | 3.67E+04 |
| LPE(24:1) | 8.64E+03 | 1.34E+04 | 2.32E+04 | 1.09E+04 | 5.25E+03 | 4.34E+03 | 1.07E+04 | 1.33E+04 | 2.27E+04 | 1.38E+04 | 2.03E+04 | 1.31E+04 | 9.54E+03 | 8.66E+03 | 8.42E+03 | 1.11E+04 |
| LPE(24:5) | 4.50E+03 | 6.81E+03 | 1.18E+04 | 5.43E+03 | 1.15E+04 | 4.03E+04 | 7.66E+03 | 2.45E+04 | 5.57E+03 | 6.36E+03 | 4.25E+03 | 6.63E+03 | 3.52E+03 | 1.98E+03 | 5.34E+03 | 5.58E+03 |
| LPI(18:0) | 3.60E+03 | 5.24E+03 | 1.79E+04 | 9.49E+03 | 1.00E+04 | 2.61E+04 | 1.09E+04 | 3.06E+04 | 8.39E+03 | 5.76E+03 | 1.04E+04 | 1.01E+04 | 6.80E+03 | 7.22E+03 | 1.13E+04 | 1.43E+04 |
| LPS(18:0) | 1.39E+04 | 2.04E+04 | 2.98E+04 | 1.93E+04 | 1.96E+04 | 2.80E+04 | 1.95E+04 | 4.03E+04 | 2.05E+04 | 1.66E+04 | 2.27E+04 | 1.94E+04 | 2.01E+04 | 2.17E+04 | 2.92E+04 | 3.70E+04 |
| PC(2:0_16:0) | NP | 5.64E+02 | 1.56E-03 | 0.00E+00 | 4.28E+02 | 1.90E+03 | 3.18E+04 | 3.90E+03 | 6.97E+02 | 0.00E+00 | NP | NP | 3.85E+02 | NP | 3.19E-02 | NP |
| PC(14:0_14:0) | 1.68E+05 | 4.46E+05 | 4.52E+05 | 3.15E+05 | 5.18E+05 | 5.37E+05 | 1.00E+06 | 1.23E+06 | 3.43E+05 | 2.90E+05 | 2.85E+05 | 2.40E+05 | 5.65E+05 | 5.14E+05 | 5.32E+05 | 6.21E+05 |
| PC(14:0_15:0) | 3.45E+04 | 3.33E+04 | 9.27E+04 | 4.87E+04 | 2.08E+05 | 1.48E+05 | 2.17E+05 | 3.80E+05 | 4.49E+04 | 4.59E+04 | 5.17E+04 | 3.66E+04 | 1.09E+05 | 1.18E+05 | 9.03E+04 | 1.19E+05 |
| PC(14:0_16:0) | 2.89E+06 | 4.65E+06 | 5.93E+06 | 2.95E+06 | 4.36E+06 | 4.22E+06 | 2.92E+06 | 7.05E+06 | 4.67E+06 | 4.00E+06 | 4.65E+06 | 3.21E+06 | 2.92E+06 | 3.10E+06 | 3.24E+06 | 3.36E+06 |
| PC(14:0_16:1) | 3.92E+05 | 8.68E+05 | 1.00E+06 | 6.87E+05 | 9.65E+05 | 9.30E+05 | 1.53E+06 | 2.14E+06 | 9.30E+05 | 7.52E+05 | 7.93E+05 | 6.03E+05 | 1.26E+06 | 1.24E+06 | 7.99E+05 | 1.77E+06 |
| PC(15:0_16:0) | 4.57E+04 | 9.03E+04 | 2.28E+05 | 1.15E+05 | 1.98E+05 | 2.57E+05 | 2.59E+05 | 4.59E+05 | 7.79E+04 | 5.41E+04 | 1.37E+05 | 3.67E+04 | 2.13E+05 | 1.76E+05 | 1.69E+05 | 1.73E+05 |
| PC(16:0_16:0) | 4.12E+06 | 5.09E+06 | 8.09E+06 | 4.86E+06 | 4.31E+06 | 3.97E+06 | 3.64E+06 | 6.97E+06 | 6.72E+06 | 5.72E+06 | 6.82E+06 | 4.66E+06 | 4.53E+06 | 3.92E+06 | 4.25E+06 | 4.32E+06 |
| PC(16:0_16:1) | 7.24E+06 | 1.14E+07 | 1.47E+07 | 1.08E+07 | 1.17E+07 | 1.07E+07 | 1.42E+07 | 1.95E+07 | 1.36E+07 | 1.16E+07 | 1.37E+07 | 9.21E+06 | 1.72E+07 | 1.61E+07 | 1.71E+07 | 2.10E+07 |
| PC(16:1_16:1) | 4.18E+05 | 1.08E+06 | 1.19E+06 | 8.06E+05 | 6.88E+05 | 7.50E+05 | 6.98E+05 | 1.71E+06 | 1.38E+06 | 1.03E+06 | 1.12E+06 | 8.33E+05 | 1.20E+05 | 3.01E+05 | 7.00E+05 | 7.44E+05 |
| PC(16:0_16:2) | 8.59E+04 | 2.20E+05 | 2.88E+05 | 1.64E+05 | 2.63E+05 | 2.88E+05 | 3.13E+05 | 5.50E+05 | 1.96E+05 | 1.59E+05 | 1.69E+05 | 1.48E+05 | 2.67E+05 | 2.21E+05 | 2.71E+05 | 2.13E+05 |
| PC(16:1_16:2) | 9.20E+03 | 3.52E+04 | 4.09E+04 | 1.91E+04 | 6.81E+04 | 4.47E+04 | 3.63E+04 | 9.03E+04 | 1.68E+04 | 2.21E+04 | 3.08E+04 | 1.54E+04 | 2.61E+04 | 5.20E+04 | 4.22E+04 | 5.98E+04 |
| PC(16:0_17:0) | 3.98E+04 | 7.25E+04 | 1.11E+05 | 6.77E+04 | 1.06E+05 | 1.50E+05 | 8.00E+04 | 4.03E+05 | 7.50E+04 | 9.38E+04 | 6.74E+04 | 8.14E+04 | 1.88E+05 | 1.05E+05 | 1.56E+05 | 1.35E+05 |
| PC(16:0_17:1) | 9.60E+05 | 1.36E+06 | 2.38E+06 | 1.56E+06 | 3.05E+06 | 2.35E+06 | 3.08E+06 | 4.97E+06 | 2.14E+06 | 1.68E+06 | 1.89E+06 | 1.27E+06 | 2.54E+06 | 2.24E+06 | 2.25E+06 | 2.22E+06 |
| PC(15:0_18:1)+D7:(s) | 3.80E+06 | 3.81E+06 | 6.79E+06 | 4.27E+06 | 3.85E+06 | 3.62E+06 | 4.28E+06 | 7.16E+06 | 3.67E+06 | 3.81E+06 | 4.39E+06 | 3.70E+06 | 3.46E+06 | 3.82E+06 | 3.95E+06 | 4.14E+06 |
| PC(16:0_18:0) | 8.70E+05 | 1.04E+06 | 2.17E+06 | 1.09E+06 | 9.95E+05 | 7.92E+05 | 7.59E+05 | 1.76E+06 | 1.71E+06 | 1.19E+06 | 1.79E+06 | 9.85E+05 | 2.12E+06 | 1.80E+06 | 1.94E+06 | 1.86E+06 |
| PC(16:0_18:1) | 2.24E+07 | 2.88E+07 | 4.66E+07 | 2.93E+07 | 4.10E+07 | 3.51E+07 | 3.66E+07 | 5.89E+07 | 4.11E+07 | 3.40E+07 | 4.07E+07 | 2.93E+07 | 4.40E+07 | 4.23E+07 | 4.27E+07 | 4.61E+07 |
| PC(16:1_18:2) | 1.92E+05 | 3.20E+05 | 5.04E+05 | 2.82E+05 | 4.60E+05 | 2.42E+05 | 2.46E+05 | 5.54E+05 | 5.02E+05 | 3.56E+05 | 4.43E+05 | 3.45E+05 | 3.39E+05 | 2.37E+05 | 3.12E+05 | 2.33E+05 |
| PC(16:0_18:3) | 1.07E+05 | 2.36E+05 | 3.09E+05 | 9.60E+04 | 2.91E+05 | 2.13E+05 | 2.72E+05 | 3.57E+05 | 2.18E+05 | 2.76E+05 | 2.38E+05 | 2.13E+05 | 2.82E+05 | 2.79E+05 | 3.06E+05 | 3.10E+05 |
| PC(14:0_20:4) | 3.70E+04 | 8.78E+04 | 1.41E+05 | 6.35E+04 | 1.73E+05 | 1.41E+05 | 1.04E+05 | 3.23E+05 | 9.48E+04 | 9.73E+04 | 7.38E+04 | 7.09E+04 | 9.40E+04 | 7.67E+04 | 8.31E+04 | 1.03E+05 |
| PC(14:0_20:5) | 1.55E+04 | 4.26E+04 | 8.39E+04 | 3.73E+04 | 6.46E+04 | 5.10E+04 | 3.17E+04 | 1.25E+05 | 5.79E+04 | 5.69E+04 | 3.78E+04 | 2.49E+04 | 2.39E+04 | 2.03E+04 | 1.73E+04 | 2.08E+04 |
| PC(17:0_18:0) | 1.28E+04 | 1.90E+04 | 3.57E+04 | 1.51E+04 | 5.00E+04 | 4.79E+04 | 3.80E+04 | 8.05E+04 | 3.27E+04 | 2.76E+04 | 3.33E+04 | 1.92E+04 | 1.75E+05 | 1.03E+05 | 1.61E+05 | 8.25E+04 |
| PC(17:0_18:1) | 7.05E+05 | 8.31E+05 | 1.92E+06 | 1.14E+06 | 1.89E+06 | 1.54E+06 | 1.68E+06 | 2.87E+06 | 1.45E+06 | 1.19E+06 | 1.41E+06 | 9.91E+05 | 3.02E+06 | 2.59E+06 | 2.74E+06 | 2.32E+06 |
| PC(17:1_18:1) | 5.67E+05 | 8.26E+05 | 1.41E+06 | 8.76E+05 | 1.23E+06 | 1.09E+06 | 8.39E+05 | 1.84E+06 | 1.26E+06 | 1.00E+06 | 1.10E+06 | 8.29E+05 | 8.54E+05 | 7.30E+05 | 7.36E+05 | 6.86E+05 |
| PC(17:1_18:2) | 4.51E+04 | 7.52E+04 | 1.53E+05 | 7.19E+04 | 6.26E+04 | 1.07E+05 | 4.05E+04 | 2.13E+05 | 1.09E+05 | 6.32E+04 | 9.98E+04 | 8.46E+04 | 9.51E+04 | 3.18E+04 | 2.14E+04 | 1.02E+05 |
| PC(15:0_20:5) | 3.45E+03 | 4.05E+03 | 1.75E+04 | 5.13E+03 | 2.76E+04 | 1.93E+04 | 1.56E+04 | 5.05E+04 | 1.09E+04 | 6.60E+03 | 6.00E+03 | 6.19E+03 | 4.11E+03 | 3.00E+03 | 1.52E+03 | 2.18E+01 |
| PC(18:0_18:1) | 4.33E+06 | 5.45E+06 | 9.93E+06 | 5.82E+06 | 8.80E+06 | 7.38E+06 | 8.49E+06 | 1.38E+07 | 9.08E+06 | 6.87E+06 | 9.12E+06 | 5.71E+06 | 1.79E+07 | 1.81E+07 | 1.75E+07 | 1.65E+07 |
| PC(18:1_18:2) | 1.13E+06 | 1.51E+06 | 2.48E+06 | 1.52E+06 | 1.66E+06 | 1.36E+06 | 1.07E+06 | 2.28E+06 | 2.24E+06 | 1.86E+06 | 2.04E+06 | 1.48E+06 | 1.13E+06 | 1.06E+06 | 1.09E+06 | 1.48E+06 |

| | | | | | | | | | | | | | | | | |
|---|---|---|---|---|---|---|---|---|---|---|---|---|---|---|---|---|
| PC(16:0_20:3) | 6.47E+05 | 5.91E+05 | 1.28E+06 | 8.54E+05 | 9.99E+05 | 8.75E+05 | 7.08E+05 | 1.73E+06 | 1.27E+06 | 1.08E+06 | 1.35E+06 | 8.32E+05 | 1.65E+06 | 1.37E+06 | 1.45E+06 | 1.66E+06 |
| PC(16:0_20:4) | 1.46E+06 | 1.69E+06 | 3.43E+06 | 1.59E+06 | 2.51E+06 | 2.24E+06 | 1.35E+06 | 3.79E+06 | 2.21E+06 | 2.47E+06 | 2.50E+06 | 2.09E+06 | 1.96E+06 | 1.47E+06 | 1.51E+06 | 1.42E+06 |
| PC(16:1_20:4) | 1.23E+05 | 1.89E+05 | 4.84E+05 | 1.74E+05 | 2.84E+05 | 2.06E+05 | 1.41E+05 | 4.76E+05 | 2.48E+05 | 2.79E+05 | 2.30E+05 | 1.97E+05 | 1.73E+05 | 1.09E+05 | 1.31E+05 | 1.49E+05 |
| PC(16:0_20:5) | 5.75E+05 | 1.24E+06 | 2.63E+06 | 1.02E+06 | 1.10E+06 | 8.68E+05 | 5.99E+05 | 2.08E+06 | 1.88E+06 | 2.04E+06 | 1.60E+06 | 1.60E+06 | 6.17E+05 | 4.52E+05 | 4.53E+05 | 4.21E+05 |
| PC(16:1_20:5) | 4.44E+04 | 7.88E+04 | 1.39E+05 | 5.86E+04 | 1.31E+05 | 1.29E+05 | 9.69E+04 | 2.80E+05 | 8.41E+04 | 9.33E+04 | 7.11E+04 | 6.63E+04 | 5.52E+04 | 5.88E+04 | 7.55E+04 | 5.21E+04 |
| PC(19:0_18:1) | 8.82E+04 | 1.37E+05 | 2.65E+05 | 1.31E+05 | 1.84E+05 | 1.55E+05 | 1.88E+05 | 3.11E+05 | 1.79E+05 | 1.28E+05 | 1.77E+05 | 1.43E+05 | 5.15E+05 | 3.81E+05 | 4.36E+05 | 3.52E+05 |
| PC(18:1_19:1) | 2.29E+05 | 3.19E+05 | 5.71E+05 | 3.44E+05 | 4.66E+05 | 3.82E+05 | 4.05E+05 | 7.01E+05 | 4.84E+05 | 3.90E+05 | 4.61E+05 | 3.18E+05 | 1.02E+06 | 9.44E+05 | 9.98E+05 | 1.01E+06 |
| PC(17:0_20:3) | 5.73E+04 | 7.55E+04 | 1.43E+05 | 7.75E+04 | 1.22E+05 | 9.52E+04 | 6.59E+04 | 1.60E+05 | 1.01E+05 | 9.04E+04 | 1.06E+05 | 8.28E+04 | 2.55E+05 | 2.40E+05 | 2.13E+05 | 2.96E+05 |
| PC(15:0_22:6) | 2.60E+04 | 3.09E+04 | 7.54E+04 | 3.17E+04 | 5.06E+04 | 4.71E+04 | 2.33E+04 | 1.27E+05 | 3.62E+04 | 5.63E+04 | 3.87E+04 | 4.75E+04 | 2.79E+04 | 2.56E+04 | 2.61E+04 | 2.05E+04 |
| PC(20:0_18:1) | 1.69E+05 | 2.44E+05 | 4.29E+05 | 2.20E+05 | 2.89E+05 | 2.00E+05 | 3.56E+05 | 4.87E+05 | 3.50E+05 | 2.55E+05 | 3.29E+05 | 2.08E+05 | 1.06E+06 | 8.37E+05 | 1.00E+06 | 9.13E+05 |
| PC(18:1_20:1) | 7.51E+05 | 1.04E+06 | 1.64E+06 | 1.10E+06 | 1.40E+06 | 1.09E+06 | 1.37E+06 | 1.92E+06 | 1.64E+06 | 1.09E+06 | 1.51E+06 | 9.35E+05 | 2.62E+06 | 2.57E+06 | 2.40E+06 | 2.92E+06 |
| PC(18:1_20:2) | 3.09E+05 | 4.13E+05 | 7.90E+05 | 4.22E+05 | 6.06E+05 | 5.19E+05 | 3.70E+05 | 8.35E+05 | 6.28E+05 | 5.02E+05 | 6.17E+05 | 4.31E+05 | 9.56E+05 | 8.00E+05 | 8.25E+05 | 1.11E+06 |
| PC(18:0_20:3) | 3.02E+05 | 3.73E+05 | 8.22E+05 | 4.00E+05 | 6.43E+05 | 5.30E+05 | 3.53E+05 | 9.95E+05 | 5.39E+05 | 4.82E+05 | 6.39E+05 | 4.22E+05 | 1.18E+06 | 9.80E+05 | 1.14E+06 | 9.92E+05 |
| PC(18:1_20:5) | 3.45E+05 | 7.57E+05 | 1.58E+06 | 7.23E+05 | 5.57E+05 | 2.70E+05 | 2.60E+05 | 1.16E+06 | 1.20E+06 | 1.26E+06 | 9.32E+05 | 1.00E+06 | 2.21E+05 | 2.41E+05 | 2.14E+05 | 2.02E+05 |
| PC(16:0_22:6) | 5.08E+05 | 1.00E+06 | 1.68E+06 | 8.56E+05 | 1.56E+06 | 1.48E+06 | 1.20E+06 | 2.91E+06 | 1.52E+06 | 1.34E+06 | 1.52E+06 | 1.01E+06 | 1.38E+06 | 1.11E+06 | 1.20E+06 | 1.02E+06 |
| PC(16:1_22:6) | 7.85E+04 | 1.96E+05 | 3.28E+05 | 1.54E+05 | 1.75E+05 | 1.57E+05 | 1.27E+05 | 3.54E+05 | 2.96E+05 | 2.73E+05 | 2.32E+05 | 2.16E+05 | 1.15E+05 | 5.13E+04 | 1.07E+05 | 1.06E+05 |
| PC(18:1_21:2) | 1.72E+04 | 2.46E+04 | 4.97E+04 | 2.17E+04 | 3.84E+04 | 2.91E+04 | 2.41E+04 | 4.54E+04 | 3.72E+04 | 2.18E+04 | 2.34E+04 | 2.99E+04 | 1.18E+05 | 6.46E+04 | 7.04E+04 | 5.59E+04 |
| PC(17:1_22:6) | 1.12E+04 | 2.69E+04 | 4.97E+04 | 2.30E+04 | 4.15E+04 | 3.50E+04 | 2.08E+04 | 8.33E+04 | 4.68E+04 | 4.24E+04 | 3.15E+04 | 3.33E+04 | 2.25E+04 | 5.47E+04 | 1.65E+04 | 1.44E+04 |
| PC(16:1_24:0) | 1.05E+05 | 2.13E+05 | 3.44E+05 | 1.75E+05 | 1.32E+05 | 9.97E+04 | 1.79E+05 | 2.06E+05 | 2.56E+05 | 1.74E+05 | 2.26E+05 | 1.73E+05 | 4.79E+05 | 4.11E+05 | 4.64E+05 | 4.38E+05 |
| PC(18:1_22:1) | 9.16E+04 | 1.44E+05 | 2.34E+05 | 1.22E+05 | 1.88E+05 | 1.41E+05 | 1.73E+05 | 2.42E+05 | 2.20E+05 | 1.35E+05 | 1.64E+05 | 1.21E+05 | 4.02E+05 | 3.13E+05 | 3.51E+05 | 4.05E+05 |
| PC(18:1_22:2) | 4.46E+04 | 6.56E+04 | 1.18E+05 | 6.04E+04 | 8.05E+04 | 6.58E+04 | 6.12E+04 | 1.06E+05 | 9.28E+04 | 6.95E+04 | 8.91E+04 | 6.34E+04 | 1.94E+05 | 1.40E+05 | 1.45E+05 | 1.76E+05 |
| PC(20:1_20:3) | 9.92E+04 | 1.22E+05 | 1.53E+05 | 4.42E+04 | 1.35E+05 | 1.21E+05 | 9.87E+04 | 2.72E+05 | 8.90E+04 | 1.47E+05 | 1.68E+05 | 1.24E+05 | 1.56E+05 | 1.75E+05 | 1.83E+05 | 1.83E+05 |
| PC(18:0_22:4) | 4.36E+04 | 4.84E+04 | 1.08E+05 | 4.30E+04 | 1.61E+05 | 1.55E+05 | 9.70E+04 | 2.96E+05 | 6.76E+04 | 6.95E+04 | 7.80E+04 | 6.42E+04 | 1.46E+05 | 1.32E+05 | 1.36E+05 | 1.38E+05 |
| PC(18:1_22:4) | 2.40E+05 | 2.37E+05 | 4.84E+05 | 2.03E+05 | 5.52E+05 | 4.60E+05 | 3.14E+05 | 8.23E+05 | 3.63E+05 | 3.48E+05 | 3.24E+05 | 2.98E+05 | 7.30E+05 | 5.04E+05 | 6.29E+05 | 1.40E+05 |
| PC(18:0_22:5) | 1.48E+05 | 1.83E+05 | 3.16E+05 | 1.92E+05 | 3.65E+05 | 3.60E+05 | 1.90E+05 | 5.50E+05 | 2.98E+05 | 2.42E+05 | 2.84E+05 | 2.09E+05 | 6.64E+05 | 5.39E+05 | 5.57E+05 | 4.35E+05 |
| PC(18:1_22:5) | 3.03E+05 | 3.96E+05 | 7.35E+05 | 3.66E+05 | 4.11E+05 | 3.36E+05 | 1.70E+05 | 5.60E+05 | 6.40E+05 | 5.18E+05 | 5.63E+05 | 4.44E+05 | 4.30E+05 | 3.29E+05 | 3.67E+05 | 3.75E+05 |
| PC(18:0_22:6) | 1.77E+05 | 2.41E+05 | 3.56E+05 | 2.17E+05 | 6.45E+05 | 5.36E+05 | 4.15E+05 | 9.72E+05 | 2.87E+05 | 2.42E+05 | 2.96E+05 | 2.16E+05 | 7.84E+05 | 6.70E+05 | 6.56E+05 | 5.24E+05 |
| PC(18:1_22:6) | 2.94E+05 | 5.58E+05 | 8.84E+05 | 4.90E+05 | 8.19E+05 | 6.68E+05 | 4.84E+05 | 1.34E+06 | 9.27E+05 | 7.57E+05 | 7.76E+05 | 6.51E+05 | 6.00E+05 | 5.20E+05 | 5.60E+05 | 5.11E+05 |
| PC(18:2_22:6) | 6.21E+04 | 1.03E+05 | 1.49E+05 | 5.94E+04 | 8.36E+04 | 5.89E+04 | 2.88E+04 | 1.14E+05 | 1.50E+05 | 1.43E+05 | 1.22E+05 | 1.13E+05 | 6.31E+04 | 3.63E+04 | 4.25E+04 | 5.25E+04 |
| PC(20:4_20:5) | 2.09E+04 | 3.58E+04 | 4.35E+04 | 1.51E+04 | 1.44E+04 | 9.08E+03 | 4.15E+03 | 2.08E+04 | 3.91E+04 | 3.87E+04 | 3.36E+04 | 3.13E+04 | 1.23E+04 | 6.33E+03 | 5.10E+03 | 7.18E+03 |
| PC(16:0_26:0) | 2.36E+04 | 9.12E+04 | 8.04E+04 | 5.60E+04 | 6.83E+03 | 3.93E+03 | 2.17E+04 | 1.84E+04 | 6.77E+04 | 5.93E+04 | 3.90E+04 | 6.57E+04 | 1.21E+05 | 9.42E+04 | 1.61E+05 | 5.44E+04 |
| PC(16:0_26:1) | 1.34E+05 | 3.72E+05 | 5.22E+05 | 1.56E+05 | 9.40E+04 | 8.07E+04 | 1.36E+05 | 1.57E+05 | 3.77E+05 | 2.82E+05 | 2.51E+05 | 3.14E+05 | 8.34E+05 | 5.55E+05 | 7.69E+05 | 6.61E+05 |
| PC(20:4_22:6) | 4.67E+04 | 7.29E+04 | 1.03E+05 | 4.13E+04 | 3.39E+04 | 2.72E+04 | 1.26E+04 | 4.54E+04 | 1.03E+05 | 1.02E+05 | 8.63E+04 | 8.09E+04 | 4.79E+04 | 2.53E+04 | 2.72E+04 | 3.25E+04 |

| | | | | | | | | | | | | | | | | |
|---|---|---|---|---|---|---|---|---|---|---|---|---|---|---|---|---|
| PC(20:5_22:6) | 1.48E+04 | 3.58E+04 | 5.50E+04 | 1.92E+04 | 1.27E+04 | 8.24E+03 | 4.70E+03 | 1.76E+04 | 5.88E+04 | 5.91E+04 | 3.86E+04 | 4.58E+04 | 1.05E+04 | 4.86E+03 | 4.84E+03 | 6.10E+03 |
| PC(18:1_24:1) | 7.84E+04 | 2.88E+05 | 4.07E+05 | 1.88E+05 | 1.43E+05 | 8.25E+04 | 1.28E+05 | 1.45E+05 | 2.64E+05 | 2.74E+05 | 3.18E+05 | 1.69E+05 | 3.79E+05 | 4.18E+05 | 4.18E+05 | 2.31E+05 |
| PC(18:0_24:5) | 3.42E+04 | 5.57E+04 | 9.90E+04 | 4.62E+04 | 1.33E+05 | 1.11E+05 | 7.13E+04 | 1.96E+05 | 7.77E+04 | 6.36E+04 | 6.28E+04 | 5.66E+04 | 7.44E+04 | 4.68E+04 | 5.17E+04 | 5.49E+04 |
| PC(18:1_24:5) | 4.74E+04 | 1.14E+05 | 1.89E+05 | 8.26E+04 | 2.12E+05 | 1.83E+05 | 1.00E+05 | 2.96E+05 | 1.54E+05 | 1.40E+05 | 1.05E+05 | 1.27E+05 | 6.69E+04 | 3.73E+04 | 5.29E+04 | 5.30E+04 |
| PC(18:1_24:6) | 1.02E+04 | 2.07E+04 | 3.56E+04 | 1.68E+04 | 5.12E+04 | 4.61E+04 | 3.57E+04 | 8.53E+04 | 3.33E+04 | 2.82E+04 | 2.86E+04 | 2.84E+04 | 4.02E+04 | 3.40E+04 | 3.67E+04 | 3.29E+04 |
| PC(20:1_22:6) | 1.85E+04 | 4.11E+04 | 4.47E+04 | 3.23E+04 | 9.04E+04 | 9.74E+04 | 7.29E+04 | 1.55E+05 | 3.72E+04 | 3.28E+04 | 3.29E+04 | 3.10E+04 | 8.57E+04 | 7.01E+04 | 7.34E+04 | 7.26E+04 |
| PC(20:4_22:5) | 5.00E+04 | 7.56E+04 | 1.32E+05 | 4.48E+04 | 5.02E+04 | 2.46E+04 | 1.32E+04 | 6.45E+04 | 9.74E+04 | 1.02E+05 | 8.36E+04 | 8.03E+04 | 6.22E+04 | 3.41E+04 | 3.83E+04 | 4.22E+04 |
| PC(22:5_22:6) | 1.25E+04 | 2.40E+04 | 2.92E+04 | 1.38E+04 | 1.72E+04 | 1.46E+04 | 6.62E+03 | 2.76E+04 | 2.48E+04 | 2.37E+04 | 2.09E+04 | 1.89E+04 | 2.69E+04 | 1.49E+04 | 1.67E+04 | 1.83E+04 |
| PC(22:6_22:6) | 7.72E+03 | 1.67E+04 | 1.86E+04 | 7.87E+03 | 1.31E+04 | 9.75E+03 | 7.47E+03 | 1.84E+04 | 1.71E+04 | 1.80E+04 | 1.39E+04 | 1.37E+04 | 1.70E+04 | 8.82E+03 | 9.42E+03 | 1.18E+04 |
| PC(18:1_26:1) | 9.84E+04 | 2.95E+05 | 3.13E+05 | 1.38E+05 | 3.39E+04 | 2.59E+04 | 5.99E+04 | 4.43E+04 | 3.23E+05 | 2.80E+05 | 2.46E+05 | 2.81E+05 | 3.23E+05 | 3.89E+05 | 3.19E+05 | 4.10E+05 |
| PC(26:1_18:2) | 1.39E+04 | 4.29E+04 | 8.02E+04 | 2.78E+04 | 1.19E+04 | 5.53E+03 | 7.26E+03 | 1.53E+04 | 5.41E+04 | 5.12E+04 | 5.10E+04 | 4.72E+04 | 2.34E+04 | 2.46E+04 | 3.51E+04 | 2.40E+04 |
| PC(18:1_28:1) | 1.25E+04 | 6.73E+04 | 6.59E+04 | 2.74E+04 | 9.27E+03 | 8.34E+03 | 1.41E+04 | 1.14E+04 | 6.71E+04 | 5.33E+04 | 2.84E+04 | 5.92E+04 | 8.56E+04 | 5.13E+04 | 1.03E+05 | 9.39E+04 |
| PC(26:0_20:4) | 9.53E+03 | 1.10E+05 | 1.68E+05 | 3.93E+04 | NP | NP | 2.83E+03 | 6.01E+03 | 4.77E+04 | 8.23E+04 | 5.69E+04 | 8.91E+04 | 1.01E+05 | 6.04E+04 | 7.59E+04 | 4.19E+04 |
| PC(O-12:0_16:0) | 2.47E+04 | 2.39E+04 | 5.94E+04 | 3.32E+04 | 4.57E+04 | 3.19E+04 | 2.43E+04 | 6.42E+04 | 4.89E+04 | 3.60E+04 | 5.68E+04 | 2.99E+04 | 4.45E+04 | 3.82E+04 | 3.63E+04 | 3.97E+04 |
| PC(O-17:0_16:0) | 1.17E+01 | 0.00E+00 | 0.00E+00 | 8.34E+00 | 0.00E+00 | 0.00E+00 | 0.00E+00 | 0.00E+00 | 1.05E+01 | 8.44E-01 | 6.26E+02 | 1.12E+01 | 1.20E+04 | 1.49E+04 | 1.83E+04 | 2.82E+04 |
| PC(O-18:0_16:0) | 1.89E+04 | 2.30E+04 | 6.23E+04 | 2.73E+04 | 3.44E+04 | 2.26E+04 | 6.99E+04 | 9.94E+04 | 4.62E+04 | 2.17E+04 | 4.27E+04 | 2.87E+04 | 1.73E+06 | 1.35E+06 | 1.48E+06 | 1.44E+06 |
| PC(O-16:0_18:1) | 1.04E+06 | 7.13E+05 | 1.41E+06 | 1.02E+06 | 1.01E+06 | 8.25E+05 | 8.82E+05 | 1.49E+06 | 1.34E+06 | 7.37E+05 | 1.64E+06 | 6.40E+05 | 7.45E+06 | 6.80E+06 | 7.07E+06 | 7.69E+06 |
| PC(O-16:1_18:2) | 5.15E+04 | 3.30E+04 | 9.14E+04 | 5.99E+04 | 9.92E+04 | 6.96E+04 | 6.52E+04 | 8.98E+04 | 8.85E+04 | 5.60E+04 | 9.15E+04 | 4.36E+04 | 1.51E+05 | 1.35E+05 | 1.28E+05 | 1.35E+05 |
| PC(O-17:0_18:1) | 1.27E+03 | 0.00E+00 | 0.00E+00 | 0.00E+00 | 2.64E+00 | 1.32E-02 | 8.53E-02 | 0.00E+00 | 7.05E-02 | 4.64E-02 | 0.00E+00 | 0.00E+00 | 2.21E+05 | 1.91E+05 | 2.01E+05 | 1.39E+05 |
| PC(O-17:1_18:1) | 1.05E+04 | 5.20E+03 | 1.07E+04 | 1.15E+04 | 1.36E+04 | 1.02E+04 | 1.34E+04 | 1.84E+04 | 1.06E+04 | 8.66E+03 | 1.31E+04 | 6.97E+03 | 5.10E+04 | 6.23E+04 | 5.91E+04 | 4.83E+04 |
| PC(O-20:0_16:0) | 7.45E+03 | 1.18E+04 | 1.70E+04 | 9.79E+03 | 3.41E+04 | 2.49E+04 | 4.48E+04 | 4.93E+04 | 1.44E+04 | 7.33E+03 | 2.01E+04 | 9.51E+03 | 4.61E+05 | 3.39E+05 | 3.89E+05 | 3.47E+05 |
| PC(O-18:0_18:1) | 6.65E+04 | 4.47E+04 | 1.06E+05 | 6.73E+04 | 1.49E+05 | 8.93E+04 | 1.50E+05 | 2.25E+05 | 9.86E+04 | 6.44E+04 | 1.47E+05 | 6.47E+04 | 2.50E+06 | 2.23E+06 | 2.43E+06 | 2.04E+06 |
| PC(O-16:0_20:3) | 3.92E+04 | 1.12E+04 | 5.20E+04 | 2.39E+04 | 1.52E+04 | 1.86E+04 | 2.23E+04 | 2.34E+04 | 4.28E+04 | 2.41E+04 | 4.30E+04 | 1.29E+04 | 3.66E+05 | 3.04E+05 | 3.12E+05 | 3.99E+05 |
| PC(O-18:2_18:1) | 9.26E+04 | 6.56E+04 | 1.15E+05 | 1.12E+05 | 2.10E+05 | 1.41E+05 | 1.64E+05 | 2.77E+05 | 1.43E+05 | 7.74E+04 | 1.70E+05 | 7.28E+04 | 3.13E+05 | 2.82E+05 | 3.38E+05 | 4.01E+05 |
| PC(O-16:0_20:4) | 2.67E+05 | 1.19E+05 | 3.43E+05 | 1.86E+05 | 2.30E+05 | 9.24E+04 | 6.72E+04 | 1.64E+05 | 2.82E+05 | 2.07E+05 | 3.73E+05 | 1.86E+05 | 7.61E+05 | 6.76E+05 | 6.60E+05 | 6.90E+05 |
| PC(O-16:0_20:5) | 6.06E+04 | 5.03E+04 | 1.08E+05 | 6.28E+04 | 2.55E+04 | 1.97E+04 | 2.29E+04 | 4.18E+04 | 9.05E+04 | 7.70E+04 | 1.01E+05 | 5.52E+04 | 1.28E+05 | 9.70E+04 | 1.10E+05 | 1.08E+05 |
| PC(O-16:1_20:4) | 2.49E+05 | 1.66E+05 | 4.50E+05 | 2.30E+05 | 5.55E+05 | 3.24E+05 | 2.06E+05 | 5.12E+05 | 3.92E+05 | 3.17E+05 | 4.93E+05 | 2.40E+05 | 2.32E+05 | 1.85E+05 | 1.91E+05 | 1.93E+05 |
| PC(O-18:0_20:3) | 2.51E+04 | 2.52E+04 | 5.60E+04 | 3.14E+04 | 3.54E+04 | 2.66E+04 | 3.41E+04 | 5.06E+04 | 4.28E+04 | 3.00E+04 | 5.85E+04 | 2.53E+04 | 2.69E+05 | 2.52E+05 | 2.07E+05 | 2.34E+05 |
| PC(O-18:0_20:4) | 8.32E+04 | 3.91E+04 | 1.37E+05 | 7.58E+04 | 1.15E+05 | 8.04E+04 | 5.90E+04 | 1.19E+05 | 1.16E+05 | 7.95E+04 | 1.33E+05 | 7.00E+04 | 4.21E+05 | 3.64E+05 | 3.62E+05 | 3.56E+05 |
| PC(O-18:1_20:4) | 8.12E+04 | 4.78E+04 | 1.51E+05 | 7.42E+04 | 1.52E+05 | 1.06E+05 | 5.34E+04 | 1.83E+05 | 1.27E+05 | 1.03E+05 | 1.42E+05 | 7.91E+04 | 1.52E+05 | 1.58E+05 | 1.48E+05 | 1.38E+05 |
| PC(O-20:6_18:0) | 4.28E+04 | 2.71E+04 | 7.24E+04 | 3.70E+04 | 2.47E+04 | 2.11E+04 | 8.31E+03 | 2.80E+04 | 6.32E+04 | 4.81E+04 | 8.21E+04 | 3.31E+04 | 5.23E+03 | 9.88E+03 | 4.81E+03 | 1.18E+04 |
| PC(O-16:1_22:5) | 1.85E+05 | 1.33E+05 | 2.97E+05 | 1.80E+05 | 4.78E+05 | 3.11E+05 | 2.14E+05 | 4.84E+05 | 3.11E+05 | 2.02E+05 | 3.58E+05 | 1.64E+05 | 5.41E+05 | 4.52E+05 | 4.70E+05 | 4.80E+05 |
| PC(O-18:1_20:5) | 3.23E+04 | 4.40E+04 | 9.09E+04 | 2.80E+04 | 7.35E+04 | 3.64E+04 | 2.67E+04 | 7.48E+04 | 6.42E+04 | 4.39E+04 | 5.23E+04 | 3.65E+04 | 5.12E+04 | 5.58E+04 | 5.67E+04 | 5.05E+04 |

| | | | | | | | | | | | | | | | | |
|---|---|---|---|---|---|---|---|---|---|---|---|---|---|---|---|---|
| PC(O-18:2_20:5) | 2.88E+04 | 2.65E+04 | 8.01E+04 | 3.01E+04 | 5.48E+04 | 3.60E+04 | 1.76E+04 | 6.31E+04 | 6.37E+04 | 4.85E+04 | 6.45E+04 | 4.34E+04 | 6.36E+03 | 5.47E+03 | 3.27E+03 | 4.82E+03 |
| PC(O-16:1_22:6) | 4.35E+04 | 4.03E+04 | 1.00E+05 | 5.76E+04 | 2.49E+05 | 1.37E+05 | 1.28E+05 | 2.52E+05 | 9.71E+04 | 6.07E+04 | 9.54E+04 | 5.01E+04 | 1.33E+05 | 1.07E+05 | 1.18E+05 | 1.10E+05 |
| PC(O-18:0_22:5) | 1.97E+04 | 1.26E+04 | 3.53E+04 | 2.07E+04 | 3.92E+04 | 3.27E+04 | 2.51E+04 | 6.57E+04 | 3.80E+04 | 2.40E+04 | 3.89E+04 | 3.04E+04 | 1.83E+05 | 1.49E+05 | 1.59E+05 | 1.39E+05 |
| PC(O-18:0_22:6) | 3.64E+04 | 2.89E+04 | 5.27E+04 | 3.80E+04 | 7.06E+04 | 6.54E+04 | 4.40E+04 | 9.34E+04 | 5.53E+04 | 4.17E+04 | 7.65E+04 | 4.38E+04 | 3.13E+05 | 2.45E+05 | 2.48E+05 | 2.38E+05 |
| PC(O-18:1_22:6) | 2.92E+04 | 1.83E+04 | 6.29E+04 | 4.19E+04 | 1.13E+05 | 7.70E+04 | 5.82E+04 | 1.36E+05 | 6.19E+04 | 3.84E+04 | 8.86E+04 | 3.46E+04 | 9.37E+04 | 8.77E+04 | 9.51E+04 | 8.29E+04 |
| PC(O-18:2_22:6) | 1.36E+04 | 1.19E+04 | 2.72E+04 | 7.73E+03 | 7.63E+04 | 4.67E+04 | 4.25E+04 | 8.32E+04 | 2.92E+04 | 1.57E+04 | 3.15E+04 | 1.51E+04 | 2.79E+04 | 2.12E+04 | 2.32E+04 | 1.57E+04 |
| PC(O-24:1_18:1) | 3.01E+03 | 7.06E+02 | 7.20E+03 | 2.84E+03 | 1.82E+04 | 1.33E+04 | 2.81E+04 | 2.95E+04 | 4.02E+03 | 1.17E+03 | 8.28E+03 | 2.15E+03 | 6.77E+04 | 1.02E+05 | 1.09E+05 | 2.16E+05 |
| PE(14:0_16:0) | 7.31E+04 | 8.39E+04 | 1.01E+05 | 9.77E+04 | 4.34E+04 | 4.34E+04 | 5.66E+04 | 8.26E+04 | 8.29E+04 | 6.40E+04 | 9.03E+04 | 5.84E+04 | 3.39E+04 | 2.46E+04 | 2.90E+04 | 3.71E+04 |
| PE(14:0_16:1) | 1.66E+04 | 4.16E+04 | 4.28E+04 | 2.90E+04 | 2.16E+04 | 2.16E+04 | 5.21E+04 | 5.98E+04 | 3.94E+04 | 3.60E+04 | 3.56E+04 | 2.98E+04 | 1.95E+04 | 1.62E+04 | 1.60E+04 | 2.54E+04 |
| PE(16:0_16:1) | 6.43E+05 | 1.04E+06 | 1.31E+06 | 1.06E+06 | 6.73E+05 | 6.26E+05 | 1.24E+06 | 1.43E+06 | 1.25E+06 | 8.93E+05 | 1.31E+06 | 8.58E+05 | 4.58E+05 | 4.83E+05 | 5.15E+05 | 6.00E+05 |
| PE(16:1_16:1) | 5.60E+04 | 1.65E+05 | 1.57E+05 | 1.13E+05 | 3.34E+04 | 4.03E+04 | 1.09E+05 | 8.90E+04 | 1.76E+05 | 1.31E+05 | 1.33E+05 | 1.12E+05 | 3.65E+04 | 3.72E+04 | 3.85E+04 | 5.94E+04 |
| PE(16:0_17:1) | 1.81E+05 | 2.48E+05 | 3.92E+05 | 3.02E+05 | 3.57E+05 | 3.21E+05 | 4.57E+05 | 5.71E+05 | 3.25E+05 | 2.64E+05 | 3.20E+05 | 2.57E+05 | 2.36E+05 | 1.94E+05 | 1.98E+05 | 2.12E+05 |
| PE(15:0_18:1)+D7:(s) | 2.36E+05 | 2.18E+05 | 4.11E+05 | 2.47E+05 | 2.48E+05 | 2.36E+05 | 2.67E+05 | 4.01E+05 | 1.77E+05 | 2.02E+05 | 1.97E+05 | 2.37E+05 | 2.01E+05 | 2.12E+05 | 2.37E+05 | 2.33E+05 |
| PE(16:1_17:1) | 2.31E+04 | 6.13E+04 | 7.27E+04 | 4.26E+04 | 3.49E+04 | 3.34E+04 | 5.22E+04 | 8.46E+04 | 6.48E+04 | 6.05E+04 | 5.65E+04 | 4.89E+04 | 1.72E+04 | 1.60E+04 | 2.27E+04 | 2.59E+04 |
| PE(16:0_18:0) | 1.34E+05 | 1.55E+05 | 2.89E+05 | 1.58E+05 | 2.22E+05 | 2.04E+05 | 1.96E+05 | 3.35E+05 | 1.83E+05 | 1.39E+05 | 2.10E+05 | 1.48E+05 | 1.40E+05 | 1.07E+05 | 1.27E+05 | 1.08E+05 |
| PE(16:0_18:1) | 5.02E+06 | 6.90E+06 | 1.03E+07 | 6.84E+06 | 6.84E+06 | 5.44E+06 | 7.30E+06 | 1.08E+07 | 9.00E+06 | 6.86E+06 | 1.02E+07 | 6.96E+06 | 4.62E+06 | 4.57E+06 | 4.69E+06 | 5.37E+06 |
| PE(16:1_18:1) | 9.71E+05 | 2.14E+06 | 2.23E+06 | 2.07E+06 | 9.08E+05 | 8.41E+05 | 1.40E+06 | 1.89E+06 | 2.69E+06 | 2.16E+06 | 2.75E+06 | 1.91E+06 | 7.30E+05 | 7.39E+05 | 8.17E+05 | 1.07E+06 |
| PE(16:1_18:2) | 3.32E+04 | 9.18E+04 | 1.21E+05 | 6.69E+04 | 4.86E+04 | 4.72E+04 | 6.17E+04 | 1.12E+05 | 1.17E+05 | 9.92E+04 | 1.17E+05 | 8.67E+04 | 3.48E+04 | 2.80E+04 | 2.94E+04 | 4.77E+04 |
| PE(14:0_20:4) | 1.06E+04 | 2.63E+04 | 3.68E+04 | 1.79E+04 | 2.32E+04 | 2.43E+04 | 2.47E+04 | 6.29E+04 | 2.47E+04 | 2.61E+04 | 2.10E+04 | 2.18E+04 | 1.94E+04 | 1.57E+04 | 1.79E+04 | 2.01E+04 |
| PE(17:0_18:1) | 4.08E+05 | 6.06E+05 | 1.20E+06 | 5.83E+05 | 9.65E+05 | 7.69E+05 | 9.77E+05 | 1.47E+06 | 7.85E+05 | 7.55E+05 | 8.05E+05 | 7.08E+05 | 7.61E+05 | 6.55E+05 | 7.39E+05 | 6.47E+05 |
| PE(17:1_18:1) | 3.11E+05 | 5.78E+05 | 8.90E+05 | 5.16E+05 | 4.80E+05 | 4.51E+05 | 5.94E+05 | 8.52E+05 | 6.92E+05 | 6.59E+05 | 6.23E+05 | 5.76E+05 | 3.00E+05 | 2.60E+05 | 3.06E+05 | 3.09E+05 |
| PE(18:0_18:1) | 5.84E+06 | 8.24E+06 | 1.38E+07 | 8.38E+06 | 9.36E+06 | 7.39E+06 | 8.41E+06 | 1.39E+07 | 1.10E+07 | 9.15E+06 | 1.18E+07 | 8.70E+06 | 1.01E+07 | 8.64E+06 | 1.01E+07 | 8.86E+06 |
| PE(18:1_18:1) | 6.38E+06 | 9.77E+06 | 1.48E+07 | 8.96E+06 | 8.39E+06 | 6.21E+06 | 8.21E+06 | 1.26E+07 | 1.33E+07 | 9.65E+06 | 1.22E+07 | 1.01E+07 | 5.27E+06 | 5.10E+06 | 5.68E+06 | 6.07E+06 |
| PE(18:0_18:2) | 1.01E+06 | 1.36E+06 | 2.73E+06 | 1.75E+06 | 1.36E+06 | 1.28E+06 | 1.17E+06 | 2.51E+06 | 1.80E+06 | 1.65E+06 | 2.47E+06 | 1.58E+06 | 1.33E+06 | 1.17E+06 | 1.34E+06 | 1.20E+06 |
| PE(18:1_18:2) | 5.92E+05 | 1.05E+06 | 1.64E+06 | 1.02E+06 | 1.01E+06 | 7.08E+05 | 9.15E+05 | 1.84E+06 | 1.38E+06 | 1.14E+06 | 1.22E+06 | 1.15E+06 | 4.80E+05 | 4.43E+05 | 5.02E+05 | 5.66E+05 |
| PE(16:0_20:3)_Rt10.5 | 2.81E+05 | 3.26E+05 | 4.83E+05 | 3.81E+05 | 3.59E+05 | 2.70E+05 | 3.14E+05 | 6.72E+05 | 3.30E+05 | 3.57E+05 | 4.35E+05 | 3.41E+05 | 4.27E+05 | 3.47E+05 | 3.64E+05 | 4.43E+05 |
| PE(16:0_20:3)_Rt11.0 | 5.09E+04 | 9.14E+04 | 1.56E+05 | 8.53E+04 | 6.48E+04 | 5.08E+04 | 9.09E+04 | 1.07E+05 | 9.29E+04 | 8.04E+04 | 9.99E+04 | 7.46E+04 | 1.61E+05 | 1.47E+05 | 1.51E+05 | 1.71E+05 |
| PE(16:0_20:4) | 1.08E+06 | 1.39E+06 | 2.87E+06 | 1.58E+06 | 1.12E+06 | 1.07E+06 | 1.08E+06 | 2.62E+06 | 2.02E+06 | 1.93E+06 | 2.01E+06 | 1.82E+06 | 1.11E+06 | 9.90E+05 | 1.10E+06 | 1.05E+06 |
| PE(16:1_20:4) | 9.14E+04 | 2.72E+05 | 3.53E+05 | 1.93E+05 | 8.27E+04 | 9.81E+04 | 6.32E+04 | 2.13E+05 | 3.05E+05 | 3.27E+05 | 2.56E+05 | 2.81E+05 | 5.82E+04 | 4.20E+04 | 5.03E+04 | 7.14E+04 |
| PE(16:0_20:5) | 2.53E+05 | 6.70E+05 | 1.04E+06 | 5.41E+05 | 2.46E+05 | 2.73E+05 | 3.39E+05 | 6.89E+05 | 7.82E+05 | 7.89E+05 | 7.01E+05 | 7.29E+05 | 1.84E+05 | 1.90E+05 | 2.11E+05 | 2.01E+05 |
| PE(16:1_20:5) | 2.66E+04 | 1.31E+05 | 1.42E+05 | 7.36E+04 | 2.84E+04 | 3.29E+04 | 2.93E+04 | 7.72E+04 | 1.28E+05 | 1.39E+05 | 8.11E+04 | 1.19E+05 | 1.33E+04 | 1.31E+04 | 1.28E+04 | 1.29E+04 |

| | | | | | | | | | | | | | | | | |
|---|---|---|---|---|---|---|---|---|---|---|---|---|---|---|---|---|
| PE(19:0_18:1)_Rt13.1 | 2.42E+04 | 3.86E+04 | 7.52E+04 | 3.56E+04 | 6.52E+04 | 4.06E+04 | 4.81E+04 | 8.48E+04 | 4.58E+04 | 3.90E+04 | 4.65E+04 | 4.09E+04 | 4.34E+04 | 3.79E+04 | 4.04E+04 | 3.71E+04 |
| PE(19:0_18:1)_Rt13.5 | 7.23E+04 | 1.05E+05 | 2.20E+05 | 9.83E+04 | 1.55E+05 | 1.13E+05 | 1.14E+05 | 2.07E+05 | 1.35E+05 | 1.15E+05 | 1.30E+05 | 1.29E+05 | 1.50E+05 | 1.20E+05 | 1.48E+05 | 1.03E+05 |
| PE(18:1_19:1) | 1.79E+05 | 2.64E+05 | 5.00E+05 | 2.60E+05 | 3.12E+05 | 2.37E+05 | 2.73E+05 | 4.65E+05 | 3.73E+05 | 3.23E+05 | 3.80E+05 | 2.97E+05 | 2.59E+05 | 2.46E+05 | 2.64E+05 | 2.39E+05 |
| PE(17:0_20:3) | 2.90E+04 | 3.58E+04 | 7.52E+04 | 3.80E+04 | 6.23E+04 | 4.30E+04 | 4.04E+04 | 8.20E+04 | 4.47E+04 | 4.30E+04 | 5.16E+04 | 4.30E+04 | 5.87E+04 | 5.42E+04 | 5.54E+04 | 5.29E+04 |
| PE(17:0_20:4) | 1.64E+05 | 1.91E+05 | 4.30E+05 | 1.94E+05 | 2.83E+05 | 2.73E+05 | 1.79E+05 | 4.43E+05 | 2.40E+05 | 2.65E+05 | 2.44E+05 | 2.70E+05 | 2.30E+05 | 1.78E+05 | 1.84E+05 | 1.32E+05 |
| PE(17:0_20:5) | 2.47E+04 | 4.77E+04 | 9.61E+04 | 3.76E+04 | 3.78E+04 | 4.40E+04 | 2.31E+04 | 6.34E+04 | 6.66E+04 | 7.28E+04 | 4.10E+04 | 6.76E+04 | 1.77E+04 | 1.10E+04 | 1.13E+04 | 1.12E+04 |
| PE(20:0_18:1) | 2.22E+05 | 3.20E+05 | 5.44E+05 | 3.11E+05 | 3.87E+05 | 2.86E+05 | 2.97E+05 | 4.84E+05 | 3.59E+05 | 2.43E+05 | 4.03E+05 | 3.07E+05 | 3.72E+05 | 3.03E+05 | 3.93E+05 | 3.24E+05 |
| PE(18:1_20:1) | 6.04E+05 | 8.52E+05 | 1.37E+06 | 8.24E+05 | 9.81E+05 | 8.05E+05 | 7.96E+05 | 1.30E+06 | 1.15E+06 | 8.37E+05 | 1.14E+06 | 8.21E+05 | 6.06E+05 | 5.32E+05 | 5.40E+05 | 6.41E+05 |
| PE(18:0_20:2) | 6.79E+04 | 6.34E+04 | 1.88E+05 | 6.77E+04 | 1.35E+05 | 8.55E+04 | 6.53E+04 | 1.56E+05 | 8.68E+04 | 1.10E+05 | 8.41E+04 | 9.92E+04 | 5.75E+05 | 4.52E+05 | 4.83E+05 | 5.04E+05 |
| PE(18:0_20:3)_Rt12.0 | 8.57E+05 | 1.10E+06 | 2.34E+06 | 1.22E+06 | 1.50E+06 | 1.10E+06 | 8.36E+05 | 1.92E+06 | 1.69E+06 | 1.55E+06 | 1.69E+06 | 1.39E+06 | 1.51E+06 | 1.10E+06 | 1.30E+06 | 1.12E+06 |
| PE(18:0_20:3)_Rt12.5 | 1.71E+05 | 2.41E+05 | 4.97E+05 | 2.49E+05 | 3.85E+05 | 2.43E+05 | 2.74E+05 | 3.87E+05 | 3.47E+05 | 2.98E+05 | 3.34E+05 | 2.90E+05 | 7.76E+05 | 5.99E+05 | 7.06E+05 | 6.26E+05 |
| PE(18:0_20:4) | 5.99E+06 | 7.80E+06 | 1.71E+07 | 7.68E+06 | 7.81E+06 | 7.08E+06 | 5.57E+06 | 1.31E+07 | 1.04E+07 | 1.04E+07 | 1.09E+07 | 9.53E+06 | 6.51E+06 | 6.14E+06 | 6.44E+06 | 5.45E+06 |
| PE(18:1_20:4) | 1.93E+06 | 3.07E+06 | 5.56E+06 | 3.03E+06 | 2.54E+06 | 2.48E+06 | 1.52E+06 | 4.54E+06 | 4.15E+06 | 3.97E+06 | 3.83E+06 | 3.56E+06 | 1.37E+06 | 1.35E+06 | 1.48E+06 | 1.40E+06 |
| PE(18:0_20:5) | 9.49E+05 | 2.31E+06 | 4.84E+06 | 2.02E+06 | 1.02E+06 | 1.15E+06 | 1.25E+06 | 2.97E+06 | 2.78E+06 | 2.91E+06 | 2.26E+06 | 2.59E+06 | 8.07E+05 | 1.00E+06 | 1.15E+06 | 9.01E+05 |
| PE(18:1_20:5) | 3.93E+05 | 1.26E+06 | 1.93E+06 | 8.59E+05 | 4.27E+05 | 4.23E+05 | 3.02E+05 | 8.72E+05 | 1.40E+06 | 1.40E+06 | 1.03E+06 | 1.29E+06 | 1.29E+05 | 2.00E+05 | 1.49E+05 | 7.63E+04 |
| PE(16:1_22:6) | 4.33E+04 | 1.36E+05 | 1.52E+05 | 8.02E+04 | 1.15E+05 | 9.99E+04 | 9.32E+04 | 2.51E+05 | 1.60E+05 | 1.52E+05 | 1.36E+05 | 1.33E+05 | 5.28E+04 | 3.85E+04 | 4.40E+04 | 5.26E+04 |
| PE(18:1_21:1) | 2.47E+04 | 3.84E+04 | 6.95E+04 | 3.55E+04 | 6.28E+04 | 4.28E+04 | 3.67E+04 | 7.19E+04 | 5.02E+04 | 3.69E+04 | 4.81E+04 | 3.85E+04 | 3.52E+04 | 2.79E+04 | 3.28E+04 | 3.00E+04 |
| PE(18:0_21:3) | 4.50E+04 | 6.65E+04 | 1.29E+05 | 6.26E+04 | 9.44E+04 | 6.29E+04 | 5.65E+04 | 9.66E+04 | 8.71E+04 | 7.59E+04 | 9.68E+04 | 8.16E+04 | 8.82E+04 | 7.23E+04 | 9.22E+04 | 7.00E+04 |
| PE(19:0_20:4) | 4.85E+04 | 7.46E+04 | 1.74E+05 | 6.95E+04 | 8.40E+04 | 7.33E+04 | 5.26E+04 | 1.32E+05 | 8.25E+04 | 1.01E+05 | 8.01E+04 | 9.44E+04 | 1.72E+04 | 1.93E+04 | 4.53E+04 | 1.13E+00 |
| PE(22:0_18:1) | 2.44E+05 | 3.89E+05 | 6.71E+05 | 3.39E+05 | 3.51E+05 | 2.22E+05 | 2.45E+05 | 4.49E+05 | 4.24E+05 | 2.56E+05 | 4.84E+05 | 3.30E+05 | 3.81E+05 | 2.62E+05 | 3.65E+05 | 1.83E+05 |
| PE(18:1_22:1) | 1.71E+05 | 2.48E+05 | 4.58E+05 | 2.18E+05 | 3.87E+05 | 2.36E+05 | 1.85E+05 | 2.21E+01 | 2.89E+05 | 2.05E+05 | 3.47E+05 | 2.24E+05 | 2.04E+05 | 1.47E+05 | 1.88E+05 | 1.11E+05 |
| PE(18:0_22:3) | 3.67E+04 | 4.75E+04 | 1.01E+05 | 4.57E+04 | 1.16E+05 | 8.49E+04 | 7.13E+04 | 1.51E+05 | 6.79E+04 | 5.49E+04 | 6.60E+04 | 5.99E+04 | 1.17E+05 | 8.46E+04 | 1.01E+05 | 8.20E+04 |
| PE(18:1_22:4) | 1.28E+05 | 1.44E+05 | 2.86E+05 | 1.23E+05 | 4.29E+05 | 3.61E+05 | 1.79E+05 | 6.10E+05 | 2.44E+05 | 2.14E+05 | 2.92E+05 | 2.26E+05 | 4.99E+04 | 1.97E+04 | 4.94E+04 | 4.90E+04 |
| PE(20:1_20:4) | 1.20E+05 | 1.83E+05 | 2.69E+05 | 1.67E+05 | 2.48E+05 | 2.47E+05 | 1.73E+05 | 4.49E+05 | 2.16E+05 | 2.13E+05 | 2.38E+05 | 1.64E+05 | 9.93E+04 | 7.10E+04 | 1.12E+05 | 1.01E+05 |
| PE(18:0_22:5) | 4.99E+05 | 4.40E+05 | 7.88E+05 | 4.94E+05 | 1.78E+06 | 1.67E+06 | 1.31E+06 | 2.52E+06 | 6.66E+05 | 4.91E+05 | 8.37E+05 | 4.90E+05 | 1.80E+06 | 1.56E+06 | 1.69E+06 | 1.48E+06 |
| PE(18:1_22:5) | 3.19E+05 | 3.09E+05 | 6.20E+05 | 3.89E+05 | 6.58E+05 | 5.59E+05 | 2.73E+05 | 9.71E+05 | 5.90E+05 | 4.36E+05 | 5.90E+05 | 4.55E+05 | 3.74E+05 | 3.45E+05 | 3.83E+05 | 3.50E+05 |
| PE(18:0_22:6) | 5.55E+05 | 8.28E+05 | 1.36E+06 | 7.70E+05 | 3.07E+06 | 2.70E+06 | 3.61E+06 | 5.36E+06 | 8.28E+05 | 7.44E+05 | 9.02E+05 | 7.64E+05 | 3.38E+06 | 3.37E+06 | 3.26E+06 | 3.09E+06 |
| PE(18:1_22:6) | 4.25E+05 | 8.11E+05 | 1.36E+06 | 6.78E+05 | 1.45E+06 | 1.37E+06 | 1.17E+06 | 2.74E+06 | 1.01E+06 | 8.88E+05 | 1.03E+06 | 8.97E+05 | 8.61E+05 | 7.56E+05 | 8.15E+05 | 8.78E+05 |
| PE(18:2_22:6) | 8.04E+03 | 2.08E+04 | 3.65E+04 | 1.35E+04 | 3.81E+04 | 4.18E+04 | 2.42E+04 | 6.23E+04 | 3.20E+04 | 3.62E+04 | 2.41E+04 | 2.80E+04 | 1.27E+04 | 6.74E+03 | 1.23E+04 | 1.07E+04 |
| PE(23:0_18:1) | 2.69E+04 | 4.75E+04 | 7.97E+04 | 3.49E+04 | 3.33E+04 | 2.01E+04 | 2.35E+04 | 3.67E+04 | 3.92E+04 | 2.79E+04 | 4.37E+04 | 3.80E+04 | 4.08E+04 | 2.49E+04 | 4.28E+04 | 3.12E+04 |
| PE(24:0_18:1) | 3.11E+05 | 5.57E+05 | 9.13E+05 | 4.25E+05 | 2.91E+05 | 1.86E+05 | 2.17E+05 | 3.26E+05 | 4.66E+05 | 3.40E+05 | 5.70E+05 | 4.66E+05 | 4.42E+05 | 3.33E+05 | 5.06E+05 | 3.34E+05 |

| | | | | | | | | | | | | | | | | |
|---|---|---|---|---|---|---|---|---|---|---|---|---|---|---|---|---|
| PE(18:1_24:1) | 1.76E+05 | 3.19E+05 | 5.43E+05 | 2.44E+05 | 2.25E+05 | 1.45E+05 | 9.70E+04 | 2.25E+05 | 3.87E+05 | 2.54E+05 | 4.03E+05 | 3.24E+05 | 1.28E+05 | 1.01E+05 | 1.42E+05 | 1.12E+05 |
| PE(24:0_18:2) | 2.56E+04 | 3.93E+04 | 7.93E+04 | 3.21E+04 | 2.48E+04 | 1.51E+04 | 1.91E+04 | 3.43E+04 | 4.15E+04 | 2.78E+04 | 4.85E+04 | 4.24E+04 | 6.11E+04 | 3.85E+04 | 5.92E+04 | 3.55E+04 |
| PE(18:0_24:4) | 7.38E+03 | 1.04E+04 | 2.68E+04 | 7.09E+03 | 3.71E+04 | 3.72E+04 | 2.52E+04 | 5.85E+04 | 1.25E+04 | 1.42E+04 | 1.45E+04 | 1.64E+04 | 4.83E+03 | 2.06E+01 | 3.00E+03 | 4.96E+03 |
| PE(22:0_20:4) | 4.41E+04 | 6.85E+04 | 1.44E+05 | 5.67E+04 | 8.37E+04 | 6.20E+04 | 7.01E+04 | 1.27E+05 | 6.32E+04 | 5.08E+04 | 6.76E+04 | 6.73E+04 | 6.26E+04 | 5.00E+04 | 7.05E+04 | 5.21E+04 |
| PE(22:1_20:4) | 5.58E+04 | 8.69E+04 | 1.69E+05 | 7.54E+04 | 2.46E+05 | 2.29E+05 | 1.84E+05 | 4.03E+05 | 9.78E+04 | 9.45E+04 | 1.00E+05 | 1.01E+05 | 3.93E+04 | 2.91E+04 | 3.91E+04 | 3.35E+04 |
| PE(18:1_24:5) | 1.90E+04 | 4.80E+04 | 7.20E+04 | 3.34E+04 | 8.69E+04 | 8.97E+04 | 6.49E+04 | 1.30E+05 | 4.95E+04 | 4.81E+04 | 4.46E+04 | 6.94E+04 | NP | 6.58E-03 | 2.23E+03 | 4.88E-03 |
| PE(25:0_18:1) | 1.40E+04 | 3.14E+04 | 5.39E+04 | 1.93E+04 | 9.07E+03 | 4.41E+03 | 6.72E+03 | 8.99E+03 | 2.06E+04 | 1.87E+04 | 2.45E+04 | 2.93E+04 | 2.63E+04 | 2.16E+04 | 3.17E+04 | 2.10E+04 |
| PE(18:1_25:1) | 1.08E+04 | 3.01E+04 | 4.98E+04 | 1.75E+04 | 8.47E+03 | 4.89E+03 | 4.62E+03 | 9.14E+03 | 2.91E+04 | 2.47E+04 | 2.64E+04 | 3.24E+04 | 1.50E+04 | 1.16E+04 | 1.77E+04 | 1.15E+04 |
| PE(26:0_18:1) | 5.38E+04 | 1.60E+05 | 2.33E+05 | 8.84E+04 | 2.13E+04 | 1.08E+04 | 2.06E+04 | 2.34E+04 | 1.16E+05 | 1.06E+05 | 1.19E+05 | 1.49E+05 | 1.22E+05 | 9.61E+04 | 1.74E+05 | 1.04E+05 |
| PE(18:1_26:1) | 6.96E+04 | 1.57E+05 | 2.23E+05 | 8.96E+04 | 2.58E+04 | 1.83E+04 | 1.52E+04 | 2.92E+04 | 1.62E+05 | 1.27E+05 | 1.49E+05 | 1.69E+05 | 5.71E+04 | 4.33E+04 | 6.42E+04 | 5.91E+04 |
| PE(24:0_20:4) | 5.51E+04 | 9.81E+04 | 2.17E+05 | 8.07E+04 | 9.16E+04 | 5.98E+04 | 8.34E+04 | 1.43E+05 | 8.66E+04 | 7.66E+04 | 1.02E+05 | 9.58E+04 | 1.38E+05 | 1.16E+05 | 1.78E+05 | 9.51E+04 |
| PE(24:1_20:4) | 3.28E+04 | 5.09E+04 | 1.09E+05 | 4.57E+04 | 6.17E+04 | 4.82E+04 | 4.66E+04 | 9.35E+04 | 5.28E+04 | 4.60E+04 | 5.91E+04 | 5.83E+04 | 3.60E+04 | 3.03E+04 | 3.17E+04 | 2.58E+04 |
| PE(24:0_20:5) | 1.02E+04 | 2.65E+04 | 4.53E+04 | 1.89E+04 | 1.05E+04 | 6.68E+03 | 1.43E+04 | 2.17E+04 | 2.25E+04 | 2.09E+04 | 2.02E+04 | 2.44E+04 | 6.89E+03 | 6.84E+03 | 9.35E+03 | 5.02E+03 |
| PE(26:0_20:4) | 7.12E+03 | 2.17E+04 | 3.63E+04 | 1.28E+04 | 5.62E+03 | 4.77E+03 | 5.79E+03 | 9.47E+03 | 1.65E+04 | 1.64E+04 | 1.44E+04 | 2.16E+04 | 2.70E+04 | 1.87E+04 | 2.92E+04 | 1.88E+04 |
| PE(24:0_22:5) | 1.31E+04 | 3.63E+04 | 5.91E+04 | 1.95E+04 | 2.37E+04 | 1.52E+04 | 2.13E+04 | 3.23E+04 | 2.96E+04 | 2.81E+04 | 3.21E+04 | 3.65E+04 | 4.81E+04 | 3.55E+04 | 5.35E+04 | 3.31E+04 |
| PE(O-16:1_20:5) | 1.88E+06 | 3.62E+06 | 6.11E+06 | 3.25E+06 | 1.64E+06 | 1.11E+06 | 1.39E+06 | 2.81E+06 | 5.09E+06 | 3.78E+06 | 4.85E+06 | 3.51E+06 | 2.04E+06 | 1.73E+06 | 1.86E+06 | 1.62E+06 |
| PE(O-18:2_20:5) | 6.79E+05 | 1.29E+06 | 2.36E+06 | 1.05E+06 | 8.60E+05 | 6.32E+05 | 5.95E+05 | 1.51E+06 | 1.73E+06 | 1.28E+06 | 1.78E+06 | 1.20E+06 | 3.53E+05 | 3.14E+05 | 3.44E+05 | 2.48E+05 |
| PE(O-18:1_22:6) | 5.24E+05 | 8.55E+05 | 1.45E+06 | 8.66E+05 | 1.65E+06 | 1.18E+06 | 1.64E+06 | 2.53E+06 | 9.02E+05 | 6.80E+05 | 1.01E+06 | 6.27E+05 | 3.30E+06 | 3.21E+06 | 3.66E+06 | 2.80E+06 |
| PE(O-18:2_22:6) | 5.57E+05 | 6.94E+05 | 1.08E+06 | 7.10E+05 | 1.55E+06 | 1.15E+06 | 1.28E+06 | 2.34E+06 | 1.06E+06 | 6.29E+05 | 1.08E+06 | 6.20E+05 | 7.69E+05 | 6.49E+05 | 7.27E+05 | 5.89E+05 |
| PG(15:0_18:1)+D7:(s) | 8.51E+05 | 8.83E+05 | 1.70E+06 | 9.53E+05 | 9.53E+05 | 8.50E+05 | 9.57E+05 | 1.69E+06 | 8.67E+05 | 8.73E+05 | 9.34E+05 | 8.71E+05 | 8.03E+05 | 8.31E+05 | 9.52E+05 | 9.39E+05 |
| PG(16:0_18:1) | 5.77E+05 | 6.70E+05 | 1.16E+06 | 8.80E+05 | 5.36E+05 | 5.42E+05 | 5.30E+05 | 1.11E+06 | 1.01E+06 | 7.16E+05 | 9.91E+05 | 6.84E+05 | 7.22E+05 | 6.40E+05 | 7.23E+05 | 7.40E+05 |
| PG(16:1_18:1) | 4.12E+04 | 6.06E+04 | 9.08E+04 | 5.96E+04 | 3.55E+04 | 2.25E+04 | 3.25E+04 | 4.66E+04 | 8.55E+04 | 6.18E+04 | 7.16E+04 | 4.73E+04 | 3.23E+04 | 4.71E+04 | 4.92E+04 | 4.24E+04 |
| PG(18:1_18:1) | 5.12E+04 | 5.82E+04 | 1.29E+05 | 6.87E+04 | 6.71E+04 | 5.54E+04 | 5.39E+04 | 1.04E+05 | 8.55E+04 | 6.07E+04 | 8.81E+04 | 5.20E+04 | 6.85E+04 | 5.61E+04 | 7.93E+04 | 7.92E+04 |
| PG(16:0_22:6) | 4.63E+04 | 1.02E+05 | 8.76E+04 | 8.31E+04 | 3.61E+04 | 3.29E+04 | 2.86E+04 | 6.49E+04 | 1.46E+05 | 9.49E+04 | 1.62E+05 | 5.33E+04 | 6.96E+04 | 7.58E+04 | 8.10E+04 | 5.65E+04 |
| PG(16:1_22:6) | 3.51E+04 | 1.05E+05 | 9.38E+04 | 6.78E+04 | 3.03E+04 | 2.28E+04 | 1.35E+04 | 4.93E+04 | 1.53E+05 | 9.35E+04 | 1.24E+05 | 8.00E+04 | 2.52E+04 | 2.60E+04 | 1.96E+04 | 2.08E+04 |
| PG(18:1_22:5) | 8.62E+04 | 1.04E+05 | 1.73E+05 | 1.16E+05 | 7.07E+04 | 4.36E+04 | 2.74E+04 | 8.82E+04 | 1.98E+05 | 1.21E+05 | 2.11E+05 | 9.18E+04 | 2.45E+04 | 3.34E+04 | 3.14E+04 | 2.34E+04 |
| PG(18:1_22:6) | 3.69E+05 | 5.97E+05 | 8.89E+05 | 5.74E+05 | 4.48E+05 | 2.93E+05 | 2.58E+05 | 5.67E+05 | 9.37E+05 | 5.87E+05 | 1.01E+06 | 2.14E+05 | 2.04E+05 | 2.42E+05 | 2.32E+05 | 1.62E+05 |
| PG(18:2_22:6) | 4.13E+04 | 7.25E+04 | 1.24E+05 | 5.88E+04 | 6.84E+04 | 5.43E+04 | 2.48E+04 | 9.66E+04 | 1.24E+05 | 8.48E+04 | 8.93E+04 | 6.84E+04 | 1.83E+04 | 1.79E+04 | 1.15E+04 | 1.39E+04 |
| PG(20:4_22:6) | 3.46E+04 | 4.63E+04 | 9.79E+04 | 4.05E+04 | 3.92E+04 | 2.51E+04 | 8.97E+03 | 5.16E+04 | 9.61E+04 | 6.95E+04 | 8.91E+04 | 5.56E+04 | 8.67E+03 | 8.52E+03 | 7.09E+03 | 2.84E+03 |
| PG(20:5_22:6) | 1.62E+04 | 4.34E+04 | 7.36E+04 | 2.19E+04 | 1.72E+04 | 1.10E+04 | 1.68E+03 | 2.49E+04 | 8.30E+04 | 5.96E+04 | 5.37E+04 | 4.78E+04 | 7.72E+02 | 9.10E+02 | 1.26E+03 | 4.12E+02 |
| PG(22:5_22:6) | 2.47E+05 | 4.06E+05 | 8.72E+05 | 3.73E+05 | 1.39E+05 | 1.12E+05 | 2.42E+04 | 1.73E+05 | 6.51E+05 | 5.04E+05 | 5.27E+05 | 4.22E+05 | 2.66E+04 | 2.39E+04 | 2.76E+04 | 1.77E+04 |
| PG(22:6_22:6) | 5.17E+05 | 1.39E+06 | 2.10E+06 | 9.93E+05 | 6.06E+05 | 4.65E+05 | 1.38E+05 | 8.60E+05 | 1.73E+06 | 1.38E+06 | 1.37E+06 | 1.11E+06 | 1.22E+05 | 1.11E+05 | 1.22E+05 | 6.26E+04 |

| | | | | | | | | | | | | | | | | |
|---|---|---|---|---|---|---|---|---|---|---|---|---|---|---|---|---|
| PI(14:0_16:0) | 4.88E+03 | 9.71E+03 | 1.01E+04 | 7.78E+03 | 4.79E+03 | 4.75E+03 | 1.58E+04 | 9.81E+03 | 1.11E+04 | 1.26E+04 | 1.47E+04 | 1.12E+04 | 3.82E+04 | 2.94E+04 | 3.44E+04 | 2.90E+04 |
| PI(16:0_16:0) | 6.85E+04 | 1.08E+05 | 1.70E+05 | 1.19E+05 | 4.67E+04 | 7.15E+04 | 2.34E+05 | 1.26E+05 | 1.56E+05 | 1.28E+05 | 1.61E+05 | 1.50E+05 | 4.04E+05 | 3.37E+05 | 4.19E+05 | 2.99E+05 |
| PI(16:0_16:1) | 1.46E+05 | 2.59E+05 | 3.09E+05 | 2.36E+05 | 1.15E+05 | 1.19E+05 | 2.95E+05 | 2.76E+05 | 3.65E+05 | 3.15E+05 | 3.64E+05 | 3.17E+05 | 3.73E+05 | 3.30E+05 | 4.03E+05 | 4.16E+05 |
| PI(16:1_16:1) | 5.58E+03 | 2.32E+04 | 1.43E+04 | 9.18E+03 | 1.80E+03 | 1.29E+01 | 1.09E+04 | 8.05E+03 | 1.90E+04 | 2.52E+04 | 1.91E+04 | 2.57E+04 | 2.94E+04 | 2.50E+04 | 2.83E+04 | 4.31E+04 |
| PI(16:0_17:1) | 2.10E+04 | 2.81E+04 | 5.44E+04 | 3.55E+04 | 3.79E+04 | 3.79E+04 | 6.77E+04 | 7.36E+04 | 4.10E+04 | 4.00E+04 | 4.19E+04 | 5.09E+04 | 9.61E+04 | 6.91E+04 | 9.57E+04 | 8.81E+04 |
| PI(15:0_18:1)+D7:(s) | 1.73E+05 | 1.97E+05 | 3.62E+05 | 1.98E+05 | 2.10E+05 | 1.76E+05 | 2.24E+05 | 3.67E+05 | 1.61E+05 | 1.70E+05 | 1.78E+05 | 2.29E+05 | 1.57E+05 | 1.52E+05 | 2.04E+05 | 1.82E+05 |
| PI(16:0_18:0) | 8.28E+04 | 1.35E+05 | 2.53E+05 | 1.30E+05 | 8.55E+04 | 8.07E+04 | 2.34E+05 | 2.04E+05 | 2.53E+05 | 1.79E+05 | 2.89E+05 | 1.80E+05 | 6.10E+05 | 5.44E+05 | 6.82E+05 | 4.40E+05 |
| PI(18:0_16:1) | 1.14E+06 | 1.45E+06 | 2.50E+06 | 1.76E+06 | 1.35E+06 | 1.27E+06 | 2.15E+06 | 2.42E+06 | 2.37E+06 | 1.83E+06 | 2.73E+06 | 2.12E+06 | 4.89E+06 | 3.81E+06 | 5.29E+06 | 4.54E+06 |
| PI(16:1_18:1)_Rt8.02 | 2.09E+05 | 3.59E+05 | 4.69E+05 | 3.36E+05 | 2.04E+05 | 1.73E+05 | 2.92E+05 | 3.63E+05 | 6.49E+05 | 5.11E+05 | 4.93E+05 | 5.24E+05 | 3.21E+05 | 2.82E+05 | 3.46E+05 | 4.65E+05 |
| PI(16:1_18:1)_Rt8.24 | 4.91E+04 | 9.84E+04 | 1.60E+05 | 7.91E+04 | 9.81E+04 | 8.21E+04 | 1.14E+05 | 1.82E+05 | 1.39E+05 | 1.48E+05 | 1.52E+05 | 1.43E+05 | 2.39E+05 | 1.78E+05 | 2.34E+05 | 3.32E+05 |
| PI(17:0_18:1) | 6.63E+04 | 6.49E+04 | 1.60E+05 | 9.15E+04 | 7.92E+04 | 7.40E+04 | 1.43E+05 | 1.23E+05 | 1.03E+05 | 1.05E+05 | 1.19E+05 | 1.13E+05 | 3.31E+05 | 2.67E+05 | 3.17E+05 | 2.50E+05 |
| PI(17:1_18:1) | 2.73E+04 | 3.77E+04 | 8.40E+04 | 4.48E+04 | 4.35E+04 | 3.51E+04 | 5.92E+04 | 7.99E+04 | 7.24E+04 | 6.53E+04 | 7.46E+04 | 7.30E+04 | 7.02E+04 | 5.55E+04 | 6.91E+04 | 7.41E+04 |
| PI(16:0_19:2) | 6.26E+03 | 7.65E+03 | 1.73E+04 | 9.16E+03 | 1.74E+04 | 1.13E+04 | 2.09E+04 | 3.01E+04 | 1.18E+04 | 1.14E+04 | 1.23E+04 | 8.21E+03 | 5.59E+04 | 4.18E+04 | 5.31E+04 | 5.68E+04 |
| PI(18:0_18:0) | 3.34E+03 | 3.82E+01 | 5.67E+04 | 2.92E+04 | 1.78E+03 | 2.50E+03 | 2.39E+03 | 5.05E+04 | 6.17E+01 | 8.75E+02 | 2.71E+00 | 3.22E+03 | 6.59E+04 | 6.55E+04 | 8.39E+04 | 5.76E+04 |
| PI(18:0_18:1) | 1.25E+06 | 1.59E+06 | 3.59E+06 | 2.16E+06 | 1.17E+06 | 9.46E+05 | 2.08E+06 | 2.15E+06 | 3.27E+06 | 2.11E+06 | 3.36E+06 | 2.16E+06 | 7.48E+06 | 6.35E+06 | 7.17E+06 | 7.10E+06 |
| PI(18:1_18:1) | 5.41E+05 | 6.66E+05 | 1.05E+06 | 8.18E+05 | 6.17E+05 | 5.09E+05 | 1.10E+06 | 8.99E+05 | 1.20E+06 | 8.91E+05 | 1.31E+06 | 9.86E+05 | 1.74E+06 | 1.36E+06 | 1.72E+06 | 2.07E+06 |
| PI(18:0_18:2) | 2.24E+05 | 2.44E+05 | 5.38E+05 | 3.20E+05 | 4.52E+05 | 3.80E+05 | 4.07E+05 | 8.32E+05 | 4.55E+05 | 3.93E+05 | 4.41E+05 | 5.25E+05 | 8.27E+05 | 6.67E+05 | 8.56E+05 | 8.86E+05 |
| PI(18:1_18:2) | 5.44E+04 | 6.80E+04 | 1.51E+05 | 7.39E+04 | 1.31E+05 | 9.53E+04 | 8.84E+04 | 2.18E+05 | 1.25E+05 | 1.20E+05 | 1.45E+05 | 1.31E+05 | 9.84E+04 | 7.05E+04 | 9.07E+04 | 1.22E+05 |
| PI(16:0_20:3) | 9.44E+04 | 1.39E+05 | 2.64E+05 | 1.57E+05 | 1.61E+05 | 1.42E+05 | 2.54E+05 | 3.93E+05 | 1.73E+05 | 1.54E+05 | 1.89E+05 | 1.65E+05 | 4.21E+05 | 3.09E+05 | 3.72E+05 | 4.55E+05 |
| PI(16:0_20:4) | 3.23E+05 | 4.63E+05 | 8.48E+05 | 4.72E+05 | 6.78E+05 | 6.21E+05 | 8.60E+05 | 1.63E+06 | 4.91E+05 | 5.17E+05 | 5.44E+05 | 5.21E+05 | 6.31E+05 | 4.78E+05 | 5.66E+05 | 5.48E+05 |
| PI(16:1_20:4) | 8.72E+03 | 1.87E+04 | 2.64E+04 | 1.24E+04 | 2.49E+04 | 2.08E+04 | 2.38E+04 | 4.27E+04 | 2.10E+04 | 2.44E+04 | 2.41E+04 | 2.43E+04 | 9.07E+03 | 8.21E+03 | 1.03E+04 | 7.15E+03 |
| PI(16:0_20:5) | 1.94E+04 | 3.77E+04 | 8.77E+04 | 3.79E+04 | 5.89E+04 | 5.48E+04 | 6.72E+04 | 1.42E+05 | 6.04E+04 | 5.68E+04 | 5.24E+04 | 5.41E+04 | 4.98E+04 | 3.78E+04 | 4.73E+04 | 4.54E+04 |
| PI(18:0_19:2) | 1.13E+04 | 1.67E+04 | 2.77E+04 | 2.77E+04 | 2.37E+04 | 3.96E+04 | 7.58E+04 | 8.68E+04 | 1.67E+04 | 1.85E+04 | 2.38E+04 | 2.57E+04 | 1.33E+05 | 1.38E+05 | 1.80E+05 | 1.15E+05 |
| PI(17:0_20:3) | 1.22E+04 | 1.63E+04 | 3.15E+04 | 1.87E+04 | 3.30E+04 | 2.66E+04 | 4.04E+04 | 6.92E+04 | 1.77E+04 | 1.68E+04 | 2.15E+04 | 1.81E+04 | 5.28E+04 | 4.23E+04 | 4.96E+04 | 4.62E+04 |
| PI(18:0_19:3) | 1.28E+04 | 2.57E+04 | 4.38E+04 | 2.85E+04 | 3.43E+04 | 2.59E+04 | 3.92E+04 | 6.40E+04 | 1.96E+04 | 1.92E+04 | 2.45E+04 | 2.64E+04 | 6.83E+04 | 5.17E+04 | 4.74E+04 | 6.63E+04 |
| PI(17:0_20:4) | 4.02E+04 | 4.59E+04 | 1.37E+05 | 5.52E+04 | 2.14E+05 | 1.55E+05 | 1.51E+05 | 3.51E+05 | 5.63E+04 | 6.50E+04 | 6.76E+04 | 7.10E+04 | 9.21E+04 | 6.10E+04 | 7.23E+04 | 4.08E+04 |
| PI(17:1_20:4) | 4.44E+03 | 8.13E+03 | 1.76E+04 | 6.58E+03 | 2.66E+04 | 1.94E+04 | 1.70E+04 | 4.46E+04 | 9.72E+03 | 1.19E+04 | 1.10E+04 | 1.24E+04 | 7.16E+03 | 4.06E+03 | 3.95E+03 | 5.21E+03 |
| PI(18:0_20:1) | 3.26E+04 | 4.51E+04 | 7.00E+04 | 6.00E+04 | 5.45E+04 | 5.08E+04 | 1.64E+05 | 1.14E+05 | 5.76E+04 | 4.43E+04 | 6.85E+04 | 4.92E+04 | 2.10E+05 | 1.79E+05 | 2.51E+05 | 1.75E+05 |
| PI(18:0_20:2) | 1.07E+05 | 1.21E+05 | 2.31E+05 | 1.81E+05 | 2.53E+05 | 2.46E+05 | 4.31E+05 | 4.65E+05 | 1.72E+05 | 1.15E+05 | 1.78E+05 | 1.43E+05 | 1.90E+06 | 1.43E+06 | 1.79E+06 | 2.08E+06 |
| PI(18:0_20:3)_Rt9.76 | 4.29E+05 | 5.45E+05 | 1.27E+06 | 6.98E+05 | 1.00E+06 | 8.68E+05 | 1.01E+06 | 2.04E+06 | 7.27E+05 | 5.92E+05 | 8.30E+05 | 6.73E+05 | 1.22E+06 | 8.16E+05 | 9.87E+05 | 9.96E+05 |
| PI(18:0_20:3)_Rt10.1 | 1.67E+05 | 1.84E+05 | 4.96E+05 | 2.87E+05 | 4.89E+05 | 3.52E+05 | 4.50E+05 | 7.81E+05 | 2.33E+05 | 2.03E+05 | 2.49E+05 | 2.39E+05 | 8.26E+05 | 6.19E+05 | 8.02E+05 | 1.03E+06 |

| | | | | | | | | | | | | | | | | |
|---|---|---|---|---|---|---|---|---|---|---|---|---|---|---|---|---|
| PI(18:0_20:4) | 3.82E+06 | 3.81E+06 | 9.47E+06 | 4.70E+06 | 1.27E+07 | 9.17E+06 | 7.76E+06 | 1.94E+07 | 4.97E+06 | 4.29E+06 | 5.58E+06 | 5.20E+06 | 4.88E+06 | 3.63E+06 | 5.32E+06 | 4.02E+06 |
| PI(18:1_20:4) | 4.67E+05 | 6.65E+05 | 1.28E+06 | 7.04E+05 | 1.39E+06 | 1.06E+06 | 1.08E+06 | 2.41E+06 | 8.00E+05 | 8.57E+05 | 9.40E+05 | 8.73E+05 | 5.71E+05 | 4.41E+05 | 5.16E+05 | 5.94E+05 |
| PI(18:0_20:5) | 1.06E+05 | 2.12E+05 | 5.32E+05 | 2.10E+05 | 4.55E+05 | 3.60E+05 | 3.41E+05 | 7.81E+05 | 3.58E+05 | 3.21E+05 | 3.16E+05 | 3.47E+05 | 3.40E+05 | 2.30E+05 | 2.86E+05 | 2.61E+05 |
| PI(18:1_20:5) | 2.65E+04 | 5.71E+04 | 1.19E+05 | 5.32E+04 | 1.14E+05 | 8.06E+04 | 7.44E+04 | 1.97E+05 | 9.08E+04 | 7.77E+04 | 7.71E+04 | 8.53E+04 | 4.03E+04 | 2.55E+04 | 3.62E+04 | 4.68E+04 |
| PI(16:0_22:6) | 4.52E+04 | 1.08E+05 | 1.49E+05 | 9.64E+04 | 1.15E+05 | 1.45E+05 | 1.87E+05 | 3.15E+05 | 1.19E+05 | 1.16E+05 | 1.17E+05 | 1.22E+05 | 1.47E+05 | 1.09E+05 | 1.27E+05 | 1.17E+05 |
| PI(18:0_21:2) | 1.79E+03 | 2.34E+03 | 4.28E+03 | 3.53E+03 | 5.48E+03 | 3.67E+03 | 9.42E+03 | 1.30E+04 | 5.23E-02 | 1.85E+03 | 2.16E+03 | 3.06E+03 | 3.79E+04 | 3.47E+04 | 4.62E+04 | 3.74E+04 |
| PI(18:0_21:3) | 3.67E+04 | 5.06E+04 | 1.45E+05 | 7.42E+04 | 8.76E+04 | 7.25E+04 | 9.18E+04 | 2.20E+05 | 8.36E+04 | 5.81E+04 | 8.21E+04 | 5.80E+04 | 1.42E+05 | 1.24E+05 | 1.32E+05 | 1.36E+05 |
| PI(19:0_20:4) | 2.27E+04 | 2.72E+04 | 6.70E+04 | 3.38E+04 | 1.29E+05 | 9.75E+04 | 7.96E+04 | 1.79E+05 | 3.04E+04 | 3.79E+04 | 3.12E+04 | 3.86E+04 | 2.39E+04 | 1.44E+04 | 1.99E+04 | 1.44E+04 |
| PI(19:1_20:4) | 2.40E+04 | 2.94E+04 | 7.30E+04 | 3.19E+04 | 6.64E+04 | 5.76E+04 | 5.34E+04 | 1.09E+05 | 3.59E+04 | 3.80E+04 | 4.12E+04 | 3.80E+04 | 4.74E+04 | 4.52E+04 | 5.19E+04 | 5.33E+04 |
| PI(17:0_22:6) | 3.94E+03 | 6.23E+03 | 1.30E+04 | 6.17E+03 | 1.91E+04 | 1.60E+04 | 2.26E+04 | 3.63E+04 | 7.59E+03 | 6.84E+03 | 7.67E+03 | 9.11E+03 | 1.44E+04 | 1.02E+04 | 1.25E+04 | 4.01E+03 |
| PI(18:0_22:2) | 5.30E+03 | 8.75E+03 | 1.22E+04 | 1.13E+04 | 1.70E+04 | 1.46E+04 | 3.54E+04 | 2.80E+04 | 5.39E+03 | 7.37E+03 | 6.62E+03 | 5.61E+03 | 6.13E+04 | 5.57E+04 | 6.67E+04 | 6.89E+04 |
| PI(18:0_22:3) | 8.84E+03 | 9.10E+03 | 3.57E+04 | 1.82E+04 | 3.69E+04 | 2.37E+04 | 3.21E+04 | 5.17E+04 | 7.90E+03 | 1.27E+04 | 1.88E+04 | 1.93E+04 | 5.67E+04 | 3.84E+04 | 4.49E+04 | 5.16E+04 |
| PI(18:0_22:4) | 1.60E+05 | 1.49E+05 | 3.79E+05 | 2.18E+05 | 3.10E+05 | 2.75E+05 | 2.68E+05 | 6.67E+05 | 2.33E+05 | 1.96E+05 | 2.76E+05 | 2.57E+05 | 2.16E+05 | 1.56E+05 | 1.91E+05 | 1.59E+05 |
| PI(20:0_20:4) | 1.59E+04 | 2.08E+04 | 5.09E+04 | 2.60E+04 | 1.17E+05 | 7.24E+04 | 7.07E+04 | 1.88E+05 | 2.50E+04 | 2.33E+04 | 2.57E+04 | 3.24E+04 | 3.63E+04 | 2.22E+04 | 3.23E+04 | 2.74E+04 |
| PI(18:1_22:4) | 1.82E+04 | 1.87E+04 | 4.05E+04 | 2.16E+04 | 4.57E+04 | 3.41E+04 | 3.33E+04 | 7.85E+04 | 3.44E+04 | 3.47E+04 | 4.12E+04 | 3.37E+04 | 1.44E+04 | 1.33E+04 | 1.16E+04 | 1.63E+04 |
| PI(18:0_22:5)_Rt9.30 | 4.16E+05 | 3.77E+05 | 9.37E+05 | 6.40E+05 | 6.00E+05 | 5.48E+05 | 5.05E+05 | 1.13E+06 | 6.05E+05 | 4.71E+05 | 6.89E+05 | 5.91E+05 | 5.81E+05 | 4.30E+05 | 5.80E+05 | 4.67E+05 |
| PI(18:0_22:5)_Rt9.75 | 1.01E+04 | 1.93E+04 | 4.11E+04 | 2.44E+04 | 3.01E+04 | 3.48E+04 | 3.41E+04 | 6.90E+04 | 1.65E+04 | 1.32E+04 | 2.08E+04 | 2.20E+04 | 4.38E+04 | 2.60E+04 | 3.73E+04 | 3.46E+04 |
| PI(18:1_22:5) | 6.06E+04 | 7.32E+04 | 1.59E+05 | 9.25E+04 | 9.51E+04 | 7.98E+04 | 5.77E+04 | 1.62E+05 | 1.33E+05 | 1.10E+05 | 1.45E+05 | 1.18E+05 | 4.85E+04 | 5.42E+04 | 3.95E+04 | 5.24E+04 |
| PI(18:0_22:6) | 1.56E+05 | 2.90E+05 | 5.37E+05 | 3.52E+05 | 5.62E+05 | 4.73E+05 | 6.88E+05 | 1.16E+06 | 3.82E+05 | 3.27E+05 | 3.73E+05 | 3.67E+05 | 5.62E+05 | 4.00E+05 | 4.57E+05 | 4.37E+05 |
| PI(18:1_22:6) | 2.41E+04 | 5.53E+04 | 8.59E+04 | 5.47E+04 | 9.23E+04 | 8.12E+04 | 9.56E+04 | 1.70E+05 | 7.57E+04 | 6.71E+04 | 7.33E+04 | 6.89E+04 | 4.49E+04 | 3.20E+04 | 3.92E+04 | 4.60E+04 |
| PI(O-16:0_16:0) | 6.80E+03 | 1.08E+04 | 9.00E+03 | 1.18E+04 | 6.53E+03 | 1.12E+04 | 2.95E+04 | 1.15E+04 | 9.32E+03 | 1.15E+04 | 1.04E+04 | 9.35E+03 | 7.70E+04 | 6.18E+04 | 7.61E+04 | 7.06E+04 |
| PI(O-16:0_20:4) | 5.74E+04 | 2.28E+04 | 8.15E+04 | 4.59E+04 | 1.41E+04 | 6.18E+03 | 1.13E+04 | 2.00E+04 | 4.42E+04 | 3.52E+04 | 6.52E+04 | 3.73E+04 | 8.94E+04 | 6.30E+04 | 7.59E+04 | 6.63E+04 |
| PI(O-16:0_22:5) | 4.78E+04 | 2.59E+04 | 5.60E+04 | 3.84E+04 | 4.96E+04 | 3.85E+04 | 4.68E+04 | 7.86E+04 | 3.20E+04 | 2.83E+04 | 4.84E+04 | 2.98E+04 | 2.45E+05 | 1.83E+05 | 2.15E+05 | 1.93E+05 |
| PI(O-16:0_22:6) | 1.64E+04 | 2.03E+04 | 2.63E+04 | 2.23E+04 | 3.91E+04 | 3.67E+04 | 7.44E+04 | 7.65E+04 | 1.52E+04 | 1.61E+04 | 2.27E+04 | 1.69E+04 | 2.26E+05 | 1.71E+05 | 2.10E+05 | 1.80E+05 |
| PI(O-18:0_22:5) | 1.05E+04 | 9.59E+03 | 1.23E+04 | 1.19E+04 | 1.94E+04 | 2.07E+04 | 1.76E+04 | 2.50E+04 | 2.28E+04 | 1.76E+04 | 1.91E+04 | 1.58E+04 | 1.16E+05 | 5.63E+04 | 6.83E+04 | 4.48E+04 |
| PI(O-18:1_22:5) | 5.13E+03 | 2.04E+03 | 4.95E+03 | 4.36E+03 | 1.74E+04 | 1.55E+04 | 8.34E+03 | 2.74E+04 | 5.77E+03 | 3.18E+03 | 6.56E+03 | 4.46E+03 | 3.29E+04 | 2.32E+04 | 2.55E+04 | 2.17E+04 |
| PI(O-18:0_22:6) | 5.85E-03 | 3.29E-03 | 6.43E-03 | 5.06E-03 | 2.84E+03 | 8.96E+03 | 1.36E+04 | 6.41E+03 | 2.68E-02 | 2.90E-03 | 2.32E-02 | 9.09E-03 | 9.90E+04 | 7.16E+04 | 8.37E+04 | 6.28E+04 |
| PIP(P-22:6_18:1) | 5.91E+04 | 6.86E+04 | 9.32E+04 | 8.47E+04 | 3.84E+04 | 3.77E+04 | 8.29E+04 | 4.42E+04 | 9.31E+04 | 8.32E+04 | 1.13E+05 | 1.00E+05 | 1.53E+05 | 1.27E+05 | 1.70E+05 | 1.64E+05 |
| PIP(P-24:7_18:1) | 3.18E+04 | 3.45E+04 | 5.23E+04 | 4.22E+04 | 3.09E+04 | 2.65E+04 | 5.12E+04 | 3.28E+04 | 5.90E+04 | 4.56E+04 | 6.39E+04 | 5.00E+04 | 7.44E+04 | 5.40E+04 | 6.97E+04 | 9.08E+04 |
| PS(16:0_16:0) | 2.98E+04 | 5.32E+04 | 7.88E+04 | 4.69E+04 | 2.61E+04 | 2.03E+04 | 1.17E+04 | 3.22E+04 | 6.69E+04 | 5.68E+04 | 7.23E+04 | 5.67E+04 | 1.90E+04 | 1.25E+04 | 1.37E+04 | 1.38E+04 |
| PS(16:0_16:1) | 1.38E+05 | 2.42E+05 | 2.77E+05 | 2.16E+05 | 7.18E+04 | 5.86E+04 | 1.60E+05 | 1.40E+05 | 2.80E+05 | 2.15E+05 | 3.39E+05 | 1.97E+05 | 1.69E+05 | 1.51E+05 | 9.16E+04 | 3.09E+05 |
| PS(16:0_17:1) | 3.66E+04 | 6.32E+04 | 1.09E+05 | 6.49E+04 | 4.27E+04 | 2.87E+04 | 5.59E+04 | 6.72E+04 | 8.40E+04 | 7.10E+04 | 8.68E+04 | 5.98E+04 | 3.25E+04 | 3.39E+04 | 4.28E+04 | 4.29E+04 |

| | | | | | | | | | | | | | | | | |
|---|---|---|---|---|---|---|---|---|---|---|---|---|---|---|---|---|
| PS(15:0_18:1)+D7:(s) | 6.64E+04 | 7.16E+04 | 1.35E+05 | 7.42E+04 | 8.10E+04 | 6.65E+04 | 7.77E+04 | 1.35E+05 | 6.29E+04 | 6.54E+04 | 7.15E+04 | 7.52E+04 | 7.30E+04 | 7.18E+04 | 7.75E+04 | 6.71E+04 |
| PS(16:0_18:1) | 1.23E+06 | 1.81E+06 | 2.84E+06 | 1.96E+06 | 1.00E+06 | 7.53E+05 | 1.30E+06 | 1.67E+06 | 2.63E+06 | 1.93E+06 | 3.07E+06 | 2.15E+06 | 1.42E+06 | 1.37E+06 | 1.64E+06 | 1.68E+06 |
| PS(16:1_18:1)_Rt8.27 | 6.64E+04 | 1.24E+05 | 1.81E+05 | 1.15E+05 | 3.28E+04 | 2.58E+04 | 8.12E+04 | 9.00E+04 | 1.94E+05 | 1.34E+05 | 1.78E+05 | 1.14E+05 | 9.11E+04 | 4.85E+04 | 9.49E+04 | 1.65E+05 |
| PS(16:1_18:1)_Rt8.48 | 4.72E+04 | 6.39E+04 | 1.15E+05 | 6.82E+04 | 3.48E+04 | 2.21E+04 | 1.66E+04 | 3.96E+04 | 1.09E+05 | 7.31E+04 | 1.04E+05 | 7.99E+04 | 4.49E+04 | 5.18E+04 | 4.74E+04 | 6.93E+04 |
| PS(18:0_17:1) | 2.26E+05 | 3.47E+05 | 6.73E+05 | 3.25E+05 | 2.17E+05 | 1.47E+05 | 3.55E+05 | 7.85E+05 | 5.24E+05 | 4.10E+05 | 5.02E+05 | 4.40E+05 | 5.15E+05 | 4.50E+05 | 5.62E+05 | 4.73E+05 |
| PS(18:0_18:1) | 3.91E+06 | 5.82E+06 | 9.98E+06 | 5.41E+06 | 5.46E+06 | 4.17E+06 | 4.81E+06 | 7.86E+06 | 7.83E+06 | 6.59E+06 | 7.73E+06 | 7.68E+06 | 1.09E+07 | 9.35E+06 | 9.99E+06 | 9.40E+06 |
| PS(18:1_18:1) | 4.51E+05 | 8.22E+05 | 1.25E+06 | 7.24E+05 | 4.40E+05 | 2.99E+05 | 4.40E+05 | 6.09E+05 | 1.23E+06 | 9.88E+05 | 1.13E+06 | 8.61E+05 | 5.73E+05 | 5.19E+05 | 6.40E+05 | 7.46E+05 |
| PS(18:0_18:2) | 3.18E+05 | 4.12E+05 | 7.87E+05 | 4.16E+05 | 3.01E+05 | 2.32E+05 | 3.15E+05 | 4.76E+05 | 5.85E+05 | 6.45E+05 | 6.38E+05 | 5.25E+05 | 5.43E+05 | 4.90E+05 | 5.65E+05 | 4.91E+05 |
| PS(18:1_18:2) | 6.65E+04 | 1.05E+05 | 1.84E+05 | 9.88E+04 | 5.07E+04 | 4.25E+04 | 5.80E+04 | 1.11E+05 | 1.62E+05 | 1.39E+05 | 1.55E+05 | 1.25E+05 | 8.26E+04 | 6.36E+04 | 7.96E+04 | 1.04E+05 |
| PS(18:0_18:3) | 9.33E+03 | 2.22E+04 | 3.22E+04 | 1.46E+04 | 1.14E+04 | 6.57E+03 | 1.95E+04 | 2.38E+04 | 2.76E+04 | 2.23E+04 | 2.84E+04 | 2.37E+04 | 5.07E+04 | 4.54E+04 | 5.39E+04 | 5.69E+04 |
| PS(16:0_20:4)_Rt8.36 | 2.50E+04 | 3.71E+04 | 7.08E+04 | 3.16E+04 | 1.90E+04 | 1.36E+04 | 2.33E+04 | 3.63E+04 | 5.40E+04 | 5.53E+04 | 6.26E+04 | 5.37E+04 | 2.26E+04 | 2.14E+04 | 2.42E+04 | 2.96E+04 |
| PS(16:0_20:4)_Rt9.47 | 1.40E+05 | 1.84E+05 | 1.92E+05 | 1.70E+05 | 9.99E+04 | 7.08E+04 | 1.33E+05 | 1.50E+05 | 2.76E+05 | 2.09E+05 | 3.00E+05 | 2.03E+05 | 1.48E+05 | 1.45E+05 | 1.56E+05 | 1.55E+05 |
| PS(17:0_20:3) | 1.44E+04 | 2.16E+04 | 5.14E+04 | 2.73E+04 | 3.31E+04 | 2.57E+04 | 2.74E+04 | 3.89E+04 | 3.51E+04 | 2.60E+04 | 3.33E+04 | 2.58E+04 | 3.90E+04 | 3.50E+04 | 4.06E+04 | 3.04E+04 |
| PS(20:0_18:1) | 3.10E+05 | 4.46E+05 | 8.13E+05 | 4.51E+05 | 4.48E+05 | 3.14E+05 | 5.24E+05 | 6.82E+05 | 4.93E+05 | 3.86E+05 | 4.96E+05 | 4.19E+05 | 6.24E+05 | 5.92E+05 | 7.20E+05 | 6.61E+05 |
| PS(18:1_20:1) | 2.80E+05 | 3.59E+05 | 5.79E+05 | 3.62E+05 | 4.34E+05 | 2.22E+05 | 2.65E+05 | 4.83E+05 | 4.63E+05 | 3.42E+05 | 4.40E+05 | 3.48E+05 | 3.32E+05 | 4.02E+05 | 4.03E+05 | 3.76E+05 |
| PS(18:0_20:3) | 8.43E+05 | 9.75E+05 | 1.75E+06 | 1.14E+06 | 1.47E+06 | 9.76E+05 | 1.19E+06 | 1.84E+06 | 1.59E+06 | 1.15E+06 | 1.48E+06 | 1.23E+06 | 1.76E+06 | 1.46E+06 | 1.66E+06 | 1.42E+06 |
| PS(18:1_20:3) | 1.56E+04 | 5.52E+04 | 1.45E+05 | 7.01E+04 | 3.58E+04 | 2.18E+04 | 2.52E+04 | 9.21E+04 | 6.75E+04 | 9.89E+04 | 7.13E+04 | 7.89E+04 | 4.83E+04 | 2.25E+04 | 1.59E+04 | 7.14E+04 |
| PS(18:0_20:4)_Rt9.55 | 2.32E+05 | 2.74E+05 | 4.69E+05 | 2.59E+05 | 2.94E+05 | 2.25E+05 | 2.55E+05 | 4.95E+05 | 4.50E+05 | 4.61E+05 | 4.56E+05 | 4.79E+05 | 3.11E+05 | 2.94E+05 | 3.52E+05 | 2.81E+05 |
| PS(18:0_20:4)_Rt10.7 | 4.41E+05 | 6.87E+05 | 9.41E+05 | 6.05E+05 | 5.69E+05 | 4.41E+05 | 5.46E+05 | 7.55E+05 | 7.81E+05 | 7.22E+05 | 8.25E+05 | 7.82E+05 | 9.49E+05 | 9.43E+05 | 9.87E+05 | 9.91E+05 |
| PS(18:0_20:5) | 9.39E+03 | 3.27E+04 | 7.26E+04 | 1.64E+04 | 1.42E+04 | 7.58E+03 | 1.50E+04 | 2.99E+04 | 1.14E+05 | 1.14E+05 | 8.82E+04 | 9.64E+04 | 3.70E+03 | 6.23E+03 | 1.22E+04 | 6.23E+03 |
| PS(18:1_20:4) | 6.06E+04 | 9.91E+04 | 1.25E+05 | 8.38E+04 | 5.25E+04 | 3.90E+04 | 4.87E+04 | 5.60E+04 | 1.44E+05 | 1.09E+05 | 1.34E+05 | 1.07E+05 | 5.75E+04 | 6.02E+04 | 7.26E+04 | 7.21E+04 |
| PS(16:0_22:6) | 1.57E+04 | 3.66E+04 | 4.64E+04 | 2.96E+04 | 5.73E+04 | 4.89E+04 | 8.40E+04 | 1.12E+05 | 4.79E+04 | 4.49E+04 | 4.72E+04 | 4.06E+04 | 4.98E+04 | 4.38E+04 | 5.22E+04 | 5.27E+04 |
| PS(17:0_22:5) | 1.39E+04 | 1.69E+04 | 1.44E+04 | 9.98E+03 | 3.64E+04 | 2.29E+04 | 2.77E+04 | 6.39E+04 | 1.39E+04 | 8.23E+03 | 9.58E+03 | 9.29E+03 | 4.77E+04 | 3.65E+04 | 4.47E+04 | 3.31E+04 |
| PS(22:0_18:1) | 2.87E+05 | 4.91E+05 | 8.99E+05 | 4.21E+05 | 3.45E+05 | 2.59E+05 | 4.80E+05 | 5.82E+05 | 4.36E+05 | 3.65E+05 | 4.47E+05 | 4.44E+05 | 5.29E+05 | 4.80E+05 | 6.22E+05 | 4.84E+05 |
| PS(18:1_22:1) | 2.50E+05 | 4.00E+05 | 6.52E+05 | 3.61E+05 | 4.44E+05 | 3.08E+05 | 3.28E+05 | 5.92E+05 | 4.04E+05 | 3.55E+05 | 4.07E+05 | 3.76E+05 | 2.66E+05 | 2.24E+05 | 2.71E+05 | 3.02E+05 |
| PS(18:0_22:4) | 1.39E+05 | 1.60E+05 | 2.69E+05 | 1.79E+05 | 4.48E+05 | 4.59E+05 | 4.53E+05 | 9.67E+05 | 2.80E+05 | 2.51E+05 | 3.28E+05 | 2.32E+05 | 5.48E+05 | 4.93E+05 | 4.52E+05 | 3.83E+05 |
| PS(18:0_22:5) | 2.02E+05 | 1.70E+05 | 3.13E+05 | 2.01E+05 | 1.01E+06 | 7.87E+05 | 7.72E+05 | 1.78E+06 | 2.18E+05 | 2.03E+05 | 2.58E+05 | 1.94E+05 | 2.01E+06 | 1.70E+06 | 2.02E+06 | 1.54E+06 |
| PS(18:1_22:5) | 4.37E+04 | 4.92E+04 | 7.56E+04 | 3.72E+04 | 1.00E+05 | 7.51E+04 | 8.21E+04 | 1.59E+05 | 7.51E+04 | 7.94E+04 | 6.51E+04 | 5.97E+04 | 1.04E+05 | 1.20E+05 | 1.13E+05 | 1.20E+05 |
| PS(18:0_22:6) | 2.45E+05 | 2.73E+05 | 3.84E+05 | 3.00E+05 | 1.26E+06 | 1.14E+06 | 1.54E+06 | 2.31E+06 | 3.20E+05 | 2.36E+05 | 3.46E+05 | 2.69E+05 | 2.66E+06 | 2.27E+06 | 2.94E+06 | 2.04E+06 |

| | | | | | | | | | | | | | | | | |
|---|---|---|---|---|---|---|---|---|---|---|---|---|---|---|---|---|
| PS(18:1_22:6) | 1.70E+04 | 4.48E+04 | 5.02E+04 | 2.98E+04 | 7.84E+04 | 6.15E+04 | 7.77E+04 | 1.42E+05 | 5.80E+04 | 5.19E+04 | 4.74E+04 | 4.75E+04 | 7.60E+04 | 5.05E+04 | 7.15E+04 | 7.40E+04 |
| PS(20:3_20:4) | 2.90E+04 | 2.90E+04 | 6.32E+04 | 3.16E+04 | 3.56E+04 | 2.81E+04 | 2.38E+04 | 4.40E+04 | 5.30E+04 | 4.89E+04 | 4.28E+04 | 5.48E+04 | 3.40E+04 | 3.21E+04 | 3.61E+04 | 2.71E+04 |
| PS(24:0_18:1) | 1.38E+05 | 3.28E+05 | 5.49E+05 | 2.95E+05 | 1.13E+05 | 7.35E+04 | 1.89E+05 | 2.17E+05 | 2.21E+05 | 2.06E+05 | 2.32E+05 | 2.56E+05 | 4.68E+05 | 2.85E+05 | 5.57E+05 | 2.77E+05 |
| PS(18:1_24:1) | 1.21E+05 | 2.61E+05 | 4.42E+05 | 2.04E+05 | 1.37E+05 | 1.10E+05 | 1.33E+05 | 1.95E+05 | 2.39E+05 | 2.04E+05 | 1.92E+05 | 2.62E+05 | 1.47E+05 | 1.32E+05 | 1.74E+05 | 1.59E+05 |
| PS(20:3_22:6)_Rt9.25 | 2.68E+04 | 2.58E+04 | 3.80E+04 | 3.23E+04 | 1.11E+05 | 1.18E+05 | 1.66E+05 | 1.54E+05 | 3.10E+04 | 2.50E+04 | 3.62E+04 | 1.47E+04 | 9.81E+02 | 1.77E+02 | 5.19E+02 | 9.56E+02 |
| PS(20:3_22:6)_Rt9.58 | 6.38E+04 | 1.06E+05 | 1.29E+05 | 8.49E+04 | 1.72E+05 | 1.94E+05 | 1.67E+05 | 2.83E+05 | 1.19E+05 | 1.16E+05 | 1.22E+05 | 1.23E+05 | 2.94E+05 | 2.69E+05 | 3.00E+05 | 2.28E+05 |
| PS(18:1_26:3) | 7.99E+04 | 1.35E+05 | 2.36E+05 | 1.05E+05 | 7.99E+04 | 5.39E+04 | 6.96E+04 | 9.59E+04 | 1.29E+05 | 1.00E+05 | 1.45E+05 | 1.35E+05 | 7.99E+04 | 7.35E+04 | 9.19E+04 | 7.67E+04 |
| SM(d34:0) | 8.41E+05 | 1.14E+06 | 2.27E+06 | 1.24E+06 | 1.28E+05 | 9.63E+04 | 7.80E+04 | 1.84E+05 | 1.45E+06 | 9.90E+05 | 1.48E+06 | 7.88E+05 | 1.15E+06 | 1.00E+06 | 1.10E+06 | 9.86E+05 |
| SM(d18:1_17:0) | 2.75E+04 | 5.68E+04 | 1.11E+05 | 4.61E+04 | 1.48E+05 | 1.18E+05 | 1.18E+05 | 2.37E+05 | 7.76E+04 | 8.07E+04 | 5.85E+04 | 6.31E+04 | 1.70E+05 | 1.68E+05 | 1.75E+05 | 1.97E+05 |
| SM(d18:1_22:0) | 2.17E+05 | 3.97E+05 | 6.90E+05 | 3.50E+05 | 5.60E+05 | 4.50E+05 | 5.31E+05 | 1.02E+06 | 6.46E+05 | 4.73E+05 | 6.44E+05 | 4.29E+05 | 1.26E+06 | 1.08E+06 | 1.33E+06 | 1.45E+06 |
| SM(d41:1) | 8.80E+04 | 1.76E+05 | 3.16E+05 | 1.45E+05 | 1.98E+05 | 1.42E+05 | 1.93E+05 | 3.83E+05 | 1.60E+05 | 1.99E+05 | 2.30E+05 | 1.68E+05 | 4.93E+05 | 4.67E+05 | 4.67E+05 | 5.34E+05 |
| SM(d42:1) | 6.87E+05 | 1.69E+06 | 2.98E+06 | 1.13E+06 | 1.60E+06 | 1.13E+06 | 1.67E+06 | 3.07E+06 | 2.43E+06 | 1.97E+06 | 2.28E+06 | 1.79E+06 | 3.57E+06 | 2.92E+06 | 3.44E+06 | 3.54E+06 |
| SM(d43:1) | 2.13E+04 | 3.42E+04 | 1.01E+05 | 4.26E+04 | 3.40E+04 | 1.98E+04 | 3.44E+04 | 6.78E+04 | 6.39E+04 | 7.35E+04 | 6.67E+04 | 6.33E+04 | 1.06E+05 | 1.06E+05 | 1.36E+05 | 1.14E+05 |
| SM(d44:2) | 4.87E+04 | 2.05E+05 | 3.39E+05 | 1.54E+05 | 6.53E+04 | 4.00E+04 | 3.28E+04 | 2.61E+04 | 3.38E+05 | 2.52E+05 | 1.78E+05 | 2.46E+05 | 1.53E+05 | 1.17E+05 | 2.34E+05 | 1.34E+05 |

| LipidID | HCT116 shNPC2 Avg | HCT116 SCR Avg | FC (shNPC2 /SCR HCT116) | p val HCT116 | FDR p val HCT116 | KM12SM shNPC2 Avg | KM12SM SCR Avg | FC (shNPC2 /SCR KM12SM) | p value KM12SM | FDR p val KM12SM | Blank | QC #1 | QC #2 | QC #3 | QC #4 | QC #5 |
|---|---|---|---|---|---|---|---|---|---|---|---|---|---|---|---|---|
| Cer(d18:0_14:0) | 3.27E+04 | 9.97E+03 | 3.28E+00 | 5.55E-02 | 2.38E-01 | 2.58E+04 | 1.88E+04 | 1.37E+00 | 2.39E-01 | 2.83E-01 | NP | 1.81E+04 | 2.05E+04 | 2.22E+04 | 1.92E+04 | 2.20E+04 |
| Cer(d18:1_14:0) | 2.72E+05 | 4.18E+05 | 6.51E-01 | 1.54E-01 | 3.53E-01 | 3.07E+05 | 3.52E+05 | 8.72E-01 | 1.21E-01 | 1.54E-01 | NP | 2.69E+05 | 3.09E+05 | 3.21E+05 | 3.32E+05 | 3.30E+05 |
| Cer(d18:2_14:0) | 1.70E+04 | 1.59E+04 | 1.07E+00 | 8.85E-01 | 9.43E-01 | 2.02E+04 | 1.38E+04 | 1.46E+00 | 1.58E-02 | 2.71E-02 | 2.47E-02 | 1.14E+04 | 1.43E+04 | 1.46E+04 | 1.58E+04 | 1.67E+04 |
| Cer(d18:1_15:0) | 1.69E+05 | 3.07E+05 | 5.49E-01 | 1.30E-01 | 3.29E-01 | 1.69E+05 | 2.61E+05 | 6.47E-01 | 1.47E-02 | 2.55E-02 | NP | 1.77E+05 | 1.70E+05 | 2.18E+05 | 2.13E+05 | 2.41E+05 |
| Cer(d18:1_16:0) | 3.35E+06 | 6.04E+06 | 5.55E-01 | 2.94E-02 | 1.93E-01 | 3.55E+06 | 5.97E+06 | 5.95E-01 | 4.98E-04 | 1.61E-03 | NP | 4.08E+06 | 4.72E+06 | 4.53E+06 | 4.85E+06 | 4.66E+06 |
| Cer(d18:2_16:0) | 4.99E+05 | 4.45E+05 | 1.12E+00 | 7.15E-01 | 8.58E-01 | 4.87E+05 | 3.35E+05 | 1.45E+00 | 8.76E-03 | 1.68E-02 | NP | 3.79E+05 | 4.02E+05 | 4.17E+05 | 4.23E+05 | 4.26E+05 |
| Cer(d18:1_17:0) | 5.10E+04 | 2.20E+05 | 2.32E-01 | 8.69E-03 | 1.75E-01 | 4.73E+04 | 9.48E+04 | 4.99E-01 | 4.94E-04 | 1.61E-03 | NP | 8.18E+04 | 9.07E+04 | 9.54E+04 | 1.03E+05 | 1.03E+05 |
| Cer(d18:1_18:0) | 1.34E+05 | 1.45E+06 | 9.27E-02 | 6.90E-04 | 5.28E-02 | 1.25E+05 | 1.65E+05 | 7.56E-01 | 9.51E-02 | 1.24E-01 | 0.00E+00 | 4.01E+05 | 4.60E+05 | 4.34E+05 | 4.49E+05 | 4.68E+05 |
| Cer(d18:2_18:0) | 2.93E+04 | 2.63E+05 | 1.12E-01 | 1.14E-05 | 2.17E-03 | 3.09E+04 | 4.29E+04 | 7.20E-01 | 1.47E-02 | 2.55E-02 | NP | 8.46E+04 | 7.59E+04 | 9.36E+04 | 9.99E+04 | 8.91E+04 |
| Cer(d18:1_20:0) | 4.99E+04 | 1.34E+05 | 3.72E-01 | 3.86E-02 | 2.11E-01 | 5.00E+04 | 7.39E+04 | 6.76E-01 | 3.24E-02 | 4.94E-02 | NP | 5.86E+04 | 6.56E+04 | 7.09E+04 | 6.90E+04 | 6.76E+04 |
| Cer(d18:0_22:0) | 4.68E+04 | 2.05E+04 | 2.28E+00 | 1.60E-02 | 1.93E-01 | 3.40E+04 | 5.06E+04 | 6.71E-01 | 1.21E-01 | 1.54E-01 | 6.59E+02 | 3.38E+04 | 3.72E+04 | 3.48E+04 | 3.66E+04 | 3.67E+04 |
| Cer(d18:1_22:0) | 5.00E+05 | 1.00E+06 | 4.98E-01 | 1.25E-01 | 3.25E-01 | 5.27E+05 | 7.83E+05 | 6.72E-01 | 4.18E-02 | 6.07E-02 | 4.05E-01 | 5.77E+05 | 5.82E+05 | 6.41E+05 | 6.31E+05 | 6.02E+05 |
| Cer(d16:1_24:1) | 7.36E+04 | 1.44E+05 | 5.11E-01 | 1.78E-01 | 3.78E-01 | 8.14E+04 | 9.00E+04 | 9.04E-01 | 5.79E-01 | 6.21E-01 | NP | 7.49E+04 | 8.04E+04 | 8.21E+04 | 8.94E+04 | 8.07E+04 |
| Cer(d18:2_22:0) | 7.94E+04 | 1.08E+05 | 7.36E-01 | 4.07E-01 | 6.25E-01 | 8.59E+04 | 7.91E+04 | 1.09E+00 | 5.93E-01 | 6.29E-01 | NP | 6.94E+04 | 7.44E+04 | 8.25E+04 | 7.65E+04 | 8.03E+04 |
| Cer(d18:1_23:0) | 1.43E+05 | 2.13E+05 | 6.75E-01 | 2.95E-01 | 5.15E-01 | 1.36E+05 | 2.01E+05 | 6.78E-01 | 1.15E-02 | 2.08E-02 | NP | 1.39E+05 | 1.51E+05 | 1.42E+05 | 1.55E+05 | 1.46E+05 |
| Cer(d18:1_23:1) | 5.81E+04 | 1.06E+05 | 5.47E-01 | 1.94E-01 | 3.91E-01 | 6.70E+04 | 8.36E+04 | 8.01E-01 | 1.33E-01 | 1.66E-01 | NP | 5.59E+04 | 6.38E+04 | 6.81E+04 | 7.32E+04 | 7.18E+04 |
| Cer(d18:0_24:0) | 1.40E+05 | 2.58E+04 | 5.44E+00 | 6.01E-03 | 1.43E-01 | 9.99E+04 | 1.68E+05 | 5.96E-01 | 6.99E-02 | 9.40E-02 | 7.58E-01 | 8.72E+04 | 9.53E+04 | 9.76E+04 | 1.01E+05 | 1.05E+05 |
| Cer(d18:0_24:1) | 1.45E+05 | 3.50E+04 | 4.15E+00 | 2.44E-02 | 1.93E-01 | 1.13E+05 | 1.31E+05 | 8.63E-01 | 6.67E-01 | 6.96E-01 | NP | 8.80E+04 | 9.53E+04 | 9.72E+04 | 1.01E+05 | 9.89E+04 |
| Cer(d18:1_24:0) | 2.15E+06 | 2.30E+06 | 9.36E-01 | 8.59E-01 | 9.32E-01 | 2.31E+06 | 2.96E+06 | 7.81E-01 | 1.40E-01 | 1.72E-01 | NP | 2.08E+06 | 2.07E+06 | 2.20E+06 | 2.12E+06 | 2.05E+06 |
| Cer(d18:1_24:1) | 1.71E+06 | 2.36E+06 | 7.25E-01 | 4.02E-01 | 6.22E-01 | 1.96E+06 | 2.36E+06 | 8.28E-01 | 1.89E-01 | 2.28E-01 | NP | 1.84E+06 | 1.70E+06 | 2.00E+06 | 1.94E+06 | 1.94E+06 |
| Cer(d18:2_24:0) | 6.07E+05 | 3.63E+05 | 1.67E+00 | 3.14E-01 | 5.40E-01 | 6.27E+05 | 4.36E+05 | 1.44E+00 | 2.25E-02 | 3.65E-02 | NP | 4.28E+05 | 4.29E+05 | 4.56E+05 | 4.47E+05 | 4.66E+05 |
| Cer(d18:2_24:1) | 3.99E+05 | 3.66E+05 | 1.09E+00 | 8.53E-01 | 9.32E-01 | 4.34E+05 | 2.72E+05 | 1.60E+00 | 2.74E-02 | 4.29E-02 | NP | 3.05E+05 | 3.17E+05 | 3.39E+05 | 3.54E+05 | 3.37E+05 |
| Cer(d19:1_24:0) | 8.59E+04 | 5.49E+04 | 1.57E+00 | 3.45E-01 | 5.68E-01 | 8.30E+04 | 1.50E+05 | 5.55E-01 | 2.75E-03 | 6.26E-03 | NP | 7.78E+04 | 8.82E+04 | 8.82E+04 | 8.32E+04 | 8.96E+04 |
| Cer(d19:1_24:1) | 7.90E+04 | 6.25E+04 | 1.26E+00 | 6.44E-01 | 8.17E-01 | 9.56E+04 | 1.37E+05 | 6.97E-01 | 2.77E-03 | 6.27E-03 | NP | 7.45E+04 | 7.65E+04 | 7.31E+04 | 7.65E+04 | 8.12E+04 |
| Cer(d20:1_24:0) | 2.28E+05 | 6.43E+04 | 3.54E+00 | 6.09E-02 | 2.38E-01 | 2.45E+05 | 3.80E+05 | 6.44E-01 | 1.02E-02 | 1.92E-02 | NP | 2.00E+05 | 2.21E+05 | 2.20E+05 | 2.26E+05 | 2.23E+05 |
| Cer(d20:1_24:1) | 2.32E+05 | 7.67E+04 | 3.02E+00 | 6.55E-02 | 2.40E-01 | 3.08E+05 | 2.83E+05 | 1.09E+00 | 3.87E-01 | 4.33E-01 | NP | 1.90E+05 | 1.84E+05 | 2.01E+05 | 2.00E+05 | 1.95E+05 |
| Cer(t16:0_18:1) | 1.39E+04 | 3.16E+04 | 4.42E-01 | 9.62E-04 | 6.12E-02 | 1.23E+04 | 1.13E+05 | 1.09E-01 | 5.56E-06 | 1.25E-04 | 1.79E+03 | 4.46E+04 | 4.45E+04 | 4.21E+04 | 5.21E+04 | 4.43E+04 |
| Cer(t18:1_23:0) | 2.87E+03 | 7.25E+03 | 3.96E-01 | 8.37E-03 | 1.75E-01 | 4.51E+03 | 3.33E+04 | 1.35E-01 | 2.56E-06 | 7.54E-05 | NP | 9.00E+03 | 1.28E+04 | 1.15E+04 | 1.03E+04 | 1.03E+04 |
| Cer(t18:0_24:1) | 6.89E+03 | 1.08E+04 | 6.41E-01 | 1.47E-01 | 3.47E-01 | 5.84E+03 | 1.67E+05 | 3.49E-02 | 2.37E-08 | 9.05E-06 | 2.74E-02 | 3.41E+04 | 3.53E+04 | 4.97E+04 | 3.64E+04 | 3.48E+04 |

| | | | | | | | | | | | | | | | | |
|---|---|---|---|---|---|---|---|---|---|---|---|---|---|---|---|---|
| CL(14:0_16:1_18:1_18:1) | 5.60E+04 | 7.24E+04 | 7.74E-01 | 3.30E-01 | 5.53E-01 | 4.31E+04 | 7.02E+04 | 6.14E-01 | 3.83E-03 | 8.36E-03 | NP | 5.69E+04 | 5.64E+04 | 5.30E+04 | 5.21E+04 | 5.57E+04 |
| CL(16:1_16:1_16:1_18:1) | 9.71E+04 | 8.26E+04 | 1.18E+00 | 4.61E-01 | 6.72E-01 | 6.87E+04 | 6.71E+04 | 1.02E+00 | 9.16E-01 | 9.23E-01 | NP | 7.38E+04 | 7.14E+04 | 7.16E+04 | 7.03E+04 | 7.35E+04 |
| CL(16:0_16:0_18:1_18:1) | 7.04E+04 | 7.31E+04 | 9.63E-01 | 9.06E-01 | 9.54E-01 | 4.84E+04 | 9.14E+04 | 5.29E-01 | 2.41E-02 | 3.88E-02 | 9.31E+02 | 7.52E+04 | 7.40E+04 | 5.88E+04 | 5.07E+04 | 4.92E+04 |
| CL(16:0_16:1_18:1_18:1) | 1.14E+05 | 1.28E+05 | 8.95E-01 | 6.87E-01 | 8.38E-01 | 9.26E+04 | 1.34E+05 | 6.91E-01 | 1.92E-02 | 3.21E-02 | NP | 1.09E+05 | 1.00E+05 | 1.05E+05 | 1.04E+05 | 1.13E+05 |
| CL(16:1_16:1_18:1_18:1) | 2.62E+05 | 2.77E+05 | 9.48E-01 | 7.60E-01 | 8.77E-01 | 2.26E+05 | 3.06E+05 | 7.39E-01 | 6.32E-02 | 8.69E-02 | NP | 2.36E+05 | 2.34E+05 | 2.50E+05 | 2.37E+05 | 2.49E+05 |
| CL(16:1_16:1_18:1_18:2) | 1.57E+05 | 1.34E+05 | 1.18E+00 | 5.28E-01 | 7.31E-01 | 1.25E+05 | 9.68E+04 | 1.29E+00 | 1.13E-01 | 1.45E-01 | NP | 1.02E+05 | 1.15E+05 | 1.21E+05 | 1.14E+05 | 1.15E+05 |
| CL(16:1_17:1_18:1_18:1) | 3.88E+04 | 8.19E+04 | 4.74E-01 | 5.47E-02 | 2.38E-01 | 2.66E+04 | 6.81E+04 | 3.90E-01 | 5.80E-03 | 1.19E-02 | NP | 4.80E+04 | 5.48E+04 | 5.33E+04 | 5.05E+04 | 4.74E+04 |
| CL(16:1_17:1_18:1_18:2) | 2.44E+04 | 3.98E+04 | 6.13E-01 | 1.73E-01 | 3.75E-01 | 1.90E+04 | 2.75E+04 | 6.91E-01 | 1.35E-01 | 1.68E-01 | NP | 1.97E+04 | 2.33E+04 | 2.39E+04 | 2.49E+04 | 2.51E+04 |
| CL(16:0_18:1_18:1_18:1) | 5.49E+04 | 8.88E+04 | 6.18E-01 | 3.79E-02 | 2.10E-01 | 3.96E+04 | 7.23E+04 | 5.47E-01 | 4.11E-03 | 8.72E-03 | 5.52E-03 | 7.16E+04 | 5.67E+04 | 5.51E+04 | 6.12E+04 | 5.91E+04 |
| CL(16:1_18:1_18:1_18:1) | 1.94E+05 | 3.10E+05 | 6.27E-01 | 4.45E-02 | 2.16E-01 | 1.36E+05 | 3.13E+05 | 4.35E-01 | 4.62E-04 | 1.54E-03 | 5.04E-03 | 2.25E+05 | 1.73E+05 | 2.25E+05 | 2.38E+05 | 2.11E+05 |
| CL(16:1_18:1_18:1_18:2) | 3.19E+05 | 3.72E+05 | 8.57E-01 | 5.62E-01 | 7.57E-01 | 2.77E+05 | 3.70E+05 | 7.47E-01 | 5.37E-02 | 7.55E-02 | 8.81E-03 | 3.00E+05 | 2.75E+05 | 3.04E+05 | 2.95E+05 | 3.09E+05 |
| CL(16:1_18:1_18:2_18:2) | 1.29E+05 | 1.14E+05 | 1.13E+00 | 7.09E-01 | 8.57E-01 | 1.03E+05 | 8.93E+04 | 1.16E+00 | 2.43E-01 | 2.87E-01 | NP | 9.60E+04 | 1.02E+05 | 9.35E+04 | 9.61E+04 | 9.53E+04 |
| CL(17:1_18:1_18:1_18:1) | 1.65E+04 | 5.90E+04 | 2.80E-01 | 5.14E-03 | 1.40E-01 | 9.52E+03 | 5.00E+04 | 1.91E-01 | 3.73E-05 | 2.57E-04 | 5.81E-03 | 3.47E+04 | 2.84E+04 | 3.15E+04 | 2.85E+04 | 3.29E+04 |
| CL(17:1_18:1_18:1_18:2) | 3.00E+04 | 6.39E+04 | 4.69E-01 | 5.27E-02 | 2.37E-01 | 2.13E+04 | 6.16E+04 | 3.46E-01 | 2.00E-03 | 4.75E-03 | 9.05E+02 | 4.37E+04 | 3.65E+04 | 4.27E+04 | 3.75E+04 | 4.13E+04 |
| CL(18:1_18:1_18:1_18:1) | 3.53E+04 | 1.21E+05 | 2.92E-01 | 3.56E-03 | 1.16E-01 | 1.84E+04 | 9.23E+04 | 1.99E-01 | 2.93E-05 | 2.27E-04 | 4.57E-03 | 6.37E+04 | 6.15E+04 | 6.50E+04 | 6.48E+04 | 5.79E+04 |
| CL(18:1_18:1_18:1_18:2) | 9.87E+04 | 2.16E+05 | 4.58E-01 | 2.74E-02 | 1.93E-01 | 8.82E+04 | 2.06E+05 | 4.29E-01 | 5.21E-05 | 3.16E-04 | 6.29E-03 | 1.53E+05 | 1.46E+05 | 1.46E+05 | 1.44E+05 | 1.50E+05 |
| CL(18:1_18:1_18:2_18:2) | 1.56E+05 | 1.94E+05 | 8.00E-01 | 5.59E-01 | 7.57E-01 | 1.30E+05 | 2.24E+05 | 5.80E-01 | 1.06E-02 | 1.98E-02 | NP | 1.50E+05 | 1.56E+05 | 1.50E+05 | 1.56E+05 | 1.44E+05 |
| CL(18:1_18:2_18:2_18:2) | 6.53E+04 | 5.72E+04 | 1.14E+00 | 7.58E-01 | 8.77E-01 | 4.82E+04 | 5.49E+04 | 8.77E-01 | 5.22E-01 | 5.68E-01 | NP | 4.63E+04 | 4.93E+04 | 1.79E+04 | 4.40E+04 | 5.26E+04 |
| CL(16:1_18:2_18:2_20:4) | 5.20E+04 | 4.74E+04 | 1.10E+00 | 7.84E-01 | 8.89E-01 | 3.88E+04 | 3.62E+04 | 1.07E+00 | 5.52E-01 | 5.95E-01 | NP | 3.65E+04 | 3.83E+04 | 3.51E+04 | 3.83E+04 | 3.96E+04 |
| CL(18:1_18:1_18:2_20:3) | 2.12E+04 | 4.05E+04 | 5.23E-01 | 1.18E-01 | 3.16E-01 | 1.55E+04 | 6.14E+04 | 2.52E-01 | 2.16E-04 | 8.51E-04 | 1.62E+00 | 3.21E+04 | 3.04E+04 | 3.36E+04 | 3.45E+04 | 2.55E+04 |
| DGDG(16:0_16:1) | 1.40E+04 | 1.34E+03 | 1.04E+01 | 2.73E-05 | 3.48E-03 | 2.33E+04 | 6.30E+04 | 3.70E-01 | 2.03E-04 | 8.26E-04 | 1.09E+03 | 2.15E+04 | 2.45E+04 | 2.98E+04 | 2.38E+04 | 2.45E+04 |
| DGDG(18:0_16:1) | 9.57E+03 | #¡DIV/0! | #¡DIV/0! | #¡DIV/0! | 0.00E+00 | 2.61E+04 | 1.22E+05 | 2.14E-01 | 2.31E-06 | 7.54E-05 | NP | 3.97E+04 | 4.36E+04 | 4.10E+04 | 4.16E+04 | 4.65E+04 |

| | | | | | | | | | | | | | | | | |
|---|---|---|---|---|---|---|---|---|---|---|---|---|---|---|---|---|
| LPC(14:0) | 2.75E+04 | 4.82E+04 | 5.70E-01 | 1.93E-01 | 3.91E-01 | 2.40E+04 | 3.19E+04 | 7.52E-01 | 1.35E-01 | 1.68E-01 | NP | 2.16E+04 | 2.54E+04 | 2.49E+04 | 3.47E+04 | 3.99E+04 |
| LPC(15:0) | 1.25E+04 | 2.31E+04 | 5.42E-01 | 2.01E-01 | 4.00E-01 | 1.07E+04 | 1.40E+04 | 7.64E-01 | 2.08E-01 | 2.48E-01 | 2.41E+03 | 8.10E+03 | 1.14E+04 | 1.35E+04 | 1.45E+04 | 1.88E+04 |
| LPC(16:0) | 4.25E+05 | 6.69E+05 | 6.36E-01 | 3.24E-01 | 5.51E-01 | 4.76E+05 | 7.38E+05 | 6.45E-01 | 9.03E-04 | 2.41E-03 | 1.47E+03 | 4.08E+05 | 4.68E+05 | 4.71E+05 | 5.87E+05 | 5.13E+05 |
| LPC(16:1) | 6.39E+04 | 7.86E+04 | 8.13E-01 | 6.13E-01 | 7.94E-01 | 6.49E+04 | 9.75E+04 | 6.65E-01 | 1.78E-01 | 2.15E-01 | NP | 5.27E+04 | 5.58E+04 | 6.74E+04 | 8.00E+04 | 9.47E+04 |
| LPC(17:0) | 9.49E+03 | 1.97E+04 | 4.82E-01 | 1.17E-01 | 3.16E-01 | 1.05E+04 | 2.27E+04 | 4.64E-01 | 6.54E-04 | 1.91E-03 | NP | 1.33E+04 | 1.51E+04 | 1.36E+04 | 1.55E+04 | 1.48E+04 |
| LPC(17:1) | 1.11E+04 | 1.67E+04 | 6.65E-01 | 4.02E-01 | 6.22E-01 | 1.08E+04 | 1.49E+04 | 7.23E-01 | 6.04E-02 | 8.36E-02 | NP | 7.57E+03 | 1.03E+04 | 1.03E+04 | 1.34E+04 | 1.49E+04 |
| LPC(18:0) | 1.93E+05 | 2.60E+05 | 7.42E-01 | 4.11E-01 | 6.29E-01 | 2.41E+05 | 4.66E+05 | 5.16E-01 | 4.05E-03 | 8.64E-03 | 5.68E+02 | 2.54E+05 | 2.60E+05 | 2.55E+05 | 2.71E+05 | 2.76E+05 |
| LPC(18:1) | 3.09E+05 | 3.85E+05 | 8.01E-01 | 6.53E-01 | 8.20E-01 | 3.36E+05 | 4.97E+05 | 6.75E-01 | 5.04E-02 | 7.16E-02 | NP | 2.14E+05 | 2.72E+05 | 3.48E+05 | 4.09E+05 | 3.94E+05 |
| LPC(18:1)+D7:(s) | 1.28E+06 | 1.38E+06 | 9.33E-01 | 8.19E-01 | 9.15E-01 | 1.01E+06 | 1.03E+06 | 9.83E-01 | 8.13E-01 | 8.37E-01 | 8.98E+05 | 1.10E+06 | 1.08E+06 | 1.13E+06 | 1.12E+06 | 1.02E+06 |
| LPC(18:2) | 2.36E+04 | 2.46E+04 | 9.58E-01 | 9.32E-01 | 9.59E-01 | 2.11E+04 | 2.40E+04 | 8.79E-01 | 5.87E-01 | 6.28E-01 | NP | 1.32E+04 | 1.91E+04 | 2.35E+04 | 2.01E+04 | 3.02E+04 |
| LPC(20:0) | 9.92E+03 | 1.07E+04 | 9.26E-01 | 8.53E-01 | 9.32E-01 | 1.54E+04 | 1.90E+04 | 8.10E-01 | 9.89E-02 | 1.29E-01 | NP | 1.41E+04 | 1.24E+04 | 1.31E+04 | 1.25E+04 | 1.19E+04 |
| LPC(20:1) | 2.13E+04 | 2.66E+04 | 8.01E-01 | 5.63E-01 | 7.57E-01 | 2.29E+04 | 3.71E+04 | 6.18E-01 | 8.11E-03 | 1.57E-02 | NP | 2.18E+04 | 2.47E+04 | 2.27E+04 | 2.32E+04 | 2.62E+04 |
| LPC(20:4) | 2.96E+04 | 3.79E+04 | 7.81E-01 | 5.04E-01 | 7.23E-01 | 3.21E+04 | 2.85E+04 | 1.13E+00 | 1.10E-01 | 1.42E-01 | NP | 2.22E+04 | 2.36E+04 | 2.83E+04 | 3.58E+04 | 3.56E+04 |
| LPC(22:5) | 1.18E+04 | 1.29E+04 | 9.17E-01 | 7.56E-01 | 8.77E-01 | 1.28E+04 | 1.56E+04 | 8.21E-01 | 1.29E-01 | 1.62E-01 | NP | 9.49E+03 | 1.21E+04 | 1.32E+04 | 1.23E+04 | 1.41E+04 |
| LPC(22:6) | 1.53E+04 | 2.27E+04 | 6.74E-01 | 2.08E-01 | 4.08E-01 | 1.64E+04 | 2.28E+04 | 7.17E-01 | 2.53E-03 | 5.81E-03 | NP | 1.57E+04 | 1.71E+04 | 1.60E+04 | 1.88E+04 | 2.30E+04 |
| LPC(26:0) | 2.36E+04 | 1.93E+03 | 1.23E+01 | 2.92E-02 | 1.93E-01 | 2.81E+04 | 5.85E+04 | 4.79E-01 | 2.23E-04 | 8.71E-04 | NP | 2.32E+04 | 2.44E+04 | 2.41E+04 | 2.64E+04 | 3.05E+04 |
| LPC(26:1) | 2.09E+04 | 2.83E+03 | 7.39E+00 | 3.93E-02 | 2.11E-01 | 3.14E+04 | 2.63E+04 | 1.19E+00 | 3.13E-01 | 3.57E-01 | NP | 1.89E+04 | 1.10E+04 | 1.96E+04 | 2.19E+04 | 1.91E+04 |
| LPC(O-16:0) | 1.14E+04 | 1.07E+04 | 1.07E+00 | 8.69E-01 | 9.35E-01 | 1.16E+04 | 1.16E+05 | 9.96E-02 | 9.09E-07 | 5.59E-05 | NP | 3.28E+04 | 3.41E+04 | 3.34E+04 | 3.68E+04 | 3.97E+04 |
| LPC(O-16:1) | 3.32E+04 | 2.82E+04 | 1.18E+00 | 6.28E-01 | 8.03E-01 | 4.01E+04 | 3.87E+04 | 1.04E+00 | 8.20E-01 | 8.42E-01 | NP | 2.84E+04 | 2.86E+04 | 3.08E+04 | 3.02E+04 | 3.10E+04 |
| LPC(O-18:1) | 2.46E+04 | 1.65E+04 | 1.49E+00 | 3.26E-01 | 5.51E-01 | 3.02E+04 | 3.35E+04 | 9.02E-01 | 3.99E-01 | 4.45E-01 | NP | 2.35E+04 | 2.37E+04 | 2.38E+04 | 2.60E+04 | 2.62E+04 |
| LPE(16:0) | 1.45E+05 | 2.42E+05 | 6.01E-01 | 2.27E-01 | 4.30E-01 | 1.69E+05 | 1.33E+05 | 1.27E+00 | 9.53E-02 | 1.24E-01 | 3.32E+02 | 1.26E+05 | 1.25E+05 | 1.50E+05 | 1.66E+05 | 1.85E+05 |
| LPE(16:1) | 3.65E+04 | 3.22E+04 | 1.13E+00 | 7.20E-01 | 8.58E-01 | 3.64E+04 | 2.49E+04 | 1.46E+00 | 1.38E-01 | 1.70E-01 | NP | 1.68E+04 | 2.53E+04 | 3.01E+04 | 3.58E+04 | 4.31E+04 |
| LPE(17:0) | 1.07E+04 | 2.14E+04 | 5.00E-01 | 1.16E-01 | 3.16E-01 | 1.19E+04 | 1.56E+04 | 7.64E-01 | 1.28E-02 | 2.28E-02 | NP | 1.12E+04 | 1.08E+04 | 1.37E+04 | 1.51E+04 | 1.47E+04 |
| LPE(17:1) | 7.53E+03 | 9.01E+03 | 8.35E-01 | 6.76E-01 | 8.28E-01 | 7.36E+03 | 4.82E+03 | 1.53E+00 | 1.97E-02 | 3.29E-02 | NP | 3.84E+03 | 6.38E+03 | 5.81E+03 | 6.55E+03 | 7.15E+03 |
| LPE(18:0) | 3.81E+05 | 5.46E+05 | 6.97E-01 | 3.81E-01 | 6.02E-01 | 5.02E+05 | 5.24E+05 | 9.59E-01 | 7.53E-01 | 7.79E-01 | 9.72E+02 | 3.80E+05 | 4.22E+05 | 4.63E+05 | 5.00E+05 | 4.74E+05 |
| LPE(18:1) | 3.37E+05 | 3.23E+05 | 1.04E+00 | 9.11E-01 | 9.56E-01 | 3.65E+05 | 2.55E+05 | 1.43E+00 | 5.49E-02 | 7.67E-02 | 8.05E+02 | 1.77E+05 | 2.37E+05 | 2.94E+05 | 3.54E+05 | 3.61E+05 |
| LPE(18:1)+D7:(s) | 4.00E+05 | 4.45E+05 | 8.99E-01 | 7.42E-01 | 8.72E-01 | 3.51E+05 | 3.53E+05 | 9.95E-01 | 9.15E-01 | 9.23E-01 | 2.57E+05 | 3.49E+05 | 3.33E+05 | 3.52E+05 | 3.57E+05 | 3.60E+05 |
| LPE(18:2) | 1.66E+04 | 2.06E+04 | 8.08E-01 | 6.00E-01 | 7.88E-01 | 1.80E+04 | 1.41E+04 | 1.27E+00 | 2.78E-01 | 3.24E-01 | NP | 8.29E+03 | 1.20E+04 | 1.55E+04 | 1.67E+04 | 2.19E+04 |
| LPE(20:0) | 1.20E+04 | 1.23E+04 | 9.71E-01 | 9.40E-01 | 9.60E-01 | 1.28E+04 | 1.26E+04 | 1.02E+00 | 8.89E-01 | 9.05E-01 | NP | 1.05E+04 | 1.13E+04 | 1.03E+04 | 1.33E+04 | 1.17E+04 |
| LPE(20:1) | 1.73E+04 | 3.29E+04 | 5.25E-01 | 6.67E-02 | 2.40E-01 | 2.17E+04 | 1.61E+04 | 1.35E+00 | 4.90E-02 | 6.99E-02 | NP | 1.59E+04 | 1.62E+04 | 2.09E+04 | 2.24E+04 | 2.43E+04 |
| LPE(20:4) | 1.52E+05 | 1.86E+05 | 8.18E-01 | 6.70E-01 | 8.25E-01 | 1.33E+05 | 7.80E+04 | 1.71E+00 | 5.18E-02 | 7.33E-02 | NP | 8.73E+04 | 1.03E+05 | 1.45E+05 | 1.53E+05 | 1.66E+05 |
| LPE(22:5) | 3.03E+04 | 8.79E+04 | 3.44E-01 | 2.03E-01 | 4.01E-01 | 2.77E+04 | 4.50E+04 | 6.15E-01 | 3.16E-02 | 4.85E-02 | NP | 3.14E+04 | 5.04E+04 | 5.64E+04 | 5.30E+04 | 5.73E+04 |
| LPE(22:6) | 4.02E+04 | 1.25E+05 | 3.22E-01 | 1.19E-01 | 3.16E-01 | 4.13E+04 | 5.76E+04 | 7.18E-01 | 7.63E-02 | 1.02E-01 | NP | 4.16E+04 | 5.46E+04 | 5.98E+04 | 6.50E+04 | 7.83E+04 |

| | | | | | | | | | | | | | | | | |
|---|---|---|---|---|---|---|---|---|---|---|---|---|---|---|---|---|
| LPE(24:0) | 2.76E+04 | 1.42E+04 | 1.95E+00 | 1.21E-01 | 3.16E-01 | 3.22E+04 | 4.04E+04 | 7.96E-01 | 8.03E-02 | 1.07E-01 | NP | 2.66E+04 | 2.70E+04 | 2.75E+04 | 2.79E+04 | 2.65E+04 |
| LPE(24:1) | 1.40E+04 | 8.40E+03 | 1.67E+00 | 1.95E-01 | 3.91E-01 | 1.75E+04 | 9.42E+03 | 1.85E+00 | 1.68E-02 | 2.84E-02 | NP | 9.83E+03 | 9.77E+03 | 1.10E+04 | 1.08E+04 | 1.22E+04 |
| LPE(24:5) | 7.14E+03 | 2.10E+04 | 3.40E-01 | 1.16E-01 | 3.16E-01 | 5.70E+03 | 4.11E+03 | 1.39E+00 | 1.62E-01 | 1.96E-01 | NP | 6.62E+03 | 4.64E+03 | 1.01E+04 | 1.04E+04 | 9.93E+03 |
| LPI(18:0) | 9.07E+03 | 1.94E+04 | 4.67E-01 | 1.43E-01 | 3.42E-01 | 8.67E+03 | 9.90E+03 | 8.76E-01 | 5.75E-01 | 6.18E-01 | NP | 4.74E+03 | 7.62E+03 | 1.00E+04 | 1.24E+04 | 1.37E+04 |
| LPS(18:0) | 2.09E+04 | 2.69E+04 | 7.77E-01 | 3.51E-01 | 5.73E-01 | 1.98E+04 | 2.70E+04 | 7.33E-01 | 1.27E-01 | 1.61E-01 | 5.30E+03 | 1.48E+04 | 2.00E+04 | 1.86E+04 | 2.40E+04 | 2.97E+04 |
| PC(2:0_16:0) | 1.88E+02 | 9.51E+03 | 1.98E-02 | 3.40E-01 | 5.62E-01 | 3.49E+02 | 1.93E+02 | 1.81E+00 | 7.33E-01 | 7.61E-01 | NP | 1.71E+03 | 2.27E+03 | 2.90E+03 | 3.48E+03 | 1.66E+03 |
| PC(14:0_14:0) | 3.45E+05 | 8.23E+05 | 4.20E-01 | 4.49E-02 | 2.16E-01 | 2.89E+05 | 5.58E+05 | 5.19E-01 | 1.45E-04 | 6.74E-04 | NP | 4.40E+05 | 4.52E+05 | 4.52E+05 | 4.53E+05 | 4.56E+05 |
| PC(14:0_15:0) | 5.23E+04 | 2.38E+05 | 2.19E-01 | 1.14E-02 | 1.91E-01 | 4.48E+04 | 1.09E+05 | 4.10E-01 | 1.24E-04 | 6.07E-04 | NP | 9.04E+04 | 6.34E+04 | 8.98E+04 | 9.10E+04 | 8.68E+04 |
| PC(14:0_16:0) | 4.10E+06 | 4.64E+06 | 8.85E-01 | 6.55E-01 | 8.20E-01 | 4.13E+06 | 3.16E+06 | 1.31E+00 | 3.37E-02 | 5.10E-02 | 2.06E+03 | 3.70E+06 | 3.85E+06 | 3.51E+06 | 2.37E+05 | 4.89E+05 |
| PC(14:0_16:1) | 7.37E+05 | 1.39E+06 | 5.30E-01 | 8.26E-02 | 2.68E-01 | 7.69E+05 | 1.27E+06 | 6.06E-01 | 5.50E-02 | 7.67E-02 | NP | 1.01E+06 | 1.01E+06 | 1.04E+06 | 1.05E+06 | 1.05E+06 |
| PC(15:0_16:0) | 1.20E+05 | 2.93E+05 | 4.09E-01 | 4.58E-02 | 2.16E-01 | 7.64E+04 | 1.83E+05 | 4.18E-01 | 4.52E-03 | 9.49E-03 | NP | 1.99E+05 | 5.88E+04 | 6.44E+04 | 1.81E+05 | 1.42E+05 |
| PC(16:0_16:0) | 5.54E+06 | 4.72E+06 | 1.17E+00 | 5.07E-01 | 7.23E-01 | 5.98E+06 | 4.25E+06 | 1.41E+00 | 1.61E-02 | 2.74E-02 | 8.58E-01 | 4.41E+06 | 4.91E+06 | 4.79E+06 | 5.25E+06 | 4.73E+06 |
| PC(16:0_16:1) | 1.10E+07 | 1.40E+07 | 7.87E-01 | 2.76E-01 | 4.98E-01 | 1.20E+07 | 1.78E+07 | 6.75E-01 | 8.18E-03 | 1.58E-02 | 4.99E+03 | 1.28E+07 | 1.43E+07 | 1.30E+07 | 1.43E+07 | 1.40E+07 |
| PC(16:1_16:1) | 8.74E+05 | 9.61E+05 | 9.10E-01 | 7.85E-01 | 8.89E-01 | 1.09E+06 | 4.66E+05 | 2.34E+00 | 1.68E-02 | 2.84E-02 | 0.00E+00 | 8.93E+05 | 4.92E+05 | 8.71E+05 | 1.06E+06 | 9.05E+05 |
| PC(16:0_16:2) | 1.90E+05 | 3.54E+05 | 5.36E-01 | 8.36E-02 | 2.68E-01 | 1.68E+05 | 2.43E+05 | 6.93E-01 | 6.57E-03 | 1.32E-02 | 4.59E-02 | 2.49E+05 | 2.28E+05 | 2.07E+05 | 2.48E+05 | 2.43E+05 |
| PC(16:1_16:2) | 2.61E+04 | 5.98E+04 | 4.36E-01 | 5.51E-02 | 2.38E-01 | 2.13E+04 | 4.50E+04 | 4.73E-01 | 2.60E-02 | 4.13E-02 | NP | 4.27E+04 | 3.07E+04 | 4.54E+04 | 5.03E+04 | 3.46E+04 |
| PC(16:0_17:0) | 7.28E+04 | 1.85E+05 | 3.94E-01 | 1.88E-01 | 3.89E-01 | 7.94E+04 | 1.46E+05 | 5.44E-01 | 1.10E-02 | 2.04E-02 | 0.00E+00 | 8.09E+04 | 6.07E+04 | 1.15E+05 | 1.20E+05 | 1.50E+05 |
| PC(16:0_17:1) | 1.56E+06 | 3.36E+06 | 4.65E-01 | 2.98E-02 | 1.93E-01 | 1.75E+06 | 2.31E+06 | 7.54E-01 | 2.90E-02 | 4.50E-02 | 0.00E+00 | 2.18E+06 | 2.05E+06 | 2.12E+06 | 1.98E+06 | 2.05E+06 |
| PC(15:0_18:1)+D7:(s) | 4.67E+06 | 4.72E+06 | 9.88E-01 | 9.60E-01 | 9.68E-01 | 3.89E+06 | 3.84E+06 | 1.01E+00 | 8.31E-01 | 8.51E-01 | 3.69E+06 | 4.04E+06 | 3.89E+06 | 4.06E+06 | 3.97E+06 | 4.07E+06 |
| PC(16:0_18:0) | 1.29E+06 | 1.08E+06 | 1.20E+00 | 5.89E-01 | 7.81E-01 | 1.42E+06 | 1.93E+06 | 7.35E-01 | 4.90E-02 | 6.99E-02 | NP | 1.36E+06 | 1.35E+06 | 1.34E+06 | 1.41E+06 | 1.33E+06 |
| PC(16:0_18:1) | 3.18E+07 | 4.29E+07 | 7.41E-01 | 1.92E-01 | 3.91E-01 | 3.63E+07 | 4.38E+07 | 8.29E-01 | 4.52E-02 | 6.48E-02 | 1.61E+04 | 3.59E+07 | 3.76E+07 | 3.81E+07 | 3.74E+07 | 3.50E+07 |
| PC(16:1_18:2) | 3.25E+05 | 3.75E+05 | 8.64E-01 | 6.35E-01 | 8.09E-01 | 4.12E+05 | 2.80E+05 | 1.47E+00 | 2.85E-02 | 4.45E-02 | 4.13E-02 | 3.84E+05 | 3.87E+05 | 1.41E+05 | 4.59E+05 | 4.07E+05 |
| PC(16:0_18:3) | 1.87E+05 | 2.83E+05 | 6.61E-01 | 1.58E-01 | 3.53E-01 | 2.36E+05 | 2.94E+05 | 8.04E-01 | 1.26E-02 | 2.26E-02 | 4.89E+02 | 2.73E+05 | 2.16E+05 | 2.71E+05 | 1.82E+05 | 2.48E+05 |
| PC(14:0_20:4) | 8.23E+04 | 1.85E+05 | 4.45E-01 | 9.97E-02 | 2.91E-01 | 8.42E+04 | 8.92E+04 | 9.43E-01 | 5.96E-01 | 6.30E-01 | NP | 8.91E+04 | 9.03E+04 | 9.54E+04 | 9.58E+04 | 1.01E+05 |
| PC(14:0_20:5) | 4.48E+04 | 6.81E+04 | 6.58E-01 | 3.83E-01 | 6.02E-01 | 4.44E+04 | 2.06E+04 | 2.16E+00 | 2.57E-02 | 4.13E-02 | NP | 4.14E+04 | 4.42E+04 | 4.37E+04 | 3.95E+04 | 4.83E+04 |
| PC(17:0_18:0) | 2.06E+04 | 5.41E+04 | 3.82E-01 | 1.92E-02 | 1.93E-01 | 2.82E+04 | 1.30E+05 | 2.16E-01 | 4.05E-03 | 8.64E-03 | NP | 3.77E+04 | 3.42E+04 | 2.92E+04 | 4.60E+04 | 5.56E+04 |
| PC(17:0_18:1) | 1.15E+06 | 1.99E+06 | 5.77E-01 | 8.32E-02 | 2.68E-01 | 1.26E+06 | 2.67E+06 | 4.73E-01 | 2.34E-04 | 8.79E-04 | NP | 1.58E+06 | 1.52E+06 | 1.56E+06 | 1.53E+06 | 1.66E+06 |
| PC(17:1_18:1) | 9.20E+05 | 1.25E+06 | 7.38E-01 | 2.82E-01 | 5.03E-01 | 1.05E+06 | 7.52E+05 | 1.39E+00 | 2.24E-02 | 3.64E-02 | 0.00E+00 | 1.08E+06 | 9.55E+05 | 1.11E+06 | 1.13E+06 | 1.12E+06 |
| PC(17:1_18:2) | 8.62E+04 | 1.06E+05 | 8.14E-01 | 6.76E-01 | 8.28E-01 | 8.91E+04 | 6.26E+04 | 1.42E+00 | 2.96E-01 | 3.43E-01 | 0.00E+00 | 8.98E+04 | 9.50E+04 | 9.50E+04 | 9.48E+04 | 9.71E+04 |
| PC(15:0_20:5) | 7.52E+03 | 2.83E+04 | 2.66E-01 | 5.08E-02 | 2.34E-01 | 7.43E+03 | 2.16E+03 | 3.43E+00 | 1.16E-02 | 2.09E-02 | 0.00E+00 | 5.31E+03 | 8.40E+03 | 6.10E+03 | 7.65E+03 | 6.30E+03 |
| PC(18:0_18:1) | 6.38E+06 | 9.62E+06 | 6.63E-01 | 1.36E-01 | 3.36E-01 | 7.70E+06 | 1.75E+07 | 4.40E-01 | 3.97E-05 | 2.57E-04 | 1.97E+03 | 9.52E+06 | 9.43E+06 | 9.60E+06 | 1.00E+07 | 9.62E+06 |
| PC(18:1_18:2) | 1.66E+06 | 1.59E+06 | 1.04E+00 | 8.64E-01 | 9.35E-01 | 1.91E+06 | 1.19E+06 | 1.60E+00 | 9.54E-03 | 1.81E-02 | NP | 1.48E+06 | 1.50E+06 | 1.42E+06 | 1.43E+06 | 1.47E+06 |

| | | | | | | | | | | | | | | | | |
|---|---|---|---|---|---|---|---|---|---|---|---|---|---|---|---|---|
| PC(16:0_20:3) | 8.44E+05 | 1.08E+06 | 7.83E-01 | 4.26E-01 | 6.38E-01 | 1.13E+06 | 1.53E+06 | 7.40E-01 | 2.65E-02 | 4.19E-02 | NP | 1.15E+06 | 1.10E+06 | 1.12E+06 | 1.20E+06 | 1.13E+06 |
| PC(16:0_20:4) | 2.04E+06 | 2.47E+06 | 8.26E-01 | 5.55E-01 | 7.57E-01 | 2.32E+06 | 1.59E+06 | 1.45E+00 | 3.97E-03 | 8.61E-03 | NP | 1.79E+06 | 1.94E+06 | 1.89E+06 | 1.99E+06 | 1.97E+06 |
| PC(16:1_20:4) | 2.43E+05 | 2.76E+05 | 8.77E-01 | 7.67E-01 | 8.81E-01 | 2.38E+05 | 1.40E+05 | 1.70E+00 | 4.26E-03 | 8.99E-03 | 8.13E+03 | 2.61E+05 | 2.58E+05 | 1.63E+05 | 2.01E+05 | 2.05E+05 |
| PC(16:0_20:5) | 1.36E+06 | 1.16E+06 | 1.17E+00 | 7.26E-01 | 8.59E-01 | 1.78E+06 | 4.86E+05 | 3.67E+00 | 3.16E-05 | 2.32E-04 | 5.03E-02 | 1.10E+06 | 1.07E+06 | 1.12E+06 | 1.04E+06 | 1.04E+06 |
| PC(16:1_20:5) | 8.03E+04 | 1.59E+05 | 5.04E-01 | 1.37E-01 | 3.36E-01 | 7.87E+04 | 6.04E+04 | 1.30E+00 | 6.33E-02 | 8.69E-02 | 1.52E+03 | 7.82E+04 | 8.81E+04 | 8.08E+04 | 1.07E+05 | 9.10E+04 |
| PC(19:0_18:1) | 1.55E+05 | 2.09E+05 | 7.41E-01 | 3.34E-01 | 5.58E-01 | 1.57E+05 | 4.21E+05 | 3.72E-01 | 4.32E-04 | 1.47E-03 | NP | 2.02E+05 | 2.05E+05 | 2.25E+05 | 2.48E+05 | 2.24E+05 |
| PC(18:1_19:1) | 3.66E+05 | 4.89E+05 | 7.48E-01 | 2.77E-01 | 4.98E-01 | 4.13E+05 | 9.91E+05 | 4.17E-01 | 7.97E-06 | 1.41E-04 | 0.00E+00 | 5.25E+05 | 5.35E+05 | 5.40E+05 | 5.35E+05 | 5.45E+05 |
| PC(17:0_20:3) | 8.83E+04 | 1.11E+05 | 7.98E-01 | 4.45E-01 | 6.58E-01 | 9.51E+04 | 2.51E+05 | 3.79E-01 | 1.39E-04 | 6.56E-04 | 5.08E-03 | 1.22E+05 | 1.32E+05 | 1.13E+05 | 1.39E+05 | 1.19E+05 |
| PC(15:0_22:6) | 4.10E+04 | 6.21E+04 | 6.60E-01 | 4.37E-01 | 6.52E-01 | 4.47E+04 | 2.50E+04 | 1.79E+00 | 6.65E-03 | 1.33E-02 | 4.70E-01 | 3.88E+04 | 3.71E+04 | 4.14E+04 | 4.52E+04 | 3.78E+04 |
| PC(20:0_18:1) | 2.65E+05 | 3.33E+05 | 7.97E-01 | 4.45E-01 | 6.58E-01 | 2.85E+05 | 9.52E+05 | 3.00E-01 | 2.69E-05 | 2.23E-04 | NP | 3.80E+05 | 3.80E+05 | 4.11E+05 | 3.38E+05 | 4.12E+05 |
| PC(18:1_20:1) | 1.13E+06 | 1.45E+06 | 7.83E-01 | 2.61E-01 | 4.76E-01 | 1.30E+06 | 2.63E+06 | 4.93E-01 | 5.61E-04 | 1.71E-03 | NP | 1.52E+06 | 1.55E+06 | 1.63E+06 | 1.60E+06 | 1.42E+06 |
| PC(18:1_20:2) | 4.83E+05 | 5.82E+05 | 8.30E-01 | 5.15E-01 | 7.24E-01 | 5.45E+05 | 9.24E+05 | 5.90E-01 | 4.57E-03 | 9.50E-03 | NP | 6.25E+05 | 6.59E+05 | 5.83E+05 | 5.99E+05 | 5.80E+05 |
| PC(18:0_20:3) | 4.74E+05 | 6.30E+05 | 7.52E-01 | 4.18E-01 | 6.30E-01 | 5.20E+05 | 1.07E+06 | 4.86E-01 | 1.91E-04 | 8.04E-04 | NP | 5.95E+05 | 6.30E+05 | 6.31E+05 | 6.35E+05 | 6.61E+05 |
| PC(18:1_20:5) | 8.51E+05 | 5.62E+05 | 1.51E+00 | 4.21E-01 | 6.34E-01 | 1.10E+06 | 2.20E+05 | 5.00E+00 | 2.92E-05 | 2.27E-04 | 1.73E+00 | 5.42E+05 | 5.83E+05 | 5.90E+05 | 5.60E+05 | 5.69E+05 |
| PC(16:0_22:6) | 1.01E+06 | 1.79E+06 | 5.67E-01 | 1.39E-01 | 3.36E-01 | 1.35E+06 | 1.18E+06 | 1.14E+00 | 2.80E-01 | 3.26E-01 | 1.48E+03 | 1.15E+06 | 1.08E+06 | 1.25E+06 | 1.19E+06 | 1.26E+06 |
| PC(16:1_22:6) | 1.89E+05 | 2.03E+05 | 9.30E-01 | 8.51E-01 | 9.32E-01 | 2.54E+05 | 9.46E+04 | 2.69E+00 | 4.82E-04 | 1.59E-03 | NP | 1.75E+05 | 1.74E+05 | 1.74E+05 | 1.88E+05 | 1.71E+05 |
| PC(18:1_21:2) | 2.83E+04 | 3.43E+04 | 8.26E-01 | 5.19E-01 | 7.26E-01 | 2.81E+04 | 7.73E+04 | 3.63E-01 | 1.42E-02 | 2.49E-02 | 0.00E+00 | 3.52E+04 | 3.01E+04 | 4.00E+04 | 3.99E+04 | 3.98E+04 |
| PC(17:1_22:6) | 2.77E+04 | 4.52E+04 | 6.13E-01 | 3.07E-01 | 5.34E-01 | 3.85E+04 | 2.70E+04 | 1.43E+00 | 2.97E-01 | 3.43E-01 | 5.08E-02 | 3.00E+04 | 4.06E+04 | 2.81E+04 | 3.11E+04 | 2.92E+04 |
| PC(16:1_24:0) | 2.09E+05 | 1.54E+05 | 1.36E+00 | 3.62E-01 | 5.82E-01 | 2.07E+05 | 4.48E+05 | 4.63E-01 | 7.69E-05 | 4.45E-04 | NP | 2.38E+05 | 2.27E+05 | 2.43E+05 | 2.28E+05 | 1.15E+05 |
| PC(18:1_22:1) | 1.48E+05 | 1.86E+05 | 7.96E-01 | 3.47E-01 | 5.69E-01 | 1.60E+05 | 3.68E+05 | 4.35E-01 | 5.51E-04 | 1.71E-03 | NP | 1.83E+05 | 1.74E+05 | 1.97E+05 | 1.98E+05 | 1.96E+05 |
| PC(18:1_22:2) | 7.20E+04 | 7.84E+04 | 9.18E-01 | 7.45E-01 | 8.72E-01 | 7.87E+04 | 1.64E+05 | 4.81E-01 | 1.18E-03 | 3.03E-03 | NP | 9.28E+04 | 9.38E+04 | 9.15E+04 | 1.00E+05 | 9.34E+04 |
| PC(20:1_20:3) | 1.04E+05 | 1.57E+05 | 6.68E-01 | 2.94E-01 | 5.15E-01 | 1.32E+05 | 1.74E+05 | 7.59E-01 | 6.02E-02 | 8.36E-02 | NP | 1.34E+05 | 1.30E+05 | 5.85E+04 | 1.33E+05 | 1.48E+05 |
| PC(18:0_22:4) | 6.09E+04 | 1.77E+05 | 3.43E-01 | 4.14E-02 | 2.13E-01 | 6.98E+04 | 1.38E+05 | 5.05E-01 | 3.22E-06 | 8.62E-05 | NP | 1.09E+05 | 1.04E+05 | 9.79E+04 | 1.11E+05 | 1.10E+05 |
| PC(18:1_22:4) | 2.91E+05 | 5.37E+05 | 5.41E-01 | 9.61E-02 | 2.88E-01 | 3.33E+05 | 5.01E+05 | 6.65E-01 | 2.44E-01 | 2.87E-01 | NP | 3.07E+05 | 2.56E+05 | 2.76E+05 | 3.09E+05 | 2.88E+05 |
| PC(18:0_22:5) | 2.10E+05 | 3.66E+05 | 5.73E-01 | 1.06E-01 | 3.02E-01 | 2.58E+05 | 5.49E+05 | 4.70E-01 | 1.27E-03 | 3.22E-03 | NP | 2.71E+05 | 2.86E+05 | 2.87E+05 | 2.72E+05 | 2.81E+05 |
| PC(18:1_22:5) | 4.50E+05 | 3.69E+05 | 1.22E+00 | 5.46E-01 | 7.51E-01 | 5.41E+05 | 3.75E+05 | 1.44E+00 | 1.12E-02 | 2.06E-02 | NP | 4.05E+05 | 4.01E+05 | 3.90E+05 | 4.06E+05 | 4.03E+05 |
| PC(18:0_22:6) | 2.48E+05 | 6.42E+05 | 3.86E-01 | 2.02E-02 | 1.93E-01 | 2.60E+05 | 6.59E+05 | 3.95E-01 | 4.11E-04 | 1.43E-03 | NP | 3.80E+05 | 4.03E+05 | 4.16E+05 | 4.49E+05 | 3.94E+05 |
| PC(18:1_22:6) | 5.56E+05 | 8.27E+05 | 6.73E-01 | 2.66E-01 | 4.82E-01 | 7.78E+05 | 5.48E+05 | 1.42E+00 | 8.88E-03 | 1.70E-02 | NP | 6.02E+05 | 6.35E+05 | 6.05E+05 | 6.17E+05 | 6.55E+05 |
| PC(18:2_22:6) | 9.35E+04 | 7.12E+04 | 1.31E+00 | 4.53E-01 | 6.63E-01 | 1.32E+05 | 4.86E+04 | 2.71E+00 | 2.14E-04 | 8.51E-04 | NP | 7.83E+04 | 7.72E+04 | 7.94E+04 | 8.53E+04 | 8.26E+04 |
| PC(20:4_20:5) | 2.88E+04 | 1.21E+04 | 2.38E+00 | 6.58E-02 | 2.40E-01 | 3.57E+04 | 7.73E+03 | 4.61E+00 | 2.97E-05 | 2.27E-04 | NP | 1.87E+04 | 2.04E+04 | 1.73E+04 | 2.05E+04 | 1.82E+04 |
| PC(16:0_26:0) | 6.28E+04 | 1.27E+04 | 4.94E+00 | 1.84E-02 | 1.93E-01 | 5.79E+04 | 1.08E+05 | 5.38E-01 | 7.72E-02 | 1.03E-01 | 0.00E+00 | 7.27E+04 | 7.38E+04 | 7.58E+04 | 9.05E+04 | 7.78E+04 |
| PC(16:0_26:1) | 2.96E+05 | 1.17E+05 | 2.53E+00 | 1.06E-01 | 3.02E-01 | 3.06E+05 | 7.05E+05 | 4.34E-01 | 1.00E-03 | 2.64E-03 | NP | 3.11E+05 | 2.03E+05 | 3.55E+05 | 3.42E+05 | 3.55E+05 |
| PC(20:4_22:6) | 6.61E+04 | 2.98E+04 | 2.22E+00 | 6.11E-02 | 2.38E-01 | 9.29E+04 | 3.32E+04 | 2.79E+00 | 2.12E-04 | 8.51E-04 | 5.51E+02 | 4.93E+04 | 4.87E+04 | 5.34E+04 | 5.13E+04 | 5.41E+04 |

| | | | | | | | | | | | | | | | | |
|---|---|---|---|---|---|---|---|---|---|---|---|---|---|---|---|---|
| PC(20:5_22:6) | 3.12E+04 | 1.08E+04 | 2.89E+00 | 7.64E-02 | 2.63E-01 | 5.06E+04 | 6.58E+03 | 7.69E+00 | 1.55E-04 | 7.05E-04 | 0.00E+00 | 2.23E+04 | 2.05E+04 | 2.47E+04 | 2.41E+04 | 2.24E+04 |
| PC(18:1_24:1) | 2.40E+05 | 1.25E+05 | 1.93E+00 | 1.58E-01 | 3.53E-01 | 2.56E+05 | 3.61E+05 | 7.09E-01 | 1.01E-01 | 1.30E-01 | NP | 2.73E+05 | 2.66E+05 | 1.90E+05 | 2.86E+05 | 2.49E+05 |
| PC(18:0_24:5) | 5.88E+04 | 1.28E+05 | 4.60E-01 | 5.85E-02 | 2.38E-01 | 6.51E+04 | 5.69E+04 | 1.14E+00 | 3.16E-01 | 3.60E-01 | NP | 7.60E+04 | 7.66E+04 | 6.92E+04 | 7.38E+04 | 6.15E+04 |
| PC(18:1_24:5) | 1.08E+05 | 1.98E+05 | 5.47E-01 | 1.26E-01 | 3.26E-01 | 1.32E+05 | 5.25E+04 | 2.51E+00 | 5.86E-04 | 1.78E-03 | NP | 7.14E+04 | 1.11E+05 | 7.20E+04 | 8.23E+04 | 1.10E+05 |
| PC(18:1_24:6) | 2.08E+04 | 5.46E+04 | 3.82E-01 | 3.08E-02 | 1.93E-01 | 2.96E+04 | 3.60E+04 | 8.24E-01 | 2.08E-02 | 3.43E-02 | NP | 3.30E+04 | 3.15E+04 | 3.38E+04 | 3.41E+04 | 3.24E+04 |
| PC(20:1_22:6) | 3.42E+04 | 1.04E+05 | 3.29E-01 | 9.77E-03 | 1.87E-01 | 3.35E+04 | 7.55E+04 | 4.43E-01 | 2.94E-05 | 2.27E-04 | NP | 5.48E+04 | 5.42E+04 | 5.66E+04 | 5.69E+04 | 5.10E+04 |
| PC(20:4_22:5) | 7.57E+04 | 3.81E+04 | 1.99E+00 | 1.57E-01 | 3.53E-01 | 9.10E+04 | 4.42E+04 | 2.06E+00 | 1.25E-03 | 3.18E-03 | NP | 5.82E+04 | 5.49E+04 | 6.06E+04 | 6.01E+04 | 6.02E+04 |
| PC(22:5_22:6) | 1.99E+04 | 1.65E+04 | 1.21E+00 | 5.88E-01 | 7.81E-01 | 2.21E+04 | 1.92E+04 | 1.15E+00 | 3.70E-01 | 4.16E-01 | 0.00E+00 | 1.81E+04 | 1.57E+04 | 1.81E+04 | 2.07E+04 | 1.69E+04 |
| PC(22:6_22:6) | 1.27E+04 | 1.22E+04 | 1.04E+00 | 8.94E-01 | 9.49E-01 | 1.57E+04 | 1.18E+04 | 1.33E+00 | 1.24E-01 | 1.57E-01 | NP | 1.28E+04 | 1.05E+04 | 1.25E+04 | 1.12E+04 | 1.23E+04 |
| PC(18:1_26:1) | 2.11E+05 | 4.10E+04 | 5.15E+00 | 2.10E-02 | 1.93E-01 | 2.83E+05 | 3.60E+05 | 7.84E-01 | 3.16E-02 | 4.85E-02 | NP | 1.01E+05 | 2.38E+05 | 2.13E+05 | 2.49E+05 | 2.43E+05 |
| PC(26:1_18:2) | 4.12E+04 | 9.99E+03 | 4.13E+00 | 7.41E-02 | 2.58E-01 | 5.09E+04 | 2.68E+04 | 1.90E+00 | 2.44E-04 | 9.06E-04 | NP | 2.01E+04 | 1.53E+04 | 1.95E+04 | 2.72E+04 | 1.94E+04 |
| PC(18:1_28:1) | 4.33E+04 | 1.08E+04 | 4.02E+00 | 5.74E-02 | 2.38E-01 | 5.20E+04 | 8.34E+04 | 6.23E-01 | 6.67E-02 | 9.03E-02 | 9.75E+02 | 3.65E+04 | 4.23E+04 | 4.87E+04 | 3.76E+04 | 4.51E+04 |
| PC(26:0_20:4) | 8.18E+04 | 4.42E+03 | 1.85E+01 | 2.22E-01 | 4.24E-01 | 6.90E+04 | 6.97E+04 | 9.90E-01 | 9.67E-01 | 9.67E-01 | NP | 5.19E+04 | 5.60E+04 | 4.75E+04 | 4.96E+04 | 5.38E+04 |
| PC(O-12:0_16:0) | 3.53E+04 | 4.16E+04 | 8.49E-01 | 6.23E-01 | 8.01E-01 | 4.29E+04 | 3.97E+04 | 1.08E+00 | 6.30E-01 | 6.61E-01 | NP | 3.80E+04 | 3.62E+04 | 3.71E+04 | 3.74E+04 | 3.71E+04 |
| PC(O-17:0_16:0) | 5.02E+00 | 0.00E+00 | #¡DIV/0! | 1.43E-01 | 3.42E-01 | 1.62E+02 | 1.84E+04 | 8.83E-03 | 2.11E-03 | 4.92E-03 | NP | 5.28E+03 | 4.31E+03 | 4.98E+03 | 6.72E+03 | 4.78E+03 |
| PC(O-18:0_16:0) | 3.29E+04 | 5.66E+04 | 5.81E-01 | 2.83E-01 | 5.03E-01 | 3.48E+04 | 1.50E+06 | 2.32E-02 | 1.92E-06 | 7.32E-05 | NP | 2.44E+05 | 2.76E+05 | 3.77E+05 | 4.06E+05 | 3.88E+05 |
| PC(O-16:0_18:1) | 1.05E+06 | 1.05E+06 | 9.95E-01 | 9.79E-01 | 9.84E-01 | 1.09E+06 | 7.25E+06 | 1.50E-01 | 1.04E-06 | 5.59E-05 | NP | 2.65E+06 | 2.71E+06 | 2.84E+06 | 2.64E+06 | 2.93E+06 |
| PC(O-16:1_18:2) | 5.90E+04 | 8.09E+04 | 7.29E-01 | 1.84E-01 | 3.86E-01 | 6.99E+04 | 1.37E+05 | 5.10E-01 | 1.92E-03 | 4.58E-03 | 2.13E+03 | 7.35E+04 | 7.14E+04 | 6.60E+04 | 8.46E+04 | 6.33E+04 |
| PC(O-17:0_18:1) | 3.17E+02 | 6.85E-01 | 4.63E+02 | 3.57E-01 | 5.80E-01 | 2.92E-02 | 1.88E+05 | 1.55E-07 | 3.83E-05 | 2.57E-04 | NP | 2.74E+04 | 2.14E+04 | 2.10E+04 | 1.93E+04 | 1.84E+04 |
| PC(O-17:1_18:1) | 9.48E+03 | 1.39E+04 | 6.82E-01 | 9.26E-02 | 2.83E-01 | 9.84E+03 | 5.52E+04 | 1.78E-01 | 1.44E-05 | 1.78E-04 | NP | 1.67E+04 | 2.15E+04 | 2.89E+04 | 1.96E+04 | 2.45E+04 |
| PC(O-20:0_16:0) | 1.15E+04 | 3.83E+04 | 3.01E-01 | 3.79E-03 | 1.16E-01 | 1.28E+04 | 3.84E+05 | 3.35E-02 | 1.16E-05 | 1.55E-04 | NP | 1.07E+05 | 1.04E+05 | 1.01E+05 | 1.01E+05 | 9.24E+04 |
| PC(O-18:0_18:1) | 7.11E+04 | 1.53E+05 | 4.64E-01 | 3.60E-02 | 2.06E-01 | 9.38E+04 | 2.30E+06 | 4.08E-02 | 7.93E-07 | 5.59E-05 | NP | 4.89E+05 | 6.94E+05 | 5.24E+05 | 4.74E+05 | 5.28E+05 |
| PC(O-16:0_20:3) | 3.16E+04 | 1.99E+04 | 1.59E+00 | 2.45E-01 | 4.60E-01 | 3.07E+04 | 3.45E+05 | 8.89E-02 | 1.18E-05 | 1.55E-04 | 1.22E+01 | 1.17E+05 | 1.25E+05 | 1.30E+05 | 1.05E+05 | 1.08E+05 |
| PC(O-18:2_18:1) | 9.63E+04 | 1.98E+05 | 4.87E-01 | 1.96E-02 | 1.93E-01 | 1.16E+05 | 3.33E+05 | 3.47E-01 | 7.86E-04 | 2.12E-03 | 4.14E-03 | 1.88E+05 | 1.90E+05 | 1.91E+05 | 1.65E+05 | 1.91E+05 |
| PC(O-16:0_20:4) | 2.29E+05 | 1.38E+05 | 1.66E+00 | 1.87E-01 | 3.88E-01 | 2.62E+05 | 6.97E+05 | 3.76E-01 | 1.01E-04 | 5.09E-04 | NP | 3.15E+05 | 3.26E+05 | 3.41E+05 | 3.45E+05 | 3.21E+05 |
| PC(O-16:0_20:5) | 7.06E+04 | 2.74E+04 | 2.57E+00 | 2.07E-02 | 1.93E-01 | 8.09E+04 | 1.11E+05 | 7.29E-01 | 4.35E-02 | 6.29E-02 | NP | 6.87E+04 | 7.09E+04 | 7.20E+04 | 7.29E+04 | 7.25E+04 |
| PC(O-16:1_20:4) | 2.74E+05 | 3.99E+05 | 6.86E-01 | 2.66E-01 | 4.82E-01 | 3.60E+05 | 2.00E+05 | 1.80E+00 | 2.71E-02 | 4.26E-02 | NP | 2.91E+05 | 2.78E+05 | 2.85E+05 | 3.07E+05 | 2.78E+05 |
| PC(O-18:0_20:3) | 3.44E+04 | 3.67E+04 | 9.39E-01 | 8.10E-01 | 9.10E-01 | 3.92E+04 | 2.40E+05 | 1.63E-01 | 1.12E-05 | 1.55E-04 | NP | 1.14E+05 | 8.89E+04 | 1.09E+05 | 9.00E+04 | 4.82E+04 |
| PC(O-18:0_20:4) | 8.39E+04 | 9.33E+04 | 8.99E-01 | 7.17E-01 | 8.58E-01 | 9.95E+04 | 3.76E+05 | 2.65E-01 | 1.26E-05 | 1.61E-04 | NP | 1.48E+05 | 1.51E+05 | 1.61E+05 | 1.57E+05 | 1.36E+05 |
| PC(O-18:1_20:4) | 8.86E+04 | 1.23E+05 | 7.18E-01 | 3.68E-01 | 5.86E-01 | 1.13E+05 | 1.49E+05 | 7.56E-01 | 4.50E-02 | 6.48E-02 | NP | 1.13E+05 | 1.13E+05 | 1.04E+05 | 1.13E+05 | 1.25E+05 |
| PC(O-20:6_18:0) | 4.48E+04 | 2.05E+04 | 2.18E+00 | 6.28E-02 | 2.39E-01 | 5.66E+04 | 7.93E+03 | 7.14E+00 | 3.76E-03 | 8.26E-03 | NP | 2.64E+04 | 2.46E+04 | 2.51E+04 | 3.57E+04 | 3.32E+04 |
| PC(O-16:1_22:5) | 1.99E+05 | 3.72E+05 | 5.34E-01 | 5.98E-02 | 2.38E-01 | 2.59E+05 | 4.86E+05 | 5.33E-01 | 3.71E-03 | 8.24E-03 | NP | 3.25E+05 | 2.96E+05 | 3.17E+05 | 3.18E+05 | 3.32E+05 |
| PC(O-18:1_20:5) | 4.88E+04 | 5.28E+04 | 9.23E-01 | 8.39E-01 | 9.32E-01 | 4.92E+04 | 5.35E+04 | 9.20E-01 | 5.10E-01 | 5.57E-01 | NP | 5.40E+04 | 4.47E+04 | 4.16E+04 | 4.47E+04 | 3.88E+04 |

| | | | | | | | | | | | | | | | | |
|---|---|---|---|---|---|---|---|---|---|---|---|---|---|---|---|---|
| PC(O-18:2_20:5) | 4.14E+04 | 4.29E+04 | 9.66E-01 | 9.31E-01 | 9.59E-01 | 5.50E+04 | 4.98E+03 | 1.10E+01 | 8.79E-05 | 4.94E-04 | NP | 3.54E+04 | 3.31E+04 | 3.26E+04 | 3.35E+04 | 3.38E+04 |
| PC(O-16:1_22:6) | 6.04E+04 | 1.91E+05 | 3.15E-01 | 1.17E-02 | 1.91E-01 | 7.58E+04 | 1.17E+05 | 6.47E-01 | 2.14E-02 | 3.49E-02 | NP | 9.89E+04 | 1.06E+05 | 9.89E+04 | 1.12E+05 | 1.07E+05 |
| PC(O-18:0_22:5) | 2.21E+04 | 4.07E+04 | 5.43E-01 | 1.13E-01 | 3.16E-01 | 3.28E+04 | 1.58E+05 | 2.08E-01 | 1.71E-05 | 1.86E-04 | NP | 6.29E+04 | 6.53E+04 | 7.19E+04 | 6.48E+04 | 6.71E+04 |
| PC(O-18:0_22:6) | 3.90E+04 | 6.83E+04 | 5.71E-01 | 4.07E-02 | 2.13E-01 | 5.43E+04 | 2.61E+05 | 2.08E-01 | 3.91E-05 | 2.57E-04 | NP | 9.90E+04 | 9.22E+04 | 1.03E+05 | 6.80E+04 | 1.03E+05 |
| PC(O-18:1_22:6) | 3.81E+04 | 9.61E+04 | 3.96E-01 | 2.72E-02 | 1.93E-01 | 5.59E+04 | 8.99E+04 | 6.22E-01 | 3.77E-02 | 5.54E-02 | NP | 6.91E+04 | 6.21E+04 | 7.15E+04 | 5.59E+04 | 6.71E+04 |
| PC(O-18:2_22:6) | 1.51E+04 | 6.22E+04 | 2.43E-01 | 5.48E-03 | 1.40E-01 | 2.29E+04 | 2.20E+04 | 1.04E+00 | 8.64E-01 | 8.83E-01 | NP | 2.25E+04 | 3.16E+04 | 2.68E+04 | 2.69E+04 | 3.22E+04 |
| PC(O-24:1_18:1) | 3.44E+03 | 2.23E+04 | 1.54E-01 | 3.94E-03 | 1.16E-01 | 3.91E+03 | 1.24E+05 | 3.16E-02 | 9.71E-03 | 1.84E-02 | NP | 5.86E+04 | 3.47E+04 | 4.99E+04 | 5.96E+04 | 8.44E+04 |
| PE(14:0_16:0) | 8.90E+04 | 5.65E+04 | 1.57E+00 | 2.81E-02 | 1.93E-01 | 7.39E+04 | 3.11E+04 | 2.37E+00 | 1.80E-03 | 4.33E-03 | NP | 5.42E+04 | 6.01E+04 | 6.06E+04 | 5.78E+04 | 5.85E+04 |
| PE(14:0_16:1) | 3.25E+04 | 3.88E+04 | 8.38E-01 | 6.12E-01 | 7.94E-01 | 3.52E+04 | 1.93E+04 | 1.82E+00 | 1.70E-03 | 4.15E-03 | NP | 2.99E+04 | 2.54E+04 | 2.56E+04 | 3.00E+04 | 3.09E+04 |
| PE(16:0_16:1) | 1.01E+06 | 9.92E+05 | 1.02E+00 | 9.38E-01 | 9.60E-01 | 1.08E+06 | 5.14E+05 | 2.10E+00 | 3.62E-03 | 8.08E-03 | NP | 8.32E+05 | 8.28E+05 | 8.12E+05 | 8.55E+05 | 8.33E+05 |
| PE(16:1_16:1) | 1.23E+05 | 6.79E+04 | 1.81E+00 | 1.28E-01 | 3.28E-01 | 1.38E+05 | 4.29E+04 | 3.22E+00 | 6.67E-04 | 1.91E-03 | NP | 8.77E+04 | 9.62E+04 | 7.16E+04 | 9.95E+04 | 9.38E+04 |
| PE(16:0_17:1) | 2.81E+05 | 4.27E+05 | 6.58E-01 | 8.85E-02 | 2.78E-01 | 2.92E+05 | 2.10E+05 | 1.39E+00 | 7.06E-03 | 1.40E-02 | NP | 2.76E+05 | 2.79E+05 | 2.77E+05 | 3.08E+05 | 2.78E+05 |
| PE(15:0_18:1)+D7:(s) | 2.78E+05 | 2.88E+05 | 9.66E-01 | 8.74E-01 | 9.38E-01 | 2.03E+05 | 2.21E+05 | 9.21E-01 | 2.98E-01 | 3.43E-01 | 1.83E+05 | 2.17E+05 | 2.24E+05 | 2.20E+05 | 2.30E+05 | 2.10E+05 |
| PE(16:1_17:1) | 4.99E+04 | 5.13E+04 | 9.74E-01 | 9.37E-01 | 9.60E-01 | 5.77E+04 | 2.04E+04 | 2.82E+00 | 1.01E-04 | 5.09E-04 | NP | 3.57E+04 | 3.56E+04 | 3.43E+04 | 3.85E+04 | 4.53E+04 |
| PE(16:0_18:0) | 1.84E+05 | 2.39E+05 | 7.69E-01 | 2.92E-01 | 5.15E-01 | 1.70E+05 | 1.21E+05 | 1.41E+00 | 3.45E-02 | 5.16E-02 | NP | 1.66E+05 | 1.55E+05 | 1.56E+05 | 1.60E+05 | 1.57E+05 |
| PE(16:0_18:1) | 7.27E+06 | 7.59E+06 | 9.57E-01 | 8.46E-01 | 9.32E-01 | 8.25E+06 | 4.81E+06 | 1.72E+00 | 6.16E-03 | 1.25E-02 | 8.92E+03 | 6.17E+06 | 6.60E+06 | 6.27E+06 | 6.77E+06 | 6.47E+06 |
| PE(16:1_18:1) | 1.85E+06 | 1.26E+06 | 1.47E+00 | 1.74E-01 | 3.77E-01 | 2.38E+06 | 8.38E+05 | 2.84E+00 | 4.15E-04 | 1.43E-03 | 1.57E+03 | 1.47E+06 | 1.53E+06 | 1.66E+06 | 1.53E+06 | 1.53E+06 |
| PE(16:1_18:2) | 7.82E+04 | 6.73E+04 | 1.16E+00 | 6.66E-01 | 8.25E-01 | 1.05E+05 | 3.50E+04 | 3.00E+00 | 1.87E-04 | 8.04E-04 | NP | 6.19E+04 | 6.31E+04 | 6.33E+04 | 6.41E+04 | 5.94E+04 |
| PE(14:0_20:4) | 2.29E+04 | 3.38E+04 | 6.78E-01 | 3.69E-01 | 5.86E-01 | 2.34E+04 | 1.83E+04 | 1.28E+00 | 1.58E-02 | 2.71E-02 | NP | 2.00E+04 | 2.04E+04 | 1.95E+04 | 2.33E+04 | 2.23E+04 |
| PE(17:0_18:1) | 7.00E+05 | 1.05E+06 | 6.70E-01 | 1.83E-01 | 3.85E-01 | 7.63E+05 | 7.00E+05 | 1.09E+00 | 1.30E-01 | 1.63E-01 | 5.18E+02 | 7.41E+05 | 7.23E+05 | 6.39E+05 | 7.75E+05 | 8.05E+05 |
| PE(17:1_18:1) | 5.74E+05 | 5.94E+05 | 9.66E-01 | 8.97E-01 | 9.49E-01 | 6.38E+05 | 2.94E+05 | 2.17E+00 | 1.56E-05 | 1.84E-04 | NP | 4.44E+05 | 4.60E+05 | 4.76E+05 | 4.49E+05 | 4.64E+05 |
| PE(18:0_18:1) | 9.06E+06 | 9.77E+06 | 9.27E-01 | 7.59E-01 | 8.77E-01 | 1.02E+07 | 9.42E+06 | 1.08E+00 | 4.06E-01 | 4.51E-01 | 1.60E+04 | 8.91E+06 | 8.73E+06 | 8.73E+06 | 8.92E+06 | 8.70E+06 |
| PE(18:1_18:1) | 9.97E+06 | 8.85E+06 | 1.13E+00 | 6.29E-01 | 8.03E-01 | 1.13E+07 | 5.53E+06 | 2.05E+00 | 6.18E-04 | 1.83E-03 | 1.39E+04 | 8.34E+06 | 8.04E+06 | 8.16E+06 | 8.64E+06 | 8.37E+06 |
| PE(18:0_18:2) | 1.71E+06 | 1.58E+06 | 1.08E+00 | 7.94E-01 | 8.95E-01 | 1.87E+06 | 1.26E+06 | 1.49E+00 | 2.60E-02 | 4.13E-02 | 1.95E+03 | 1.87E+06 | 2.13E+06 | 1.64E+06 | 1.53E+06 | 1.48E+06 |
| PE(18:1_18:2) | 1.08E+06 | 1.12E+06 | 9.62E-01 | 9.01E-01 | 9.51E-01 | 1.22E+06 | 4.98E+05 | 2.46E+00 | 2.19E-05 | 2.09E-04 | 6.51E+02 | 8.77E+05 | 8.85E+05 | 7.52E+05 | 8.55E+05 | 9.17E+05 |
| PE(16:0_20:3)_Rt10.5 | 3.68E+05 | 4.04E+05 | 9.11E-01 | 7.35E-01 | 8.66E-01 | 3.66E+05 | 3.95E+05 | 9.26E-01 | 4.16E-01 | 4.60E-01 | 8.32E-01 | 3.52E+05 | 3.39E+05 | 3.06E+05 | 3.82E+05 | 3.74E+05 |
| PE(16:0_20:3)_Rt11.0 | 9.59E+04 | 7.84E+04 | 1.22E+00 | 5.14E-01 | 7.24E-01 | 8.70E+04 | 1.58E+05 | 5.52E-01 | 1.12E-04 | 5.54E-04 | 5.22E-03 | 8.52E+04 | 1.01E+05 | 1.05E+05 | 1.02E+05 | 1.01E+05 |
| PE(16:0_20:4) | 1.73E+06 | 1.47E+06 | 1.18E+00 | 6.53E-01 | 8.20E-01 | 1.95E+06 | 1.06E+06 | 1.83E+00 | 3.38E-06 | 8.62E-05 | 2.15E+03 | 1.28E+06 | 1.49E+06 | 1.23E+06 | 1.45E+06 | 1.39E+06 |
| PE(16:1_20:4) | 2.27E+05 | 1.14E+05 | 1.99E+00 | 1.34E-01 | 3.34E-01 | 2.92E+05 | 5.55E+04 | 5.27E+00 | 7.38E-06 | 1.41E-04 | NP | 1.50E+05 | 1.48E+05 | 1.55E+05 | 1.56E+05 | 1.56E+05 |
| PE(16:0_20:5) | 6.26E+05 | 3.87E+05 | 1.62E+00 | 2.60E-01 | 4.76E-01 | 7.50E+05 | 1.96E+05 | 3.83E+00 | 2.56E-07 | 3.26E-05 | NP | 4.47E+05 | 4.60E+05 | 4.22E+05 | 4.26E+05 | 4.05E+05 |
| PE(16:1_20:5) | 9.33E+04 | 4.20E+04 | 2.22E+00 | 1.30E-01 | 3.29E-01 | 1.17E+05 | 1.30E+04 | 8.96E+00 | 1.76E-04 | 7.73E-04 | NP | 5.77E+04 | 5.76E+04 | 5.91E+04 | 6.09E+04 | 6.14E+04 |

| | | | | | | | | | | | | | | | | |
|---|---|---|---|---|---|---|---|---|---|---|---|---|---|---|---|---|
| PE(19:0_18:1)_Rt13.1 | 4.34E+04 | 5.97E+04 | 7.27E-01 | 3.12E-01 | 5.40E-01 | 4.31E+04 | 3.97E+04 | 1.09E+00 | 1.93E-01 | 2.31E-01 | NP | 3.95E+04 | 3.97E+04 | 4.37E+04 | 4.67E+04 | 4.01E+04 |
| PE(19:0_18:1)_Rt13.5 | 1.24E+05 | 1.47E+05 | 8.41E-01 | 5.75E-01 | 7.68E-01 | 1.27E+05 | 1.30E+05 | 9.76E-01 | 8.05E-01 | 8.31E-01 | NP | 1.03E+05 | 1.15E+05 | 1.18E+05 | 1.14E+05 | 1.11E+05 |
| PE(18:1_19:1) | 3.01E+05 | 3.22E+05 | 9.35E-01 | 8.15E-01 | 9.13E-01 | 3.43E+05 | 2.52E+05 | 1.36E+00 | 4.57E-03 | 9.50E-03 | NP | 2.72E+05 | 2.71E+05 | 2.80E+05 | 2.99E+05 | 2.74E+05 |
| PE(17:0_20:3) | 4.45E+04 | 5.69E+04 | 7.82E-01 | 4.16E-01 | 6.30E-01 | 4.56E+04 | 5.53E+04 | 8.25E-01 | 6.77E-03 | 1.35E-02 | NP | 4.10E+04 | 4.72E+04 | 4.37E+04 | 4.84E+04 | 4.51E+04 |
| PE(17:0_20:4) | 2.45E+05 | 2.95E+05 | 8.32E-01 | 5.72E-01 | 7.66E-01 | 2.55E+05 | 1.81E+05 | 1.41E+00 | 1.37E-02 | 2.41E-02 | NP | 2.15E+05 | 2.22E+05 | 2.31E+05 | 2.33E+05 | 2.07E+05 |
| PE(17:0_20:5) | 5.15E+04 | 4.21E+04 | 1.22E+00 | 6.15E-01 | 7.94E-01 | 6.20E+04 | 1.28E+04 | 4.83E+00 | 5.33E-04 | 1.70E-03 | 2.84E-02 | 3.61E+04 | 4.45E+04 | 4.10E+04 | 3.40E+04 | 4.10E+04 |
| PE(20:0_18:1) | 3.49E+05 | 3.64E+05 | 9.61E-01 | 8.69E-01 | 9.35E-01 | 3.28E+05 | 3.48E+05 | 9.43E-01 | 6.41E-01 | 6.70E-01 | NP | 2.91E+05 | 3.24E+05 | 3.15E+05 | 3.19E+05 | 3.02E+05 |
| PE(18:1_20:1) | 9.11E+05 | 9.71E+05 | 9.39E-01 | 7.76E-01 | 8.85E-01 | 9.88E+05 | 5.80E+05 | 1.70E+00 | 5.33E-03 | 1.10E-02 | 6.07E+02 | 7.68E+05 | 7.70E+05 | 8.17E+05 | 8.07E+05 | 7.14E+05 |
| PE(18:0_20:2) | 9.68E+04 | 1.10E+05 | 8.76E-01 | 7.25E-01 | 8.59E-01 | 9.51E+04 | 5.03E+05 | 1.89E-01 | 5.11E-06 | 1.22E-04 | 4.82E-03 | 1.65E+05 | 1.84E+05 | 1.79E+05 | 1.78E+05 | 2.11E+05 |
| PE(18:0_20:3)_Rt12.0 | 1.38E+06 | 1.34E+06 | 1.03E+00 | 9.28E-01 | 9.59E-01 | 1.58E+06 | 1.26E+06 | 1.26E+00 | 3.47E-02 | 5.18E-02 | 1.33E+03 | 1.21E+06 | 1.24E+06 | 1.25E+06 | 1.28E+06 | 1.25E+06 |
| PE(18:0_20:3)_Rt12.5 | 2.90E+05 | 3.22E+05 | 8.98E-01 | 6.98E-01 | 8.47E-01 | 3.17E+05 | 6.77E+05 | 4.69E-01 | 1.47E-04 | 6.79E-04 | 3.69E-02 | 3.64E+05 | 3.85E+05 | 3.89E+05 | 3.55E+05 | 3.39E+05 |
| PE(18:0_20:4) | 9.65E+06 | 8.40E+06 | 1.15E+00 | 6.93E-01 | 8.44E-01 | 1.03E+07 | 6.14E+06 | 1.68E+00 | 3.22E-05 | 2.32E-04 | 1.35E+04 | 7.93E+06 | 7.00E+06 | 8.37E+06 | 8.12E+06 | 8.40E+06 |
| PE(18:1_20:4) | 3.40E+06 | 2.77E+06 | 1.23E+00 | 5.53E-01 | 7.57E-01 | 3.88E+06 | 1.40E+06 | 2.76E+00 | 1.17E-06 | 5.59E-05 | 2.35E+03 | 2.53E+06 | 2.53E+06 | 2.49E+06 | 2.67E+06 | 2.54E+06 |
| PE(18:0_20:5) | 2.53E+06 | 1.60E+06 | 1.58E+00 | 3.61E-01 | 5.82E-01 | 2.64E+06 | 9.66E+05 | 2.73E+00 | 4.60E-05 | 2.88E-04 | 2.69E+03 | 1.68E+06 | 1.61E+06 | 1.67E+06 | 1.77E+06 | 1.78E+06 |
| PE(18:1_20:5) | 1.11E+06 | 5.06E+05 | 2.20E+00 | 1.34E-01 | 3.34E-01 | 1.28E+06 | 1.39E+05 | 9.23E+00 | 1.64E-05 | 1.84E-04 | 4.11E+00 | 6.25E+05 | 7.26E+05 | 7.30E+05 | 6.21E+05 | 6.99E+05 |
| PE(16:1_22:6) | 1.03E+05 | 1.40E+05 | 7.38E-01 | 4.48E-01 | 6.58E-01 | 1.45E+05 | 4.70E+04 | 3.09E+00 | 1.10E-05 | 1.55E-04 | NP | 9.43E+04 | 8.95E+04 | 9.63E+04 | 8.48E+04 | 9.01E+04 |
| PE(18:1_21:1) | 4.20E+04 | 5.36E+04 | 7.84E-01 | 3.98E-01 | 6.20E-01 | 4.34E+04 | 3.15E+04 | 1.38E+00 | 1.84E-02 | 3.09E-02 | NP | 3.38E+04 | 3.97E+04 | 3.76E+04 | 3.93E+04 | 3.57E+04 |
| PE(18:0_21:3) | 7.57E+04 | 7.76E+04 | 9.75E-01 | 9.29E-01 | 9.59E-01 | 8.53E+04 | 8.07E+04 | 1.06E+00 | 5.40E-01 | 5.84E-01 | NP | 7.44E+04 | 6.92E+04 | 7.37E+04 | 7.37E+04 | 7.24E+04 |
| PE(19:0_20:4) | 9.16E+04 | 8.54E+04 | 1.07E+00 | 8.56E-01 | 9.32E-01 | 8.94E+04 | 2.05E+04 | 4.37E+00 | 6.10E-04 | 1.83E-03 | NP | 5.72E+04 | 5.02E+04 | 6.14E+04 | 6.63E+04 | 7.46E+04 |
| PE(22:0_18:1) | 4.11E+05 | 3.17E+05 | 1.30E+00 | 4.07E-01 | 6.25E-01 | 3.73E+05 | 2.98E+05 | 1.25E+00 | 3.12E-01 | 3.57E-01 | 3.44E-03 | 2.99E+05 | 3.00E+05 | 3.22E+05 | 3.07E+05 | 2.87E+05 |
| PE(18:1_22:1) | 2.74E+05 | 2.02E+05 | 1.35E+00 | 5.10E-01 | 7.24E-01 | 2.66E+05 | 1.62E+05 | 1.64E+00 | 3.61E-02 | 5.36E-02 | 1.69E+03 | 2.13E+05 | 2.15E+05 | 2.17E+05 | 2.38E+05 | 2.12E+05 |
| PE(18:0_22:3) | 5.76E+04 | 1.06E+05 | 5.44E-01 | 8.00E-02 | 2.66E-01 | 6.22E+04 | 9.61E+04 | 6.47E-01 | 7.62E-03 | 1.49E-02 | NP | 7.23E+04 | 7.29E+04 | 6.66E+04 | 7.22E+04 | 7.32E+04 |
| PE(18:1_22:4) | 1.70E+05 | 3.95E+05 | 4.31E-01 | 6.01E-02 | 2.38E-01 | 2.44E+05 | 4.20E+04 | 5.81E+00 | 3.87E-05 | 2.57E-04 | 5.99E-02 | 2.14E+05 | 1.74E+05 | 2.14E+05 | 1.91E+05 | 1.70E+05 |
| PE(20:1_20:4) | 1.85E+05 | 2.79E+05 | 6.62E-01 | 2.08E-01 | 4.08E-01 | 2.07E+05 | 9.57E+04 | 2.17E+00 | 7.84E-04 | 2.12E-03 | 5.99E-02 | 1.44E+05 | 1.83E+05 | 1.85E+05 | 1.49E+05 | 1.63E+05 |
| PE(18:0_22:5) | 5.55E+05 | 1.82E+06 | 3.05E-01 | 3.14E-03 | 1.16E-01 | 6.21E+05 | 1.63E+06 | 3.80E-01 | 9.19E-05 | 4.94E-04 | 6.57E+02 | 1.12E+06 | 1.06E+06 | 1.16E+06 | 1.16E+06 | 1.16E+06 |
| PE(18:1_22:5) | 4.09E+05 | 6.16E+05 | 6.65E-01 | 2.48E-01 | 4.64E-01 | 5.17E+05 | 3.63E+05 | 1.43E+00 | 1.13E-02 | 2.07E-02 | NP | 3.99E+05 | 4.46E+05 | 3.02E+05 | 4.76E+05 | 4.42E+05 |
| PE(18:0_22:6) | 8.78E+05 | 3.68E+06 | 2.38E-01 | 3.78E-03 | 1.16E-01 | 8.09E+05 | 3.27E+06 | 2.47E-01 | 5.79E-08 | 1.11E-05 | NP | 1.92E+06 | 1.90E+06 | 2.11E+06 | 2.00E+06 | 2.12E+06 |
| PE(18:1_22:6) | 8.20E+05 | 1.68E+06 | 4.88E-01 | 7.99E-02 | 2.66E-01 | 9.56E+05 | 8.27E+05 | 1.16E+00 | 3.10E-02 | 4.79E-02 | NP | 8.89E+05 | 9.43E+05 | 8.98E+05 | 9.41E+05 | 9.24E+05 |
| PE(18:2_22:6) | 1.97E+04 | 4.16E+04 | 4.73E-01 | 7.11E-02 | 2.49E-01 | 3.01E+04 | 1.06E+04 | 2.84E+00 | 5.55E-04 | 1.71E-03 | NP | 2.37E+04 | 2.06E+04 | 1.96E+04 | 2.90E+04 | 2.14E+04 |
| PE(23:0_18:1) | 4.72E+04 | 2.84E+04 | 1.66E+00 | 1.76E-01 | 3.78E-01 | 3.72E+04 | 3.49E+04 | 1.07E+00 | 6.83E-01 | 7.11E-01 | NP | 3.44E+04 | 3.22E+04 | 2.82E+04 | 3.10E+04 | 2.99E+04 |
| PE(24:0_18:1) | 5.52E+05 | 2.55E+05 | 2.16E+00 | 6.97E-02 | 2.48E-01 | 4.60E+05 | 4.04E+05 | 1.14E+00 | 4.07E-01 | 4.51E-01 | NP | 3.64E+05 | 3.76E+05 | 3.56E+05 | 3.64E+05 | 3.52E+05 |

| | | | | | | | | | | | | | | | | |
|---|---|---|---|---|---|---|---|---|---|---|---|---|---|---|---|---|
| PE(18:1_24:1) | 3.21E+05 | 1.73E+05 | 1.85E+00 | 1.36E-01 | 3.36E-01 | 3.42E+05 | 1.21E+05 | 2.83E+00 | 7.67E-04 | 2.12E-03 | NP | 2.18E+05 | 1.96E+05 | 2.11E+05 | 2.12E+05 | 2.17E+05 |
| PE(24:0_18:2) | 4.41E+04 | 2.33E+04 | 1.89E+00 | 1.56E-01 | 3.53E-01 | 4.00E+04 | 4.86E+04 | 8.25E-01 | 3.28E-01 | 3.71E-01 | NP | 2.99E+04 | 3.27E+04 | 3.32E+04 | 3.32E+04 | 2.90E+04 |
| PE(18:0_24:4) | 1.29E+04 | 3.95E+04 | 3.27E-01 | 1.92E-02 | 1.93E-01 | 1.44E+04 | 3.20E+03 | 4.50E+00 | 2.02E-04 | 8.26E-04 | NP | 1.50E+04 | 1.40E+04 | 1.71E+04 | 1.70E+04 | 1.49E+04 |
| PE(22:0_20:4) | 7.83E+04 | 8.57E+04 | 9.13E-01 | 7.90E-01 | 8.93E-01 | 6.22E+04 | 5.88E+04 | 1.06E+00 | 6.00E-01 | 6.32E-01 | NP | 6.29E+04 | 6.26E+04 | 6.34E+04 | 6.41E+04 | 6.18E+04 |
| PE(22:1_20:4) | 9.68E+04 | 2.65E+05 | 3.64E-01 | 2.00E-02 | 1.93E-01 | 9.85E+04 | 3.53E+04 | 2.79E+00 | 5.71E-07 | 5.46E-05 | NP | 1.07E+05 | 1.08E+05 | 1.09E+05 | 1.05E+05 | 1.07E+05 |
| PE(18:1_24:5) | 4.31E+04 | 9.29E+04 | 4.64E-01 | 3.05E-02 | 1.93E-01 | 5.29E+04 | 7.45E+02 | 7.10E+01 | 5.41E-04 | 1.71E-03 | NP | 4.09E+04 | 4.16E+04 | 3.95E+04 | 4.58E+04 | 4.35E+04 |
| PE(25:0_18:1) | 2.96E+04 | 7.30E+03 | 4.06E+00 | 4.65E-02 | 2.17E-01 | 2.33E+04 | 2.52E+04 | 9.25E-01 | 5.97E-01 | 6.30E-01 | NP | 1.79E+04 | 1.77E+04 | 1.97E+04 | 1.72E+04 | 1.95E+04 |
| PE(18:1_25:1) | 2.70E+04 | 6.78E+03 | 3.99E+00 | 5.80E-02 | 2.38E-01 | 2.81E+04 | 1.40E+04 | 2.02E+00 | 7.62E-04 | 2.12E-03 | NP | 1.64E+04 | 1.41E+04 | 1.80E+04 | 1.55E+04 | 1.41E+04 |
| PE(26:0_18:1) | 1.34E+05 | 1.90E+04 | 7.02E+00 | 2.81E-02 | 1.93E-01 | 1.23E+05 | 1.24E+05 | 9.89E-01 | 9.46E-01 | 9.48E-01 | NP | 9.08E+04 | 8.49E+04 | 8.99E+04 | 9.79E+04 | 8.32E+04 |
| PE(18:1_26:1) | 1.35E+05 | 2.21E+04 | 6.09E+00 | 1.80E-02 | 1.93E-01 | 1.52E+05 | 5.59E+04 | 2.71E+00 | 8.98E-05 | 4.94E-04 | NP | 8.26E+04 | 8.08E+04 | 8.71E+04 | 7.94E+04 | 7.41E+04 |
| PE(24:0_20:4) | 1.13E+05 | 9.46E+04 | 1.19E+00 | 6.67E-01 | 8.25E-01 | 9.02E+04 | 1.32E+05 | 6.84E-01 | 6.66E-02 | 9.03E-02 | NP | 9.20E+04 | 9.34E+04 | 9.84E+04 | 9.55E+04 | 9.30E+04 |
| PE(24:1_20:4) | 5.95E+04 | 6.25E+04 | 9.52E-01 | 8.85E-01 | 9.43E-01 | 5.40E+04 | 3.09E+04 | 1.75E+00 | 7.74E-04 | 2.12E-03 | NP | 4.43E+04 | 4.03E+04 | 4.27E+04 | 4.63E+04 | 4.30E+04 |
| PE(24:0_20:5) | 2.52E+04 | 1.33E+04 | 1.90E+00 | 1.93E-01 | 3.91E-01 | 2.20E+04 | 7.03E+03 | 3.13E+00 | 2.46E-05 | 2.18E-04 | NP | 1.08E+04 | 1.22E+04 | 1.29E+04 | 1.36E+04 | 1.17E+04 |
| PE(26:0_20:4) | 1.95E+04 | 6.41E+03 | 3.04E+00 | 8.86E-02 | 2.78E-01 | 1.72E+04 | 2.34E+04 | 7.35E-01 | 9.48E-02 | 1.24E-01 | NP | 1.47E+04 | 1.85E+04 | 1.38E+04 | 1.41E+04 | 1.42E+04 |
| PE(24:0_22:5) | 3.20E+04 | 2.32E+04 | 1.38E+00 | 4.47E-01 | 6.58E-01 | 3.16E+04 | 4.25E+04 | 7.43E-01 | 8.18E-02 | 1.08E-01 | NP | 1.87E+04 | 1.66E+04 | 1.75E+04 | 1.69E+04 | 1.70E+04 |
| PE(O-16:1_20:5) | 3.72E+06 | 1.73E+06 | 2.14E+00 | 8.35E-02 | 2.68E-01 | 4.31E+06 | 1.81E+06 | 2.38E+00 | 7.78E-04 | 2.12E-03 | 2.25E+03 | 2.37E+06 | 2.55E+06 | 2.60E+06 | 2.54E+06 | 2.72E+06 |
| PE(O-18:2_20:5) | 1.35E+06 | 9.00E+05 | 1.50E+00 | 3.28E-01 | 5.51E-01 | 1.50E+06 | 3.15E+05 | 4.76E+00 | 2.30E-04 | 8.79E-04 | 1.28E-01 | 7.80E+05 | 8.73E+05 | 8.60E+05 | 9.18E+05 | 9.10E+05 |
| PE(O-18:1_22:6) | 9.22E+05 | 1.75E+06 | 5.27E-01 | 5.19E-02 | 2.36E-01 | 8.05E+05 | 3.24E+06 | 2.48E-01 | 1.79E-05 | 1.90E-04 | 1.40E+03 | 1.57E+06 | 1.47E+06 | 1.55E+06 | 1.60E+06 | 1.49E+06 |
| PE(O-18:2_22:6) | 7.59E+05 | 1.58E+06 | 4.81E-01 | 2.94E-02 | 1.93E-01 | 8.46E+05 | 6.84E+05 | 1.24E+00 | 2.71E-01 | 3.18E-01 | NP | 9.13E+05 | 8.87E+05 | 8.58E+05 | 8.62E+05 | 8.89E+05 |
| PG(15:0_18:1)+D7:(s) | 1.10E+06 | 1.11E+06 | 9.85E-01 | 9.54E-01 | 9.64E-01 | 8.87E+05 | 8.81E+05 | 1.01E+00 | 8.99E-01 | 9.11E-01 | 7.07E+05 | 8.96E+05 | 8.78E+05 | 8.75E+05 | 9.26E+05 | 9.70E+05 |
| PG(16:0_18:1) | 8.22E+05 | 6.80E+05 | 1.21E+00 | 4.94E-01 | 7.15E-01 | 8.49E+05 | 7.06E+05 | 1.20E+00 | 1.61E-01 | 1.95E-01 | 5.64E-02 | 6.65E+05 | 6.79E+05 | 6.91E+05 | 7.04E+05 | 7.45E+05 |
| PG(16:1_18:1) | 6.30E+04 | 3.43E+04 | 1.84E+00 | 4.52E-02 | 2.16E-01 | 6.65E+04 | 4.28E+04 | 1.56E+00 | 3.68E-02 | 5.42E-02 | NP | 4.77E+04 | 4.41E+04 | 5.23E+04 | 4.90E+04 | 5.28E+04 |
| PG(18:1_18:1) | 7.68E+04 | 7.02E+04 | 1.09E+00 | 7.68E-01 | 8.81E-01 | 7.16E+04 | 7.08E+04 | 1.01E+00 | 9.42E-01 | 9.47E-01 | NP | 5.35E+04 | 6.67E+04 | 6.76E+04 | 5.13E+04 | 6.83E+04 |
| PG(16:0_22:6) | 7.98E+04 | 4.06E+04 | 1.96E+00 | 3.55E-02 | 2.06E-01 | 1.14E+05 | 7.08E+04 | 1.61E+00 | 1.39E-01 | 1.70E-01 | NP | 7.01E+04 | 7.27E+04 | 7.42E+04 | 7.81E+04 | 8.16E+04 |
| PG(16:1_22:6) | 7.54E+04 | 2.90E+04 | 2.60E+00 | 3.61E-02 | 2.06E-01 | 1.13E+05 | 2.29E+04 | 4.92E+00 | 1.55E-03 | 3.85E-03 | NP | 5.54E+04 | 5.40E+04 | 5.71E+04 | 5.76E+04 | 5.92E+04 |
| PG(18:1_22:5) | 1.20E+05 | 5.75E+04 | 2.08E+00 | 3.60E-02 | 2.06E-01 | 1.55E+05 | 2.82E+04 | 5.52E+00 | 4.67E-03 | 9.64E-03 | 9.49E-02 | 7.20E+04 | 7.77E+04 | 9.02E+04 | 8.73E+04 | 8.20E+04 |
| PG(18:1_22:6) | 6.07E+05 | 3.92E+05 | 1.55E+00 | 1.45E-01 | 3.43E-01 | 6.86E+05 | 2.10E+05 | 3.26E+00 | 4.06E-02 | 5.91E-02 | NP | 4.42E+05 | 4.49E+05 | 4.44E+05 | 4.50E+05 | 4.87E+05 |
| PG(18:2_22:6) | 7.42E+04 | 6.10E+04 | 1.22E+00 | 5.91E-01 | 7.81E-01 | 9.17E+04 | 1.54E+04 | 5.96E+00 | 6.71E-04 | 1.91E-03 | NP | 5.23E+04 | 5.94E+04 | 5.82E+04 | 5.90E+04 | 5.96E+04 |
| PG(20:4_22:6) | 5.48E+04 | 3.12E+04 | 1.76E+00 | 2.19E-01 | 4.21E-01 | 7.76E+04 | 6.78E+03 | 1.14E+01 | 2.72E-04 | 9.82E-04 | NP | 3.60E+04 | 3.33E+04 | 3.64E+04 | 3.87E+04 | 4.20E+04 |
| PG(20:5_22:6) | 3.88E+04 | 1.37E+04 | 2.83E+00 | 1.21E-01 | 3.16E-01 | 6.10E+04 | 8.38E+02 | 7.28E+01 | 2.35E-04 | 8.79E-04 | NP | 2.70E+04 | 2.70E+04 | 2.32E+04 | 2.75E+04 | 2.64E+04 |
| PG(22:5_22:6) | 4.75E+05 | 1.12E+05 | 4.24E+00 | 4.16E-02 | 2.13E-01 | 5.26E+05 | 2.40E+04 | 2.19E+01 | 4.16E-05 | 2.65E-04 | NP | 2.40E+05 | 2.47E+05 | 2.62E+05 | 2.59E+05 | 2.66E+05 |
| PG(22:6_22:6) | 1.25E+06 | 5.17E+05 | 2.42E+00 | 9.28E-02 | 2.83E-01 | 1.40E+06 | 1.04E+05 | 1.34E+01 | 5.88E-05 | 3.45E-04 | NP | 6.87E+05 | 7.08E+05 | 7.46E+05 | 7.25E+05 | 7.56E+05 |

| | | | | | | | | | | | | | | | | |
|---|---|---|---|---|---|---|---|---|---|---|---|---|---|---|---|---|
| PI(14:0_16:0) | 8.12E+03 | 8.78E+03 | 9.25E-01 | 8.27E-01 | 9.21E-01 | 1.24E+04 | 3.27E+04 | 3.79E-01 | 1.30E-04 | 6.21E-04 | NP | 1.34E+04 | 1.34E+04 | 1.48E+04 | 1.51E+04 | 1.42E+04 |
| PI(16:0_16:0) | 1.16E+05 | 1.20E+05 | 9.73E-01 | 9.47E-01 | 9.62E-01 | 1.49E+05 | 3.65E+05 | 4.08E-01 | 3.16E-04 | 1.11E-03 | NP | 1.75E+05 | 1.82E+05 | 1.79E+05 | 1.75E+05 | 1.80E+05 |
| PI(16:0_16:1) | 2.38E+05 | 2.01E+05 | 1.18E+00 | 5.60E-01 | 7.57E-01 | 3.40E+05 | 3.81E+05 | 8.94E-01 | 1.38E-01 | 1.70E-01 | 7.49E+02 | 2.67E+05 | 2.89E+05 | 2.83E+05 | 2.84E+05 | 2.79E+05 |
| PI(16:1_16:1) | 1.31E+04 | 5.20E+03 | 2.51E+00 | 1.38E-01 | 3.36E-01 | 2.23E+04 | 3.15E+04 | 7.07E-01 | 8.07E-02 | 1.07E-01 | NP | 1.69E+04 | 1.07E+04 | 1.58E+04 | 1.73E+04 | 2.12E+04 |
| PI(16:0_17:1) | 3.48E+04 | 5.43E+04 | 6.40E-01 | 1.53E-01 | 3.53E-01 | 4.35E+04 | 8.72E+04 | 4.98E-01 | 6.78E-04 | 1.92E-03 | NP | 4.97E+04 | 5.04E+04 | 4.82E+04 | 4.32E+04 | 4.97E+04 |
| PI(15:0_18:1)+D7:(s) | 2.32E+05 | 2.44E+05 | 9.52E-01 | 8.53E-01 | 9.32E-01 | 1.85E+05 | 1.74E+05 | 1.06E+00 | 5.93E-01 | 6.29E-01 | 9.17E+04 | 1.93E+05 | 1.73E+05 | 1.96E+05 | 1.95E+05 | 1.95E+05 |
| PI(16:0_18:0) | 1.50E+05 | 1.51E+05 | 9.93E-01 | 9.85E-01 | 9.86E-01 | 2.25E+05 | 5.69E+05 | 3.96E-01 | 1.06E-03 | 2.75E-03 | 4.79E-03 | 2.31E+05 | 2.16E+05 | 2.35E+05 | 2.53E+05 | 2.23E+05 |
| PI(18:0_16:1) | 1.71E+06 | 1.80E+06 | 9.54E-01 | 8.47E-01 | 9.32E-01 | 2.27E+06 | 4.63E+06 | 4.89E-01 | 6.64E-04 | 1.91E-03 | 1.05E+04 | 2.42E+06 | 2.52E+06 | 2.47E+06 | 2.49E+06 | 2.59E+06 |
| PI(16:1_18:1)_Rt8.02 | 3.43E+05 | 2.58E+05 | 1.33E+00 | 2.59E-01 | 4.76E-01 | 5.44E+05 | 3.54E+05 | 1.54E+00 | 1.13E-02 | 2.07E-02 | 5.42E+02 | 3.39E+05 | 3.97E+05 | 3.39E+05 | 3.52E+05 | 3.55E+05 |
| PI(16:1_18:1)_Rt8.24 | 9.66E+04 | 1.19E+05 | 8.10E-01 | 5.07E-01 | 7.23E-01 | 1.45E+05 | 2.46E+05 | 5.92E-01 | 2.07E-02 | 3.43E-02 | 4.40E+02 | 1.26E+05 | 1.31E+05 | 1.30E+05 | 1.41E+05 | 1.30E+05 |
| PI(17:0_18:1) | 9.56E+04 | 1.05E+05 | 9.12E-01 | 7.52E-01 | 8.77E-01 | 1.10E+05 | 2.91E+05 | 3.77E-01 | 9.56E-05 | 5.05E-04 | NP | 1.28E+05 | 1.41E+05 | 1.32E+05 | 1.37E+05 | 1.47E+05 |
| PI(17:1_18:1) | 4.84E+04 | 5.44E+04 | 8.90E-01 | 7.18E-01 | 8.58E-01 | 7.13E+04 | 6.72E+04 | 1.06E+00 | 4.03E-01 | 4.49E-01 | 0.00E+00 | 5.85E+04 | 5.20E+04 | 5.22E+04 | 6.10E+04 | 5.92E+04 |
| PI(16:0_19:2) | 1.01E+04 | 2.00E+04 | 5.05E-01 | 7.71E-02 | 2.63E-01 | 1.09E+04 | 5.19E+04 | 2.11E-01 | 2.66E-05 | 2.23E-04 | 0.00E+00 | 2.03E+04 | 1.84E+04 | 2.00E+04 | 2.07E+04 | 1.92E+04 |
| PI(18:0_18:0) | 2.23E+04 | 1.43E+04 | 1.56E+00 | 6.69E-01 | 8.25E-01 | 1.04E+03 | 6.82E+04 | 1.52E-02 | 2.06E-05 | 2.07E-04 | NP | 4.21E+04 | 1.02E+01 | 4.61E+04 | 4.44E+04 | 4.56E+04 |
| PI(18:0_18:1) | 2.15E+06 | 1.59E+06 | 1.36E+00 | 3.85E-01 | 6.03E-01 | 2.73E+06 | 7.02E+06 | 3.88E-01 | 4.88E-05 | 3.01E-04 | 7.91E+03 | 2.91E+06 | 3.29E+06 | 3.08E+06 | 2.89E+06 | 2.97E+06 |
| PI(18:1_18:1) | 7.69E+05 | 7.80E+05 | 9.86E-01 | 9.51E-01 | 9.63E-01 | 1.10E+06 | 1.72E+06 | 6.37E-01 | 1.15E-02 | 2.08E-02 | 1.37E+03 | 8.62E+05 | 1.09E+06 | 9.83E+05 | 1.04E+06 | 1.07E+06 |
| PI(18:0_18:2) | 3.32E+05 | 5.17E+05 | 6.41E-01 | 1.96E-01 | 3.92E-01 | 4.53E+05 | 8.09E+05 | 5.60E-01 | 7.05E-04 | 1.98E-03 | 6.28E+00 | 4.10E+05 | 4.05E+05 | 5.93E+05 | 4.37E+05 | 4.69E+05 |
| PI(18:1_18:2) | 8.67E+04 | 1.33E+05 | 6.52E-01 | 2.55E-01 | 4.73E-01 | 1.30E+05 | 9.53E+04 | 1.36E+00 | 2.63E-02 | 4.17E-02 | NP | 8.64E+04 | 1.19E+05 | 1.01E+05 | 1.04E+05 | 9.67E+04 |
| PI(16:0_20:3) | 1.64E+05 | 2.37E+05 | 6.89E-01 | 3.16E-01 | 5.42E-01 | 1.70E+05 | 3.89E+05 | 4.37E-01 | 5.25E-04 | 1.68E-03 | NP | 2.15E+05 | 2.09E+05 | 2.27E+05 | 2.15E+05 | 2.23E+05 |
| PI(16:0_20:4) | 5.26E+05 | 9.48E+05 | 5.55E-01 | 1.55E-01 | 3.53E-01 | 5.18E+05 | 5.56E+05 | 9.33E-01 | 3.04E-01 | 3.49E-01 | 3.46E+02 | 5.55E+05 | 5.73E+05 | 5.34E+05 | 5.92E+05 | 5.78E+05 |
| PI(16:1_20:4) | 1.66E+04 | 2.81E+04 | 5.91E-01 | 1.18E-01 | 3.16E-01 | 2.34E+04 | 8.68E+03 | 2.70E+00 | 8.12E-06 | 1.41E-04 | NP | 1.62E+04 | 1.69E+04 | 1.46E+04 | 1.36E+04 | 1.59E+04 |
| PI(16:0_20:5) | 4.57E+04 | 8.07E+04 | 5.66E-01 | 2.15E-01 | 4.14E-01 | 5.60E+04 | 4.51E+04 | 1.24E+00 | 1.33E-02 | 2.36E-02 | NP | 5.60E+04 | 5.16E+04 | 4.63E+04 | 5.14E+04 | 5.11E+04 |
| PI(18:0_19:2) | 2.08E+04 | 5.65E+04 | 3.69E-01 | 6.00E-02 | 2.38E-01 | 2.12E+04 | 1.41E+05 | 1.50E-01 | 1.30E-04 | 6.21E-04 | NP | 6.66E+04 | 5.84E+04 | 5.49E+04 | 5.99E+04 | 6.15E+04 |
| PI(17:0_20:3) | 1.97E+04 | 4.23E+04 | 4.65E-01 | 7.01E-02 | 2.48E-01 | 1.85E+04 | 4.77E+04 | 3.88E-01 | 2.34E-05 | 2.13E-04 | NP | 2.78E+04 | 2.55E+04 | 2.86E+04 | 2.74E+04 | 3.02E+04 |
| PI(18:0_19:3) | 2.77E+04 | 4.08E+04 | 6.78E-01 | 2.51E-01 | 4.68E-01 | 2.24E+04 | 5.84E+04 | 3.84E-01 | 6.14E-04 | 1.83E-03 | NP | 2.78E+04 | 3.44E+04 | 3.27E+04 | 4.17E+04 | 3.54E+04 |
| PI(17:0_20:4) | 6.95E+04 | 2.18E+05 | 3.19E-01 | 2.91E-02 | 1.93E-01 | 6.50E+04 | 6.66E+04 | 9.76E-01 | 8.92E-01 | 9.06E-01 | NP | 7.57E+04 | 8.79E+04 | 9.56E+04 | 9.42E+04 | 8.73E+04 |
| PI(17:1_20:4) | 9.19E+03 | 2.69E+04 | 3.42E-01 | 4.23E-02 | 2.13E-01 | 1.13E+04 | 5.09E+03 | 2.21E+00 | 6.24E-04 | 1.83E-03 | NP | 9.59E+03 | 1.06E+04 | 1.05E+04 | 1.10E+04 | 9.56E+03 |
| PI(18:0_20:1) | 5.19E+04 | 9.59E+04 | 5.42E-01 | 1.69E-01 | 3.69E-01 | 5.49E+04 | 2.04E+05 | 2.70E-01 | 1.92E-04 | 8.04E-04 | NP | 1.26E+05 | 1.38E+05 | 1.42E+05 | 1.17E+05 | 9.18E+04 |
| PI(18:0_20:2) | 1.60E+05 | 3.49E+05 | 4.58E-01 | 2.62E-02 | 1.93E-01 | 1.52E+05 | 1.80E+06 | 8.45E-02 | 2.12E-05 | 2.07E-04 | 7.59E+02 | 5.60E+05 | 5.27E+05 | 2.85E+05 | 5.89E+05 | 6.00E+05 |
| PI(18:0_20:3)_Rt9.76 | 7.36E+05 | 1.23E+06 | 5.99E-01 | 1.85E-01 | 3.87E-01 | 7.06E+05 | 1.00E+06 | 7.03E-01 | 2.08E-02 | 3.43E-02 | 1.53E+03 | 7.51E+05 | 7.95E+05 | 7.55E+05 | 8.25E+05 | 8.43E+05 |
| PI(18:0_20:3)_Rt10.1 | 2.83E+05 | 5.18E+05 | 5.47E-01 | 9.67E-02 | 2.88E-01 | 2.31E+05 | 8.20E+05 | 2.82E-01 | 4.62E-04 | 1.54E-03 | 5.79E+02 | 4.49E+05 | 4.31E+05 | 4.11E+05 | 4.29E+05 | 4.32E+05 |

| | | | | | | | | | | | | | | | | |
|---|---|---|---|---|---|---|---|---|---|---|---|---|---|---|---|---|
| PI(18:0_20:4) | 5.45E+06 | 1.22E+07 | 4.45E-01 | 5.91E-02 | 2.38E-01 | 5.01E+06 | 4.46E+06 | 1.12E+00 | 2.92E-01 | 3.39E-01 | 6.25E+03 | 5.48E+06 | 6.01E+06 | 6.08E+06 | 6.01E+06 | 6.87E+06 |
| PI(18:1_20:4) | 7.78E+05 | 1.49E+06 | 5.23E-01 | 9.80E-02 | 2.89E-01 | 8.68E+05 | 5.30E+05 | 1.64E+00 | 2.80E-04 | 1.00E-03 | NP | 7.79E+05 | 8.20E+05 | 8.08E+05 | 8.34E+05 | 8.61E+05 |
| PI(18:0_20:5) | 2.65E+05 | 4.84E+05 | 5.47E-01 | 1.62E-01 | 3.55E-01 | 3.36E+05 | 2.79E+05 | 1.20E+00 | 6.81E-02 | 9.20E-02 | NP | 3.00E+05 | 2.82E+05 | 3.10E+05 | 3.16E+05 | 3.08E+05 |
| PI(18:1_20:5) | 6.40E+04 | 1.16E+05 | 5.51E-01 | 1.78E-01 | 3.78E-01 | 8.27E+04 | 3.72E+04 | 2.23E+00 | 1.76E-04 | 7.73E-04 | NP | 6.58E+04 | 5.94E+04 | 5.93E+04 | 6.41E+04 | 6.51E+04 |
| PI(16:0_22:6) | 9.96E+04 | 1.91E+05 | 5.23E-01 | 1.12E-01 | 3.15E-01 | 1.19E+05 | 1.25E+05 | 9.47E-01 | 4.61E-01 | 5.04E-01 | NP | 1.14E+05 | 1.18E+05 | 1.18E+05 | 1.14E+05 | 1.14E+05 |
| PI(18:0_21:2) | 2.98E+03 | 7.88E+03 | 3.78E-01 | 6.31E-02 | 2.39E-01 | 1.77E+03 | 3.91E+04 | 4.52E-02 | 6.67E-06 | 1.41E-04 | NP | 7.81E+03 | 9.79E+03 | 1.05E+04 | 1.36E+04 | 1.02E+04 |
| PI(18:0_21:3) | 7.66E+04 | 1.18E+05 | 6.49E-01 | 3.62E-01 | 5.82E-01 | 7.05E+04 | 1.33E+05 | 5.28E-01 | 2.32E-04 | 8.79E-04 | NP | 8.21E+04 | 7.52E+04 | 9.35E+04 | 8.17E+04 | 8.60E+04 |
| PI(19:0_20:4) | 3.77E+04 | 1.21E+05 | 3.10E-01 | 1.32E-02 | 1.93E-01 | 3.45E+04 | 1.81E+04 | 1.90E+00 | 2.10E-03 | 4.92E-03 | NP | 3.92E+04 | 4.97E+04 | 4.16E+04 | 4.79E+04 | 4.80E+04 |
| PI(19:1_20:4) | 3.96E+04 | 7.17E+04 | 5.52E-01 | 1.09E-01 | 3.10E-01 | 3.83E+04 | 4.95E+04 | 7.74E-01 | 2.26E-03 | 5.24E-03 | 4.39E+03 | 4.66E+04 | 4.50E+04 | 4.93E+04 | 4.55E+04 | 4.34E+04 |
| PI(17:0_22:6) | 7.33E+03 | 2.35E+04 | 3.12E-01 | 1.62E-02 | 1.93E-01 | 7.80E+03 | 1.03E+04 | 7.58E-01 | 3.23E-01 | 3.66E-01 | NP | 1.11E+04 | 1.07E+04 | 1.13E+04 | 8.90E+03 | 1.06E+04 |
| PI(18:0_22:2) | 9.40E+03 | 2.38E+04 | 3.96E-01 | 3.06E-02 | 1.93E-01 | 6.25E+03 | 6.31E+04 | 9.89E-02 | 1.38E-06 | 5.86E-05 | 6.60E-03 | 2.10E+04 | 2.45E+04 | 2.63E+04 | 2.56E+04 | 2.96E+04 |
| PI(18:0_22:3) | 1.80E+04 | 3.61E+04 | 4.97E-01 | 7.97E-02 | 2.66E-01 | 1.47E+04 | 4.79E+04 | 3.07E-01 | 4.62E-04 | 1.54E-03 | NP | 2.45E+04 | 2.44E+04 | 1.86E+04 | 3.17E+04 | 1.20E+04 |
| PI(18:0_22:4) | 2.26E+05 | 3.80E+05 | 5.96E-01 | 2.11E-01 | 4.11E-01 | 2.41E+05 | 1.80E+05 | 1.33E+00 | 3.67E-02 | 5.42E-02 | 8.91E+00 | 2.16E+05 | 2.35E+05 | 2.00E+05 | 2.32E+05 | 2.28E+05 |
| PI(20:0_20:4) | 2.84E+04 | 1.12E+05 | 2.54E-01 | 2.66E-02 | 1.93E-01 | 2.66E+04 | 2.95E+04 | 9.00E-01 | 4.51E-01 | 4.95E-01 | 4.22E+02 | 4.38E+04 | 4.13E+04 | 4.45E+04 | 4.40E+04 | 4.70E+04 |
| PI(18:1_22:4) | 2.47E+04 | 4.79E+04 | 5.17E-01 | 9.84E-02 | 2.89E-01 | 3.60E+04 | 1.39E+04 | 2.59E+00 | 3.19E-05 | 2.32E-04 | NP | 2.43E+04 | 2.63E+04 | 2.74E+04 | 2.72E+04 | 2.56E+04 |
| PI(18:0_22:5)_Rt9.30 | 5.92E+05 | 6.96E+05 | 8.52E-01 | 6.15E-01 | 7.94E-01 | 5.89E+05 | 5.15E+05 | 1.14E+00 | 2.57E-01 | 3.02E-01 | NP | 5.08E+05 | 5.40E+05 | 5.42E+05 | 5.49E+05 | 5.90E+05 |
| PI(18:0_22:5)_Rt9.75 | 2.37E+04 | 4.20E+04 | 5.65E-01 | 1.53E-01 | 3.53E-01 | 1.81E+04 | 3.54E+04 | 5.11E-01 | 6.18E-03 | 1.25E-02 | NP | 2.64E+04 | 2.67E+04 | 2.00E+04 | 3.42E+04 | 3.14E+04 |
| PI(18:1_22:5) | 9.63E+04 | 9.86E+04 | 9.76E-01 | 9.42E-01 | 9.60E-01 | 1.26E+05 | 4.86E+04 | 2.60E+00 | 9.05E-05 | 4.94E-04 | NP | 7.65E+04 | 7.87E+04 | 7.78E+04 | 8.03E+04 | 8.24E+04 |
| PI(18:0_22:6) | 3.34E+05 | 7.21E+05 | 4.63E-01 | 6.59E-02 | 2.40E-01 | 3.62E+05 | 4.64E+05 | 7.81E-01 | 3.29E-02 | 4.99E-02 | NP | 3.50E+05 | 4.29E+05 | 4.54E+05 | 4.05E+05 | 4.14E+05 |
| PI(18:1_22:6) | 5.50E+04 | 1.10E+05 | 5.00E-01 | 6.21E-02 | 2.39E-01 | 7.13E+04 | 4.05E+04 | 1.76E+00 | 1.81E-04 | 7.85E-04 | NP | 5.84E+04 | 5.67E+04 | 5.78E+04 | 6.30E+04 | 6.47E+04 |
| PI(O-16:0_16:0) | 9.60E+03 | 1.47E+04 | 6.54E-01 | 3.64E-01 | 5.82E-01 | 1.01E+04 | 7.14E+04 | 1.42E-01 | 2.37E-06 | 7.54E-05 | NP | 2.43E+04 | 2.53E+04 | 2.75E+04 | 2.72E+04 | 2.51E+04 |
| PI(O-16:0_20:4) | 5.19E+04 | 1.29E+04 | 4.03E+00 | 2.09E-02 | 1.93E-01 | 4.55E+04 | 7.37E+04 | 6.17E-01 | 2.09E-02 | 3.43E-02 | NP | 4.09E+04 | 2.61E+04 | 3.13E+04 | 4.18E+04 | 4.46E+04 |
| PI(O-16:0_22:5) | 4.20E+04 | 5.34E+04 | 7.87E-01 | 3.37E-01 | 5.59E-01 | 3.46E+04 | 2.09E+05 | 1.66E-01 | 2.09E-05 | 2.07E-04 | NP | 7.87E+04 | 8.01E+04 | 7.83E+04 | 8.20E+04 | 7.42E+04 |
| PI(O-16:0_22:6) | 2.13E+04 | 5.67E+04 | 3.77E-01 | 1.87E-02 | 1.93E-01 | 1.77E+04 | 1.97E+05 | 9.01E-02 | 9.36E-06 | 1.55E-04 | NP | 6.64E+04 | 6.93E+04 | 6.65E+04 | 7.33E+04 | 7.02E+04 |
| PI(O-18:0_22:5) | 1.11E+04 | 2.06E+04 | 5.36E-01 | 1.31E-03 | 7.17E-02 | 1.88E+04 | 7.13E+04 | 2.64E-01 | 1.56E-02 | 2.69E-02 | NP | 1.66E+04 | 1.61E+04 | 1.48E+04 | 1.42E+04 | 1.52E+04 |
| PI(O-18:1_22:5) | 4.12E+03 | 1.72E+04 | 2.40E-01 | 1.73E-02 | 1.93E-01 | 4.99E+03 | 2.58E+04 | 1.93E-01 | 2.01E-04 | 8.26E-04 | NP | 1.19E+04 | 1.07E+04 | 1.17E+04 | 1.20E+04 | 1.13E+04 |
| PI(O-18:0_22:6) | 5.16E-03 | 7.95E+03 | 6.49E-07 | 1.25E-02 | 1.91E-01 | 1.55E-02 | 7.93E+04 | 1.95E-07 | 5.52E-05 | 3.29E-04 | NP | 1.58E+04 | 1.61E+04 | 1.69E+04 | 1.74E+04 | 1.58E+04 |
| PIP(P-22:6_18:1) | 7.64E+04 | 5.08E+04 | 1.50E+00 | 1.02E-01 | 2.94E-01 | 9.74E+04 | 1.53E+05 | 6.35E-01 | 2.75E-03 | 6.26E-03 | NP | 8.21E+04 | 9.78E+04 | 9.36E+04 | 9.58E+04 | 9.77E+04 |
| PIP(P-24:7_18:1) | 4.02E+04 | 3.53E+04 | 1.14E+00 | 5.22E-01 | 7.26E-01 | 5.46E+04 | 7.22E+04 | 7.56E-01 | 8.78E-02 | 1.16E-01 | NP | 4.94E+04 | 5.33E+04 | 4.81E+04 | 4.85E+04 | 5.10E+04 |
| PS(16:0_16:0) | 5.22E+04 | 2.26E+04 | 2.31E+00 | 3.68E-02 | 2.07E-01 | 6.32E+04 | 1.48E+04 | 4.28E+00 | 2.29E-05 | 2.13E-04 | 0.00E+00 | 3.36E+04 | 3.37E+04 | 3.34E+04 | 3.74E+04 | 3.47E+04 |
| PS(16:0_16:1) | 2.18E+05 | 1.07E+05 | 2.03E+00 | 2.83E-02 | 1.93E-01 | 2.58E+05 | 1.80E+05 | 1.43E+00 | 2.18E-01 | 2.59E-01 | 2.88E+03 | 1.79E+05 | 1.77E+05 | 1.90E+05 | 1.86E+05 | 1.89E+05 |
| PS(16:0_17:1) | 6.85E+04 | 4.86E+04 | 1.41E+00 | 2.93E-01 | 5.15E-01 | 7.54E+04 | 3.80E+04 | 1.98E+00 | 1.55E-03 | 3.85E-03 | 3.68E+02 | 5.21E+04 | 5.02E+04 | 5.87E+04 | 5.94E+04 | 5.68E+04 |

| | | | | | | | | | | | | | | | | |
|---|---|---|---|---|---|---|---|---|---|---|---|---|---|---|---|---|
| PS(15:0_18:1)+D7:(s) | 8.68E+04 | 9.01E+04 | 9.63E-01 | 8.86E-01 | 9.43E-01 | 6.87E+04 | 7.24E+04 | 9.50E-01 | 3.46E-01 | 3.90E-01 | 5.58E+04 | 7.32E+04 | 6.89E+04 | 6.91E+04 | 7.90E+04 | 6.82E+04 |
| PS(16:0_18:1) | 1.96E+06 | 1.18E+06 | 1.66E+00 | 9.03E-02 | 2.81E-01 | 2.44E+06 | 1.53E+06 | 1.60E+00 | 1.36E-02 | 2.40E-02 | 2.41E+04 | 1.63E+06 | 1.72E+06 | 1.59E+06 | 1.62E+06 | 1.81E+06 |
| PS(16:1_18:1)_Rt8.27 | 1.22E+05 | 5.74E+04 | 2.12E+00 | 6.63E-02 | 2.40E-01 | 1.55E+05 | 9.98E+04 | 1.55E+00 | 1.21E-01 | 1.54E-01 | 1.36E+03 | 8.76E+04 | 8.93E+04 | 9.37E+04 | 9.19E+04 | 8.95E+04 |
| PS(16:1_18:1)_Rt8.48 | 7.35E+04 | 2.83E+04 | 2.60E+00 | 2.61E-02 | 1.93E-01 | 9.14E+04 | 5.34E+04 | 1.71E+00 | 1.04E-02 | 1.94E-02 | 9.03E+02 | 6.35E+04 | 6.15E+04 | 5.88E+04 | 6.07E+04 | 6.11E+04 |
| PS(18:0_17:1) | 3.93E+05 | 3.76E+05 | 1.05E+00 | 9.25E-01 | 9.59E-01 | 4.69E+05 | 5.00E+05 | 9.38E-01 | 4.24E-01 | 4.67E-01 | 3.83E+03 | 4.03E+05 | 4.15E+05 | 4.48E+05 | 4.00E+05 | 4.93E+05 |
| PS(18:0_18:1) | 6.28E+06 | 5.58E+06 | 1.13E+00 | 6.62E-01 | 8.25E-01 | 7.45E+06 | 9.91E+06 | 7.53E-01 | 1.75E-03 | 4.24E-03 | 1.24E+05 | 6.66E+06 | 6.78E+06 | 7.00E+06 | 7.03E+06 | 6.26E+06 |
| PS(18:1_18:1) | 8.11E+05 | 4.47E+05 | 1.81E+00 | 8.54E-02 | 2.72E-01 | 1.05E+06 | 6.20E+05 | 1.70E+00 | 3.75E-03 | 8.26E-03 | 1.22E+04 | 6.27E+05 | 6.82E+05 | 6.86E+05 | 6.61E+05 | 7.19E+05 |
| PS(18:0_18:2) | 4.83E+05 | 3.31E+05 | 1.46E+00 | 2.38E-01 | 4.49E-01 | 5.98E+05 | 5.22E+05 | 1.15E+00 | 6.48E-02 | 8.88E-02 | 1.05E+04 | 4.11E+05 | 4.54E+05 | 4.25E+05 | 4.55E+05 | 4.48E+05 |
| PS(18:1_18:2) | 1.14E+05 | 6.55E+04 | 1.73E+00 | 1.53E-01 | 3.53E-01 | 1.45E+05 | 8.24E+04 | 1.76E+00 | 1.70E-03 | 4.15E-03 | 0.00E+00 | 9.23E+04 | 8.64E+04 | 9.51E+04 | 9.50E+04 | 9.43E+04 |
| PS(18:0_18:3) | 1.96E+04 | 1.53E+04 | 1.28E+00 | 5.22E-01 | 7.26E-01 | 2.55E+04 | 5.17E+04 | 4.93E-01 | 9.66E-05 | 5.05E-04 | 3.45E-02 | 2.67E+04 | 2.68E+04 | 2.88E+04 | 2.88E+04 | 2.47E+04 |
| PS(16:0_20:4)_Rt8.36 | 4.11E+04 | 2.30E+04 | 1.79E+00 | 1.59E-01 | 3.53E-01 | 5.64E+04 | 2.44E+04 | 2.31E+00 | 2.51E-05 | 2.18E-04 | 0.00E+00 | 3.12E+04 | 3.30E+04 | 3.34E+04 | 3.27E+04 | 3.16E+04 |
| PS(16:0_20:4)_Rt9.47 | 1.72E+05 | 1.13E+05 | 1.51E+00 | 3.24E-02 | 1.96E-01 | 2.47E+05 | 1.51E+05 | 1.63E+00 | 7.52E-03 | 1.48E-02 | 8.43E-02 | 1.56E+05 | 1.75E+05 | 1.64E+05 | 1.55E+05 | 1.63E+05 |
| PS(17:0_20:3) | 2.87E+04 | 3.13E+04 | 9.17E-01 | 7.72E-01 | 8.83E-01 | 3.01E+04 | 3.62E+04 | 8.30E-01 | 1.14E-01 | 1.46E-01 | NP | 2.90E+04 | 2.95E+04 | 3.37E+04 | 2.94E+04 | 3.19E+04 |
| PS(20:0_18:1) | 5.05E+05 | 4.92E+05 | 1.03E+00 | 9.24E-01 | 9.59E-01 | 4.49E+05 | 6.49E+05 | 6.91E-01 | 2.11E-03 | 4.92E-03 | 6.90E+03 | 4.80E+05 | 4.83E+05 | 4.73E+05 | 4.87E+05 | 5.15E+05 |
| PS(18:1_20:1) | 3.95E+05 | 3.51E+05 | 1.12E+00 | 6.46E-01 | 8.17E-01 | 3.98E+05 | 3.78E+05 | 1.05E+00 | 5.88E-01 | 6.28E-01 | 4.09E+03 | 3.33E+05 | 3.32E+05 | 3.31E+05 | 3.47E+05 | 3.23E+05 |
| PS(18:0_20:3) | 1.18E+06 | 1.37E+06 | 8.61E-01 | 5.14E-01 | 7.24E-01 | 1.37E+06 | 1.58E+06 | 8.66E-01 | 1.61E-01 | 1.96E-01 | 1.14E+04 | 1.42E+06 | 1.38E+06 | 1.26E+06 | 1.38E+06 | 1.28E+06 |
| PS(18:1_20:3) | 7.15E+04 | 4.37E+04 | 1.63E+00 | 4.14E-01 | 6.30E-01 | 7.92E+04 | 3.95E+04 | 2.00E+00 | 3.42E-02 | 5.14E-02 | 0.00E+00 | 5.18E+04 | 4.06E+04 | 5.23E+04 | 6.57E+04 | 5.58E+04 |
| PS(18:0_20:4)_Rt9.55 | 3.09E+05 | 3.17E+05 | 9.72E-01 | 9.18E-01 | 9.59E-01 | 4.62E+05 | 3.10E+05 | 1.49E+00 | 9.92E-05 | 5.09E-04 | 8.23E-01 | 3.08E+05 | 3.31E+05 | 3.16E+05 | 3.08E+05 | 3.38E+05 |
| PS(18:0_20:4)_Rt10.7 | 6.69E+05 | 5.78E+05 | 1.16E+00 | 4.89E-01 | 7.11E-01 | 7.78E+05 | 9.67E+05 | 8.04E-01 | 2.49E-04 | 9.13E-04 | 5.41E-01 | 6.83E+05 | 7.09E+05 | 7.71E+05 | 7.61E+05 | 7.05E+05 |
| PS(18:0_20:5) | 3.28E+04 | 1.67E+04 | 1.97E+00 | 3.21E-01 | 5.48E-01 | 1.03E+05 | 7.10E+03 | 1.45E+01 | 7.18E-06 | 1.41E-04 | 0.00E+00 | 2.69E+04 | 3.14E+04 | 2.49E+04 | 2.10E+04 | 2.80E+04 |
| PS(18:1_20:4) | 9.22E+04 | 4.91E+04 | 1.88E+00 | 2.21E-02 | 1.93E-01 | 1.23E+05 | 6.56E+04 | 1.88E+00 | 1.18E-03 | 3.03E-03 | 8.14E+00 | 7.89E+04 | 8.18E+04 | 8.41E+04 | 7.71E+04 | 8.46E+04 |
| PS(16:0_22:6) | 3.21E+04 | 7.55E+04 | 4.25E-01 | 3.19E-02 | 1.96E-01 | 4.51E+04 | 4.96E+04 | 9.10E-01 | 1.41E-01 | 1.73E-01 | NP | 4.52E+04 | 4.47E+04 | 4.42E+04 | 4.64E+04 | 4.44E+04 |
| PS(17:0_22:5) | 1.38E+04 | 3.77E+04 | 3.66E-01 | 4.19E-02 | 2.13E-01 | 1.03E+04 | 4.05E+04 | 2.53E-01 | 1.67E-04 | 7.52E-04 | 9.70E+00 | 2.13E+04 | 2.69E+04 | 2.39E+04 | 2.62E+04 | 3.47E+04 |
| PS(22:0_18:1) | 5.25E+05 | 4.16E+05 | 1.26E+00 | 4.98E-01 | 7.18E-01 | 4.23E+05 | 5.29E+05 | 8.00E-01 | 3.25E-02 | 4.94E-02 | 8.56E+03 | 4.11E+05 | 4.26E+05 | 4.31E+05 | 4.41E+05 | 4.25E+05 |
| PS(18:1_22:1) | 4.16E+05 | 4.18E+05 | 9.95E-01 | 9.86E-01 | 9.86E-01 | 3.85E+05 | 2.66E+05 | 1.45E+00 | 1.06E-03 | 2.75E-03 | 4.08E+03 | 3.36E+05 | 3.29E+05 | 3.36E+05 | 3.28E+05 | 3.45E+05 |
| PS(18:0_22:4) | 1.87E+05 | 5.82E+05 | 3.21E-01 | 2.39E-02 | 1.93E-01 | 2.73E+05 | 4.69E+05 | 5.82E-01 | 2.86E-03 | 6.44E-03 | 3.97E+03 | 3.63E+05 | 3.37E+05 | 3.73E+05 | 3.36E+05 | 3.25E+05 |
| PS(18:0_22:5) | 2.21E+05 | 1.09E+06 | 2.03E-01 | 1.12E-02 | 1.91E-01 | 2.18E+05 | 1.82E+06 | 1.20E-01 | 1.14E-05 | 1.55E-04 | 7.74E+03 | 7.28E+05 | 7.73E+05 | 7.07E+05 | 7.72E+05 | 7.97E+05 |
| PS(18:1_22:5) | 5.14E+04 | 1.04E+05 | 4.94E-01 | 4.45E-02 | 2.16E-01 | 6.98E+04 | 1.14E+05 | 6.12E-01 | 2.98E-04 | 1.05E-03 | 1.37E-01 | 5.49E+04 | 7.91E+04 | 6.91E+04 | 6.32E+04 | 6.68E+04 |
| PS(18:0_22:6) | 3.01E+05 | 1.56E+06 | 1.92E-01 | 3.17E-03 | 1.16E-01 | 2.93E+05 | 2.48E+06 | 1.18E-01 | 3.53E-05 | 2.49E-04 | 1.01E+04 | 1.02E+06 | 1.01E+06 | 9.99E+05 | 1.01E+06 | 1.06E+06 |

| | | | | | | | | | | | | | | | | |
|---|---|---|---|---|---|---|---|---|---|---|---|---|---|---|---|---|
| PS(18:1_22:6) | 3.55E+04 | 8.99E+04 | 3.95E-01 | 3.02E-02 | 1.93E-01 | 5.12E+04 | 6.80E+04 | 7.53E-01 | 3.94E-02 | 5.77E-02 | NP | 5.29E+04 | 5.58E+04 | 5.55E+04 | 5.63E+04 | 5.74E+04 |
| PS(20:3_20:4) | 3.82E+04 | 3.29E+04 | 1.16E+00 | 5.93E-01 | 7.81E-01 | 4.99E+04 | 3.23E+04 | 1.54E+00 | 1.74E-03 | 4.23E-03 | NP | 3.66E+04 | 3.45E+04 | 3.43E+04 | 4.03E+04 | 3.97E+04 |
| PS(24:0_18:1) | 3.27E+05 | 1.48E+05 | 2.21E+00 | 9.58E-02 | 2.88E-01 | 2.29E+05 | 3.97E+05 | 5.76E-01 | 5.33E-02 | 7.51E-02 | 4.81E+03 | 3.00E+05 | 3.10E+05 | 2.34E+05 | 2.98E+05 | 2.19E+05 |
| PS(18:1_24:1) | 2.57E+05 | 1.44E+05 | 1.78E+00 | 1.60E-01 | 3.53E-01 | 2.24E+05 | 1.53E+05 | 1.47E+00 | 8.01E-03 | 1.56E-02 | 1.80E+03 | 1.59E+05 | 1.78E+05 | 1.64E+05 | 1.73E+05 | 1.71E+05 |
| PS(20:3_22:6)_Rt9.25 | 3.07E+04 | 1.37E+05 | 2.24E-01 | 2.39E-04 | 2.29E-02 | 2.67E+04 | 6.58E+02 | 4.06E+01 | 1.33E-03 | 3.33E-03 | 6.53E+00 | 1.04E+05 | 1.12E+05 | 1.09E+05 | 1.19E+05 | 9.87E+04 |
| PS(20:3_22:6)_Rt9.58 | 9.59E+04 | 2.04E+05 | 4.71E-01 | 1.21E-02 | 1.91E-01 | 1.20E+05 | 2.73E+05 | 4.41E-01 | 8.96E-05 | 4.94E-04 | 1.03E+03 | 1.16E+05 | 1.31E+05 | 1.22E+05 | 1.18E+05 | 1.21E+05 |
| PS(18:1_26:3) | 1.39E+05 | 7.48E+04 | 1.86E+00 | 1.20E-01 | 3.16E-01 | 1.27E+05 | 8.05E+04 | 1.58E+00 | 3.99E-03 | 8.61E-03 | 7.90E+02 | 9.32E+04 | 8.96E+04 | 9.44E+04 | 9.47E+04 | 8.94E+04 |
| SM(d34:0) | 1.37E+06 | 1.22E+05 | 1.13E+01 | 7.00E-03 | 1.57E-01 | 1.18E+06 | 1.06E+06 | 1.11E+00 | 5.28E-01 | 5.73E-01 | NP | 9.19E+05 | 8.57E+05 | 8.94E+05 | 8.60E+05 | 9.13E+05 |
| SM(d18:1_17:0) | 6.04E+04 | 1.56E+05 | 3.88E-01 | 2.90E-02 | 1.93E-01 | 7.00E+04 | 1.77E+05 | 3.95E-01 | 1.61E-05 | 1.84E-04 | NP | 1.02E+05 | 8.44E+04 | 1.06E+05 | 1.10E+05 | 1.04E+05 |
| SM(d18:1_22:0) | 4.14E+05 | 6.40E+05 | 6.47E-01 | 2.14E-01 | 4.14E-01 | 5.48E+05 | 1.28E+06 | 4.28E-01 | 2.53E-04 | 9.22E-04 | NP | 5.40E+05 | 6.60E+05 | 6.45E+05 | 6.56E+05 | 6.37E+05 |
| SM(d41:1) | 1.81E+05 | 2.29E+05 | 7.91E-01 | 5.31E-01 | 7.32E-01 | 1.89E+05 | 4.90E+05 | 3.86E-01 | 1.12E-05 | 1.55E-04 | NP | 1.95E+05 | 2.36E+05 | 2.67E+05 | 2.57E+05 | 2.66E+05 |
| SM(d42:1) | 1.62E+06 | 1.87E+06 | 8.70E-01 | 7.21E-01 | 8.58E-01 | 2.12E+06 | 3.37E+06 | 6.29E-01 | 9.64E-04 | 2.56E-03 | NP | 1.98E+06 | 2.07E+06 | 2.04E+06 | 2.07E+06 | 2.04E+06 |
| SM(d43:1) | 4.98E+04 | 3.90E+04 | 1.28E+00 | 6.15E-01 | 7.94E-01 | 6.69E+04 | 1.16E+05 | 5.79E-01 | 5.60E-04 | 1.71E-03 | NP | 6.28E+04 | 6.33E+04 | 5.84E+04 | 6.89E+04 | 6.03E+04 |
| SM(d44:2) | 1.86E+05 | 4.11E+04 | 4.54E+00 | 5.41E-02 | 2.38E-01 | 2.53E+05 | 1.60E+05 | 1.59E+00 | 6.66E-02 | 9.03E-02 | NP | 1.14E+05 | 1.09E+05 | 1.86E+05 | 1.72E+05 | 1.22E+05 |
